## Supplementary fig. for "Unravelling the developmental and functional significance of an ancient Argonaute duplication"

**Content:**

**Supplementary Figure. 1:** miRNA expression in distinct developmental stages and the *Nematostella* AGOs expression profile.

**Supplementary Figure. 2:** miRNAs exhibiting alternative strand selection in NveAGO1 and NveAGO2.

**Supplementary Figure 3:** miRNA strand selection is affected by the knockdown of their carrying AGOs.

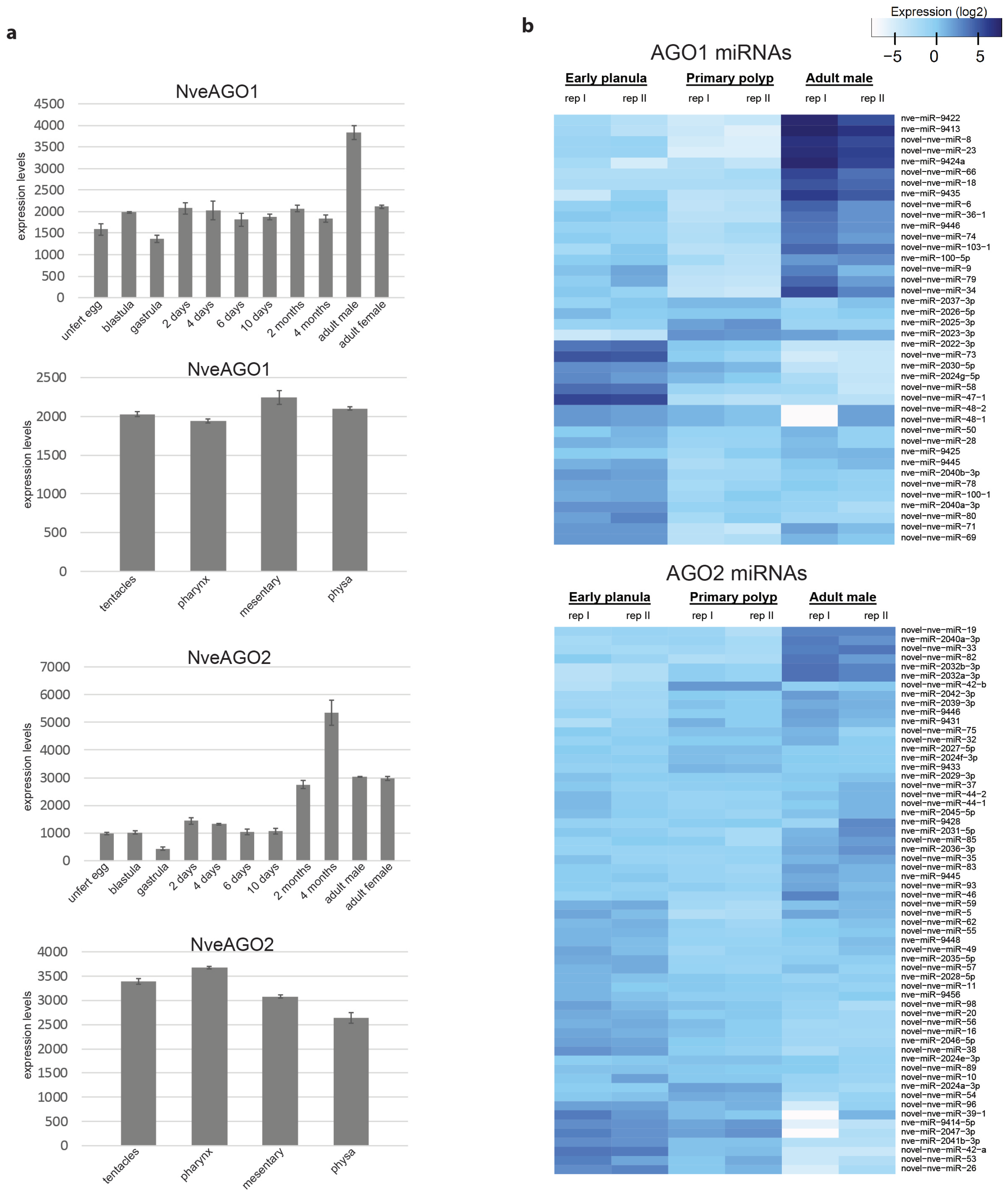


**Supplementary Fig. 1. miRNA expression in distinct developmental stages and the *Nematostella* AGOs expression profile.**

**a**, Expression of *Nematostella* AGOs throughout development and in distinct tissues. Nanostring data to generate this representation (**Supplementary table 5**) was taken from Praher et al[1]. **b**, Heatmap representing $log_2$ expression levels of known and novel miRNAs. Counts per Million (CPM) from two distinct biological replicates of each NveAGO-IP were used to generate this heatmap.

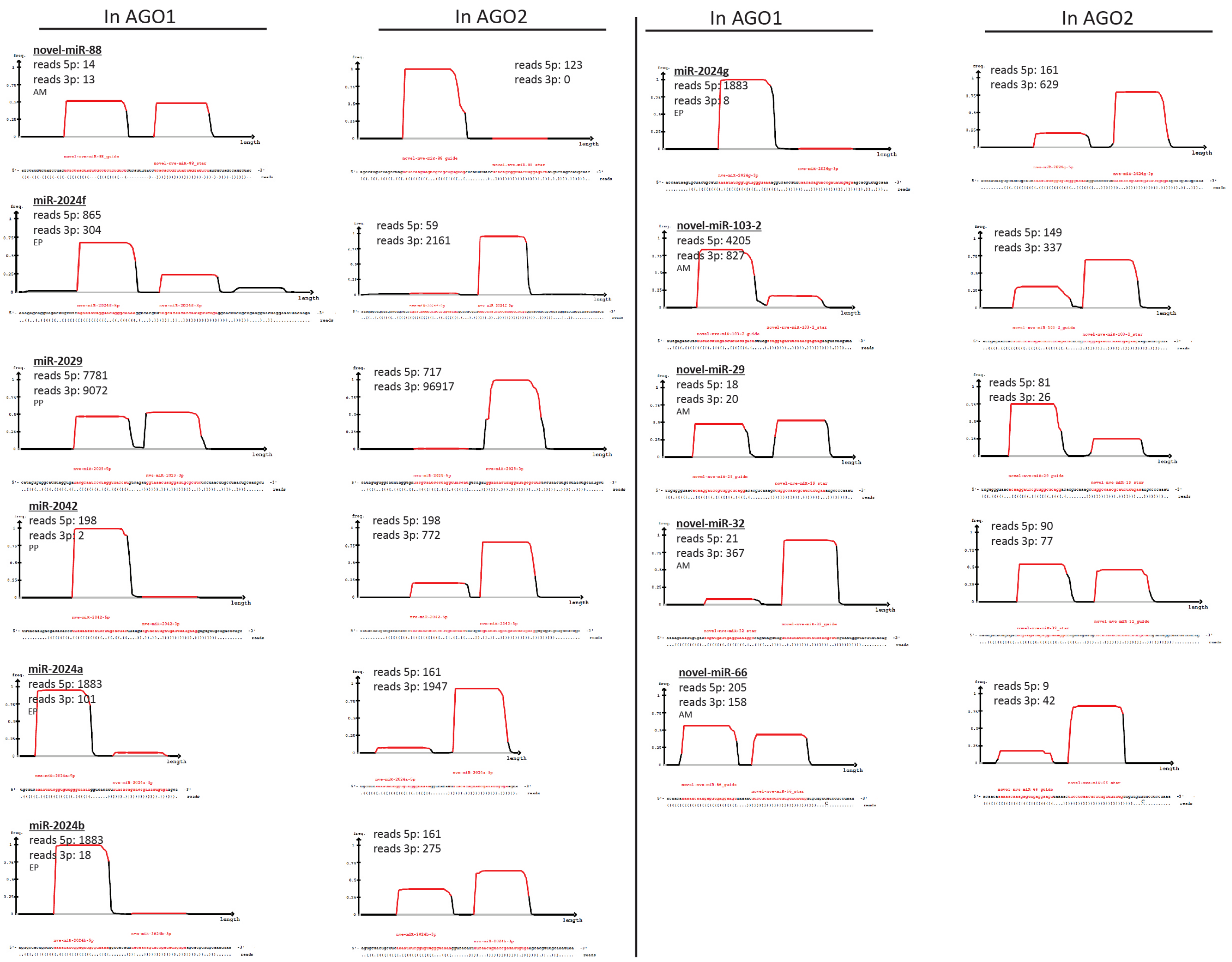


**Supplementary Fig. 2. miRNAs exhibiting alternative strand selection in NveAGO1 and NveAGO2.**

Eleven miRNAs exhibit alternative profiles of strand selection depending on their hosting AGO. Each pair represents miRNA signature as observed in NveAGO1 IP (left columns) or in NveAGO2 IP (right columns). The graphic outputs were generated using mirDeep2[2]. Abbreviations: AM, Adult male; EP, Early planula; PP, Primary polyp.

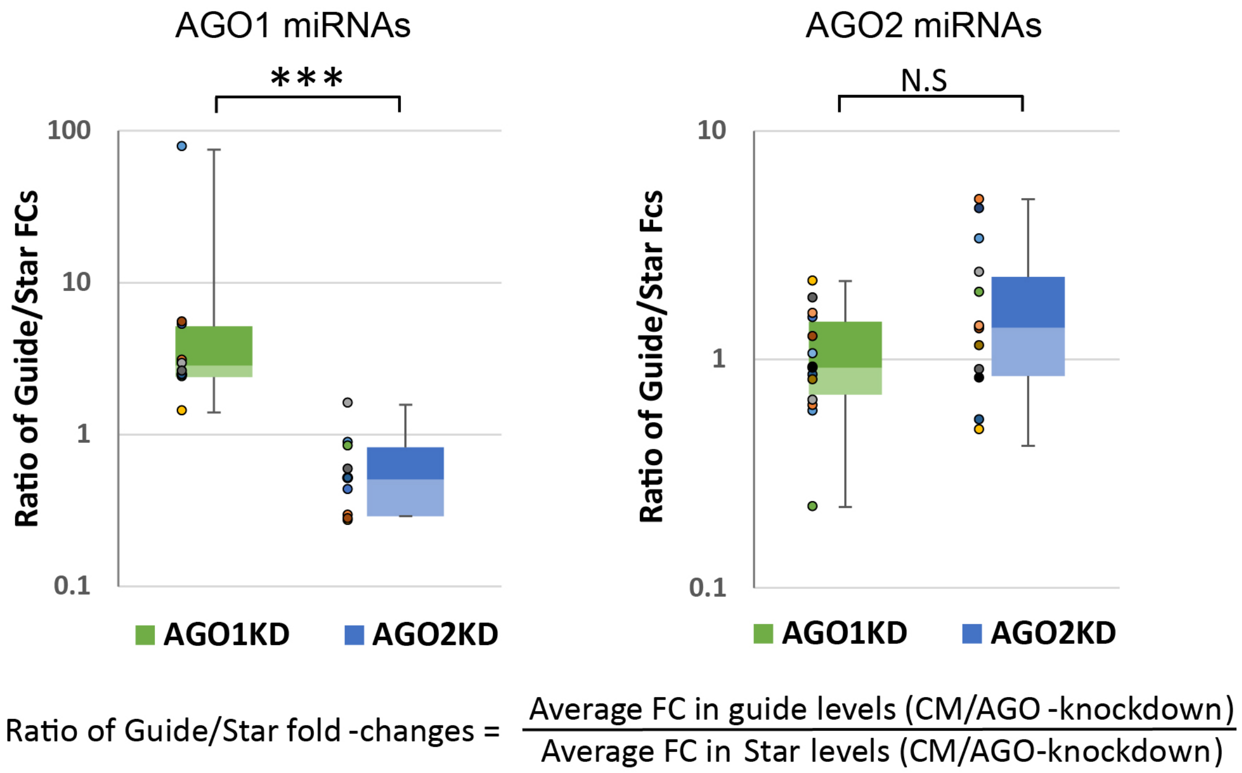


**Supplementary Figure 3. AGO knockdown effect on *Nematostella* miRNA strand selection.**

Strand selection disruption of NveAGO1 miRNAs in NveAGO1 knockdown (left panel) and NveAGO2 miRNAs in NveAGO2 knockdown (right panel). The reads mapped to individual miRNAs were normalized using spike-ins and average reads were generated from biological triplicates (**Supplementary table 5**). Weakly expressed miRNAs (less than 50 read counts for an individual miRNA) and miRNA with no star reads were excluded from this analysis. Next we calculated the ratio between the average guide fold-change to the average star fold-change (lower panel). NveAGO1 enriched miRNAs are significantly affected in NveAGO1 KD (P=0.00029, Mann-Whitney U Test). NveAGO2 enriched miRNAs did not exhibit a significant change in NveAGO2 KD (P=0.15, Mann-Whitney U Test).

**References**


1 Praher, D. *et al.* Characterization of the piRNA pathway during development of the sea anemone Nematostella vectensis. *RNA Biol* **14**, 1727-1741, doi:10.1080/15476286.2017.1349048 (2017).

2 Friedlander, M. R., Mackowiak, S. D., Li, N., Chen, W. & Rajewsky, N. miRDeep2 accurately identifies known and hundreds of novel microRNA genes in seven animal clades. *Nucleic acids research* **40**, 37-52, doi:10.1093/nar/gkr688 (2012).
