## Supplementary file 2 for "Unravelling the developmental and functional significance of an ancient Argonaute duplication"

### Novel *Nematostella* miRNAs

miRBase precursor : novel-nve-miR-5
Total read count : 208
novel-nve-miR-5_guide read count : 203
novel-nve-miR-5_star read count : 5
remaining reads : 0

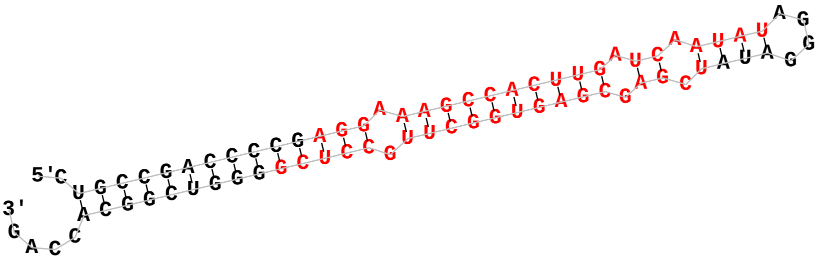


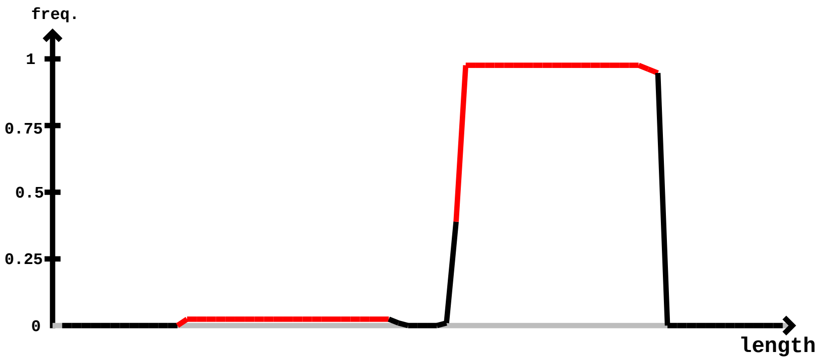


novel-nve-miR-5_star
novel-nve-miR-5_guide

```
5'- cugccgaccccgaggaaagccacuugaucaauauagggauaucgagcgaguggcuugccucggggucggcaccag   -3'    exp
    .((((((((((((((.((((((((((.((.((((....)))).)).)))))))))).))))))))))))))....   reads      mm         sample
    ............aggaaagccacuugaucaauau.........................................   1          0          seq
    ............aggaaagccacCugaucaauau.........................................   1          1          seq
    ............aggaaagccacuugaucaauaC.........................................   1          1          seq
    ............aggaaagccacuugaucaauauU........................................   2          1          seq
    .......................................uaucgagcgaguggcuugUcu...............   1          1          seq
    .......................................uaucgagcgaUuggcuugccucg.............   1          1          seq
    ........................................aucgagcgaguggcuugccu...............   2          0          seq
    ........................................aucgagcgaguggcuugccuc..............   1          0          seq
    ........................................aucgagcgaUuggcuugccucg.............   3          1          seq
    ........................................aucgagcgaguggcuugccucU.............   56         1          seq
    ........................................aucgagcgaguggcuugccucg.............   5          0          seq
    ........................................aucgagcgaguggcuugccucA.............   5          1          seq
    ........................................aucgagcgaguggcuugccucC.............   7          1          seq
    .........................................ucgagcgaguggcuugccuc..............   2          0          seq
    .........................................ucgagcgaguggcuugccucU.............   88         1          seq
    .........................................ucgagcgaguggcuugccucC.............   16         1          seq
    .........................................ucgagcgaguggcuugccucg.............   3          0          seq
    .........................................ucgagcgaguggcuugccucA.............   13         1          seq
```

miRBase precursor : novel-nve-miR-6
Total read count : 1792
novel-nve-miR-6_guide read count : 1432
novel-nve-miR-6_star read count : 213
remaining reads : 147

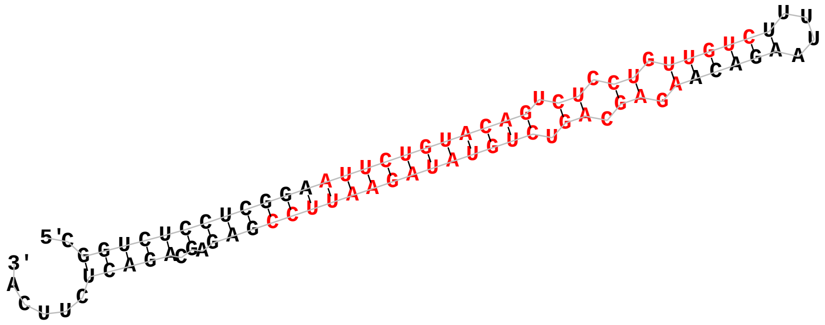

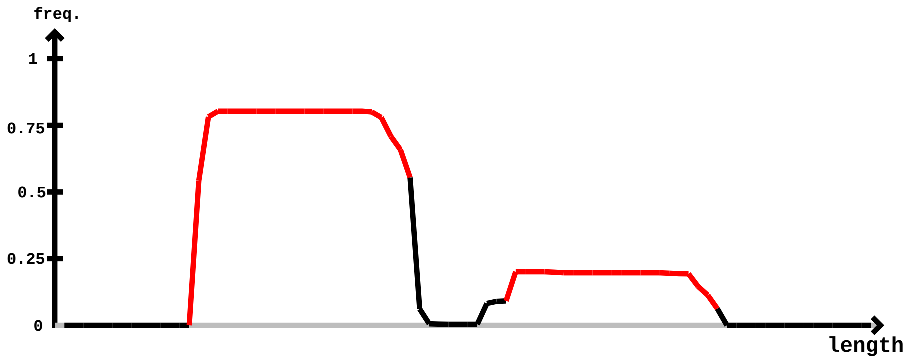


novel-nve-miR-6_guide

novel-nve-miR-6_star

| | | | |
|---|---|---|---|
| 5'- cggucuccucggaauucuguacagucuccuguugucuuuuaagacaagagcagucuguauagaauuccgagagcagacucuuca -3' | exp | | |
| .((((((((((((((((((((((.((.((.((((((....)))))).)).)).)))))))))))))))).).))))))..... | reads | mm | sample |
| .............auucuguacagucuccug..................................................... | 3 | 0 | seq |
| .............auucuguacagucuccuA..................................................... | 1 | 1 | seq |
| .............auucuguacGgucuccug..................................................... | 1 | 1 | seq |
| .............auucuguacagucuccugu.................................................... | 29 | 0 | seq |
| .............auucuguacagucuccugC.................................................... | 1 | 1 | seq |
| .............Uuucuguacagucuccugu.................................................... | 1 | 1 | seq |
| .............auucuguaUagucuccugu.................................................... | 2 | 1 | seq |
| .............auucuguGcagucuccugu.................................................... | 1 | 1 | seq |
| .............Guucuguacagucuccuguu................................................... | 1 | 1 | seq |
| .............auucuguacagucuccuguC................................................... | 11 | 1 | seq |
| .............auucuguacagucuccuguG................................................... | 2 | 1 | seq |
| .............auucuguacagucuccuguu................................................... | 93 | 0 | seq |
| .............auucCguacagucuccuguu................................................... | 2 | 1 | seq |
| .............auucugAacagucuccuguu................................................... | 1 | 1 | seq |
| .............auucuguGcagucuccuguu................................................... | 1 | 1 | seq |
| .............auucuguacagucuccugCu................................................... | 1 | 1 | seq |
| .............auucuguacagucuccuguA................................................... | 3 | 1 | seq |
| .............auucuguacUgucuccuguug.................................................. | 1 | 1 | seq |
| .............auucCguacagucuccuguug.................................................. | 1 | 1 | seq |
| .............auucuguacagucuccuguuA.................................................. | 4 | 1 | seq |
| .............auucuguacagucuccuguuC.................................................. | 4 | 1 | seq |
| .............auucuguacagucuccuguug.................................................. | 51 | 0 | seq |
| .............auucuguacagucuccuCuug.................................................. | 1 | 1 | seq |
| .............auucuguacagucuccCguug.................................................. | 1 | 1 | seq |
| .............auucuguacagucuccuguAg.................................................. | 1 | 1 | seq |
| .............auucuguacagucuccuguuU.................................................. | 10 | 1 | seq |
| .............Cuucuguacagucuccuguug.................................................. | 1 | 1 | seq |
| .............auucuguacagucuccuguugC................................................. | 18 | 1 | seq |
| .............auucuguacagucuccuguugu................................................. | 102 | 0 | seq |
| .............auucuguacagucucUuguugu................................................. | 2 | 1 | seq |
| .............auucuguacagucuccuguugA................................................. | 3 | 1 | seq |
| .............auucuguGcagucuccuguugu................................................. | 1 | 1 | seq |
| .............auucuguacagucuccuguuUu................................................. | 4 | 1 | seq |
| .............auucAguacagucuccuguugu................................................. | 1 | 1 | seq |

```
             novel-nve-miR-6_guide
                                                novel-nve-miR-6_star
cggucuccucggaauucuguacagucuccuguugucuuuuaagacaagagcagucuguauagaauuccgagagcagacucuuca

.............auucuguacagucuccuguugG.................................................    4      1      seq
.............auucuguacagucuccuguCgu.................................................    1      1      seq
.............auucuguacGgucuccuguugu.................................................    1      1      seq
.............auucuguacUgucuccuguuguc................................................    1      1      seq
.............aAucuguacagucuccuguuguc................................................    1      1      seq
.............auucuguacagucuccAguuguc................................................    1      1      seq
.............auucuguacagAcuccuguuguc................................................    1      1      seq
.............auucuguacagucuccuguugCc................................................    2      1      seq
.............auuGuguacagucuccuguuguc................................................    1      1      seq
.............auucuguGcagucuccuguuguc................................................    3      1      seq
.............auucugCacagucuccuguuguc................................................    1      1      seq
.............auucuguUcagucuccuguuguc................................................    1      1      seq
.............auucuguaUagucuccuguuguc................................................    2      1      seq
.............auucuguacagucuccuguuUuc................................................    1      1      seq
.............auucuguacagucuccugCuguc................................................    1      1      seq
.............auucuguacagucCccuguuguc................................................    2      1      seq
.............Cuucuguacagucuccuguuguc................................................    1      1      seq
.............auucuguacGgucuccuguuguc................................................    7      1      seq
.............auucuguacagucuccuguuguU................................................    55     1      seq
.............auucuguacagucuccuguCguc................................................    1      1      seq
.............auucuguacagucuAcuguuguc................................................    1      1      seq
.............Guucuguacagucuccuguuguc................................................    6      1      seq
.............auucuguacaguUuccuguuguc................................................    1      1      seq
.............auucuguacagCcuccuguuguc................................................    3      1      seq
.............auucuguacagucuccuguuguc................................................    451    0      seq
.............auucuguacagucuccuguuguA................................................    21     1      seq
.............auCcuguacagucuccuguuguc................................................    2      1      seq
.............auucuguacagucuccuguAguc................................................    2      1      seq
.............Uuucuguacagucuccuguuguc................................................    1      1      seq
.............auucuguacagucuccuCuuguc................................................    2      1      seq
.............auucuguacagucuUcuguuguc................................................    1      1      seq
.............auuAuguacagucuccuguuguc................................................    1      1      seq
.............auucuguacagucucUuguuguc................................................    3      1      seq
.............auucuguacagucuccGguuguc................................................    1      1      seq
.............auAcuguacagucuccuguuguc................................................    1      1      seq
.............auucuguacagucuccuguuguG................................................    1      1      seq
.............auucuguacagucucGuguugucu...............................................    1      1      seq
.............auucuguacagucuccuguugCcu...............................................    1      1      seq
.............auucuguacagucuccuguugucu...............................................    12     0      seq
.............auucuguacagucuccuguugucC...............................................    4      1      seq
.............auucuguacagucuccuguugucG...............................................    1      1      seq
.............auucuguacagucuccuguugucA...............................................    1      1      seq
.............auuUuguacagucuccuguugucu...............................................    2      1      seq
.............auucuguacagucuccuguugucuu..............................................    2      0      seq
.............auucuAuacagucuccuguugucuuu.............................................    1      1      seq
.............auucuguacagucuccuguugucuuuuaagacaagagc.................................    1      0      seq
..............uucuguacagucuccugu....................................................    3      0      seq
..............uucuguacagucuccuguu...................................................    11     0      seq
..............uucuguacagucuccuguA...................................................    1      1      seq
..............uucuguacagucuccuguuA..................................................    1      1      seq
..............uucuguacagucuccuguug..................................................    9      0      seq
..............uucuguacagucuccuguuU..................................................    5      1      seq
..............uucuguacagCcuccuguugu.................................................    1      1      seq
..............uucuguaAagucuccuguugu.................................................    1      1      seq
..............Gucuguacagucuccuguugu.................................................    1      1      seq
..............uucuguacagucuccugCugu.................................................    2      1      seq
..............uucuguacagucuccuguugC.................................................    5      1      seq
..............uucuguacagucuccuguCgu.................................................    2      1      seq
..............uucuguacagucuccuguugu.................................................    37     0      seq
..............uucuguacagucuccuguuUuc................................................    1      1      seq
..............uucCguacagucuccuguuguc................................................    2      1      seq
..............uucuguacagucuccuguuguc................................................    235    0      seq
..............Gucuguacagucuccuguuguc................................................    1      1      seq
..............Aucuguacagucuccuguuguc................................................    2      1      seq
..............uucuguacagucCccuguuguc................................................    1      1      seq
..............uucuguacagucuccuguuguA................................................    7      1      seq
..............uucuguacagucucUuguuguc................................................    5      1      seq
..............uucuguacGgucuccuguuguc................................................    2      1      seq
..............uucugCacagucuccuguuguc................................................    1      1      seq
```

novel-nve-miR-6_guide

novel-nve-miR-6_star

```
cggucuccucggaauucuguacagucuccuguugucuuuuaagacaagagcagucuguauagaauuccgagagcagacucuuca

..............uucuCuacagucuccuguuguc................................................	1	1	seq
..............uucuguacagucuccuguugAc................................................	1	1	seq
..............uucuguacagucuccuguuguU................................................	28	1	seq
..............uucuguacagucuccugCuguc................................................	1	1	seq
..............uAcuguacagucuccuguuguc................................................	1	1	seq
..............uucuguacagucuccAguuguc................................................	1	1	seq
..............uucuguacagucAccuguuguc................................................	1	1	seq
..............uCcuguacagucuccuguuguc................................................	1	1	seq
..............uucuguacagucuAcuguuguc................................................	2	1	seq
..............Cucuguacagucuccuguuguc................................................	1	1	seq
..............uucuguacagCcuccuguugucu...............................................	1	1	seq
..............uucuguacagucuccuguugucA...............................................	5	1	seq
..............uucuguacagucuccuguuguGu...............................................	1	1	seq
..............uucuguacagucuccuguugucC...............................................	7	1	seq
..............Aucuguacagucuccuguugucu...............................................	1	1	seq
..............uucuguacagucuccuguugucu...............................................	33	0	seq
..............uucuguacGgucuccuguugucu...............................................	1	1	seq
..............uucuguacagucuccuguugucuuuuaagacaagag..................................	2	0	seq
..............uucuguacagucuccCguugucuuuuaagacaagag..................................	1	1	seq
..............uucuguacagucuccuguugucuuuuaagacaagagc.................................	2	0	seq
..............uucuguacagucuccuguugucuuuuaagacaagagA.................................	1	1	seq
...............ucuguacagucuccuguugC.................................................	1	1	seq
...............ucuguacagucuccuguuguU................................................	2	1	seq
...............ucuguacagucuccuguugGc................................................	1	1	seq
...............ucuguaUagucuccuguuguc................................................	2	1	seq
...............ucuguacagucuccuguuguc................................................	3	0	seq
...............ucuguacagucuccuguugucA...............................................	4	1	seq
...............ucuguacagucuccuguugucu...............................................	19	0	seq
...............ucugCacagucuccuguugucu...............................................	1	1	seq
...............ucuguacagucuccCguugucu...............................................	1	1	seq
...............ucuguacagucuccuguugucC...............................................	1	1	seq
...............ucuguacagucuccuguugucG...............................................	1	1	seq
...............ucuguacGgucuccuguugucu...............................................	1	1	seq
...............ucGguacagucuccuguugucu...............................................	1	1	seq
...........................................acaagagcagucuguauaA......................	1	1	seq
...........................................acaagagcagucuguauag......................	2	0	seq
...........................................acaagagcagucuguauaga.....................	3	0	seq
...........................................acaagagcagucuguauagaa....................	1	0	seq
...........................................Ucaagagcagucuguauagaau...................	1	1	seq
...........................................acGagagcagucuguauagaau...................	1	1	seq
...........................................acaagagcagucuguauagaGu...................	1	1	seq
...........................................acaaUagcagucuguauagaau...................	1	1	seq
...........................................acaagagcagucuguauagaaC...................	15	1	seq
...........................................acaagagcagCcuguauagaau...................	1	1	seq
...........................................acaagagUagucuguauagaau...................	1	1	seq
...........................................acaagagcagucuguauagaau...................	61	0	seq
...........................................acaagagcagucuguauagaaCu..................	5	1	seq
...........................................acaagagcaguAuguauagaauu..................	1	1	seq
...........................................acaagagcagucuguauagaauA..................	2	1	seq
...........................................acaagagcagucuguauagaauu..................	37	0	seq
...........................................Gcaagagcagucuguauagaauu..................	1	1	seq
...........................................acaagagcagucuguauagaaAu..................	1	1	seq
...........................................acaagagcagucuguauagaauC..................	3	1	seq
...........................................acaagagcagucuguauagaauuU.................	1	1	seq
............................................caagagcagucuguauagaau...................	2	0	seq
............................................caagagcagucuguauagaauC..................	1	1	seq
............................................caagagcagucuguauagaauuU.................	2	1	seq
............................................caagagcagucuguauagaauuc.................	9	0	seq
.............................................aagagcagucuguauagaauuU.................	1	1	seq
.............................................aagagcagucuguauagaauuc.................	2	0	seq
..............................................agagcagucuguauagaau...................	1	0	seq
..............................................agagcagucuguauagaauC..................	1	1	seq
..............................................agagcagucuguauagaauu..................	6	0	seq
..............................................agagcagucuguauGgaauuc.................	1	1	seq
..............................................agagcagucuguauagaGuuc.................	1	1	seq
..............................................agagcagucuguUuagaauuc.................	1	1	seq
..............................................agagcagucuguauagGauuc.................	1	1	seq
..............................................agagcagucuguaCagaauuc.................	1	1	seq
..............................................agGgcagucuguauagaauuc.................	1	1	seq
```

```
             novel-nve-miR-6_guide
                                                novel-nve-miR-6_star
cggucuccucggaauucuguacagucuccuguugucuuuuaagacaagagcagucuguauagaauuccgagagcagacucuuca

.............................................agagcagucuguauagaauuc.................   70    0    seq
.............................................agagcagucuguauagaauucU................   95    1    seq
.............................................agagcagucuguauagaauucA................   6     1    seq
.............................................agagcagucuguauagaauuAc................   1     1    seq
.............................................agagcagucuguauagaauucc................   10    0    seq
```

miRBase precursor : novel-nve-miR-8
Total read count : 34694
novel-nve-miR-8_guide read count : 34269
novel-nve-miR-8_star read count : 425
remaining reads : 0

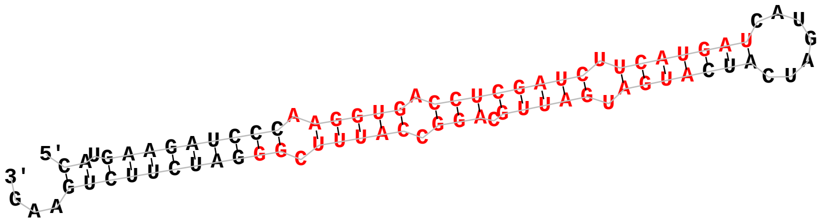

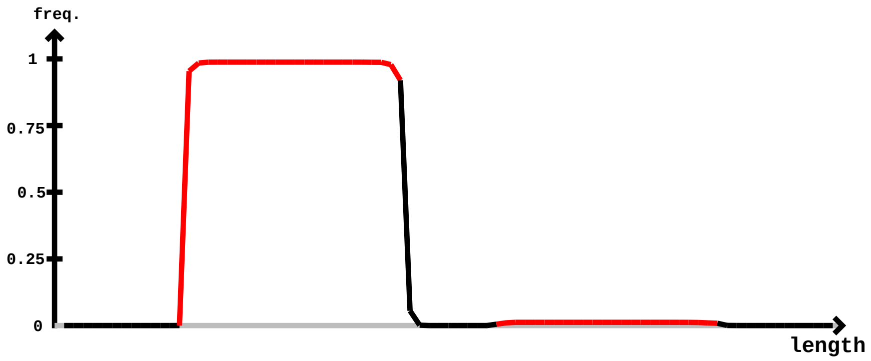


```
                                                  novel-nve-miR-8_star
                 novel-nve-miR-8_guide
5'- caugaagauccaaggugaccucgaucuucaugaucaugaucaucaugaugauugcaggccauuucgggaucuucugaag   -3'    exp
    ((.(((((((((.(((((.((((((((.(((((((.......))))))).))))).))).))))).)))))))))))...    reads      mm         sample
    ...........caaggugaccucgaucuucaugau.............................................    5          0          seq
    ...........caaggugaccucgaucuCcaugau.............................................    1          1          seq
    ............aaggugaccucgaucuuc..................................................    1          0          seq
    ............aaggugaccucgaucuuca.................................................    13         0          seq
    ............aaggugaccucgGucuuca.................................................    1          1          seq
    ............aaggugaccucgaucuucau................................................    6          0          seq
    ............Gaggugaccucgaucuucau................................................    1          1          seq
    ............aaggugaccucgaucuucGug...............................................    2          1          seq
    ............aaggugaccucgaucuucaCg...............................................    2          1          seq
    ............aaggugaccucgaucCucaug...............................................    1          1          seq
    ............aaggAgaccucgaucuucaug...............................................    3          1          seq
    ............aaggugaccuUgaucuucaug...............................................    2          1          seq
    ............aaggugaccucgaucAucaug...............................................    1          1          seq
    ............aaggugaccucgaucuucauA...............................................    23         1          seq
    ............aaggugaccCcgaucuucaug...............................................    1          1          seq
    ............aaggugaccucgauAuucaug...............................................    1          1          seq
    ............aagguCaccucgaucuucaug...............................................    1          1          seq
    ............aaggugacAucgaucuucaug...............................................    1          1          seq
    ............aaggugaUcucgaucuucaug...............................................    1          1          seq
    ............aaggugaccucgaucuCcaug...............................................    2          1          seq
    ............aaggCgaccucgaucuucaug...............................................    1          1          seq
    ............aaggugaccucgaGcuucaug...............................................    1          1          seq
    ............aaggugaccucgaucuucauU...............................................    6          1          seq
    ............aaggugaccucgaucuucaug...............................................    232        0          seq
    ............aaggugaccucgaucuuUaug...............................................    1          1          seq
    ............aaggugaccucgaucuucaGg...............................................    1          1          seq
    ............Gaggugaccucgaucuucaug...............................................    1          1          seq
    ............aaggugaccucgaucuucaugC..............................................    2          1          seq
    ............aaggugaccucgaucuucaAga..............................................    1          1          seq
    ............aaggugaccucAaucuucauga..............................................    1          1          seq
    ............aagguCaccucgaucuucauga..............................................    2          1          seq
    ............aaggugaccucgaucuuUauga..............................................    6          1          seq
    ............aagAugaccucgaucuucauga..............................................    1          1          seq
    ............aagguAaccucgaucuucauga..............................................    4          1          seq
```

```
                                                    novel-nve-miR-8_star
            novel-nve-miR-8_guide
caugaagauccc aaggugaccucgaucuucaugau caugaucauc augaugauugcaggccauuucgg gaucuucugaag
```

```
caugaagaucccaaggugaccucgaucuucaugaucaugaucaucaugaugauugcaggccauuucgggaucuucugaag

............aaggugaccucgaucuucaGga..............................................    1       1       seq
............aaggugaccucgaucuucaugU..............................................    23      1       seq
............Gaggugaccucgaucuucauga..............................................    16      1       seq
............aUggugaccucgaucuucauga..............................................    1       1       seq
............aaggugacGucgaucuucauga..............................................    2       1       seq
............aaggugGccucgaucuucauga..............................................    16      1       seq
............aaggugaccucgaucuGcauga..............................................    1       1       seq
............aaggugaccucgaucGucauga..............................................    1       1       seq
............aaggugaccucgaucCucauga..............................................    7       1       seq
............aaggGgaccucgaucuucauga..............................................    2       1       seq
............aaggugUccucgaucuucauga..............................................    2       1       seq
............aaggugaccucgaucuucGuga..............................................    4       1       seq
............aGggugaccucgaucuucauga..............................................    8       1       seq
............aaggugaccucgauUuucauga..............................................    3       1       seq
............aaggugaccucgaucuucaugG..............................................    21      1       seq
............aaggugaccucgaGcuucauga..............................................    1       1       seq
............aaggugaccuUgaucuucauga..............................................    2       1       seq
............aaggCgaccucgaucuucauga..............................................    5       1       seq
............Caggugaccucgaucuucauga..............................................    1       1       seq
............aaggugaccucgauAuucauga..............................................    1       1       seq
............aaggugaccAcgaucuucauga..............................................    1       1       seq
............aaAgugaccucgaucuucauga..............................................    3       1       seq
............aaggugaccucgaucuucUuga..............................................    2       1       seq
............aaggugaccucgaucuucCuga..............................................    1       1       seq
............aaggugaUcucgaucuucauga..............................................    2       1       seq
............aaggugaccucUaucuucauga..............................................    1       1       seq
............Uaggugaccucgaucuucauga..............................................    1       1       seq
............aaggugaccucgaucuCcauga..............................................    10      1       seq
............aaggugaccucgGucuucauga..............................................    12      1       seq
............aagCugaccucgaucuucauga..............................................    2       1       seq
............aaggugaccucgaAcuucauga..............................................    1       1       seq
............aaggugaccCcgaucuucauga..............................................    7       1       seq
............aaggugacUucgaucuucauga..............................................    1       1       seq
............aaggugaccucgaucuucauAa..............................................    3       1       seq
............aaggugaccucgaucuuGauga..............................................    2       1       seq
............aaggugaccucgaCcuucauga..............................................    3       1       seq
............aaggugacAucgaucuucauga..............................................    4       1       seq
............aaggugaccuAgaucuucauga..............................................    2       1       seq
............aaggugaccucgaucuucauga..............................................    1739    0       seq
............aaggugaccucgaucuucaCga..............................................    3       1       seq
............aaggAgaccucgaucuucauga..............................................    5       1       seq
............aaAgugaccucgaucuucaugau.............................................    31      1       seq
............aGggugaccucgaucuucaugau.............................................    89      1       seq
............aaggugaccucgaCcuucaugau.............................................    72      1       seq
............aaggugaccucgaucuAcaugau.............................................    21      1       seq
............aaUgugaccucgaucuucaugau.............................................    11      1       seq
............aaggugaccucgaucuuGaugau.............................................    1       1       seq
............aagguUaccucgaucuucaugau.............................................    10      1       seq
............aagguAaccucgaucuucaugau.............................................    25      1       seq
............aaggugaccucgaucuGcaugau.............................................    7       1       seq
............aaggugaccucgaucuucaGgau.............................................    6       1       seq
............aaggugaccucgaucuucaugCu.............................................    5       1       seq
............aaggugaccucgaucuCcaugau.............................................    91      1       seq
............aaggugaccucgaucuucaCgau.............................................    54      1       seq
............aaggugaccucgaucuucaugau.............................................    23287   0       seq
............aaggugUccucgaucuucaugau.............................................    28      1       seq
............aaggugaccucgaucuuUaugau.............................................    36      1       seq
............aaggugaAcucgaucuucaugau.............................................    10      1       seq
............aaggugaccuGgaucuucaugau.............................................    2       1       seq
............aaggugaccucgaucuuAaugau.............................................    8       1       seq
............aaggugaccucgauUuucaugau.............................................    15      1       seq
............aaggugaccucUaucuucaugau.............................................    6       1       seq
............aUggugaccucgaucuucaugau.............................................    24      1       seq
............aaggugaUcucgaucuucaugau.............................................    28      1       seq
............Uaggugaccucgaucuucaugau.............................................    34      1       seq
............aaggugaccucgaucuucaAgau.............................................    34      1       seq
............aaggugaccucgaAcuucaugau.............................................    20      1       seq
............aaggGgaccucgaucuucaugau.............................................    19      1       seq
............aaggugaccucgaucAucaugau.............................................    19      1       seq
```

novel-nve-miR-8_star

novel-nve-miR-8_guide

caugaagauccc**aaggugaccucgaucuucaugau**caugaucauc**augaugauugcaggccauuucgg**gaucuucugaag

| | | | |
|---|---|---|---|
| ............aaggugaccucgaucCucaugau............................................. | 110 | 1 | seq |
| ............aaggugaccucgaucGucaugau............................................. | 7 | 1 | seq |
| ............aaggugaccCcgaucuucaugau............................................. | 102 | 1 | seq |
| ............aaggugaccucgaucuucaugaA............................................. | 430 | 1 | seq |
| ............aaggugaccucgaGcuucaugau............................................. | 4 | 1 | seq |
| ............aaggugaccucgaucuucauCau............................................. | 6 | 1 | seq |
| ............Caggugaccucgaucuucaugau............................................. | 4 | 1 | seq |
| ............aaggugCccucgaucuucaugau............................................. | 29 | 1 | seq |
| ............aaggugaccucgGucuucaugau............................................. | 95 | 1 | seq |
| ............aaggugaccuAgaucuucaugau............................................. | 20 | 1 | seq |
| ............aaggugacUucgaucuucaugau............................................. | 65 | 1 | seq |
| ............Gaggugaccucgaucuucaugau............................................. | 199 | 1 | seq |
| ............aaggugaccucgCucuucaugau............................................. | 5 | 1 | seq |
| ............aaggugaccucgaucuucCugau............................................. | 6 | 1 | seq |
| ............aagAugaccucgaucuucaugau............................................. | 21 | 1 | seq |
| ............aCggugaccucgaucuucaugau............................................. | 6 | 1 | seq |
| ............aaCgugaccucgaucuucaugau............................................. | 11 | 1 | seq |
| ............aaggugaccucgauGuucaugau............................................. | 1 | 1 | seq |
| ............aaggugacAucgaucuucaugau............................................. | 44 | 1 | seq |
| ............aagCugaccucgaucuucaugau............................................. | 20 | 1 | seq |
| ............aagUugaccucgaucuucaugau............................................. | 11 | 1 | seq |
| ............aaggugaccucgaucuucaugUu............................................. | 13 | 1 | seq |
| ............aaggugaccucCaucuucaugau............................................. | 5 | 1 | seq |
| ............aaggugaccucgaucuucauAau............................................. | 23 | 1 | seq |
| ............aaggAgaccucgaucuucaugau............................................. | 54 | 1 | seq |
| ............aaggugaccucAaucuucaugau............................................. | 26 | 1 | seq |
| ............aaggugaccucgUucuucaugau............................................. | 8 | 1 | seq |
| ............aagguCaccucgaucuucaugau............................................. | 12 | 1 | seq |
| ............aaggugaccucgaucuucauUau............................................. | 4 | 1 | seq |
| ............aaggugaccGcgaucuucaugau............................................. | 8 | 1 | seq |
| ............aaggugaccucgaucuucUugau............................................. | 16 | 1 | seq |
| ............aaggugGccucgaucuucaugau............................................. | 124 | 1 | seq |
| ............aaggugaccAcgaucuucaugau............................................. | 33 | 1 | seq |
| ............aaggugaccucgaucuucaugaG............................................. | 210 | 1 | seq |
| ............aaggugaccucgaucuucGugau............................................. | 105 | 1 | seq |
| ............aaggugaccucgaucuucaugaC............................................. | 2960 | 1 | seq |
| ............aaggugaccuUgaucuucaugau............................................. | 44 | 1 | seq |
| ............aaggugaccucgaucuucaugGu............................................. | 134 | 1 | seq |
| ............aaggugaccucgauAuucaugau............................................. | 18 | 1 | seq |
| ............aaggugacGucgaucuucaugau............................................. | 4 | 1 | seq |
| ............aaggCgaccucgaucuucaugau............................................. | 75 | 1 | seq |
| ............aaggugaccucgaucuucaugauU............................................ | 1504 | 1 | seq |
| ............aaggugaccucgaucuucaCgauc............................................ | 1 | 1 | seq |
| ............aaggugaccucgaCcuucaugauc............................................ | 1 | 1 | seq |
| ............aaggugacUucgaucuucaugauc............................................ | 1 | 1 | seq |
| ............aaggugGccucgaucuucaugauc............................................ | 1 | 1 | seq |
| ............aaggugaccucgaucuucaugauA............................................ | 40 | 1 | seq |
| ............aaggugCccucgaucuucaugauc............................................ | 1 | 1 | seq |
| ............Uaggugaccucgaucuucaugauc............................................ | 1 | 1 | seq |
| ............aaggugaccucgaucuucaAgauc............................................ | 2 | 1 | seq |
| ............aaggugaccucgaucuucaugauc............................................ | 172 | 0 | seq |
| ............aaggugaccucgaucuuUaugauc............................................ | 1 | 1 | seq |
| ............aaggugaccucgGucuucaugauc............................................ | 1 | 1 | seq |
| ............aaggugaccucgaucuucaugauG............................................ | 7 | 1 | seq |
| ............aaggugaccucgaucuucaugGuc............................................ | 1 | 1 | seq |
| ............aagAugaccucgaucuucaugauc............................................ | 1 | 1 | seq |
| ............aaggugaccucgaucuucGugauc............................................ | 2 | 1 | seq |
| ............aaggugaccucgaucuucauCauc............................................ | 1 | 1 | seq |
| ............aaggugaccucgaucuucaugauAa........................................... | 1 | 1 | seq |
| ............aaggugaccucgaucuucaugauUa........................................... | 42 | 1 | seq |
| ............aaggugaccucgaucuucaugauca........................................... | 2 | 0 | seq |
| ............aaggugaccucgaucuucaugaucU........................................... | 2 | 1 | seq |
| ............aaggugaccucgaucuucaugauUau.......................................... | 1 | 1 | seq |
| .............Gggugaccucgaucuucaug............................................... | 1 | 1 | seq |
| .............aggugaccucgGucuucaug............................................... | 1 | 1 | seq |
| .............aggugaccucgaucuucaug............................................... | 6 | 0 | seq |
| .............aggugaccucgaucuucaugG.............................................. | 1 | 1 | seq |
| .............aAgugaccucgaucuucauga.............................................. | 2 | 1 | seq |
| .............Gggugaccucgaucuucauga.............................................. | 3 | 1 | seq |

novel-nve-miR-8_star

novel-nve-miR-8_guide

caugaagauccc**aaggugaccucgaucuucaugau**caugaucauc**augaugauugcaggccauuucgg**gaucuucugaag

| | | | |
|---|---|---|---|
| .............aggugaccucgaucuucauga.............................................. | 89 | 0 | seq |
| .............aggugaccucgaucuCcauga.............................................. | 1 | 1 | seq |
| .............Uggugaccucgaucuucauga.............................................. | 1 | 1 | seq |
| .............aggugaccucgaucuucaugU.............................................. | 2 | 1 | seq |
| .............aggugaccucgaucuucaAgau............................................. | 1 | 1 | seq |
| .............aggugaccucgaucuCcaugau............................................. | 2 | 1 | seq |
| .............aggugaccuUgaucuucaugau............................................. | 3 | 1 | seq |
| .............aggugaccucgaucuucaugau............................................. | 663 | 0 | seq |
| .............aggCgaccucgaucuucaugau............................................. | 1 | 1 | seq |
| .............aAgugaccucgaucuucaugau............................................. | 11 | 1 | seq |
| .............aggugaccucgaucGucaugau............................................. | 1 | 1 | seq |
| .............agCugaccucgaucuucaugau............................................. | 2 | 1 | seq |
| .............agguCaccucgaucuucaugau............................................. | 1 | 1 | seq |
| .............aggugaccuGgaucuucaugau............................................. | 1 | 1 | seq |
| .............aggugaccucgaucuucGugau............................................. | 3 | 1 | seq |
| .............aggugaUcucgaucuucaugau............................................. | 1 | 1 | seq |
| .............aggugaccucgaucuucaugaA............................................. | 10 | 1 | seq |
| .............aggugaccucgaucuucaCgau............................................. | 3 | 1 | seq |
| .............aggugaccucgGucuucaugau............................................. | 3 | 1 | seq |
| .............aggugCccucgaucuucaugau............................................. | 1 | 1 | seq |
| .............aggugGccucgaucuucaugau............................................. | 2 | 1 | seq |
| .............Gggugaccucgaucuucaugau............................................. | 5 | 1 | seq |
| .............aggugaccucgaucuGcaugau............................................. | 1 | 1 | seq |
| .............aggugaccucgaucuucaugaC............................................. | 71 | 1 | seq |
| .............aggugacUucgaucuucaugau............................................. | 2 | 1 | seq |
| .............Uggugaccucgaucuucaugau............................................. | 3 | 1 | seq |
| .............aggugaccucgaCcuucaugau............................................. | 1 | 1 | seq |
| .............aggugaccucgaucuucaugaG............................................. | 7 | 1 | seq |
| .............aggugaccucgaucCucaugau............................................. | 1 | 1 | seq |
| .............aggugacAucgaucuucaugau............................................. | 2 | 1 | seq |
| .............aggAgaccucgaucuucaugau............................................. | 4 | 1 | seq |
| .............aggugaccucgaucuucaugUu............................................. | 1 | 1 | seq |
| .............agguAaccucgaucuucaugau............................................. | 2 | 1 | seq |
| .............aggugaccAcgaucuucaugau............................................. | 1 | 1 | seq |
| .............aggugaccucgaucuucaugGu............................................. | 7 | 1 | seq |
| .............aggugaccucgaucuucauAau............................................. | 1 | 1 | seq |
| .............aggugaccCcgaucuucaugau............................................. | 3 | 1 | seq |
| .............aggugaccucgaucuucaugauc............................................ | 8 | 0 | seq |
| .............aggugaccucgaucuucaugauA............................................ | 5 | 1 | seq |
| .............aggugaccucgaucCucaugauc............................................ | 1 | 1 | seq |
| .............aggugaccucgaucuucaugauU............................................ | 107 | 1 | seq |
| .............aggugaccucgaucuucaugauUa........................................... | 2 | 1 | seq |
| .............aggugaccucgaucuucaugaucU........................................... | 3 | 1 | seq |
| ..............ggugaccucgaucuucaug............................................... | 2 | 0 | seq |
| ..............ggugaccucgaucuucauga.............................................. | 3 | 0 | seq |
| ..............ggugaccucgaucuucaugaC............................................. | 9 | 1 | seq |
| ..............gguCaccucgaucuucaugau............................................. | 1 | 1 | seq |
| ..............ggCgaccucgaucuucaugau............................................. | 1 | 1 | seq |
| ..............ggugaccucgauUucaugau............................................. | 1 | 1 | seq |
| ..............ggugaccucgaucuucaugaA............................................. | 1 | 1 | seq |
| ..............ggugacUucgaucuucaugau............................................. | 1 | 1 | seq |
| ..............ggugaccucgaucuucaugau............................................. | 76 | 0 | seq |
| ..............ggugaccucgaCcuucaugau............................................. | 1 | 1 | seq |
| ..............ggugaccucgaucuucaugauU............................................ | 10 | 1 | seq |
| ..............ggugaccucgaucuucaugauc............................................ | 1 | 0 | seq |
| ...............gugaccucgaucuucaugau............................................. | 5 | 0 | seq |
| ............................................caugaugauugcaggcca.................. | 1 | 0 | seq |
| ............................................caugaugauugcaggccaC................. | 1 | 1 | seq |
| ............................................caugaugGuugcaggccau................. | 1 | 1 | seq |
| ............................................caugaugauugcaggccau................. | 3 | 0 | seq |
| ............................................caugaugauugUaggccauu................ | 2 | 1 | seq |
| ............................................caugaugauugcaggccauu................ | 1 | 0 | seq |
| ............................................caugaugauugcaggccauuC............... | 1 | 1 | seq |
| ............................................Gaugaugauugcaggccauuu............... | 1 | 1 | seq |
| ............................................caugaugauugcaggccauuu............... | 12 | 0 | seq |
| ............................................caugaugauugcaggccauuuA.............. | 1 | 1 | seq |
| ............................................caugaugauugcaggccauuuU.............. | 1 | 1 | seq |
| ............................................caugaugaAugcaggccauuuc.............. | 1 | 1 | seq |
| ............................................caugaugauugcaggccauuuc.............. | 23 | 0 | seq |

novel-nve-miR-8_star

novel-nve-miR-8_guide

caugaagaucccaaggugaccucgaucuucaugaucaugaucaucaugaugauugcaggccauuucgggaucuucugaag

| | | | |
|---|---|---|---|
| ............................................caugaugauugcaggccGuuuc.............. | 1 | 1 | seq |
| ............................................caugaugauugcaggccauuucg............. | 16 | 0 | seq |
| ............................................caugaugauugcaggccauuucA............. | 2 | 1 | seq |
| ............................................caugaCgauugcaggccauuucg............. | 1 | 1 | seq |
| ............................................caugaugauugcaggccauuucU............. | 4 | 1 | seq |
| ............................................caugaugauugcaggccauuucgg............ | 82 | 0 | seq |
| ............................................caugGugauugcaggccauuucgg............ | 2 | 1 | seq |
| ............................................caugaugauugcaggccauuucgA............ | 7 | 1 | seq |
| ............................................caGgaugauugcaggccauuucgg............ | 1 | 1 | seq |
| ............................................caugaugauugcGggccauuucgg............ | 1 | 1 | seq |
| ............................................caugaugauugcaggccaAuucgg............ | 1 | 1 | seq |
| ............................................caugaAgauugcaggccauuucgg............ | 1 | 1 | seq |
| ............................................caugaugauugUaggccauuucgg............ | 1 | 1 | seq |
| ............................................caugaugauugcaggccauuucAg............ | 1 | 1 | seq |
| ............................................caugaugauugcaggccauuucgU............ | 1 | 1 | seq |
| ............................................caugaugauugcaggccauuucggU........... | 4 | 1 | seq |
| ............................................caugaugauugcaggccauuucggA........... | 4 | 1 | seq |
| ............................................caugaugauugcaggccauuucggUa.......... | 1 | 1 | seq |
| .............................................augaugauugcaggccauA................ | 1 | 1 | seq |
| .............................................augaugauugcaggccauuuA.............. | 1 | 1 | seq |
| .............................................augaugauugcaggccauuuc.............. | 18 | 0 | seq |
| .............................................augaugauugcaggccauuuU.............. | 1 | 1 | seq |
| .............................................augaugauugcUggccauuuc.............. | 1 | 1 | seq |
| .............................................augGugauugcaggccauuucg............. | 2 | 1 | seq |
| .............................................augaugauugcaggccauuucA............. | 1 | 1 | seq |
| .............................................augaugauugcaggccauuucg............. | 11 | 0 | seq |
| .............................................aAgaugauugcaggccauuucg............. | 1 | 1 | seq |
| .............................................augaugauugcaggccauuucgA............ | 9 | 1 | seq |
| .............................................Gugaugauugcaggccauuucgg............ | 3 | 1 | seq |
| .............................................augaugauCgcaggccauuucgg............ | 1 | 1 | seq |
| .............................................augaCgauugcaggccauuucgg............ | 1 | 1 | seq |
| .............................................aCgaugauugcaggccauuucgg............ | 1 | 1 | seq |
| .............................................augaugauugcaggccGuuucgg............ | 3 | 1 | seq |
| .............................................augaugauugcaggccauuucgU............ | 1 | 1 | seq |
| .............................................augaAgauugcaggccauuucgg............ | 3 | 1 | seq |
| .............................................augaugauugcaggccauuucgg............ | 110 | 0 | seq |
| .............................................augaugauugcaUgccauuucgg............ | 1 | 1 | seq |
| .............................................augaugauugcaggccauuucggA........... | 3 | 1 | seq |
| .............................................augaugauugcaggccauuucggUau......... | 1 | 1 | seq |
| ..............................................ugaugauugcaggccauuu............... | 2 | 0 | seq |
| ..............................................ugaugauugcaggccauuuc.............. | 5 | 0 | seq |
| ..............................................ugaugauugcCggccauuucg............. | 1 | 1 | seq |
| ..............................................ugaugauugcagAccauuucg............. | 1 | 1 | seq |
| ..............................................ugaugauugcaggccauuucA............. | 1 | 1 | seq |
| ..............................................ugaugauCgcaggccauuucg............. | 1 | 1 | seq |
| ..............................................ugaugauugcaggccauuucg............. | 5 | 0 | seq |
| ..............................................ugaugauugcaggccauuucAg............ | 1 | 1 | seq |
| ..............................................ugaugauugcaggccauuucgC............ | 1 | 1 | seq |
| ..............................................ugaugauugcaggccauuucgg............ | 40 | 0 | seq |
| ..............................................Agaugauugcaggccauuucgg............ | 1 | 1 | seq |
| ..............................................Cgaugauugcaggccauuucgg............ | 1 | 1 | seq |
| ..............................................ugaugauugcaggccauuucgU............ | 2 | 1 | seq |
| ..............................................ugaugauugcaggccauuucgA............ | 6 | 1 | seq |
| ..............................................ugaugauugcaggccauuucggU........... | 3 | 1 | seq |

miRBase precursor : novel-nve-miR-9
Total read count : 162
novel-nve-miR-9_guide read count : 161
novel-nve-miR-9_star read count : 1
remaining reads : 0

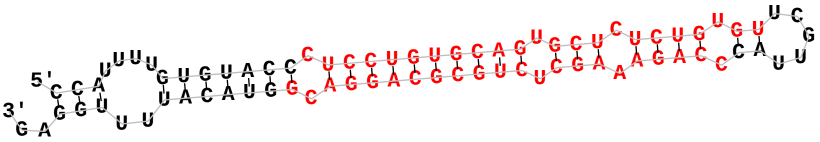

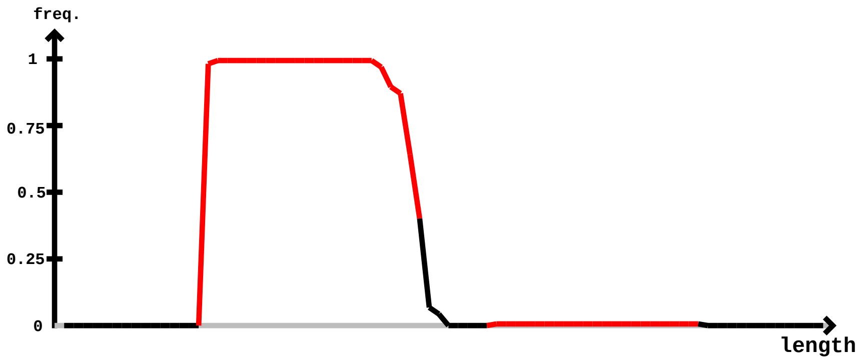


novel-nve-miR-9_guide

novel-nve-miR-9_star

```
5'- ccauuuuguguacccuccugugcagugcucucuguguucguuacccagaaagcucugcgcaggacgguacauuuuggag  -3'   exp
    (((....(((((((.((((((((((.(((.((((.((.....)).)))).))).)))))))))).)))))))..)))..     reads     mm      sample
    ..............cuccugugcagugcucuc...............................................     4         0       seq
    ..............cuccugugcagugcucucC..............................................     2         1       seq
    ..............cuccugugcagugcucucu..............................................     9         0       seq
    ..............cCccugugcagugcucucu..............................................     1         1       seq
    ..............cuccugugcagugcucucug.............................................     4         0       seq
    ..............cuccugugcagCgcucucugu............................................     1         1       seq
    ..............cuccugugcagugcucucugC............................................     1         1       seq
    ..............cuccugugcagugcucucugu............................................     35        0       seq
    ..............cuccugugcagugcucucugug...........................................     37        0       seq
    ..............cuccugugcagugcucucugugC..........................................     4         1       seq
    ..............cuccugugcagCgcucucugugu..........................................     1         1       seq
    ..............cuccugugcagugcucucugugA..........................................     1         1       seq
    ..............cuccugugcagugcucucuguUu..........................................     1         1       seq
    ..............cuccugugcagugcAcucugugu..........................................     1         1       seq
    ..............cuccugugcagugcucucugugu..........................................     44        0       seq
    ..............cuccugugcagugcucucugugG..........................................     1         1       seq
    ..............cuccugugcGgugcucucugugu..........................................     1         1       seq
    ..............cuccugugcagugcucucuguguu.........................................     2         0       seq
    ..............cuccugugcagugcucucuguguC.........................................     2         1       seq
    ..............cuccugugcagugcucucuguguuc........................................     1         0       seq
    ..............cuccugugcagugcucucuguguuU........................................     6         1       seq
    ...............uccugugcagugcucucuguA...........................................     1         1       seq
    ...............uccugugcagugcucucugug...........................................     1         0       seq
    ............................................ccagaaagcucugcgcaggacg.............     1         0       seq
```

miRBase precursor : novel-nve-miR-10
Total read count : 586
novel-nve-miR-10_guide read count : 565
novel-nve-miR-10_star read count : 11
remaining reads : 10

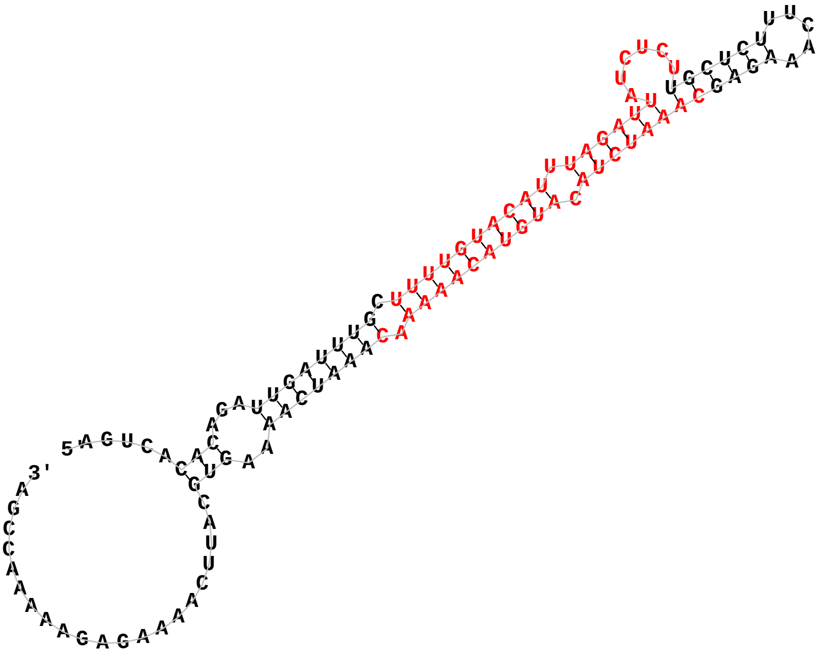

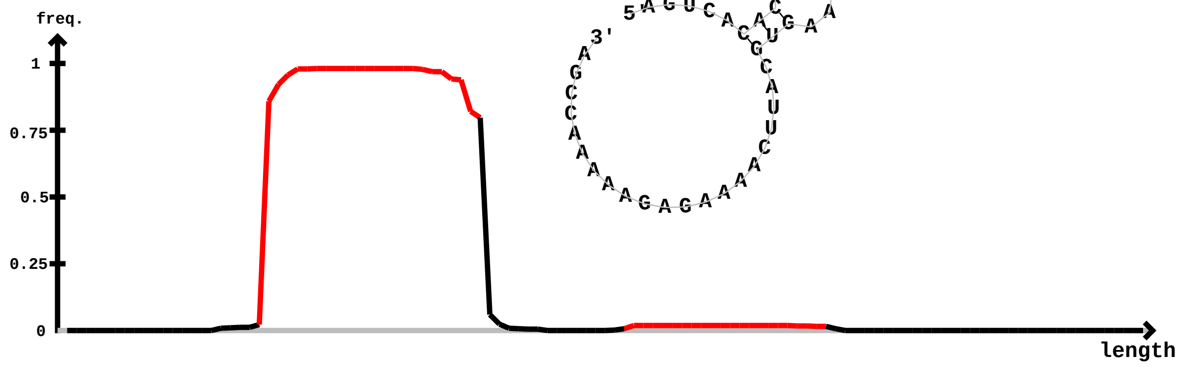


novel-nve-miR-10_guide

novel-nve-miR-10_star

```
5'- agucacacagauugauuugcuuuuguacauuuagauuaucucuugcucuuucaaagagcaaaucuacauguacaaaaacaaaucaaaagugcauucaaaagagaaaaaccga   -3'    exp
    .....(((...((((((((.((((((((((.((((((......(((((.....))))))))))).)))))))))).))))))))..)))......................   reads    mm    sample
    ...............uuugcuuuuguacauGuagau............................................................................   1        1     seq
    ...............uuugcuuuuguacauGuagauu...........................................................................   4        1     seq
    ................uugcuuuuguacauGuagauu...........................................................................   1        1     seq
    .................ugcuuuuguacauGuagau............................................................................   1        1     seq
    ...................cuuuuguacauGuagauuau.........................................................................   1        1     seq
    ...................cuuuuguacauGuagauuaucu.......................................................................   1        1     seq
    ...................cuuuuguacauuuagauuaucucu.....................................................................   1        0     seq
    ...................cuuuuguacauGuagauuaucucu.....................................................................   3        1     seq
    ....................uuuuguacauGuagauuau.........................................................................   15       1     seq
    ....................uuuuguacauGuagauuauc........................................................................   2        1     seq
    ....................uuuuguacauuuagauuaucA.......................................................................   1        1     seq
    ....................uuuuguacauuuagauuaucu.......................................................................   4        0     seq
    ....................uuuuguacauGuagauuaucu.......................................................................   59       1     seq
    ....................uuuuguacauuuagauuaucuc......................................................................   1        0     seq
    ....................uuuuguacauGuagauuaucuc......................................................................   11       1     seq
    ....................uuuuguacauAuagauuaucucu.....................................................................   1        1     seq
    ....................uuuuguacauuuagauuaucucG.....................................................................   2        1     seq
    ....................uuuuguacauuuagauuaucucu.....................................................................   37       0     seq
    ....................uuuuguacauGuagauuaucucu.....................................................................   327      1     seq
    ....................uuuuguacauuuagauuaucucC.....................................................................   7        1     seq
    ....................uuuuguacauuuagauuaucucA.....................................................................   2        1     seq
    ....................uuuuguacauuuagauuaucucuu....................................................................   2        0     seq
    ....................uuuuguacauGuagauuaucucuu....................................................................   10       1     seq
    ....................uuuuguacauuuagauuaucucuug...................................................................   2        0     seq
    ....................uuuuguacauGuagauuaucucuug...................................................................   3        1     seq
    ....................uuuuguacauGuagauuaucucuugcu.................................................................   1        1     seq
    ....................uuuuguacauGuagauuaucucuugcucu...............................................................   3        1     seq
    .....................uuuguacauGuagauuaucu.......................................................................   4        1     seq
    .....................uuuguacauGuagauuaucuc......................................................................   1        1     seq
    .....................uuuguacauGuagauuaucucu.....................................................................   25       1     seq
    .....................uuuguacauuuagauuaucucu.....................................................................   1        0     seq
    .....................uuuguacauGuagauuaucucuu....................................................................   3        1     seq
    .....................uuuguacauGuagauuaucucuug...................................................................   2        1     seq
    .....................uuuguacauGuagauuaucucuugc..................................................................   1        1     seq
```

```
                    novel-nve-miR-10_guide
                                                                          novel-nve-miR-10_star
agucacacagauugauuugcuuuuguacauuuagauuaucucuugcucuuucaaagagcaaaucuacauguacaaaaacaaaucaaaagugcauucaaaagagaaaaaccga

......................uuguacauGuagauuaucucu.....................................................................    19    1    seq
......................uuguacauuuagauuaucucu.....................................................................    1     0    seq
......................uuguacauuuagauuaucucA.....................................................................    1     1    seq
.......................uguacauuuagauuaucuc......................................................................    1     0    seq
.......................uguacauGuagauuaucucu.....................................................................    4     1    seq
.......................uguacauuuagauuaucucu.....................................................................    1     0    seq
.......................uguacauuuagauuaucucuu....................................................................    1     0    seq
.......................uguacauuuagauuaucucuC....................................................................    1     1    seq
.......................uguacauGuagauuaucucuu....................................................................    3     1    seq
.......................uguacauGuagauuaucucuug...................................................................    1     1    seq
.......................uguacauuuagauuaucucuuU...................................................................    1     1    seq
.........................uacauGuagauuaucucuu....................................................................    1     1    seq
........................................................agcaaaucuacauguacaa.....................................    1     0    seq
.........................................................gcaaaucuacauguacaaaaaca................................    1     0    seq
.........................................................gcaaaucuacauguacaaaaacU................................    2     1    seq
..........................................................caaaucuacauguacaaaa...................................    1     0    seq
..........................................................caaaucuacauguacaaaaac.................................    5     0    seq
..........................................................caaaucuacauguacaaaaacC................................    1     1    seq
```

miRBase precursor : novel-nve-miR-11
Total read count : 62
novel-nve-miR-11_guide read count : 58
novel-nve-miR-11_star read count : 4
remaining reads : 0

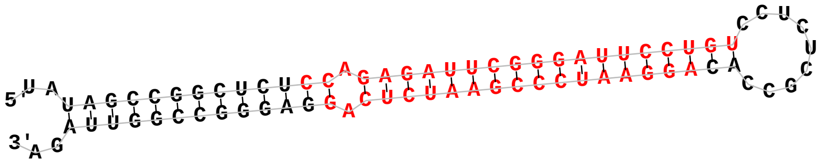


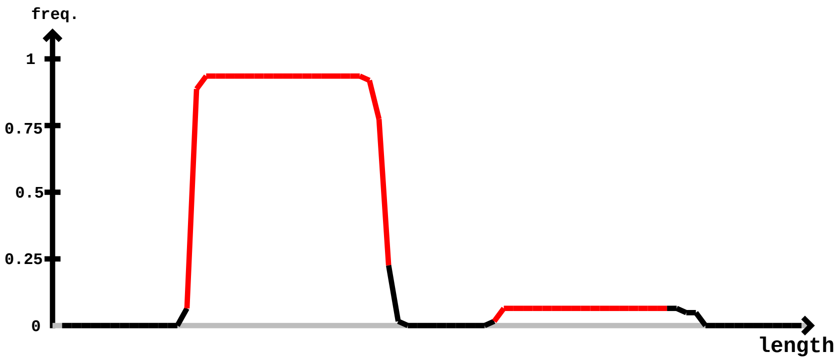


```
                                                   novel-nve-miR-11_star
                   novel-nve-miR-11_guide
5'- uauagccggcucuccagagauucgggauuccuguccucucgccacaggaaucccgaaucucaggagggccgguuaga   -3'    exp
    ..(((((((((((((.((((((((((((((((((.........)))))))))))))))))).)))))))))))))..    reads      mm         sample
    ...........uccagagauucgggauuccug.............................................    2          0          seq
    ...........uccGgagauucgggauuccug.............................................    1          1          seq
    ...........uccGgagauucgggauuccuguc...........................................    1          1          seq
    ............ccGgagauucgggauucc...............................................    1          1          seq
    ............ccagagauucgggauuccu..............................................    1          0          seq
    ............ccGgagauucgggauuccu..............................................    6          1          seq
    ............ccGgagauucgggauuccug.............................................    28         1          seq
    ............ccUgagauucgggauuccug.............................................    2          1          seq
    ............ccGgagauucgggauuccugu............................................    13         1          seq
    .............cGgagauucgggauuccu..............................................    2          1          seq
    .............cGgagauucgggauuccug.............................................    1          1          seq
    ............................................caggaaucccgaaucucagg.............    1          0          seq
    .............................................aggaaucccgaaucucaggag...........    1          0          seq
    .............................................aggaaucccgaaucucaggaU...........    1          1          seq
    .............................................aggaaucccgaaucucCggag...........    1          1          seq
```

miRBase precursor : novel-nve-miR-12
Total read count : 135
novel-nve-miR-12_guide read count : 121
novel-nve-miR-12_star read count : 8
remaining reads : 6

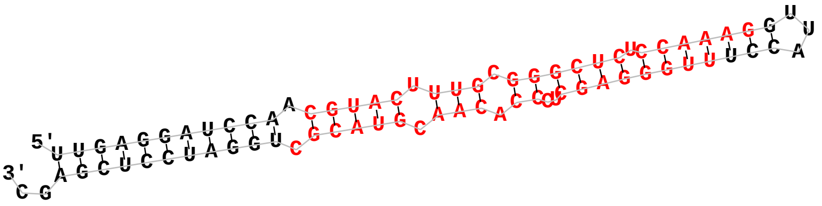

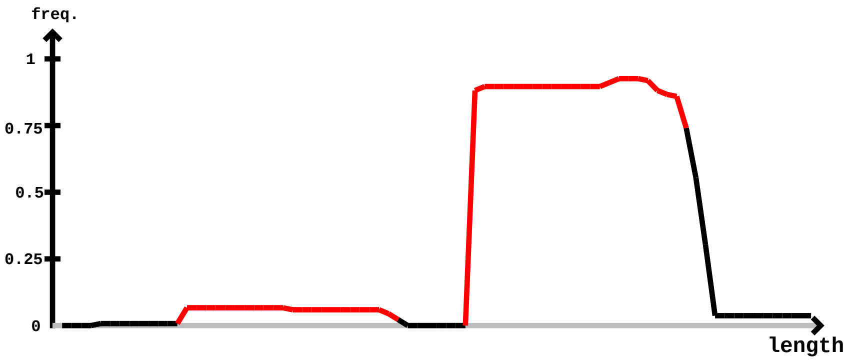


```
                      novel-nve-miR-12_star
                                              novel-nve-miR-12_guide
5'- uugaggauccaacguacuuugcgggcucuccaaagguuaccuuugggagcucccacaacguacgcuggauccucgagc   -3'    exp
    (((((((((((.(((((.(((.((((((.(((((((...)))))))))))..)).))).))))).)))))))))))..   reads      mm         sample
    ...aggauccaGcguacuuugcg.......................................................   1          1          seq
    ............cguacuuugcgggcucuccaa.............................................   2          0          seq
    ............cguacuuugcgggcucuccaaa............................................   3          0          seq
    ............cguacuuugcgggcucuccaaaU...........................................   1          1          seq
    ............cguacuuugcgggcucuccaaag...........................................   2          0          seq
    ..........................................uugggagcucccacaacg..................   2          0          seq
    ..........................................uugggagcucccacaacgA.................   1          1          seq
    ..........................................uugggagcucccacaacgC.................   2          1          seq
    ..........................................uugggagcucccacaacgu.................   2          0          seq
    ..........................................uugggagcucccacaacgua................   2          0          seq
    ..........................................uugggagcucccacaacguac...............   1          0          seq
    ..........................................uugggagcucccacaacguacU..............   1          1          seq
    ..........................................uugggagcucccacaacguacA..............   2          1          seq
    ..........................................uugggagcucccacaacguacg..............   12         0          seq
    ..........................................uugAgagcucccacaacguacgc.............   1          1          seq
    ..........................................uugggagcucccacaacguacgU.............   9          1          seq
    ..........................................uugggagcucccacaacguacgA.............   3          1          seq
    ..........................................uugggagcucccacaacguacgc.............   12         0          seq
    ..........................................uugggagcucGcacaacguacgcu............   1          1          seq
    ..........................................uugggagcucccacaacguacgcA............   1          1          seq
    ..........................................uugggagcucccacaacguacgcu............   20         0          seq
    ..........................................uugggagcucccacaacguacgcG............   3          1          seq
    ..........................................uugggagcucccacaacguacgcC............   8          1          seq
    ..........................................uugggagcuAccacaacguacgcu............   1          1          seq
    ..........................................uugggagcucccacaacguacgcuU...........   28         1          seq
    ..........................................uugggagcucccacaacguacgcuC...........   7          1          seq
    ...........................................ugggagcucccacaacguacg..............   1          0          seq
    ...........................................ugggagcucccacaacguacgcuU...........   1          1          seq
    ........................................................aaAguacgcuggauccucgagc   2          1          seq
    .........................................................aAguacgcuggauccucgagc   2          1          seq
    ............................................................uacgcuggauccucgagc   1          0          seq
```

miRBase precursor : novel-nve-miR-13
Total read count : 98
novel-nve-miR-13_guide read count : 77
novel-nve-miR-13_star read count : 5
remaining reads : 16

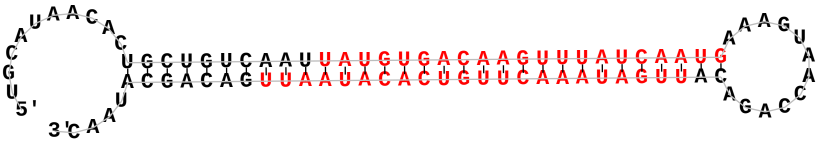

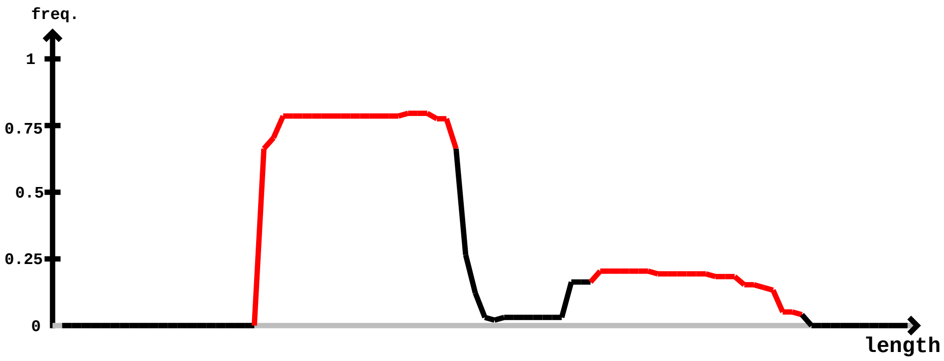


novel-nve-miR-13_star
novel-nve-miR-13_guide

```
5'- ugcauaacacugcugucaauuaugugacaaguuuaucaaugaaaguaaccagacauugauaaacuugucacauaauugacagcauaac  -3'   exp
    ..........(((((((((((((((((((((((((((((((............)))))))))))))))))))))))))))))))....   reads   mm   sample
    ....................uaugugacaaguuuaucG..................................................   1   1   seq
    ....................uaugugacaaguuuaCca..................................................   1   1   seq
    ....................uaugugacaaguuuaucaa.................................................   1   0   seq
    ....................uaugugacaaguuuaucaau................................................   8   0   seq
    ....................uGugugacaaguuuaucaau................................................   1   1   seq
    ....................uaugGgacaaguuuaucaau................................................   1   1   seq
    ....................uaugCgacaaguuuaucaaug...............................................   1   1   seq
    ....................uaugugacaaguuuaCcaaug...............................................   2   1   seq
    ....................uaugugacaaguuuaucaaug...............................................   26   0   seq
    ....................uaugugacGaguuuaucaaug...............................................   1   1   seq
    ....................uaugugacaaguuuaucaauA...............................................   4   1   seq
    ....................uaugugacaaguuuaucaauga..............................................   2   0   seq
    ....................uaugugacaaguuuaucaaugU..............................................   8   1   seq
    ....................uaugugacaaguuuaucaaugC..............................................   1   1   seq
    ....................uaugugacaaguuuaucaaugaC.............................................   1   1   seq
    ....................uaugugacaaguuuaucaaugaU.............................................   6   1   seq
    .....................Gugugacaaguuuaucaaug...............................................   1   1   seq
    .....................augugacaaguuuaucaaug...............................................   1   0   seq
    .....................augugacaaguuuaucGaug...............................................   1   1   seq
    .....................augugacaaguuuaucaaugaCa............................................   1   1   seq
    ......................ugugacaaguuuaucaau................................................   1   0   seq
    ......................ugugaAaaguuuaucaaug...............................................   1   1   seq
    ......................ugugacaaguuuaucaaug...............................................   1   0   seq
    ......................ugugacaaguuuaucaaugU..............................................   3   1   seq
    ......................ugugacaaguuuaucaaugaU.............................................   2   1   seq
    ...................................ucaaugaaaguaaccagaca.................................   1   0   seq
    .......................................ugaaaguaaccagacauugaua...........................   1   0   seq
    .............................................uaaccagacauugauaaacuuU.....................   1   1   seq
    .....................................................Ucauugauaaacuuguca.................   3   1   seq
    .....................................................Ucauugauaaacuugucaca...............   1   1   seq
    .....................................................Ucauugauaaacuugucacau..............   1   1   seq
    .....................................................Ucauugauaaacuugucacaua.............   8   1   seq
    .......................................................uugauaaacuugucacauaau............   1   0   seq
    .......................................................uugauaaacuugucacauaaCu...........   1   1   seq
```

```
                                                      novel-nve-miR-13_star
                    novel-nve-miR-13_guide
ugcauaacacugcugucaauuaugugacaaguuuaucaauggaaaguaaccagacauugauaaacuugucacauaauugacagcauaac

......................................................uugauaaacuugucacauaauu...........    1          0          seq
......................................................uugauGaacuugucacauaauu...........    2          1          seq
```

miRBase precursor : novel-nve-miR-14
Total read count : 115
novel-nve-miR-14_guide read count : 90
novel-nve-miR-14_star read count : 25
remaining reads : 0

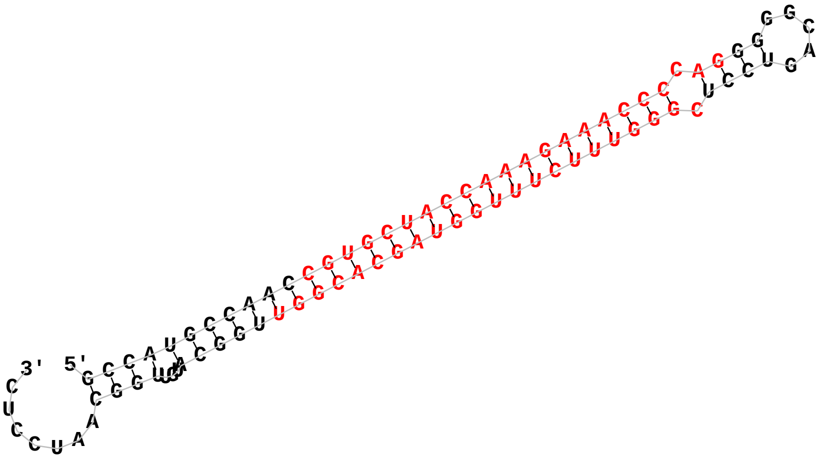

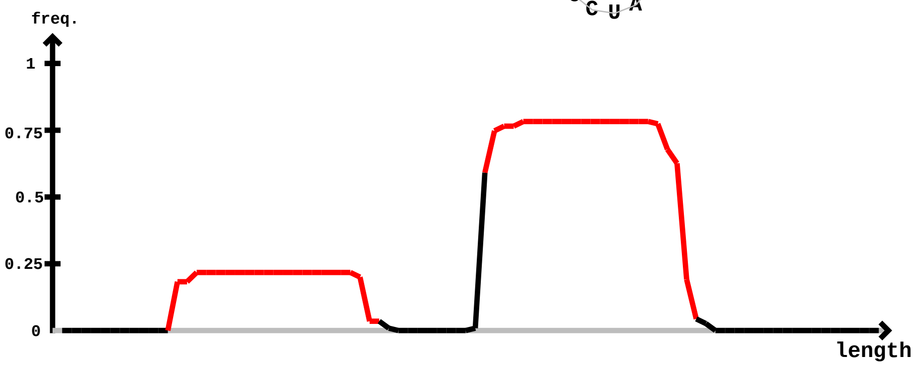


novel-nve-miR-14_guide

novel-nve-miR-14_star

```
5'- gccaugccaaccgugcuaccaaagaaaccccagggggcaguccucggguuucuuugguagcacgguuggcauguuggcaauccuc  -3'    exp
    ((((((((((((((((((((((((((((.((((.....)))).))))))))))))))))))))))))...)))).......    reads       mm          sample
    ...........cgugcuaccaaagaaaccc.......................................................    1           0           seq
    ...........cgugcuaccaaagaaaccA.......................................................    1           1           seq
    ...........cgugcuaccaaagaaacccU......................................................    8           1           seq
    ...........cgugcuaccaaagaaacccc......................................................    1           0           seq
    ...........cgugcuaccaaagaaacccG......................................................    9           1           seq
    ...........cgugcuaccaaagaaacccA......................................................    1           1           seq
    .............ugcuaccaaagaaaccccaA....................................................    1           1           seq
    .............ugcuaccaaagaaaccccag....................................................    2           0           seq
    .............ugcuaccaaagaaaccccaCg...................................................    1           1           seq
    ..........................................cucggguuucuuugguagcacU.....................    1           1           seq
    ...........................................ucggguuucuuugguagG........................    1           1           seq
    ...........................................ucggguuucuuugguagca.......................    2           0           seq
    ...........................................ucggguuucuuugguagUa.......................    1           1           seq
    ...........................................ucggguuucuuugguagcG.......................    8           1           seq
    ...........................................ucggguuucuuugguagcac......................    2           0           seq
    ...........................................ucggguuucuuugguagcaU......................    1           1           seq
    ...........................................ucggguuucuuugguagcacg.....................    25          0           seq
    ...........................................ucggguuucuuugguagcacA.....................    2           1           seq
    ...........................................ucggguuuUuuugguagcacg.....................    1           1           seq
    ...........................................uGggguuucuuugguagcacg.....................    8           1           seq
    ...........................................ucggguuucuuugguagcacC.....................    2           1           seq
    ...........................................ucggguuucuuugguagcacU.....................    6           1           seq
    ...........................................ucggguuucuuugguagcacgC....................    1           1           seq
    ...........................................ucggguuucuuugguagcacgU....................    1           1           seq
    ...........................................uGggguuucuuugguagcacgg....................    1           1           seq
    ...........................................ucggguuucuuugguagcacgA....................    1           1           seq
    ...........................................ucggguuucuuugguagcacgg....................    4           0           seq
    ............................................cggguuucuuugguagcac......................    3           0           seq
    ............................................cgggCuucuuugguagcacg.....................    1           1           seq
    ............................................cggguuucuuugguaUcacg.....................    1           1           seq
    ............................................cggguuucuuugguagcacg.....................    3           0           seq
    ............................................cggguuucuuugguagcacgg....................    6           0           seq
    ............................................cggguuucuuugguagcacggu...................    2           0           seq
    ............................................cggguuucuuugguagcacgguA..................    1           1           seq
```

```
                                            novel-nve-miR-14_guide
           novel-nve-miR-14_star
gccaugccaaccgugcuaccaaagaaaccccagggggcaguccucggguuucuuugguagcacgguuggcauguuggcaauccuc

............................................cggguuucuuugguagcacgguu..................    1          0          seq
.............................................ggguuucuuugguagcacgg....................    1          0          seq
.............................................ggguuucuuugguagcacgU....................    1          1          seq
...............................................guuucuuugguagcacgg....................    1          0          seq
...............................................guuucuuugguagcacgguu..................    1          0          seq
```

miRBase precursor : novel-nve-miR-15
Total read count : 545
novel-nve-miR-15_guide read count : 538
novel-nve-miR-15_star read count : 7
remaining reads : 0

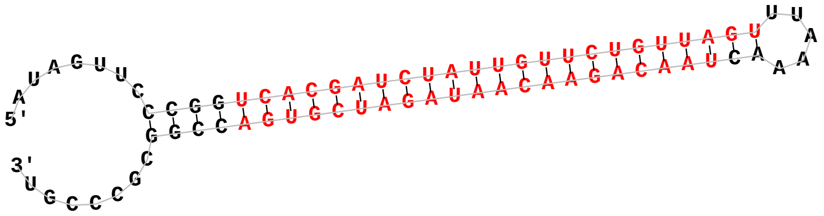


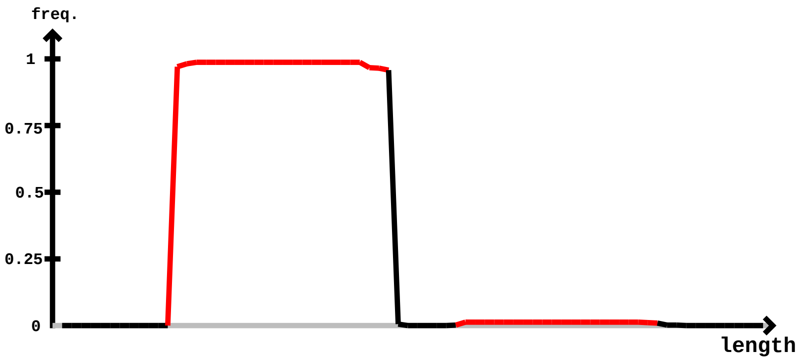


```
                novel-nve-miR-15_guide
                                     novel-nve-miR-15_star
5'- auaguucccggucacgaucuauuguucuguuaguuuaaaacuaacagaacaauagaucgugaccggcgcccgu   -3'    exp
    ...(..(((((((((((((((((((((((((((.....))))))))))))))))))))))))))).)..)..    reads      mm         sample
    ...........ucacgaucuauuguucuguu..........................................    7          0          seq
    ...........ucacgaucuauuguucuguC..........................................    3          1          seq
    ...........ucacgaucuauuguucugCu..........................................    1          1          seq
    ...........ucacgaucuauuguucuguua.........................................    1          0          seq
    ...........ucacgaucuauuguucuguuaC........................................    1          1          seq
    ...........ucacgaucuauuguucuguuaU........................................    3          1          seq
    ...........ucacgaucCauuguucuguuagu.......................................    4          1          seq
    ...........ucacgaucuauuguucugGuagu.......................................    1          1          seq
    ...........ucacgaucuauuguucuguuagG.......................................    12         1          seq
    ...........Acacgaucuauuguucuguuagu.......................................    5          1          seq
    ...........ucGcgaucuauuguucuguuagu.......................................    1          1          seq
    ...........ucacgaCcuauuguucuguuagu.......................................    2          1          seq
    ...........ucacgaucuauuguucCguuagu.......................................    1          1          seq
    ...........ucacgaucuauuguucGguuagu.......................................    1          1          seq
    ...........ucacgaucuauuguAcuguuagu.......................................    1          1          seq
    ...........uUacgaucuauuguucuguuagu.......................................    1          1          seq
    ...........ucacgaucuauuguuUuguuagu.......................................    1          1          seq
    ...........ucaUgaucuauuguucuguuagu.......................................    1          1          seq
    ...........ucacgaucuauCguucuguuagu.......................................    1          1          seq
    ...........ucacgaucuGuuguucuguuagu.......................................    1          1          seq
    ...........Ccacgaucuauuguucuguuagu.......................................    4          1          seq
    ...........ucacgGucuauuguucuguuagu.......................................    2          1          seq
    ...........ucacgaucuauuguCcuguuagu.......................................    2          1          seq
    ...........ucacgaucuauuguucugAuagu.......................................    1          1          seq
    ...........Gcacgaucuauuguucuguuagu.......................................    1          1          seq
    ...........ucacgaucuauuguucuguuagC.......................................    95         1          seq
    ...........ucacgaucuauugCucuguuagu.......................................    1          1          seq
    ...........ucaAgaucuauuguucuguuagu.......................................    1          1          seq
    ...........ucacgaucuauuguucuguuagA.......................................    2          1          seq
    ...........ucacgaucuauuguucuguuagu.......................................    368        0          seq
    ...........ucUcgaucuauuguucuguuagu.......................................    1          1          seq
    ...........ucacgaucuauuguucuguuaguu......................................    1          0          seq
    ...........ucacgaucuauuguucuguuaguC......................................    1          1          seq
    ............cacgaucuauuguucuguuagA.......................................    1          1          seq
```

```
           novel-nve-miR-15_guide
                                         novel-nve-miR-15_star
auaguucccggucacgaucuauuguucuguuaguuuaaaacuaacagaacaauagaucgugaccggcgcccgu

............cacgaucuauuguucuguuagu.......................................    5    0    seq
.............acgaucuauuguucuguuagu.......................................    2    0    seq
.............acgaucuauuguucuguuaguC......................................    1    1    seq
........................................cuaacagaacaauagaucgG.............    1    1    seq
.........................................uaacagaacaauagaucgug............    1    0    seq
.........................................uaacagaUcaauagaucguga...........    1    1    seq
.........................................uaacagaacaauagaucguga...........    3    0    seq
.........................................Aaacagaacaauagaucgugacc.........    1    1    seq
```

miRBase precursor : novel-nve-miR-16
Total read count : 333
novel-nve-miR-16_guide read count : 249
novel-nve-miR-16_star read count : 79
remaining reads : 5

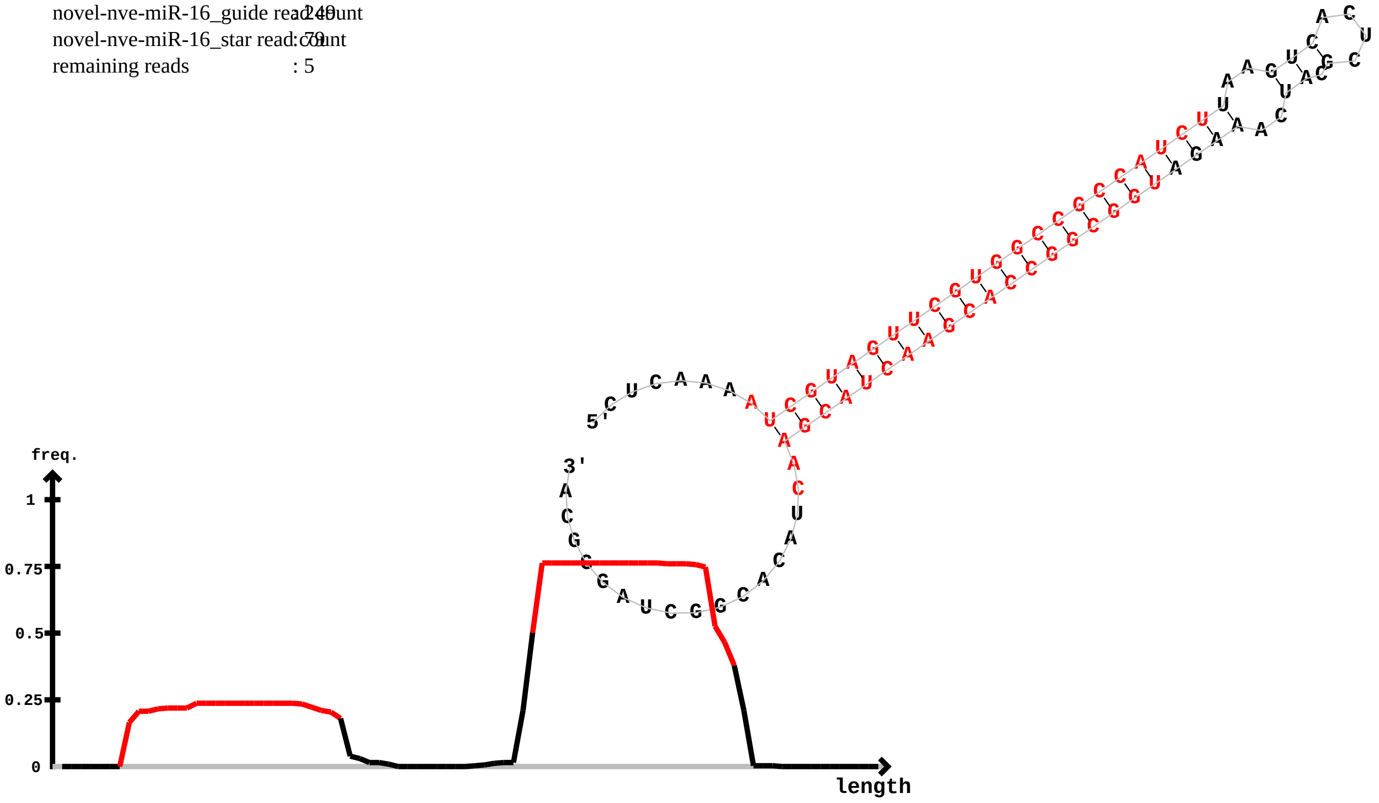


```
                                                  novel-nve-miR-16_guide
        novel-nve-miR-16_star
5'- cucaaaaucguaguucguggccgccaucuuaagucacucgcaucaaagauggcggccacgaacuacgaacuacacggcuagggca   -3'    exp
    .......(((((((((((((((((((((((..(((....).))..)))))))))))))))))))))))........((....)).     reads   mm    sample
    ......aucguaguucguggccgc.............................................................     1       0     seq
    ......aucguaguucguggccgcU............................................................     1       1     seq
    ......aucguaguucguggccgcG............................................................     1       1     seq
    ......aucguaguucguggccgcc............................................................     1       0     seq
    ......aucguaguucguggccgcGa...........................................................     1       1     seq
    ......aucguaguucguggccgccG...........................................................     2       1     seq
    ......aucguaguucguggccgcca...........................................................     1       0     seq
    ......aucguaguucguggccgccau..........................................................     2       0     seq
    ......aucguaguucguggccgccauA.........................................................     2       1     seq
    ......aucguaguucguggccgccauc.........................................................     2       0     seq
    ......aucguaguucguggccgccaucA........................................................     2       1     seq
    ......aucguaguucguggccgccGucu........................................................     1       1     seq
    ......aucguaguucguggccgccaucG........................................................     13      1     seq
    ......aucguaguucguggccgccaucC........................................................     5       1     seq
    ......aucguaguucguUgccgccaucu........................................................     1       1     seq
    ......aucguaguucguggccgccaucu........................................................     16      0     seq
    ......aucguaguucguggcUgccaucu........................................................     1       1     seq
    ......aucguaguucguggccgccaucuu.......................................................     1       0     seq
    ......aucguaguucguggccgccaucuuG......................................................     1       1     seq
    .......ucguaguucguggccgcc............................................................     1       0     seq
    .......ucguaguucguggccgccaCc.........................................................     1       1     seq
    .......ucguaguucguggccgccauU.........................................................     1       1     seq
    .......ucguaguucguggccgccauc.........................................................     2       0     seq
    .......ucguaguucguggccgccaucC........................................................     1       1     seq
    .......ucguaguucguggccgccaucG........................................................     5       1     seq
    .......ucguaguucguggccgccaucu........................................................     2       0     seq
    .......ucguaguucguggccgccaucuG.......................................................     1       1     seq
    .........guaguucguggccgccaucuu.......................................................     1       0     seq
    .........guaguucguggccgccaucuuU......................................................     2       1     seq
    ..........uaguucguggccgccaucuuU......................................................     1       1     seq
    .............uucguggccgccaucuuG......................................................     1       1     seq
    .............uucguggccgccaucuuaag....................................................     2       0     seq
    .............uucguggccgccaucuuaagG...................................................     1       1     seq
    .............uucguggccgccaucuuaagu...................................................     2       0     seq
```

novel-nve-miR-16_guide

novel-nve-miR-16_star

cucaaaaucguaguucguggccgccaucuuaagucacucgcaucaaagauggcggccacgaacuacgaacuacacggcuagggca

| | | | |
|---|---|---|---|
| ..........................................ucaaagauggcggccacgaU....................... | 1 | 1 | seq |
| ...........................................cCaagauggcggccacgaacua.................... | 1 | 1 | seq |
| ............................................Uaagauggcggccacgaacuac................... | 1 | 1 | seq |
| ............................................Uaagauggcggccacgaacuacg.................. | 1 | 1 | seq |
| .............................................aagauggcggccacgaacuaU................... | 1 | 1 | seq |
| ...............................................gauggcggccacgaacuaG................... | 1 | 1 | seq |
| ...............................................gauggcggccacgaacuacg.................. | 27 | 0 | seq |
| ...............................................gauggcggccacgaacuacC.................. | 1 | 1 | seq |
| ...............................................gauggcggUcacgaacuacg.................. | 1 | 1 | seq |
| ...............................................gauggcggccacgaGcuacg.................. | 1 | 1 | seq |
| ...............................................gauggcggccCcgaacuacg.................. | 1 | 1 | seq |
| ...............................................gauggcggccacAaacuacg.................. | 1 | 1 | seq |
| ...............................................gauggcggccacgaacuacA.................. | 1 | 1 | seq |
| ...............................................gauggcggccacgaacuacgG................. | 4 | 1 | seq |
| ...............................................gauggcggccacgCacuacga................. | 1 | 1 | seq |
| ...............................................gauggcggccacgaGcuacga................. | 1 | 1 | seq |
| ...............................................gauggcggccacgaacuacga................. | 6 | 0 | seq |
| ...............................................gauggcggccacgaacuacgaG................ | 1 | 1 | seq |
| ...............................................gauggcggccacgaacuacgaa................ | 8 | 0 | seq |
| ...............................................gauggcggccacgaacuacgaU................ | 1 | 1 | seq |
| ...............................................gauggcggccacgaacuacgaUc............... | 1 | 1 | seq |
| ...............................................gauggcggccacgaacuacgaaU............... | 4 | 1 | seq |
| ...............................................gauggcggcUacgaacuacgaac............... | 2 | 1 | seq |
| ...............................................gauggcggccacgaacuacgaaUu.............. | 2 | 1 | seq |
| ................................................auggcggAcacgaacuacg.................. | 4 | 1 | seq |
| ................................................auggcggccacgaacuacC.................. | 1 | 1 | seq |
| ................................................auggcggccacgaacuacA.................. | 1 | 1 | seq |
| ................................................auggcggccacgaacuacg.................. | 24 | 0 | seq |
| ................................................auggcggccacgaacuacgG................. | 2 | 1 | seq |
| ................................................auggcggccacgaacuacga................. | 3 | 0 | seq |
| ................................................auggcggccaUgaacuacgaa................ | 1 | 1 | seq |
| ................................................auggcggccacgaacuacgaa................ | 7 | 0 | seq |
| ................................................auggcggccacgaacuacgaG................ | 3 | 1 | seq |
| ................................................auggcggccacgaacuacgaU................ | 3 | 1 | seq |
| ................................................aAggcggccacgaacuacgaa................ | 2 | 1 | seq |
| ................................................auggcggccacgaacuacgaUc............... | 1 | 1 | seq |
| ................................................auggcggccacgaacuacgaaU............... | 9 | 1 | seq |
| ................................................auggcggccacgaacuacgaaG............... | 1 | 1 | seq |
| ................................................auggcggccacgaacuacgaac............... | 8 | 0 | seq |
| ................................................auggcgUccacgaacuacgaacu.............. | 1 | 1 | seq |
| ................................................auggcggccacgaacuacgaacu.............. | 5 | 0 | seq |
| ................................................auggcggccacgaacuacgaaUu.............. | 2 | 1 | seq |
| ................................................auggcggccacgaacuacgaacG.............. | 13 | 1 | seq |
| ................................................auggcggccacgaacuacgaacA.............. | 2 | 1 | seq |
| ................................................auggcggccacgaacuacgaacC.............. | 2 | 1 | seq |
| ................................................Guggcggccacgaacuacgaacu.............. | 1 | 1 | seq |
| ................................................auggcggccacgaacuacgaacuacG........... | 1 | 1 | seq |
| .................................................uggcggccacgaacuacg.................. | 8 | 0 | seq |
| .................................................uggcggccacgaacuacA.................. | 1 | 1 | seq |
| .................................................uggcggccacgaaAuacg.................. | 1 | 1 | seq |
| .................................................uggcggccacgaacuacga................. | 3 | 0 | seq |
| .................................................uggcggccacgaacuacgaG................ | 1 | 1 | seq |
| .................................................uggcggccacgaacuacgaU................ | 2 | 1 | seq |
| .................................................uggcggccacgaacuacgaaU............... | 22 | 1 | seq |
| .................................................uggcggccacgaacuacgaUc............... | 2 | 1 | seq |
| .................................................uggcggccacgaacuacgaac............... | 1 | 0 | seq |
| .................................................uggcggccacgaacuacgaaA............... | 2 | 1 | seq |
| .................................................uggcggccacgaacuacgaaG............... | 3 | 1 | seq |
| .................................................uggcggccacgaacuacgaaUu.............. | 2 | 1 | seq |
| .................................................uggcggccacgaacuacgaacA.............. | 1 | 1 | seq |
| .................................................uggcggUcacgaacuacgaacu.............. | 1 | 1 | seq |
| .................................................uggcggccacgaacuacgaacC.............. | 2 | 1 | seq |
| .................................................uggcggccacgaacuacgaacu.............. | 15 | 0 | seq |
| .................................................uggcggccacgaacuacgaacG.............. | 20 | 1 | seq |

miRBase precursor : novel-nve-miR-18
Total read count : 233
novel-nve-miR-18_guide read count : 224
novel-nve-miR-18_star read count : 9
remaining reads : 0

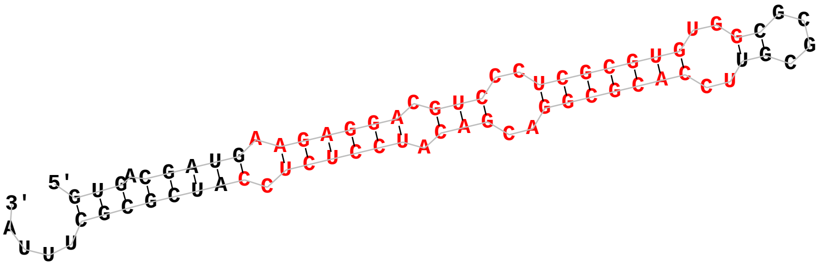


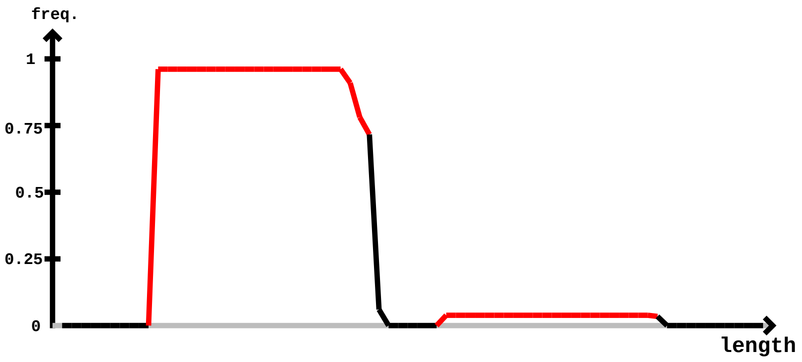


```
                                                novel-nve-miR-18_star
          novel-nve-miR-18_guide
5'- gugacgaugaagaggacgucccucgcgugugcgcgcguuccacgcggacgacauccucuccaucgcguuua   -3'    exp
    (((.(((((.((((((.(((..(((((((..((....))..)))))))..))).)))))).)))))))).... reads   mm   sample
    .........aagaggacgucccucgcguA...........................................  1       1    seq
    .........aagaggacgucccucgUgug...........................................  1       1    seq
    .........aagaggacgucccucgcgug...........................................  10      0    seq
    .........aagaggacgucccucgcgugu..........................................  21      0    seq
    .........Gagaggacgucccucgcgugu..........................................  2       1    seq
    .........aagaggacgucccucgcgugC..........................................  3       1    seq
    .........aagaggacgucccucgcAugu..........................................  1       1    seq
    .........aagaggacgCcccucgcgugu..........................................  1       1    seq
    .........aagagAacgucccucgcgugu..........................................  1       1    seq
    .........aagaggacgucccucgcgugA..........................................  1       1    seq
    .........aagaggacgucccucgcgugug.........................................  14      0    seq
    .........aagaggacgucccucgcguguA.........................................  1       1    seq
    .........aagaggacgucccucgcgugugA........................................  17      1    seq
    .........aagaggaUgucccucgcgugugg........................................  1       1    seq
    .........aagaggacgucccucCcgugugg........................................  1       1    seq
    .........aagaggacguccUucgcgugugg........................................  1       1    seq
    .........aagaggacgCcccucgcgugugg........................................  1       1    seq
    .........aagaggacAucccucgcgugugg........................................  3       1    seq
    .........aagaggacgucccCcgcgugugg........................................  1       1    seq
    .........aagaggacgucccuUgcgugugg........................................  1       1    seq
    .........aaCaggacgucccucgcgugugg........................................  1       1    seq
    .........aagaggacgucccucgcgugugg........................................  116     0    seq
    .........aagaggacgucccucgcgugugC........................................  2       1    seq
    .........aagaggacgucccucgcgugugU........................................  7       1    seq
    .........aagaggaAgucccucgcgugugg........................................  1       1    seq
    .........aagaggacgucccucgcguguggU.......................................  9       1    seq
    .........aagaggacgucccucgcguguggc.......................................  1       0    seq
    .........aagaggacgucccucgcguguggA.......................................  4       1    seq
    .......................................uccacgcggacgGcauccucuc...........  1       1    seq
    .......................................uccacgcggacgacauccucucc..........  5       0    seq
    .......................................uccacgcggacgacauccucucA..........  1       1    seq
    .......................................uccaUgcggacgacauccucucc..........  1       1    seq
    .......................................uccacgcggacgacauccucucU..........  1       1    seq
```

miRBase precursor : novel-nve-miR-19
Total read count : 1469
novel-nve-miR-19_guide read count : 1374
novel-nve-miR-19_star read count : 82
remaining reads : 13

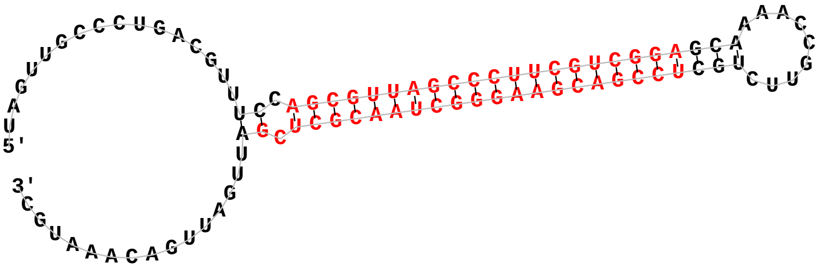

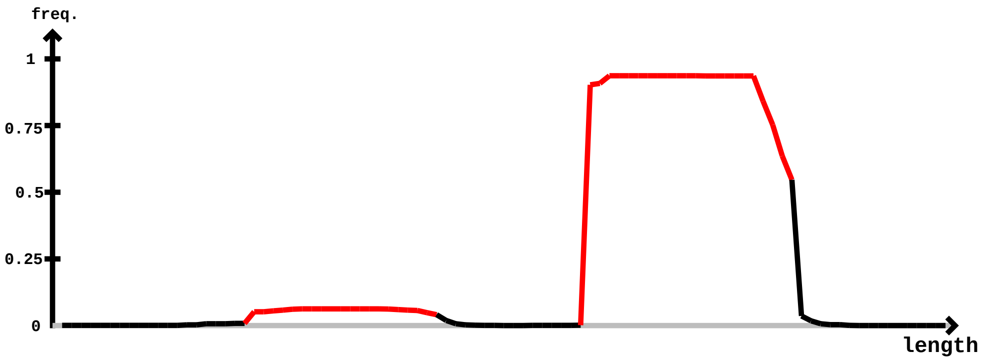


novel-nve-miR-19_star

novel-nve-miR-19_guide

```
5'- uaguugcccugacguuuccagcguuagcccuucgucggagcaaaaccguucugcuccgacgaagggcuaacgcucgauugauugacaaaugc  -3'   exp
    ..(((.....)))...((.(((((((((((((((((((((((.........))))))))))))))))))))))).))...............   reads   mm   sample
    uGguugcccugacguuuc..........................................................................   1       1    seq
    ............cguuucGagcguuagcccuuc...........................................................   1       1    seq
    ............cguuucGagcguuagcccuucg..........................................................   2       1    seq
    ..............uuucGagcguuagcccuucg..........................................................   1       1    seq
    ..............uuucGagcguuagcccuucgu.........................................................   3       1    seq
    ..............uuucGagcguuagcccuucguc........................................................   2       1    seq
    .................cGagcguuagcccuucgucg.......................................................   1       1    seq
    .................cGagcguuagcccuucgucgga.....................................................   1       1    seq
    ..................Gagcguuagcccuucgucgg......................................................   1       1    seq
    ...................agcguuagcccuucgucg.......................................................   1       0    seq
    ...................agcguuGgcccuucgucg.......................................................   10      1    seq
    ...................agcguuagcccuucgucgg......................................................   2       0    seq
    ...................agcguuGgcccuucgucgg......................................................   8       1    seq
    ...................agcguuagcccuucgucggG.....................................................   2       1    seq
    ...................agcguuGgcccuucgucgga.....................................................   25      1    seq
    ...................agcguuagcccuucgucgga.....................................................   3       0    seq
    ...................agcguuGgcccuucgucggag....................................................   9       1    seq
    ...................agcguuagcccuucgucggaA....................................................   2       1    seq
    ...................agcguuagcccuucgucggaUc...................................................   1       1    seq
    ...................agcguuagcccuucgucggagUa..................................................   1       1    seq
    .....................cguuGgcccuucgucgga.....................................................   1       1    seq
    .....................cguuagcccuucgucggag....................................................   2       0    seq
    .....................cguuagccUuucgucggag....................................................   1       1    seq
    .....................cguuGgcccuucgucggag....................................................   1       1    seq
    ......................guuagcccuucgucggag....................................................   3       0    seq
    ......................guuagcccuucgucggagA...................................................   1       1    seq
    .......................uuagcccuucgucggagU...................................................   1       1    seq
    .......................uuagcccuucgucggagG...................................................   3       1    seq
    .......................uuGgcccuucgucggagca..................................................   1       1    seq
    ........................uagcccuucgucggagcaa.................................................   1       0    seq
    ........................uGgcccuucgucggagcaaaa...............................................   1       1    seq
    ................................................uuUugcuccgacgaaggg..........................   1       1    seq
    .....................................................cuccgacgaagggcuaacgG...................   1       1    seq
    ......................................................uccgacgaGgggcuaacg....................   1       1    seq
```

```
                   novel-nve-miR-19_star
                                                      novel-nve-miR-19_guide
uaguugcccugacguuuccagcguuagcccuucgucggagcaaaaccguucugcuccgacgaagggcuaacgcucgauugauugacaaaugc

......................................................uccgacgaagggcuaGcg....................    15     1     seq
......................................................uccgacgaagggcuaacC....................    2      1     seq
......................................................uccgacgaagggcuCacg....................    1      1     seq
......................................................uccgacgaagggcuaacA....................    9      1     seq
......................................................ucGgacgaagggcuaacg....................    1      1     seq
......................................................uccgacgaagggcuaacg....................    58     0     seq
......................................................uccgacgaagggcuGacg....................    36     1     seq
......................................................uccgacgaagggcuaacU....................    16     1     seq
......................................................uccgacgaagggcuaacgc...................    11     0     seq
......................................................uccgacgaagggcuGacgc...................    3      1     seq
......................................................uccgaAgaagggcuaacgc...................    1      1     seq
......................................................uccgacgaagggcuaacUc...................    1      1     seq
......................................................uccgacgaagggcuaacgG...................    31     1     seq
......................................................uccgacgaagggcuaacAc...................    1      1     seq
......................................................uccgacgGagggcuaacgc...................    1      1     seq
......................................................uccgacgaagggcuaacgU...................    8      1     seq
......................................................uccgacgaagggcuaacgA...................    69     1     seq
......................................................uccgacgaagggcuaacgcG..................    30     1     seq
......................................................uccgacgaagggcuaaAgcu..................    1      1     seq
......................................................uccgacgaagggcuaaUgcu..................    1      1     seq
......................................................uccgacgaagggcuaacgcA..................    10     1     seq
......................................................uccgacgaagggcuaGcgcu..................    6      1     seq
......................................................uccgacgaagggcuGacgcu..................    11     1     seq
......................................................uccgacgGagggcuaacgcu..................    1      1     seq
......................................................uccgacgaagggcuaacgAu..................    52     1     seq
......................................................uccgacgaagggcuaacCcu..................    1      1     seq
......................................................uccgacgaagggcuaacgcC..................    7      1     seq
......................................................Cccgacgaagggcuaacgcu..................    1      1     seq
......................................................uccgacgaagggcuaacgcu..................    50     0     seq
......................................................uccgacgaagggcuaacgcuG.................    16     1     seq
......................................................uccgacgaagggcuaacgcuc.................    33     0     seq
......................................................uccgacgGagggcuaacgcuc.................    1      1     seq
......................................................uccgacgaGgggcuaacgcuc.................    2      1     seq
......................................................uccgacgaagggcuGacgcuc.................    11     1     seq
......................................................uccgGcgaagggcuaacgcuc.................    4      1     seq
......................................................uccgacgaagggcuaacgcuU.................    21     1     seq
......................................................uccgacgaagggcuaCcgcuc.................    1      1     seq
......................................................uccgacgaagggcuaacgcuA.................    25     1     seq
......................................................uccgacgaagggcuaGcgcuc.................    14     1     seq
......................................................uAcgacgaagggcuaacgcuc.................    1      1     seq
......................................................uccgacgaagggcuaacgcuAg................    12     1     seq
......................................................uccgacgaagggcuaacAcucg................    2      1     seq
......................................................uccgacgaagggcuaUcgcucg................    3      1     seq
......................................................uccgacgaagggcuaacgcucC................    8      1     seq
......................................................Cccgacgaagggcuaacgcucg................    3      1     seq
......................................................uccgaAgaagggcuaacgcucg................    1      1     seq
......................................................uUcgacgaagggcuaacgcucg................    1      1     seq
......................................................uccgacgaagggcCaacgcucg................    2      1     seq
......................................................uccgacgaUgggcuaacgcucg................    1      1     seq
......................................................uccgacgaagggcuaacgcAcg................    13     1     seq
......................................................Gccgacgaagggcuaacgcucg................    1      1     seq
......................................................Accgacgaagggcuaacgcucg................    6      1     seq
......................................................uccAacgaagggcuaacgcucg................    1      1     seq
......................................................uccgacgGagggcuaacgcucg................    1      1     seq
......................................................uccgacgaagggcuaacgcuUg................    4      1     seq
......................................................uccgacgaGgggcuaacgcucg................    2      1     seq
......................................................ucUgacgaagggcuaacgcucg................    2      1     seq
......................................................uAcgacgaagggcuaacgcucg................    1      1     seq
......................................................ucAgacgaagggcuaacgcucg................    2      1     seq
......................................................uccgacgaagggcuaacgcucU................    22     1     seq
......................................................uccgacgaagggcuaacgcucA................    86     1     seq
......................................................uccgacgaagggcuGacgcucg................    92     1     seq
......................................................uccgacgaagggUuaacgcucg................    1      1     seq
......................................................uccgacgaagggcuaaGgcucg................    1      1     seq
......................................................uccgacgaagggcuaacgcucg................    315    0     seq
......................................................uccgacgaagggcuaGcgcucg................    117    1     seq
......................................................uccgGcgaagggcuaacgcucg................    2      1     seq
......................................................uccgacgaagggcuaacgUucg................    1      1     seq
......................................................uccgUcgaagggcuaacgcucg................    2      1     seq
```

```
                   novel-nve-miR-19_star
                                                      novel-nve-miR-19_guide
uaguugcccugacguuuccagcguuagcccuucgucggagcaaaaccguucugcuccgacgaagggcuaacgcucgauugauugacaaaugc

......................................................uccCacgaagggcuaacgcucg................    1     1     seq
......................................................uccgacgaagggcuGacgcucga...............    2     1     seq
......................................................uccgacgaGgggcuaacgcucga...............    3     1     seq
......................................................uccgacgaagggcuaacgcucgU...............    3     1     seq
......................................................uccgacgaagggcuaacgcucgG...............    3     1     seq
......................................................uccgacgaagggcuaacgcucUa...............    1     1     seq
......................................................uccgacgaagggcuaacgcucga...............    11    0     seq
......................................................uccgacgaagggcuaacgcucgC...............    1     1     seq
......................................................uccgacgaagggcuaacgcucAa...............    3     1     seq
......................................................uccgacgaagggcuGacgcucgau..............    2     1     seq
......................................................uccgacgaagggcuaacgcucgau..............    6     0     seq
......................................................uccgacgaagggcuaacgcucgaG..............    4     1     seq
......................................................uccgacgaagggcuaacgcucgaC..............    4     1     seq
......................................................uccgacgaagggcuGacgcucgauu.............    2     1     seq
......................................................uccgacgaagggcuaacgcucgauA.............    1     1     seq
......................................................uccgacgaagggcuaacgcucgauu.............    1     0     seq
......................................................uccgacgaagggcuaacgcucgUuu.............    1     1     seq
......................................................uccgacgaagggcuaacgcucgauuAa...........    2     1     seq
......................................................uccgacgaagggcuaacgcucgauuUa...........    2     1     seq
......................................................uccgacgaagggcuaacgcucgauuUau..........    1     1     seq
.......................................................ccgacgaagggcuaacgA...................    3     1     seq
.......................................................ccgacgaagggcuaacgcucA................    1     1     seq
.......................................................ccgacgaagggcuaacgcucg................    1     0     seq
.......................................................ccgacgaagggcuaGcgcucg................    3     1     seq
........................................................cgacgaagggcuaacgcu..................    1     0     seq
........................................................cgacgaagggcuaacgcuU.................    2     1     seq
........................................................cgaAgaagggcuaacgcucg................    2     1     seq
........................................................cgacgaagggcuaacgcucg................    24    0     seq
........................................................cgacgaagggcuaGcgcucg................    1     1     seq
........................................................cgacgaaAggcuaacgcucg................    1     1     seq
........................................................Agacgaagggcuaacgcucg................    5     1     seq
........................................................cgacgaagggcuUacgcucg................    1     1     seq
........................................................cgacgaagggcuGacgcucg................    3     1     seq
........................................................cgacgaagggcuaacgcucA................    2     1     seq
```

miRBase precursor : novel-nve-miR-20
Total read count : 929
novel-nve-miR-20_guide read count : 926
novel-nve-miR-20_star read count : 0
remaining reads : 3

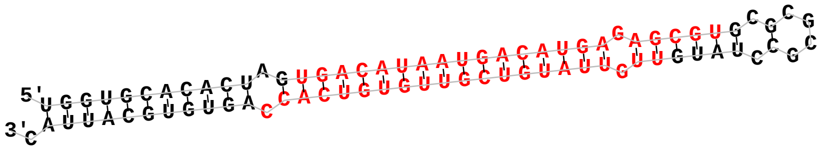

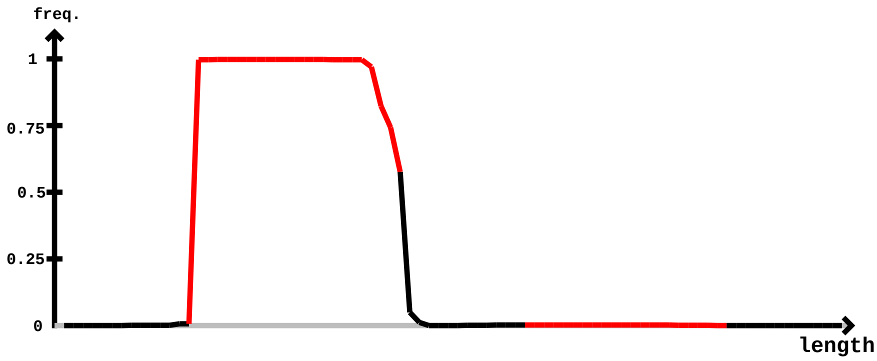


```
                 novel-nve-miR-20_guide
                                                  novel-nve-miR-20_star
5'- uggugcacacuagugacauaaugacaugagagcgugcgcgcgccuauguuguuaugucguugugucaccagugugcauuac  -3'   exp
    (((((((((((.(((((((((((((((((.((((((.(....).)))))).))))))))))))))))).))))))))))).   reads   mm   sample
    ......acacuagugacaCaaugacau......................................................   1       1    seq
    ...........agugacauCaugacaugagag.................................................   1       1    seq
    ...........agugacauaaugacaugagaA.................................................   2       1    seq
    ...........agugacauaaugacaugagag.................................................   2       0    seq
    .............ugacauaaugacaugaga..................................................   23      0    seq
    .............ugGcauaaugacaugaga..................................................   1       1    seq
    .............Agacauaaugacaugaga..................................................   1       1    seq
    .............Ggacauaaugacaugagag.................................................   1       1    seq
    .............ugacauaaugacaugagag.................................................   104     0    seq
    .............Agacauaaugacaugagag.................................................   1       1    seq
    .............ugacauaaAgacaugagag.................................................   1       1    seq
    .............ugacauaaugacaugagGg.................................................   2       1    seq
    .............ugacauaauCacaugagag.................................................   1       1    seq
    .............ugacauaaugacGugagag.................................................   1       1    seq
    .............ugacauaaugacaugagaU.................................................   2       1    seq
    .............ugacauaaugacaugaAag.................................................   1       1    seq
    .............ugacauaaugacaugagaA.................................................   14      1    seq
    .............Cgacauaaugacaugagag.................................................   1       1    seq
    .............ugGcauaaugacaugagag.................................................   2       1    seq
    .............ugacauaaugGcaugagagc................................................   1       1    seq
    .............ugacaAaaugacaugagagc................................................   1       1    seq
    .............Agacauaaugacaugagagc................................................   2       1    seq
    .............ugacauaaugacaugagagU................................................   16      1    seq
    .............ugacauaaugacaugUgagc................................................   2       1    seq
    .............ugacauaaugacaugagagA................................................   4       1    seq
    .............ugacauaaugacaugagagc................................................   49      0    seq
    .............Agacauaaugacaugagagcg...............................................   4       1    seq
    .............uAacauaaugacaugagagcg...............................................   1       1    seq
    .............uCacauaaugacaugagagcg...............................................   1       1    seq
    .............ugacauaaugacaCgagagcg...............................................   1       1    seq
    .............Ggacauaaugacaugagagcg...............................................   2       1    seq
    .............ugacauaaugacaugagagcU...............................................   9       1    seq
    .............ugacauaaugacaugagagcC...............................................   2       1    seq
    .............ugacauaaugacaugagagcA...............................................   33      1    seq
```

```
             novel-nve-miR-20_guide
                                   novel-nve-miR-20_star
uggugcacacuagugacauaaugacaugagagcgugcgcgccuauguuguuaugucguugugucaccagugugcauuac

.............ugacGuaaugacaugagagcg.............................................    1     1     seq
.............ugacauaaugacaugagagUg.............................................    1     1     seq
.............ugUcauaaugacaugagagcg.............................................    1     1     seq
.............ugacauaaugacaugagagcg.............................................    99    0     seq
.............ugacauaaugacaugagaUcgu............................................    3     1     seq
.............ugaAauaaugacaugagagcgu............................................    1     1     seq
.............Ggacauaaugacaugagagcgu............................................    1     1     seq
.............ugacauaaugacaugagagcgA............................................    7     1     seq
.............ugacGuaaugacaugagagcgu............................................    1     1     seq
.............ugacauaaAgacaugagagcgu............................................    1     1     seq
.............ugacauaGugacaugagagcgu............................................    1     1     seq
.............ugCcauaaugacaugagagcgu............................................    1     1     seq
.............ugUcauaaugacaugagagcgu............................................    1     1     seq
.............ugacauaaugaUaugagagcgu............................................    1     1     seq
.............ugacauaaugacaugagagcgC............................................    65    1     seq
.............ugacauaaugacaugagagUgu............................................    1     1     seq
.............ugacauaaugacaugagagcAu............................................    2     1     seq
.............ugacauaaugacaugGgagcgu............................................    2     1     seq
.............Agacauaaugacaugagagcgu............................................    6     1     seq
.............uAacauaaugacaugagagcgu............................................    1     1     seq
.............ugacauaaugacaugagagcgu............................................    359   0     seq
.............ugacauaaugacaugagagcCu............................................    1     1     seq
.............ugacUuaaugacaugagagcgu............................................    2     1     seq
.............ugacauaaCgacaugagagcgu............................................    1     1     seq
.............Cgacauaaugacaugagagcgu............................................    2     1     seq
.............ugacauaaugCcaugagagcgu............................................    1     1     seq
.............ugGcauaaugacaugagagcgu............................................    2     1     seq
.............ugacauaaugacaGgagagcgu............................................    1     1     seq
.............ugacauaaugacauAagagcgu............................................    2     1     seq
.............ugacauaaugaAaugagagcgu............................................    1     1     seq
.............ugacauaaugGcaugagagcgu............................................    3     1     seq
.............ugacauaaugacGugagagcgu............................................    4     1     seq
.............ugacauaaugacaugagagcgG............................................    14    1     seq
.............ugacauaaugacaugagagcguC...........................................    5     1     seq
.............ugacauaaugacaugagagcguU...........................................    28    1     seq
.............ugacauaaugacaugagagcguA...........................................    2     1     seq
.............ugacauaaugacaugagagcguAc..........................................    2     1     seq
.............ugacauaaugacaugagagcguUc..........................................    9     1     seq
...............acauaaugacaugagagcgu............................................    1     0     seq
.......................................gccuauguuguuaugucguugu..................    1     0     seq
..........................................uauguuguCaugucguuguguca..............    1     1     seq
```

miRBase precursor : novel-nve-miR-23
Total read count : 18943
novel-nve-miR-23_guide read count : 18938
novel-nve-miR-23_star read count : 5
remaining reads : 0

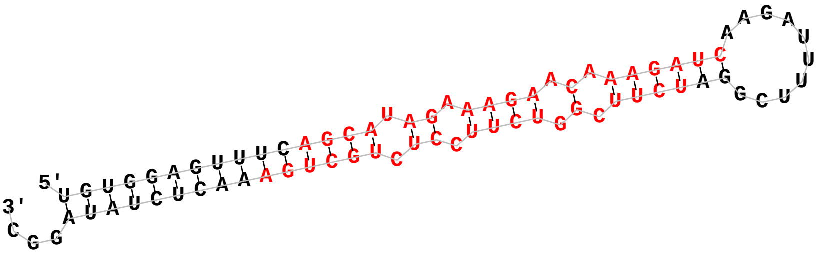


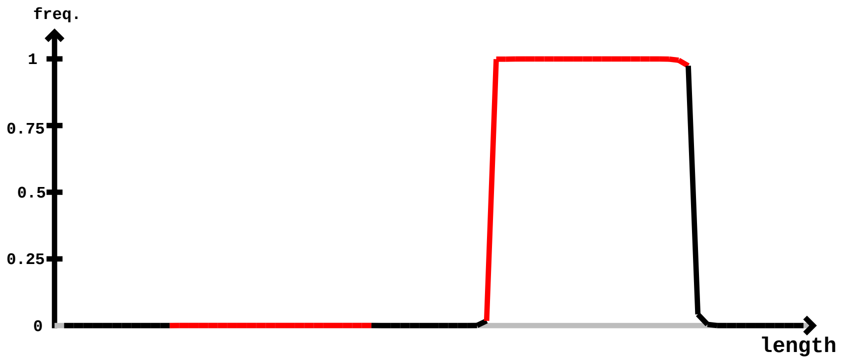


```
                novel-nve-miR-23_star
                                          novel-nve-miR-23_guide
5'- uguggaguuucagcauagaaagaacaaagaucaagauuuucggaucuucggucuuccucugcugaaacucuauaggc   -3'    exp
    ((((((((((((((((.((.((((.(.((((((..........)))))).).)))).)).)))))))))))))))...    reads    mm    sample
    ...........agcauagaaagaacaaagauU.............................................    1        1     seq
    ...........agcauagaaagaacaaagauc.............................................    4        0     seq
    ..........................................Aaucuucggucuuccucugcuga............    5        1     seq
    ...........................................aucuucggucuuccucug................    1        0     seq
    ...........................................aucuucggucuuccucugcA..............    1        1     seq
    ...........................................aucuucggucuuccucugcu..............    12       0     seq
    ...........................................aucuucggucuuccucugcC..............    4        1     seq
    ...........................................Uucuucggucuuccucugcu..............    1        1     seq
    ...........................................aucuucggucuuccucugcug.............    1        0     seq
    ...........................................Gucuucggucuuccucugcuga............    7        1     seq
    ...........................................aucuucggucuucUucugcuga............    1        1     seq
    ...........................................aucuucggucCuccucugcuga............    2        1     seq
    ...........................................aucuucggucuuccucugcugG............    6        1     seq
    ...........................................Cucuucggucuuccucugcuga............    1        1     seq
    ...........................................aucuucggucuuccucCgcuga............    2        1     seq
    ...........................................aucuucggucuCccucugcuga............    1        1     seq
    ...........................................aucuucggCcuuccucugcuga............    4        1     seq
    ...........................................aucuucAgucuuccucugcuga............    1        1     seq
    ...........................................aucuucggucuuccucugcuga............    258      0     seq
    ...........................................aucuCcggucuuccucugcuga............    1        1     seq
    ...........................................aucuucggucuuccucGgcuga............    2        1     seq
    ...........................................aucuucggucuuccucugcAga............    1        1     seq
    ...........................................aucCucggucuuccucugcuga............    2        1     seq
    ...........................................aucuucggucuuccGcugcuga............    1        1     seq
    ...........................................aucuucggucuuccCcugcuga............    2        1     seq
    ...........................................aucuucggucuuccucAgcuga............    2        1     seq
    ...........................................aucuucggucuuccAcugcuga............    1        1     seq
    ...........................................aAcuucggucuuccucugcuga............    2        1     seq
    ...........................................aucAucggucuuccucugcuga............    3        1     seq
    ...........................................aucuucggucuuccucugcugU............    10       1     seq
    ...........................................aucuucggucuuccucugcugaU...........    1        1     seq
    ............................................ucuucggucuuccucugc...............    3        0     seq
    ............................................ucuucggucuuccucugU...............    7        1     seq
    ............................................ucuucggucuuccucugA...............    1        1     seq
```

```
           novel-nve-miR-23_star
                                                novel-nve-miR-23_guide
uguggaguuucagcauagaaagaacaaagaucaagauuuucggaucuucggucuuccucugcugaaacucuauaggc

............................................ucuucggucuuccucugcG..............    1     1     seq
............................................ucuucggucuuccucCgcu..............    1     1     seq
............................................ucuucggucuuccucugcu..............    41    0     seq
............................................ucuucggucuuccucugcA..............    8     1     seq
............................................ucuucggucuuccucugcC..............    4     1     seq
............................................ucuuAggucuuccucugcu..............    1     1     seq
............................................ucuucggucuuccucugAu..............    5     1     seq
............................................ucuucgCucuuccucugcu..............    1     1     seq
............................................ucuCcggucuuccucugcug.............    3     1     seq
............................................ucuucggucuuccucugcuA.............    34    1     seq
............................................Ccuucggucuuccucugcug.............    3     1     seq
............................................ucuGcggucuuccucugcug.............    1     1     seq
............................................ucuucggAcuuccucugcug.............    1     1     seq
............................................ucuucggucuuccucugcug.............    319   0     seq
............................................ucuucggucuuAcucugcug.............    2     1     seq
............................................ucuucggucuAccucugcug.............    2     1     seq
............................................ucuucggucuuccucugcuU.............    12    1     seq
............................................ucuucggCcuuccucugcug.............    3     1     seq
............................................ucuucggucuuccucugAug.............    1     1     seq
............................................ucuucggucuuUcucugcug.............    1     1     seq
............................................ucuucggucuCccucugcug.............    1     1     seq
............................................ucuucggucuuccucugcCg.............    2     1     seq
............................................ucuucggucuuccAcugcug.............    1     1     seq
............................................Acuucggucuuccucugcug.............    2     1     seq
............................................ucuucggucCuccucugcug.............    1     1     seq
............................................ucAucggucuuccucugcug.............    2     1     seq
............................................ucuucggucuuccucugcuC.............    1     1     seq
............................................Gcuucggucuuccucugcug.............    1     1     seq
............................................ucCucggucuuccucugcuga............    72    1     seq
............................................ucuucggucuAccucugcuga............    19    1     seq
............................................ucuuAggucuuccucugcuga............    6     1     seq
............................................ucuucggucuuccucugGuga............    6     1     seq
............................................ucuucggucuucUucugcuga............    22    1     seq
............................................ucuucgUucuuccucugcuga............    6     1     seq
............................................ucuucggucuuccucugcugG............    186   1     seq
............................................ucuucgguAuuccucugcuga............    7     1     seq
............................................ucuucggucuuccucugcuCa............    6     1     seq
............................................ucuucgguUuuccucugcuga............    15    1     seq
............................................ucuucggucuuUcucugcuga............    20    1     seq
............................................ucuucAgucuuccucugcuga............    24    1     seq
............................................ucuucggucAuccucugcuga............    16    1     seq
............................................ucGucggucuuccucugcuga............    4     1     seq
............................................ucuucggCcuuccucugcuga............    60    1     seq
............................................uAuucggucuuccucugcuga............    12    1     seq
............................................ucuucggucuCccucugcuga............    69    1     seq
............................................ucuuUggucuuccucugcuga............    28    1     seq
............................................ucuucgCucuuccucugcuga............    6     1     seq
............................................ucuucggucuuccucugAuga............    5     1     seq
............................................uUuucggucuuccucugcuga............    32    1     seq
............................................ucuucggucuuccuUugcuga............    4     1     seq
............................................ucuGcggucuuccucugcuga............    9     1     seq
............................................ucuucggucuuccucugcuUa............    4     1     seq
............................................ucuucggucuuccCcugcuga............    79    1     seq
............................................ucuucggucuuccucugcGga............    6     1     seq
............................................ucuucggucuGccucugcuga............    1     1     seq
............................................Ccuucggucuuccucugcuga............    68    1     seq
............................................ucuucggucuuccucugcugC............    63    1     seq
............................................Gcuucggucuuccucugcuga............    26    1     seq
............................................ucuucgguGuuccucugcuga............    4     1     seq
............................................ucuucggucuuccuAugcuga............    5     1     seq
............................................ucuucgAucuuccucugcuga............    10    1     seq
............................................ucuucUgucuuccucugcuga............    9     1     seq
............................................ucuCcggucuuccucugcuga............    52    1     seq
............................................ucuucggucuuccucGgcuga............    8     1     seq
............................................Acuucggucuuccucugcuga............    133   1     seq
............................................ucuucggGcuuccucugcuga............    6     1     seq
............................................ucuucggucuuAcucugcuga............    11    1     seq
............................................ucuucggucuuccucugcuAa............    13    1     seq
............................................uGuucggucuuccucugcuga............    1     1     seq
```

```
           novel-nve-miR-23_star
                                 novel-nve-miR-23_guide
uguggaguuucagcauagaaagaacaaagaucaagauuuucggaucuucggucuuccucugcugaaacucuauaggc

............................................ucuucggucGuccucugcuga............    5       1       seq
............................................ucuuGggucuuccucugcuga............    2       1       seq
............................................ucuAcggucuuccucugcuga............    28      1       seq
............................................ucuucggAcuuccucugcuga............    19      1       seq
............................................ucuucggucuuccGcugcuga............    3       1       seq
............................................ucuucggucuuGcucugcuga............    3       1       seq
............................................ucuucggucuuccucugUuga............    50      1       seq
............................................ucuucggucuuccucugcAga............    9       1       seq
............................................ucuucggucuuccucCgcuga............    54      1       seq
............................................ucuucggucuuccucuUcuga............    6       1       seq
............................................ucuucCgucuuccucugcuga............    9       1       seq
............................................ucuucggucuucGucugcuga............    4       1       seq
............................................ucuucggucuuccAcugcuga............    16      1       seq
............................................ucAucggucuuccucugcuga............    67      1       seq
............................................ucuucggucuuccucuAcuga............    16      1       seq
............................................ucuucggucuuccucugcCga............    43      1       seq
............................................ucuucggucuuccucugcugU............    420     1       seq
............................................ucuucggucCuccucugcuga............    78      1       seq
............................................ucuucggucuuccucuCcuga............    5       1       seq
............................................ucuucggucuuccucugcuga............    15325   0       seq
............................................ucuucggucuuccucAgcuga............    23      1       seq
............................................ucuucggucuucAucugcuga............    9       1       seq
............................................ucuucggucuuccCcugcugaa...........    2       1       seq
............................................ucuucAgucuuccucugcugaa...........    1       1       seq
............................................ucuucggucuucUucugcugaa...........    2       1       seq
............................................uUuucggucuuccucugcugaa...........    1       1       seq
............................................ucuucggCcuuccucugcugaa...........    1       1       seq
............................................ucuucggucuuccucugUugaa...........    3       1       seq
............................................ucuAcggucuuccucugcugaa...........    1       1       seq
............................................ucuucggucuuccucugcugUa...........    4       1       seq
............................................ucuucggucuuccucugcugaC...........    14      1       seq
............................................ucuucggucuuccAcugcugaa...........    1       1       seq
............................................ucuucggucuuccucuAcugaa...........    1       1       seq
............................................ucuucggAcuuccucugcugaa...........    1       1       seq
............................................ucuucggucuuccucugcugaU...........    184     1       seq
............................................ucAucggucuuccucugcugaa...........    1       1       seq
............................................ucuucggucuuccucugcugaG...........    6       1       seq
............................................Acuucggucuuccucugcugaa...........    3       1       seq
............................................ucuucggucCuccucugcugaa...........    3       1       seq
............................................ucuuUggucuuccucugcugaa...........    1       1       seq
............................................ucuucggucuuccucugcugaa...........    478     0       seq
............................................ucuucgguUuuccucugcugaa...........    1       1       seq
............................................Ccuucggucuuccucugcugaa...........    3       1       seq
............................................ucuucggucuuccucCgcugaa...........    1       1       seq
............................................ucuucggucuuccucugcugaaa..........    3       0       seq
............................................ucuucggucuuccucugcugaaC..........    4       1       seq
............................................ucuucggucuuccucugcugaaG..........    1       1       seq
............................................ucuucggucuuccucugcugaUa..........    14      1       seq
............................................ucuucggucuuccucugcugaaU..........    59      1       seq
............................................ucuucggucuuccucugcugaUac.........    1       1       seq
.............................................cuucggucuuccucugcuga............    1       0       seq
..............................................uucggucuuccucugcuga............    8       0       seq
..............................................uucggucuuccucugcugaaU..........    2       1       seq
...............................................ucggucuuccucugcuga............    3       0       seq
```

miRBase precursor : novel-nve-miR-24
Total read count : 368
novel-nve-miR-24_guide read count : 295
novel-nve-miR-24_star read count : 54
remaining reads : 19

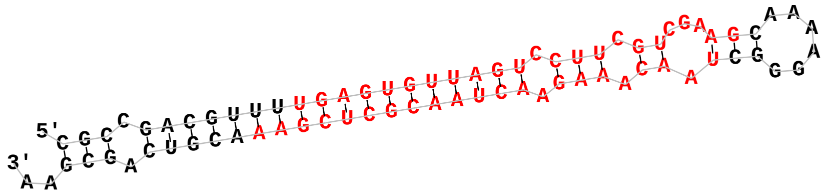

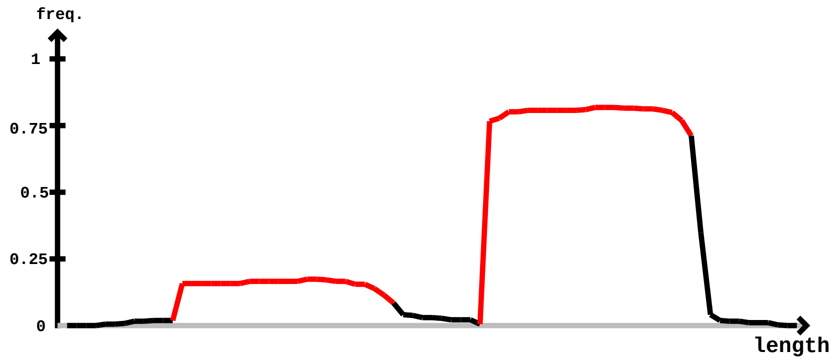


```
                 novel-nve-miR-24_star
                                              novel-nve-miR-24_guide
5'- cgccgacguuuugaguguuaguccuucgucgaagcaaaagggcuaacaaagaacuaacgcucgaaacgucagcgaa  -3'    exp
    (((.((((((((((((((((((.(((.((...(((......))).)).))).))))))))))))))))))).)))..   reads      mm         sample
    ...cgacguuuugaguguuagucA....................................................   1          1          seq
    ...cgacguuuugaguguuagucU....................................................   1          1          seq
    .....acguuuugaguguuagucc....................................................   1          0          seq
    ......cguuuugaguguuagucUuu..................................................   1          1          seq
    ......cguuuugaguguuaguccuuU.................................................   1          1          seq
    ......cguuuugaguguuaguccuuc.................................................   1          0          seq
    ........uuuugaguguuaguccuucgu...............................................   1          0          seq
    ...........ugaguguuaguccuucgu...............................................   2          0          seq
    ...........ugagAguuaguccuucgu...............................................   1          1          seq
    ...........ugaguguuaguccuucgucg.............................................   3          0          seq
    ...........ugaguguuaguccuucgucA.............................................   2          1          seq
    ...........ugaguguuaguccCucgucg.............................................   1          1          seq
    ...........ugaguguuaguccuucgucga............................................   9          0          seq
    ...........ugaguguuaguccuucgucgaC...........................................   1          1          seq
    ...........ugaguguuaguccuucgucgaa...........................................   8          0          seq
    ...........ugaguguuaguccuucgucgaG...........................................   1          1          seq
    ...........ugaguguuaguccuucgucgaU...........................................   1          1          seq
    ...........ugaguguuaguccuucgucgaag..........................................   8          0          seq
    ...........ugaguguuaguccuucgucgaaA..........................................   1          1          seq
    ...........ugaguguuaguccuucgucgaaU..........................................   5          1          seq
    ...........ugagugCuaguccuucgucgaag..........................................   1          1          seq
    ...........ugaguguuaguccuucgucgaaC..........................................   1          1          seq
    ...........ugaguguuaguccuucgucgaagc.........................................   1          0          seq
    ...........ugaguguuaguccuucgucgaagU.........................................   1          1          seq
    ...........ugaguguuaguccuucgucgaagcU........................................   3          1          seq
    ...........ugaguguuaguccuucgucgaagca........................................   1          0          seq
    .................uagCccuucgucgaagcaaa.......................................   1          1          seq
    .................uagCccuucgucgaagcaaaa......................................   2          1          seq
    ........................uucgucgaagcaaaaggg..................................   5          0          seq
    ........................uucgucgaagcaaaaggA..................................   1          1          seq
    ...................................aaaagggcuaacaaagaacuaU...................   1          1          seq
    ....................................aaagggcuaacaaagaacuaacC.................   1          1          seq
    ...........................................uaacaaagaacuaacgcu...............   2          0          seq
    ...........................................uaacaaagaacuaacgcuc..............   2          0          seq
```

```
             novel-nve-miR-24_star
                                        novel-nve-miR-24_guide
cgccgacguuuugaguguuaguccuucgucgaagcaaaagggcuaacaaagaacuaacgcucgaaacgucagcgaa

...........................................uaacaaagaacuaacgcuU..............    1    1    seq
...........................................uaacaaagaacuaacgcucg.............    9    0    seq
...........................................uaacaaagaacuaacgcucA.............    1    1    seq
...........................................uaacaaagGacuaacgcucg.............    1    1    seq
...........................................uaacaaagaacuaacgcucgU............    1    1    seq
...........................................uaacaaagaacuaacgcucga............    18   0    seq
...........................................uaacaaagaacuaacgcucgaU...........    14   1    seq
...........................................uGacaaagaacuaacgcucgaa...........    1    1    seq
...........................................uaacaaagaacuaacgcucgGa...........    1    1    seq
...........................................uaacaaagaacuaacgcucgaG...........    5    1    seq
...........................................uaacaaagaacuaacgcucgaa...........    87   0    seq
...........................................uaacaaagaacuaacgcucgaC...........    5    1    seq
...........................................uaacaaagaacuaacgcucUaa...........    1    1    seq
...........................................uaacaaagaacuaGcgcucgaa...........    3    1    seq
...........................................uaacaaagaacCaacgcucgaa...........    1    1    seq
...........................................uaacGaagaacuaacgcucgaa...........    1    1    seq
...........................................uaacaGagaacuaacgcucgaa...........    1    1    seq
...........................................uaacaaagaacuGacgcucgaa...........    1    1    seq
...........................................Aaacaaagaacuaacgcucgaa...........    3    1    seq
...........................................uaacaaagaacuaacgcAcgaa...........    1    1    seq
...........................................uaacaaagaacuaacgcucgaaa..........    13   0    seq
...........................................uaacaaagaacuaacgcucgaaG..........    1    1    seq
...........................................uaacaaagaacuaacgcucgaaU..........    77   1    seq
...........................................uaacaaagaacuaacgcucgaaC..........    20   1    seq
...........................................uaacaaagaacuaacgcucgaaUc.........    2    1    seq
...........................................uaacaaagaacuaacgcucgaaac.........    6    0    seq
...........................................uaacaaagaacuaacgcucgaaacU........    1    1    seq
............................................aacaaagaacuaacgcucga............    1    0    seq
............................................aacaaagaacuaacgcucgaa...........    3    0    seq
.............................................acaaagaacuaacgcucga............    1    0    seq
.............................................acaaagaacuaacgcucgaa...........    6    0    seq
.............................................acaaagaacuaacgcucgaaU..........    2    1    seq
...............................................aaagaacuaacgcucgaaacguU......    1    1    seq
...............................................aaagaacuaacgcucgaaacguc......    1    0    seq
.....................................................cuaacgcucgaaacgucGgc...    1    1    seq
......................................................uaacgcucgaaacgucGgc...    2    1    seq
......................................................uaacgcucgaaacgucagcU..    1    1    seq
```

miRBase precursor : novel-nve-miR-25
Total read count : 79
novel-nve-miR-25_guide read count : 74
novel-nve-miR-25_star read count : 5
remaining reads : 0

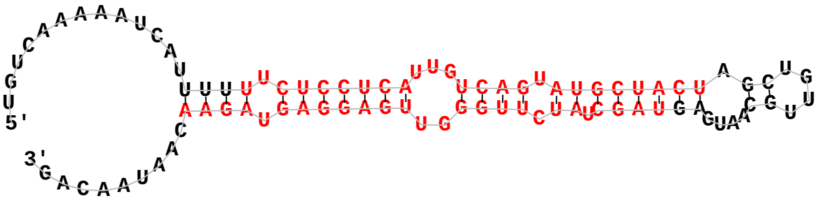

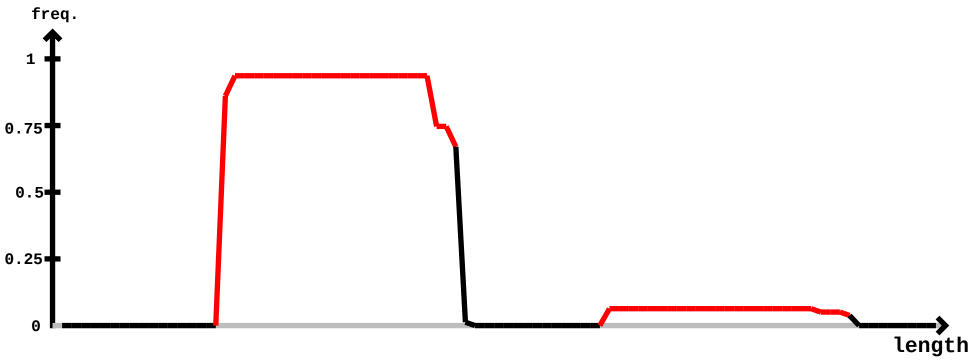


```
                                                                novel-nve-miR-25_star
                         novel-nve-miR-25_guide
5'- ugucaaaaaucauuuuuucuccucauugucaguaugcuacuagcuguugcaaugaguagcuaucuuggguugaggaguagaacaauaacag   -3'    exp
    .............((((.(((((((...((((.((((((((.((....))....)))))).)).)))).))))))).)))).........    reads      mm         sample
    ................uucuccucauugucaguaugcu.....................................................    8          0          seq
    ................uucuccucauugucaguaugAu.....................................................    1          1          seq
    ................uucuccucauugucaguaugcC.....................................................    4          1          seq
    ................Gucuccucauugucaguaugcu.....................................................    1          1          seq
    ................uucuccucauugucaguaugcuaU...................................................    1          1          seq
    ................uucuccucauugucaguaugcuac...................................................    5          0          seq
    ................uucuccGcauugucaguaugcuacu..................................................    1          1          seq
    ................uucuccucauugucaguaugcuacC..................................................    11         1          seq
    ................uCcuccucauugucaguaugcuacu..................................................    1          1          seq
    ................uucuccCcauugucaguaugcuacu..................................................    1          1          seq
    ................uucuccucauugucaguaugcuacu..................................................    33         0          seq
    ................uucuccucauugucaguaugcuacuU.................................................    1          1          seq
    .................ucuccucauugucaguaugcu.....................................................    1          0          seq
    .................ucuccucauugucaguaugcuacu..................................................    2          0          seq
    .................ucuccucauugucaguaugcuacC..................................................    1          1          seq
    .................uUuccucauugucaguaugcuacu..................................................    1          1          seq
    .................ucuccCcauugucaguaugcuacu..................................................    1          1          seq
    ........................................................uagcuaucuuggguugaggagu.............    1          0          seq
    ........................................................uagcuaucuuggguugaggaguaga..........    1          0          seq
    ........................................................uagcuaucuuggguugaggaguagaC.........    1          1          seq
    ........................................................uagcuaucuuggguugaggaguagaU.........    2          1          seq
```

miRBase precursor : novel-nve-miR-26
Total read count : 551
novel-nve-miR-26_guide read count : 449
novel-nve-miR-26_star read count : 92
remaining reads : 10

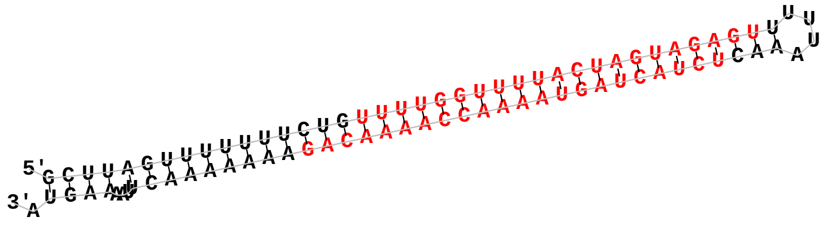

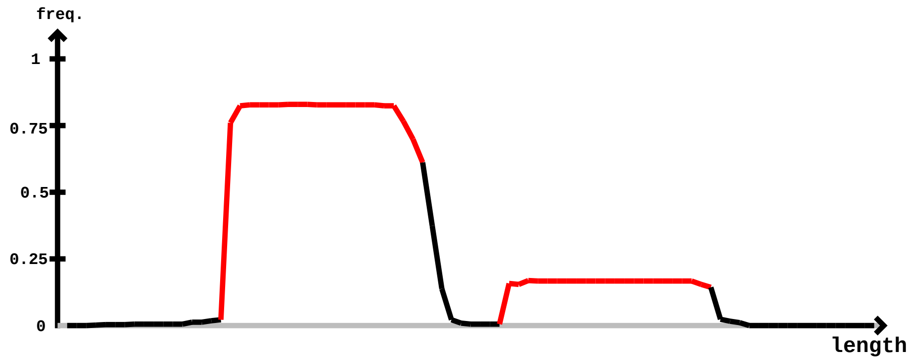


```
                                                          novel-nve-miR-26_star
                    novel-nve-miR-26_guide
5'- gcuuaguuuuuuucuguuuugguuuuacuaguagaguuuuuaaacucuacuaguaaaaccaaaacagaaaaaaacuuaaaagua  -3'   exp
    ((((((((((((((((((((((((((((((((((((((....))))))))))))))))))))))))))))))))))...)))).   reads   mm   sample
    ..uAaguuuuuuucuguuuugg..............................................................   1       1    seq
    ...uaUuuuuuuucuguuuugg..............................................................   1       1    seq
    ......uuuuuuucuguuuugguGu...........................................................   1       1    seq
    ............ucuguuuugguuuuacuagC....................................................   1       1    seq
    ............ucuguuuugguuuuacuagu....................................................   1       0    seq
    ............ucuguCuugguuuuacuaguaga.................................................   1       1    seq
    ............ucuguuuugguuuuacuaguagaU................................................   1       1    seq
    ..............uguuuugguuuuacuaguagag................................................   2       0    seq
    ..............uguuuugguuuuacuaguagagC...............................................   1       1    seq
    ...............guuuugguuuuacuaguag..................................................   1       0    seq
    ...............guuuugguuuuacuaguagag................................................   1       0    seq
    ................uuuugguuuuacuGguag..................................................   1       1    seq
    ................uuuugguuuuacuaguag..................................................   29      0    seq
    ................uuuugguuuuacuaguaA..................................................   1       1    seq
    ................uuuAgguuuuacuaguaga.................................................   1       1    seq
    ................uuuugguuuuaUuaguaga.................................................   1       1    seq
    ................Auuugguuuuacuaguaga.................................................   1       1    seq
    ................uuuugguuuuacuaguaga.................................................   29      0    seq
    ................Cuuugguuuuacuaguaga.................................................   1       1    seq
    ................uuuugguuuCacuaguaga.................................................   1       1    seq
    ................uuuugguuuuacAaguaga.................................................   1       1    seq
    ................uuuugguuuuacuaguagaA................................................   5       1    seq
    ................uuuugguuuuacuaguagaC................................................   1       1    seq
    ................uuuugguuuuacuaguagag................................................   35      0    seq
    ................uuCugguuuuacuaguagag................................................   1       1    seq
    ................uuuugguuuCacuaguagagu...............................................   1       1    seq
    ................uAuugguuuuacuaguagagu...............................................   2       1    seq
    ................Auuugguuuuacuaguagagu...............................................   1       1    seq
    ................uuuugguuuuacuaguagagA...............................................   1       1    seq
    ................Guuugguuuuacuaguagagu...............................................   1       1    seq
    ................uuuuggCuuuacuaguagagu...............................................   1       1    seq
    ................uuuugguuuuacuaguagagG...............................................   1       1    seq
    ................Cuuugguuuuacuaguagagu...............................................   1       1    seq
    ................uuuugguuuuacuaguagagu...............................................   87      0    seq
```

```
                              novel-nve-miR-26_star
               novel-nve-miR-26_guide
gcuuaguuuuuuucuguuuugguuuuacuaguagaguuuuuaaacucuacuaguaaaaccaaaacagaaaaaaacuuaaaagua

................uuuugguuuuacuaguagagC...............................................   17   1   seq
................uuuugguuuuacuGguagaguu..............................................   1    1   seq
................Cuuugguuuuacuaguagaguu..............................................   1    1   seq
................uuuugguuuuacuaguagaguA..............................................   5    1   seq
................uuuugguuuuacuaguagaguC..............................................   22   1   seq
................uuuugguuuuacuaguaAaguu..............................................   1    1   seq
................uuuCgguuuuacuaguagaguu..............................................   1    1   seq
................uuuuCguuuuacuaguagaguu..............................................   1    1   seq
................uuuugguuuuacuaguagaguu..............................................   86   0   seq
................Auuugguuuuacuaguagaguu..............................................   1    1   seq
................uuuuggCuuuacuaguagaguu..............................................   1    1   seq
................uuuugguuuuacuaguagUguu..............................................   1    1   seq
................uuuugguuuuacuaguagaguuu.............................................   39   0   seq
................uuuugguuuuacuaguagaguuC.............................................   10   1   seq
................uuuugguuuuacuaguagaguuA.............................................   8    1   seq
................uuuugguuuuacuaguagaguuAu............................................   3    1   seq
................uuuugguuuuacuaguagaguuuA............................................   1    1   seq
................uuuugguuuuacuaguagaguuuu............................................   2    0   seq
................uuuugguuuuacuaguagaguuuAu...........................................   1    1   seq
................uuuugguuuuacuaguagaguuuuu...........................................   1    0   seq
.................uuugguuuuacuaguaga.................................................   1    0   seq
.................uuugguuuuacuaguagag................................................   2    0   seq
.................uuugguuuuacuaguagagA...............................................   1    1   seq
.................uuugguuuuacuaguagagC...............................................   1    1   seq
.................uuugguuuuacuaguagagu...............................................   15   0   seq
.................uuugguuuuacuaguagaguu..............................................   6    0   seq
.................uuugguuuuacuaguagaguA..............................................   3    1   seq
.................uuugguuuuacuaguagaguuu.............................................   4    0   seq
.................uuugguuuuacuaguagaguuC.............................................   1    1   seq
.................uuugguuuuacuaguagaguuuA............................................   1    1   seq
..................uugguuuuacuaguagaguu..............................................   1    0   seq
..................uugguuuuacuaguagaguA..............................................   1    1   seq
......................uuuuacuaguagaguuuuuaaacC......................................   1    1   seq
......................uuuuacuaguagaguuuuuaaacu......................................   1    0   seq
......................uuuuacuaguagaguuuuuaaacucu....................................   1    0   seq
.............................................ucuacuaguaaaaccaaaac...................   7    0   seq
.............................................ucuacuaguaaaaccaaaaca..................   6    0   seq
.............................................ucuacuaguaaaaccaaaacag.................   47   0   seq
.............................................ucuacuaguaaaaccaaaacaA.................   9    1   seq
.............................................ucuacuaguaaaGccaaaacag.................   1    1   seq
.............................................uUuacuaguaaaaccaaaacag.................   1    1   seq
.............................................ucuacuaguaaaaccaGaacag.................   1    1   seq
.............................................ucuacuaguaaaaccaaaacaC.................   1    1   seq
.............................................ucuacuaguaaaaccaaaGcag.................   2    1   seq
.............................................ucuacAaguaaaaccaaaacag.................   1    1   seq
.............................................ucuacuaguaGaaccaaaacag.................   1    1   seq
.............................................uGuacuaguaaaaccaaaacag.................   1    1   seq
.............................................ucuacuaguaaaaccaaaacaU.................   1    1   seq
.............................................ucuacuaguaaaaccaaaacagU................   1    1   seq
.............................................ucuacuaguaaaaccaaaacagaU...............   2    1   seq
.............................................ucuacuaguaaaaccaaaacagaaU..............   2    1   seq
...............................................uacuaguaaaaccaaaacaga................   2    0   seq
...............................................uacuaguaaaaccaaaacagG................   1    1   seq
...............................................uacuaguaaaaccaaaacagaa...............   1    0   seq
...............................................uacuaguaaaaccaaaacagaaC..............   2    1   seq
...............................................uacuaguaaaaccaaaacagaaU..............   2    1   seq
```

miRBase precursor : novel-nve-miR-28
Total read count : 422
novel-nve-miR-28_guide read count : 421
novel-nve-miR-28_star read count : 1
remaining reads : 0

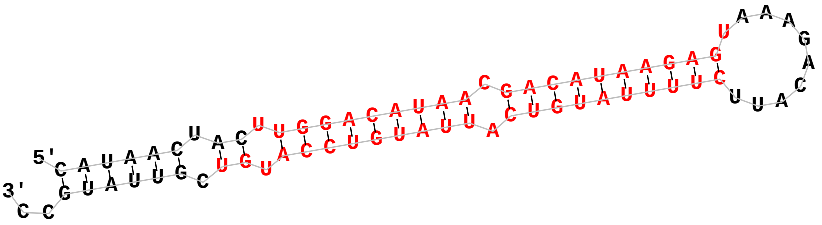

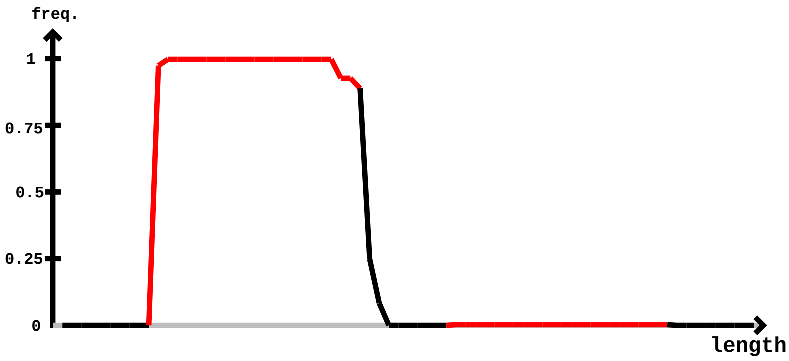


```
                    novel-nve-miR-28_guide
                                                    novel-nve-miR-28_star
5'- cauaacuacuuggacauaacgacauaagaguaaagacauucuuuuaugucauuauguccaugucguuaugcc   -3'    exp
    ((((((.((.(((((((((.((((((((((.........)))))))))).))))))))).)).))))))..     reads       mm          sample
    .........uuggUcauaacgacauaag............................................   1           1           seq
    .........uuggGcauaacgacauaag............................................   1           1           seq
    .........uuggacauaacgacauaaA............................................   1           1           seq
    .........uuggacauaacgacauaag............................................   23          0           seq
    .........uuggacauaacgacauaagaA..........................................   2           1           seq
    .........uuggacauaacgacauaagag..........................................   10          0           seq
    .........Auggacauaacgacauaagag..........................................   1           1           seq
    .........uuggacauaGcgacauaagagu.........................................   1           1           seq
    .........uuggacauaacgacauaagagA.........................................   3           1           seq
    .........uuggacauaacgacauaagagG.........................................   3           1           seq
    .........Guggacauaacgacauaagagu.........................................   1           1           seq
    .........uuggacauaacgacaCaagagu.........................................   1           1           seq
    .........uCggacauaacgacauaagagu.........................................   3           1           seq
    .........uuggacauaacgacauaagagu.........................................   218         0           seq
    .........Auggacauaacgacauaagagu.........................................   2           1           seq
    .........uuggacauaacgacauaagagC.........................................   27          1           seq
    .........uuggacauaacgacauaUgagu.........................................   1           1           seq
    .........uuggacauaacgacauaGgagu.........................................   1           1           seq
    .........uAggacauaacgacauaagagu.........................................   1           1           seq
    .........uuggacauaacgacUuaagagu.........................................   1           1           seq
    .........uuggacauaacgacGuaagagu.........................................   2           1           seq
    .........uuggacauaacgacauGagagu.........................................   1           1           seq
    .........uuggacauaacgaUauaagagu.........................................   1           1           seq
    .........uuggaAauaacgacauaagagu.........................................   2           1           seq
    .........uuggacauaacgGcauaagagua........................................   1           1           seq
    .........uuggacauaacgacauaagagua........................................   54          0           seq
    .........uuggacauaacgacauaaguU..........................................   7           1           seq
    .........uuggacauaacgacauaagaguG........................................   1           1           seq
    .........uuggacauaacgacauaagaguC........................................   4           1           seq
    .........uuCgacauaacgacauaagagua........................................   1           1           seq
    .........uuggacauaacgacauaagaguaC.......................................   4           1           seq
    .........uuggacauaacgacauaagaguaU.......................................   31          1           seq
    ..........uggacauaacgacauaag............................................   3           0           seq
    ..........uggacaGaacgacauaag............................................   1           1           seq
```

```
         novel-nve-miR-28_guide
                                        novel-nve-miR-28_star
cauaacuacuuggacauaacgacauaagaguaaagacauucuuuuaugucauuauguccaugucguuaugcc

..........uggacaGaacgacauaagag..........................................    3    1    seq
..........uggacauaacgacauaagagA.........................................    1    1    seq
..........uggacauaacgacauaagagGa........................................    2    1    seq
........................................Uuuuuaugucauuauguccaugu.........    1    1    seq
```

miRBase precursor : novel-nve-miR-29
Total read count : 108
novel-nve-miR-29_guide read count : 81
novel-nve-miR-29_star read count : 26
remaining reads : 1

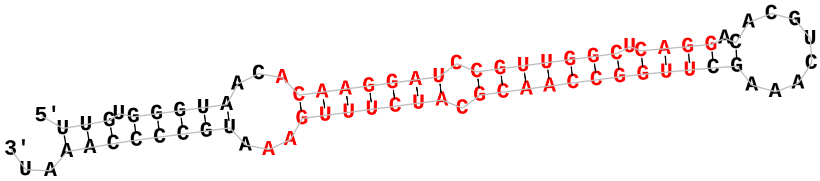

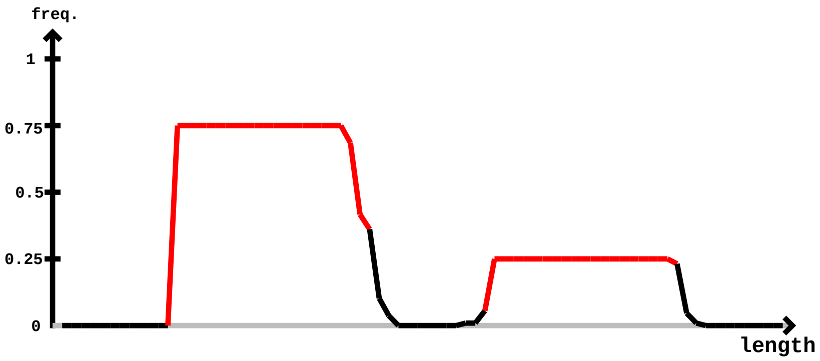


novel-nve-miR-29_guide

novel-nve-miR-29_star

```
5'- uuguggguaacacaaggauccguuggcucaggacacgucaaagcuuggccaacgcaucuuugaaaugccccaaau  -3'    exp
    (((.(((((...(((((((.(((((((.((((.(........))))))))))))).)))))))...))))))))..   reads      mm         sample
    ...........acaaggauccguuggcuG..............................................   4          1          seq
    ...........acaaggauccguuggcuc..............................................   2          0          seq
    ...........acaaggauccguuggcuU..............................................   1          1          seq
    ...........acaaggauccguuggcucG.............................................   16         1          seq
    ...........acaaggGuccguuggcuca.............................................   1          1          seq
    ...........acaaggauccguuggcuca.............................................   12         0          seq
    ...........acaaggauccguuggcucag............................................   6          0          seq
    ...........acaaggauccguuggcucagU...........................................   1          1          seq
    ...........Gcaaggauccguuggcucagg...........................................   1          1          seq
    ...........acaaggauccguuggcucagA...........................................   6          1          seq
    ...........acaaggauccguuggcAcagg...........................................   1          1          seq
    ...........acaaggauccguuggcucagg...........................................   16         0          seq
    ...........acaaggauccguuggcucaCg...........................................   3          1          seq
    ...........acaaggauccguuggcucaggC..........................................   1          1          seq
    ...........acaaggauccguuggcucaggU..........................................   1          1          seq
    ...........acaaggauccguuggcucagga..........................................   5          0          seq
    ...........acaaggauccguuggcucaggaU.........................................   4          1          seq
    .........................................agcuuggccaacgcaucuuugaC...........   1          1          seq
    ...........................................cuuggccaacgcaucuuugaa...........   1          0          seq
    ...........................................cuuggccaacgcaucuuugaG...........   2          1          seq
    ...........................................cuuggccaacgcaucuuugaU...........   1          1          seq
    ...........................................cuuggccaacgcaucuuugaaC..........   1          1          seq
    ............................................uuggccaacgcaucuuuga............   2          0          seq
    ............................................uuggccaacgcaucuuugaC...........   1          1          seq
    ............................................uuggccaacgcaucuuugaG...........   1          1          seq
    ............................................uuggccaacgcaucuuugaa...........   10         0          seq
    ............................................uuggccaacgcaucuuugaU...........   3          1          seq
    ............................................uuggccaacgcaucuuugaaa..........   2          0          seq
    ............................................uuggccaacgcaucuuugaUa..........   1          1          seq
    ............................................uuggccaacgcaucuuugaUau.........   1          1          seq
```

miRBase precursor : novel-nve-miR-30
Total read count : 82
novel-nve-miR-30_guide read count : 73
novel-nve-miR-30_star read count : 8
remaining reads : 1

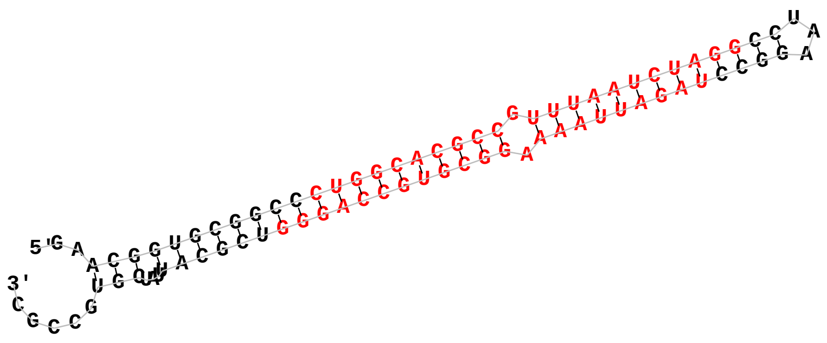

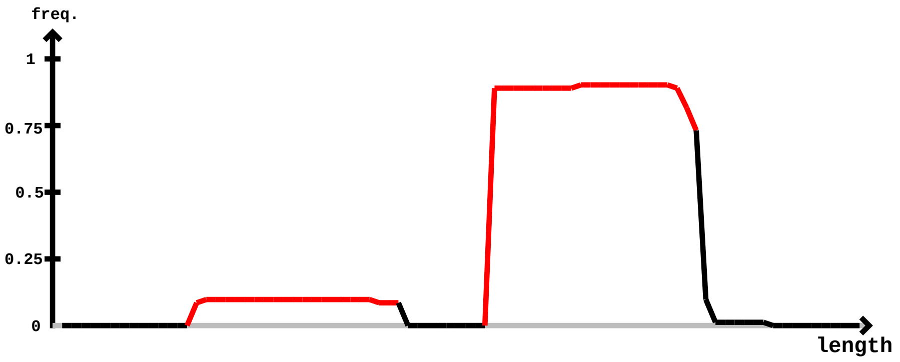


```
                                                      novel-nve-miR-30_guide
                     novel-nve-miR-30_star
5'- gaacggugcggcccuggcacgccguuuaaucuaggccuaaggccuagauuaaaaggcgugccagggucgcauuaucgugccgc  -3'   exp
    ..((((((((((((((((((((.(((((((((((((...))))))))))))).))))))))))))))))))...))).....   reads    mm    sample
    .............cuggcacgccguuuaaucu...................................................   1        0     seq
    .............cuggcacgccguuuaaucuagC................................................   1        1     seq
    .............cuggcacgccguuuaaucuagU................................................   1        1     seq
    .............cuggcacgccguuuaaucuagg................................................   4        0     seq
    ..............uggcacgccguuuaaucuagA................................................   1        1     seq
    ............................................uagauuaaaaggcgugcca....................   1        0     seq
    ............................................uagauuaaaaggcgCgccag...................   1        1     seq
    ............................................uagauuaaaaggcgugccaA...................   1        1     seq
    ............................................uagauuaaaaggcgugccaU...................   1        1     seq
    ............................................uagauuaaaaggcgugccag...................   3        0     seq
    ............................................uagauuaaaaggcgugccagA..................   2        1     seq
    ............................................uagauuaaaaggcgugccagg..................   5        0     seq
    ............................................uagauuaaaaggcgugccaggA.................   8        1     seq
    ............................................uagauuaaaaggcgugccaggC.................   2        1     seq
    ............................................uagauuaaaaggUgugccaggg.................   1        1     seq
    ............................................uagauuaaaaggcgugccaggU.................   2        1     seq
    ............................................uagauuaaaaggcgugccaggg.................   36       0     seq
    ............................................uagauuaaaagCcgugccaggg.................   1        1     seq
    ............................................Gagauuaaaaggcgugccaggg.................   2        1     seq
    ............................................uagauuaaaaggcgugccagggA................   3        1     seq
    ............................................uagauuaaaaggcgugccagggu................   3        0     seq
    ............................................uagauuaaaaggcgugccagggC................   1        1     seq
    .....................................................Cggcgugccagggucgcauu..........   1        1     seq
```

miRBase precursor : novel-nve-miR-32
Total read count : 611
novel-nve-miR-32_guide read count : 595
novel-nve-miR-32_star read count : 16
remaining reads : 0

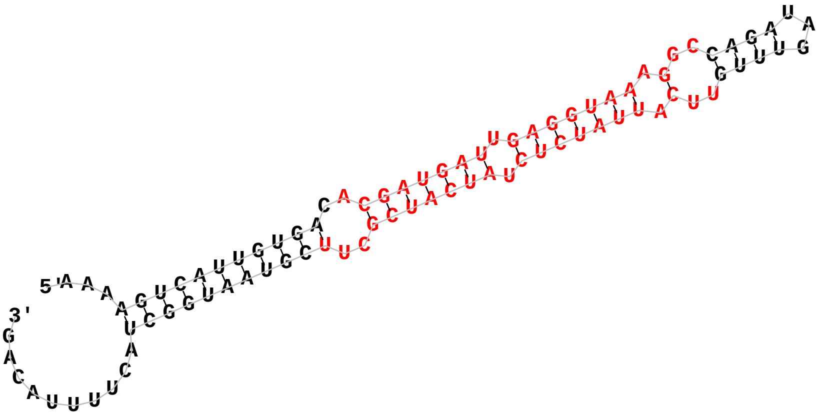

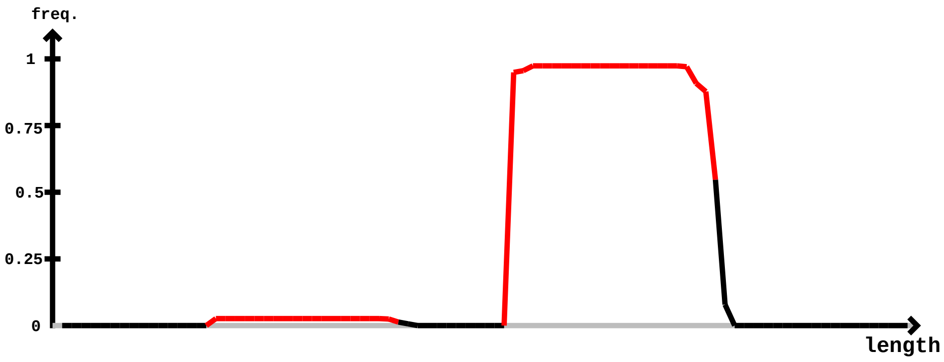


```
                                              novel-nve-miR-32_guide
               novel-nve-miR-32_star
5'- aaaagucauugugacacgaugauugagguaaaggccagauaguuuguucauuaucucuaucaucgcuucguaauggcuacuuuuacag  -3'   exp
    ...((((((((((..(((((((.(((((((.(..((((...))))..).))))))).)))))))..))))))))))............  reads  mm  sample
    ...............acgaugauugagguaaag.......................................................  1  0  seq
    ...............acgaugauugagguaaagg......................................................  6  0  seq
    ...............acgaugauugagguaaagA......................................................  1  1  seq
    ...............acgaugauugagguaaaggc.....................................................  3  0  seq
    ...............acgaugauugagguaaaggU.....................................................  1  1  seq
    ...............acgaugauugagguaaaggcU....................................................  2  1  seq
    ...............acgaugauugagguaaaggAc....................................................  1  1  seq
    ...............acgaugauugagguaaaggcc....................................................  1  0  seq
    ..............................................uucauuaucucuaucauc........................  2  0  seq
    ..............................................uucauuaucucuaucaucA.......................  5  1  seq
    ..............................................uAcauuaucucuaucaucg.......................  1  1  seq
    ..............................................uucauuaucucuaucaucU.......................  3  1  seq
    ..............................................uucauuaucucuaucaucg.......................  26  0  seq
    ..............................................Aucauuaucucuaucaucg.......................  1  1  seq
    ..............................................uucauuaucucuUucaucg.......................  1  1  seq
    ..............................................uucauuaucucuaucauAg.......................  1  1  seq
    ..............................................uucauuaucucuaucaucgc......................  7  0  seq
    ..............................................uucauuaucucuaucaucgA......................  1  1  seq
    ..............................................uucauuaucucuaucaucgU......................  8  1  seq
    ..............................................uucauuaucucuaucaucgG......................  1  1  seq
    ..............................................uucauuaucucuaCcaucgc......................  1  1  seq
    ..............................................uucauuaucCcuaucaucgc......................  1  1  seq
    ..............................................uucauuaucuAuaucaucgcu.....................  1  1  seq
    ..............................................uucauuaAcucuaucaucgcu.....................  1  1  seq
    ..............................................uucauuaucucuaucaucgcu.....................  134  0  seq
    ..............................................uucauuaucucuaucaucUcu.....................  1  1  seq
    ..............................................uucauuaucucuaucGucgcu.....................  1  1  seq
    ..............................................uuUauuaucucuaucaucgcu.....................  1  1  seq
    ..............................................uucauuaucucuaucaCcgcu.....................  2  1  seq
    ..............................................uucauuaucucuaucaucgUu.....................  1  1  seq
    ..............................................uCcauuaucucuaucaucgcu.....................  1  1  seq
    ..............................................uucaAuaucucuaucaucgcu.....................  1  1  seq
    ..............................................uucauuaucucuaucaucgcC.....................  32  1  seq
    ..............................................uucauuaucucuauAaucgcu.....................  1  1  seq
```

```
                                              novel-nve-miR-32_guide
               novel-nve-miR-32_star
aaaagucauugugacacgaugauugagguaaaggccagauaguuuguucauuaucucuaucaucgcuucguaauggcuacuuuuacag

..............................................Gucauuaucucuaucaucgcu.....................    1     1     seq
..............................................uucauuaucucuaucaucgcG.....................    4     1     seq
..............................................uucauuaucucuUucaucgcu.....................    1     1     seq
..............................................uucUuuaucucuaucaucgcu.....................    1     1     seq
..............................................uucauuaCcucuaucaucgcu.....................    1     1     seq
..............................................uuAauuaucucuaucaucgcu.....................    1     1     seq
..............................................uucauuaucucuaucaucgcA.....................    12    1     seq
..............................................uucauuUucucuaucaucgcuu....................    1     1     seq
..............................................uucauuaucucuaAcaucgcuu....................    1     1     seq
..............................................uucauuaucucuaucaucgcuA....................    9     1     seq
..............................................uucauuaCcucuaucaucgcuu....................    1     1     seq
..............................................uucauuaucucuaucaucgcGu....................    1     1     seq
..............................................Aucauuaucucuaucaucgcuu....................    1     1     seq
..............................................uucauuaucGcuaucaucgcuu....................    1     1     seq
..............................................uucauuauAucuaucaucgcuu....................    1     1     seq
..............................................uucauuaucucuaucaucgcuC....................    52    1     seq
..............................................uucauuaucucuaucaucgcuG....................    9     1     seq
..............................................uucauAaucucuaucaucgcuu....................    1     1     seq
..............................................uAcauuaucucuaucaucgcuu....................    2     1     seq
..............................................uucauuaucCcuaucaucgcuu....................    1     1     seq
..............................................uucauuaucucCaucaucgcuu....................    1     1     seq
..............................................uucauuaucucuaucaCcgcuu....................    1     1     seq
..............................................uucauuaucucuaucaucgUuu....................    1     1     seq
..............................................uucauuaucucuGucaucgcuu....................    1     1     seq
..............................................uucauuaucucuaucaucgcuu....................    190   0     seq
..............................................uucauuaucucuaucaucgcAu....................    1     1     seq
..............................................uucauuaucucuaucaucgcuuG...................    1     1     seq
..............................................uucauuaucucuaucaucgcuuA...................    2     1     seq
..............................................uucauuaucucuaucaucgcuuc...................    8     0     seq
..............................................uucauuaucucuaucaucgcuuU...................    36    1     seq
...............................................ucauuaucucuaucaucgcu.....................    2     0     seq
...............................................ucauuaucucuaucaucgcuC....................    1     1     seq
...............................................ucauuaucucuaucaucgcuu....................    1     0     seq
................................................cauuaucucuaucaucCcu.....................    1     1     seq
................................................cauuaucucuaucaucgcu.....................    1     0     seq
................................................cauuaucucuaucaucgcuC....................    2     1     seq
................................................cauuaucucuaucaucgcuu....................    6     0     seq
................................................cauuaucucuaucaucgcuAc...................    1     1     seq
```

miRBase precursor : novel-nve-miR-33
Total read count : 376
novel-nve-miR-33_guide read count : 361
novel-nve-miR-33_star read count : 1
remaining reads : 14

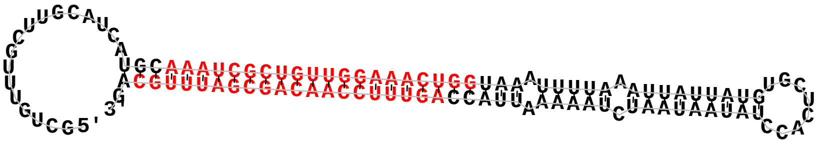

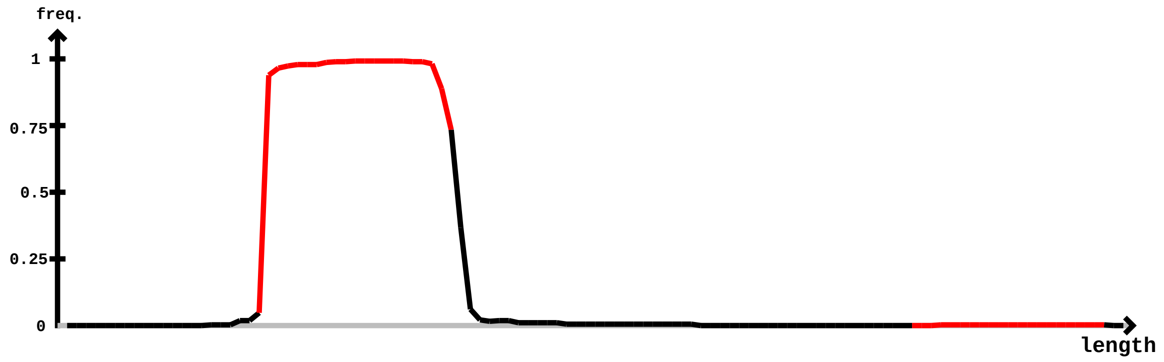


```
                                   novel-nve-miR-33_guide
                                                                                       novel-nve-miR-33_star
5'- gcuguuugcuugcaucaugcaaaucgcuguuggaaacugguaaauuuuaaauuauuaugugcucaccuauaauaaucuaaaaaauuaccaguuuccaacagcgauuugcag   -3'    exp
    ((.....))........((((((((((((((((((((((((((.(((((.(((((((((........))))))))).))))).))))))))))))))))))))))))))).   reads   mm   sample
    ..............ucGugcaaaucgcuguuggaa............................................................................   1       1    seq
    .................ugcaaaucgcuguuggaaac..........................................................................   1       0    seq
    .................ugcaaaucgcugAuggaaac..........................................................................   1       1    seq
    .................Cgcaaaucgcuguuggaaac..........................................................................   1       1    seq
    .................ugcaaaucgcuguuggaaacA.........................................................................   1       1    seq
    .................ugcaaaucgcuguuggaaacu.........................................................................   2       0    seq
    ...................caaaucgcuguuggaaacug........................................................................   1       0    seq
    ...................caaaucgcuguuggaaacugg.......................................................................   1       0    seq
    ...................cGaaucgcuguuggaaacugg.......................................................................   2       1    seq
    ...................caaaucgcuguuggaaacugA.......................................................................   1       1    seq
    ...................caaaucgcuguuggaaacuggA......................................................................   2       1    seq
    ...................caaaucgcuguuggaaacuggG......................................................................   1       1    seq
    ...................caaaucgcuguuggaaacuggu......................................................................   1       0    seq
    ...................caaaucgcuguuggaaacugguaC....................................................................   2       1    seq
    ....................aaaucgcugCuggaaacu.........................................................................   1       1    seq
    ....................aaaucgcuguuggaaacA.........................................................................   1       1    seq
    ....................aaaucgcuguuggaaacC.........................................................................   7       1    seq
    ....................aaaucgcCguuggaaacu.........................................................................   1       1    seq
    ....................aaaucgcuguuggaaacu.........................................................................   11      0    seq
    ....................aaaucgcuguuggaGacu.........................................................................   1       1    seq
    ....................aGaucgcuguuggaaacu.........................................................................   10      1    seq
    ....................aaaucgcuguuggaaacug........................................................................   33      0    seq
    ....................aGaucgcuguuggaaacug........................................................................   10      1    seq
    ....................aaaucgcuguuggaGacug........................................................................   1       1    seq
    ....................aaaucgcuguuggaaacuC........................................................................   1       1    seq
    ....................aaaucgcuguuggaaacuA........................................................................   9       1    seq
    ....................aaaucgcugCuggaaacug........................................................................   1       1    seq
    ....................aaaucgcCguuggaaacug........................................................................   1       1    seq
    ....................Gaaucgcuguuggaaacug........................................................................   1       1    seq
    ....................aaaucgcuguuggaaacugA.......................................................................   21      1    seq
    ....................Uaaucgcuguuggaaacugg.......................................................................   1       1    seq
    ....................aaaucgcugCuggaaacugg.......................................................................   1       1    seq
    ....................aaGucgcuguuggaaacugg.......................................................................   1       1    seq
    ....................aaaucgcuguuggaGacugg.......................................................................   6       1    seq
```

```
                    novel-nve-miR-33_guide
                                                                                        novel-nve-miR-33_star
gcuguuugcuugcaucaugcaaaucgcuguuggaaacugguaaauuuuaaauuauuaugugcucaccuauaauaaucuaaaaauuaccaguuuccaacagcgauuugcag

....................aaaucAcuguuggaaacugg......................................................................  1   1   seq
....................aaauAgcuguuggaaacugg......................................................................  1   1   seq
....................aaaucgcuguuggaaaUugg......................................................................  1   1   seq
....................aaaucgcuguugCaaacugg......................................................................  1   1   seq
....................aaaucgcuguuggaaacugg......................................................................  68  0   seq
....................Gaaucgcuguuggaaacugg......................................................................  3   1   seq
....................aaaucgcuguuggaaacugU......................................................................  1   1   seq
....................aGaucgcuguuggaaacugg......................................................................  19  1   seq
....................aaauUgcuguuggaaacugg......................................................................  1   1   seq
....................aGaucgcuguuggaaacuggu.....................................................................  4   1   seq
....................aaaucgcuguuggaGacuggu.....................................................................  3   1   seq
....................aaaucgcuguuggaaacuggu.....................................................................  34  0   seq
....................aaaucgcuguuggaaacuggG.....................................................................  2   1   seq
....................aaaucgcugCuggaaacuggu.....................................................................  1   1   seq
....................aaaCcgcuguuggaaacuggu.....................................................................  1   1   seq
....................aaaucgcuguuggaaaUuggu.....................................................................  1   1   seq
....................aaaucCcuguuggaaacuggu.....................................................................  1   1   seq
....................aaaucgcuguuggaaacuggC.....................................................................  6   1   seq
....................aaaucAcuguuggaaacuggu.....................................................................  1   1   seq
....................aaaucgcCguuggaaacuggu.....................................................................  2   1   seq
....................Gaaucgcuguuggaaacuggu.....................................................................  1   1   seq
....................aaaucgcuguuggaaacuggA.....................................................................  48  1   seq
....................aaauUgcuguuggaaacuggu.....................................................................  3   1   seq
....................Gaaucgcuguuggaaacuggua....................................................................  1   1   seq
....................aaaucgcuguuggaGacuggua....................................................................  1   1   seq
....................aGaucgcuguuggaaacuggua....................................................................  4   1   seq
....................aaaucgcuguuggaaacugAua....................................................................  1   1   seq
....................aaaucgcuguuggaaacugguC....................................................................  3   1   seq
....................aaaucgcuguuggaaacuggua....................................................................  2   0   seq
.....................aaucgcuguuggaaacugA......................................................................  2   1   seq
.....................aaucgcuguuggaaacAgg......................................................................  1   1   seq
.....................aaucgcuguuggaaacugg......................................................................  1   0   seq
.....................Caucgcuguuggaaacugg......................................................................  1   1   seq
.....................aaucgcuguuggaaacugU......................................................................  2   1   seq
.....................aaucgcuguuggaaacuggu.....................................................................  1   0   seq
.....................aaucgcuguuggaaacuggA.....................................................................  2   1   seq
......................aucgcuguuggaGacugg......................................................................  1   1   seq
......................aucgcuguuggaaacuggA.....................................................................  1   1   seq
......................aucgcuguuggaaacuggAa....................................................................  1   1   seq
.......................ucgcuguuggaaacuggua....................................................................  2   0   seq
..........................cuguuggaaacugguaaauu................................................................  2   0   seq
..........................cuguuggaaacugguaaauC................................................................  1   1   seq
...........................uguuggaaacugguaaauuuuaUa...........................................................  1   1   seq
.............................uuggaaacugguaaauuuuaUa...........................................................  1   1   seq
.........................................Uaauuuuaaauuauuaugugcuca.............................................  1   1   seq
............................................uuuuaaauuauuaugugcucU.............................................  1   1   seq
...........................................................................................uuccaaUagcgauuugc..  1   1   seq
```

miRBase precursor : novel-nve-miR-37
Total read count : 2843
novel-nve-miR-37_guide read count : 2819
novel-nve-miR-37_star read count : 2
remaining reads : 22

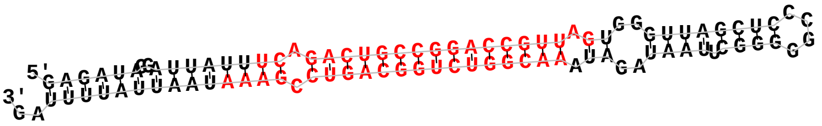

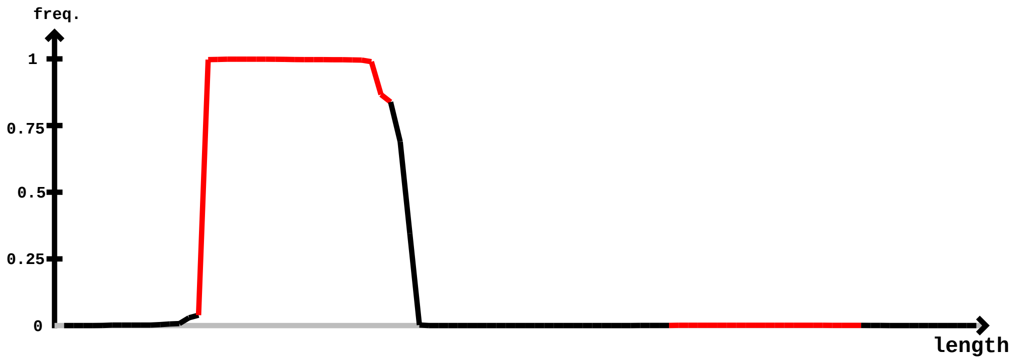


```
                                                                 novel-nve-miR-37_star
                 novel-nve-miR-37_guide
5'- gagauaggauuauuucagacugccggaccguuaguggguuagcucccgggggcuuaauagauaaacggucuggcaguccgaaauaauuauuuuag   -3'    exp
    (((((..((((((((.((((((((((((((.((..((((((((....)))).))))..)).))))))))))))))).)))))))))))))..    reads      mm         sample
    ...auaggauuauuucGgacu..........................................................................    1          1          seq
    ....uaggauuauuucagacug.........................................................................    1          0          seq
    ....uaggauuauuucagacugc........................................................................    2          0          seq
    ....uaggauuauuucagacugcc.......................................................................    1          0          seq
    .........uuauuucGgacugccgga....................................................................    1          1          seq
    .........uuauuucGgacugccggacc..................................................................    2          1          seq
    .........uuauuucagacugccggaccguua..............................................................    1          0          seq
    .........uuauuucagacugccggaccguuU..............................................................    1          1          seq
    ..........uauuucagacugccggaccgu................................................................    5          0          seq
    ..........uauuucagacugccggaccguuU..............................................................    1          1          seq
    ..........uauuucagacugccggaccguuaU.............................................................    1          1          seq
    ...........auuucGgacugccggaccgu................................................................    1          1          seq
    ...........auuucagacugccggaccgu................................................................    1          0          seq
    ...........auuucagacugccggaccguu...............................................................    2          0          seq
    ............uuucGgacugccggaccg.................................................................    2          1          seq
    ............uuucagacugccggaccgu................................................................    3          0          seq
    ............uuucGgacugccggaccgu................................................................    5          1          seq
    ............uuucagacugccggaccgC................................................................    1          1          seq
    ............Auucagacugccggaccgu................................................................    1          1          seq
    ............uuucagacugccggaccguC...............................................................    3          1          seq
    ............uuucagacugcUggaccguu...............................................................    1          1          seq
    ............uuucagacugccggaccguu...............................................................    1          0          seq
    ............uuucGgacugccggaccguu...............................................................    11         1          seq
    ............uuucagacugUcggaccguua..............................................................    1          1          seq
    ............uuucGgacugccggaccguua..............................................................    14         1          seq
    ............uuucagacugccggaccguuC..............................................................    1          1          seq
    ............uuucagacugccggaccguua..............................................................    5          0          seq
    ............uuucagacugccggaccguuU..............................................................    4          1          seq
    ............uuucagacugccggaccguuaA.............................................................    1          1          seq
    ............uuucagacugccggaccguuaU.............................................................    2          1          seq
    ............uuucagacugccggaccguuag.............................................................    6          0          seq
    .............uucagacugccggaccguC...............................................................    1          1          seq
    .............uucGgacugccggaccguu...............................................................    4          1          seq
    .............uucGgacugccggaccguua..............................................................    2          1          seq
```

```
                                                                                 novel-nve-miR-37_star
              novel-nve-miR-37_guide
gagauaggauuauuucagacugccggaccguuaguggguuagcucccgggggcuuaauagauaaacggucuggcaguccgaaauaauuauuuuag

.............uucagacugccggaccguuU..............................................................  1    1    seq
.............uucagacugccggaccguuag.............................................................  6    0    seq
.............uucagacugccggaccguuaC.............................................................  1    1    seq
.............uAcagacugccggaccguuagu............................................................  1    1    seq
.............uucagacugccggaccguuagu............................................................  10   0    seq
.............uucagacugccggaccguuagC............................................................  1    1    seq
.............uucagacugccggaccguuaguA...........................................................  1    1    seq
..............ucagacugccggaccAuu...............................................................  1    1    seq
..............ucagGcugccggaccguu...............................................................  2    1    seq
..............ucagacugccggacUguu...............................................................  1    1    seq
..............ucagacuAccggaccguu...............................................................  1    1    seq
..............ucagacugcAggaccguu...............................................................  1    1    seq
..............ucUgacugccggaccguu...............................................................  2    1    seq
..............uAagacugccggaccguu...............................................................  1    1    seq
..............ucagacugccggaccguC...............................................................  53   1    seq
..............ucagacugccggUccguu...............................................................  1    1    seq
..............Acagacugccggaccguu...............................................................  2    1    seq
..............ucagacugccggaccguu...............................................................  244  0    seq
..............ucagacugcUggaccguu...............................................................  1    1    seq
..............ucagacugccggaccguA...............................................................  13   1    seq
..............Ccagacugccggaccguu...............................................................  1    1    seq
..............ucagacuCccggaccguu...............................................................  1    1    seq
..............ucagacAgccggaccguu...............................................................  1    1    seq
..............ucagacugccggaccgCu...............................................................  1    1    seq
..............ucagacugccggaccguuU..............................................................  5    1    seq
..............uUagacugccggaccguua..............................................................  1    1    seq
..............ucagacugccggaccguAa..............................................................  1    1    seq
..............ucagacugccggaccguua..............................................................  41   0    seq
..............ucagacugcUggaccguua..............................................................  1    1    seq
..............ucagacugccggaccAuuag.............................................................  1    1    seq
..............ucagacugccggaccguuUg.............................................................  1    1    seq
..............ucagacugccggaccguuag.............................................................  349  0    seq
..............ucagacugcUggaccguuag.............................................................  3    1    seq
..............ucagacCgccggaccguuag.............................................................  1    1    seq
..............ucaUacugccggaccguuag.............................................................  1    1    seq
..............ucagacugccggaccguAag.............................................................  2    1    seq
..............ucagacugccggaccguuaA.............................................................  16   1    seq
..............ucGgacugccggaccguuag.............................................................  2    1    seq
..............ucaAacugccggaccguuag.............................................................  1    1    seq
..............Ccagacugccggaccguuag.............................................................  2    1    seq
..............ucagacuCccggaccguuag.............................................................  1    1    seq
..............Acagacugccggaccguuag.............................................................  2    1    seq
..............ucagacugAcggaccguuag.............................................................  1    1    seq
..............ucagacugccggaccgCuag.............................................................  2    1    seq
..............ucagacugccggaGcguuag.............................................................  1    1    seq
..............ucagacugccggaccguuaC.............................................................  4    1    seq
..............ucagacugccggaccguuaU.............................................................  6    1    seq
..............ucagGcugccggaccguuag.............................................................  1    1    seq
..............ucagacugccggaccguCag.............................................................  2    1    seq
..............ucagCcugccggaccguuag.............................................................  2    1    seq
..............ucagacugccAgaccguuag.............................................................  1    1    seq
..............ucagacugccggaccguuGg.............................................................  2    1    seq
..............ucagacugccggacUguuag.............................................................  1    1    seq
..............ucagacugccgUaccguuagu............................................................  1    1    seq
..............ucagGcugccggaccguuagu............................................................  3    1    seq
..............ucagacugccggacAguuagu............................................................  1    1    seq
..............ucagacugccAgaccguuagu............................................................  2    1    seq
..............ucagacugccggaUcguuagu............................................................  1    1    seq
..............ucagacuUccggaccguuagu............................................................  1    1    seq
..............ucagacugccggaccCuuagu............................................................  1    1    seq
..............ucagacugccggGccguuagu............................................................  3    1    seq
..............ucagacugccggacUguuagu............................................................  3    1    seq
..............ucagacugccggaccguuaAu............................................................  3    1    seq
..............uUagacugccggaccguuagu............................................................  1    1    seq
..............uAagacugccggaccguuagu............................................................  1    1    seq
..............Ccagacugccggaccguuagu............................................................  2    1    seq
..............ucGgacugccggaccguuagu............................................................  2    1    seq
..............ucagacugccggaccguuagA............................................................  17   1    seq
..............ucagacugccggaccguuagG............................................................  14   1    seq
```

```
                                                               novel-nve-miR-37_star
              novel-nve-miR-37_guide
gagauaggauuauuucagacugccggaccguuaguggguuagcucccgggggcuuaauagauaaacggucuggcaguccgaaauaauuauuuuag

..............ucagacugccggaccguuagu............................................................    710    0    seq
..............ucagacugccggaccguuagC............................................................    167    1    seq
..............ucagacCgccggaccguuagu............................................................    3      1    seq
..............Gcagacugccggaccguuagu............................................................    1      1    seq
..............ucagacugAcggaccguuagu............................................................    1      1    seq
..............ucaUacugccggaccguuagu............................................................    1      1    seq
..............ucagacugccggaccgAuagu............................................................    1      1    seq
..............ucagacugccggaccguuGgu............................................................    3      1    seq
..............Acagacugccggaccguuagu............................................................    12     1    seq
..............ucagacugccggCccguuagu............................................................    1      1    seq
..............ucagacugcAggaccguuagu............................................................    2      1    seq
..............ucagacugccggaccAuuagu............................................................    3      1    seq
..............ucagacugccggaccguCagu............................................................    1      1    seq
..............ucagacugccggaccguuaguC...........................................................    166    1    seq
..............ucagacugccggaccguuaguU...........................................................    767    1    seq
..............ucagacugccggaccguuaguA...........................................................    30     1    seq
..............ucagacugccggaccguuagug...........................................................    12     0    seq
..............ucagacugccggaccguuaguUg..........................................................    3      1    seq
..............ucagacugccggaccguuagugA..........................................................    2      1    seq
..............ucagacugccggaccguuagugU..........................................................    1      1    seq
...............cagacugccggaccguuag.............................................................    1      0    seq
...............cagacugccggaccguuaguU...........................................................    1      1    seq
................agacugccggaccguuagC............................................................    1      1    seq
................agacugcAggaccguuagu............................................................    1      1    seq
................agacugccggaccguuaguU...........................................................    1      1    seq
...........................................................gauaaacggucuggcaguUc................    1      1    seq
...............................................................aacggucuggcaguccgaaa............    1      0    seq
...............................................................aacggucuggcaguccgaaaua..........    1      0    seq
```

miRBase precursor : novel-nve-miR-42-a
Total read count : 107
novel-nve-miR-42-a_guide read count : 105
novel-nve-miR-42-a_star read count : 1
remaining reads : 1

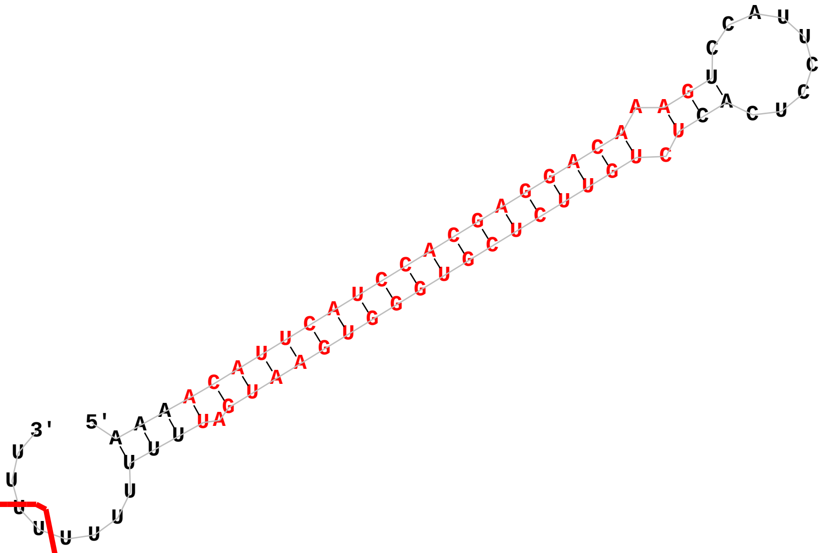

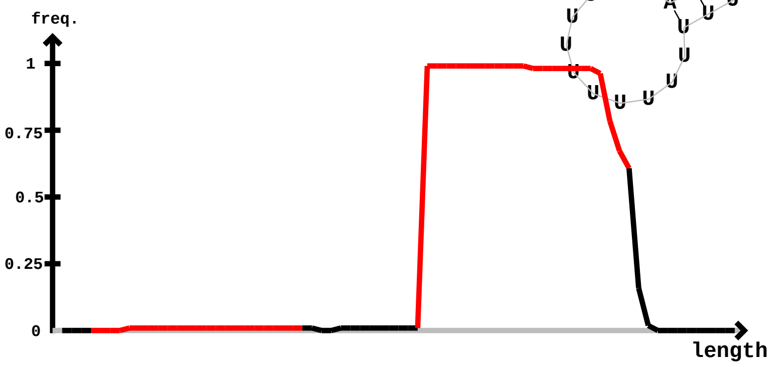


```
        novel-nve-miR-42-a_star
                                      novel-nve-miR-42-a_guide
5'- aaaacauucauccacgaggacaaaguccauuccucacucuguucucgugggugaaugauuuuuuuuuuuu   -3'    exp
    ....((((((((((((((((((.(((.........))).)))))))))))))))))).............    reads     mm        sample
    ......uucauccacgaggacaaagu............................................    1         0         seq
    ............................auuccucacucuguucucgA......................    1         1         seq
    .....................................ucuguucucgugggugaU...............    1         1         seq
    .....................................ucuguucucgugggugaa...............    1         0         seq
    .....................................ucuguucucgugggugaau..............    9         0         seq
    .....................................ucuguucucgugggugaaG..............    1         1         seq
    .....................................ucuguucCcgugggugaau..............    1         1         seq
    .....................................ucuguucucgugggugaaC..............    4         1         seq
    .....................................ucuguucucgugggugaaA..............    1         1         seq
    .....................................ucuguucucguggCugaau..............    1         1         seq
    .....................................ucuguucucgugggugaUu..............    1         1         seq
    .....................................ucuguucucgugggugGau..............    1         1         seq
    .....................................ucuguucucgugggugaauU.............    5         1         seq
    .....................................ucuguucucgugggugaaug.............    3         0         seq
    .....................................ucuguucucgugggugaauA.............    2         1         seq
    .....................................ucuguucucgugggugaauC.............    2         1         seq
    .....................................ucuguucucgugggugaauga............    6         0         seq
    .....................................ucuguucucgugggugaaugU............    1         1         seq
    .....................................ucuguAcucgugggugaaugau...........    1         1         seq
    .....................................ucuguucucgugggugaaugau...........    35        0         seq
    .....................................ucuguucucgugggugaaugaG...........    1         1         seq
    .....................................uAuguucucgugggugaaugau...........    1         1         seq
    .....................................uUuguucucgugggugaaugau...........    1         1         seq
    .....................................ucuguucucgugggCgaaugau...........    1         1         seq
    .....................................ucuguucucgugggugaaugaC...........    8         1         seq
    .....................................ucuguucucAugggugaaugauu..........    1         1         seq
    .....................................ucuguucucgugggugaaugauu..........    11        0         seq
    .....................................ucuguucucgugggugaaugauC..........    3         1         seq
    .....................................ucuguucucgugggugaaugauAu.........    1         1         seq
    .....................................ucuguucucgugggugaaugauuu.........    1         0         seq
```

miRBase precursor : novel-nve-miR-42-b
Total read count : 435
novel-nve-miR-42-b_guide read count : 434
novel-nve-miR-42-b_star read count : 1
remaining reads : 0

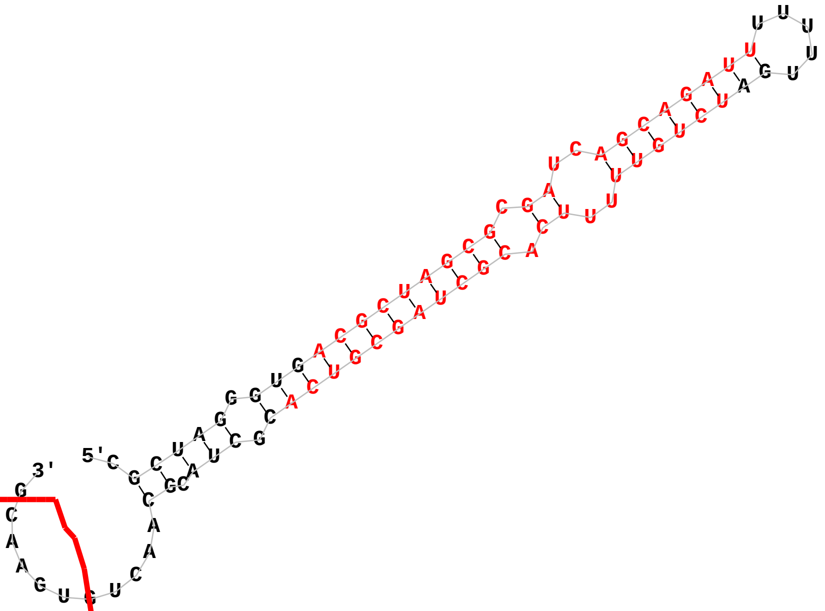


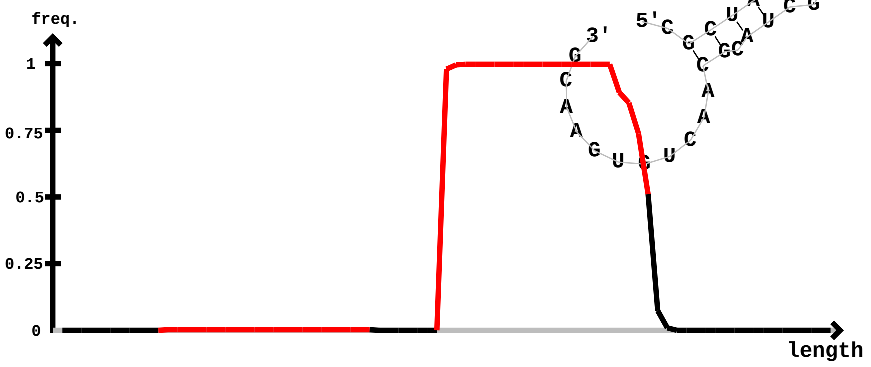


```
                                     novel-nve-miR-42-b_guide
           novel-nve-miR-42-b_star
5'- cgcuagggugacgcuagcgcgaucagcagauuuuuugaucuguuuuucacgcuagcgucacgcuacgcaacugugaacg   -3'    exp
    .(((((.((((((((((((.((..((((((((.....))))))))..)).))))))))))))).))).))...........    reads      mm         sample
    ..........acgcuagcgcgaucagcagauu................................................    1          0          seq
    .......................................ucuguuuuucacgcuagc.......................    30         0          seq
    .......................................ucuguuuuucacgcuagU.......................    12         1          seq
    .......................................ucuguuuCucacgcuagc.......................    1          1          seq
    .......................................ucuguuuuucacgcuagA.......................    2          1          seq
    .......................................ucuguuuuucaAgcuagc.......................    1          1          seq
    .......................................ucuguuuuucacgcuagcU......................    1          1          seq
    .......................................ucAguuuuucacgcuagcg......................    1          1          seq
    .......................................ucuguuuuucacgcuagcA......................    4          1          seq
    .......................................ucugAuuuucacgcuagcg......................    1          1          seq
    .......................................ucuguuuuucacgcuagcg......................    10         0          seq
    .......................................ucuguuuuucGcgcuagcgu.....................    1          1          seq
    .......................................ucuguuCuucacgcuagcgu.....................    1          1          seq
    .......................................ucuguuuuucacgcuagcgC.....................    5          1          seq
    .......................................uUuguuuuucacgcuagcgu.....................    1          1          seq
    .......................................ucuguAuuucacgcuagcgu.....................    1          1          seq
    .......................................ucuguuuuucacgcuagcgG.....................    2          1          seq
    .......................................ucuguuuuucacgcuagcgu.....................    35         0          seq
    .......................................ucuguuuuucacgcuagcgA.....................    2          1          seq
    .......................................ucuUuuuuucacgcuagcguc....................    1          1          seq
    .......................................Ccuguuuuucacgcuagcguc....................    1          1          seq
    .......................................ucuguuuuucacUcuagcguc....................    1          1          seq
    .......................................ucuguuuuucacgcuagcguA....................    4          1          seq
    .......................................ucuguuuuuAacgcuagcguc....................    1          1          seq
    .......................................ucuguuuuucacgcuagcguc....................    62         0          seq
    .......................................ucuguuuuCcacgcuagcguc....................    1          1          seq
    .......................................ucuguuuuucacgcuagcguU....................    24         1          seq
    .......................................ucuguuuuucacgcuagcgCc....................    1          1          seq
    .......................................ucuguuAuucacgcuagcguca...................    1          1          seq
    .......................................ucuguuuuucacgcuagcgucC...................    2          1          seq
    .......................................ucuguuuuCcacgcuagcguca...................    1          1          seq
    .......................................ucuguuuuucacAcuagcguca...................    2          1          seq
    .......................................ucuguuCuucacgcuagcguca...................    2          1          seq
    .......................................ucuguuuuucacgcuagcgCca...................    2          1          seq
```

novel-nve-miR-42-b_guide

novel-nve-miR-42-b_star

cgcuagggugacgcuagcgcgaucagcagauuuuuugaucuguuuuucacgcuagcgucacgcuacgcaacugugaacg

```
......................................ucuguuuuucacgcGagcguca...................   1     1   seq
......................................ucuguuuuucacgcuagcguca...................   154   0   seq
......................................ucugGuuuucacgcuagcguca...................   1     1   seq
......................................ucuguuuuucGcgcuagcguca...................   1     1   seq
......................................ucuguuuuucaAgcuagcguca...................   1     1   seq
......................................ucugCuuuucacgcuagcguca...................   1     1   seq
......................................ucuguuuuucacgcuagcgucU...................   13    1   seq
......................................Ccuguuuuucacgcuagcguca...................   1     1   seq
......................................ucuguuuuucacgcuagcgucG...................   5     1   seq
......................................ucuguuuuucacgcuagcgucaA..................   2     1   seq
......................................ucuguuuuucacgcuagcgucac..................   7     0   seq
......................................ucuguuuuucacgcuagcgucaU..................   18    1   seq
......................................ucuguuuuucacgcuagcgucacC.................   1     1   seq
......................................ucuguuuuucacgcuagcgucacU.................   2     1   seq
......................................ucuguuuuucacgcuagcgucacA.................   1     1   seq
.......................................cuguuuuucacgcuagUgu.....................   1     1   seq
.......................................cuguuuuucacgcuagcgu.....................   1     0   seq
.......................................cuguuuuucacgcuagcguA....................   1     1   seq
.......................................cuguuuuucacgcuagcguc....................   2     0   seq
.......................................cuguuuCucacgcuagcguca...................   1     1   seq
.......................................cuguuuuucacgcuagcgucU...................   1     1   seq
........................................uguuuuucacgcuagcgucaU..................   1     1   seq
```

miRBase precursor : novel-nve-miR-43
Total read count : 81
novel-nve-miR-43_guide read count : 81
novel-nve-miR-43_star read count : 0
remaining reads : 0

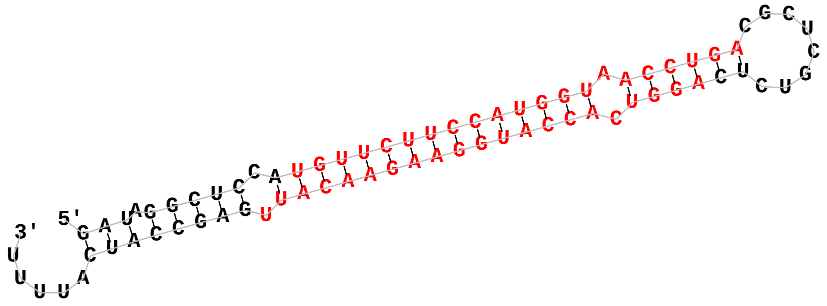

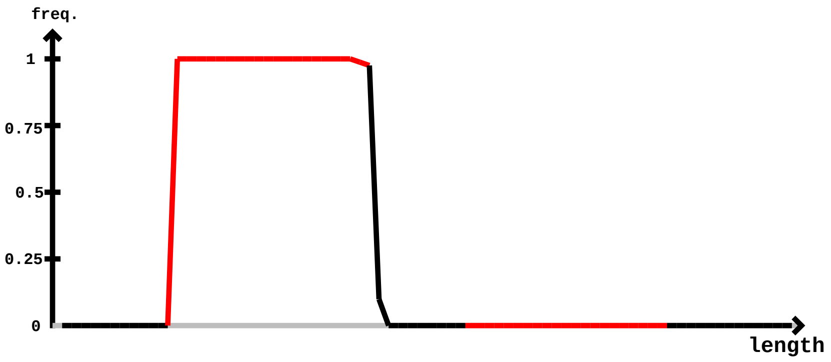


```
                                                  novel-nve-miR-43_star
                novel-nve-miR-43_guide
5'- gauaggcuccauguucuuccaugguaaccugacgcucgucucaggucaccauggaagaacauugagccaucauuuu   -3'    exp
    (((.(((((.(((((((((((((((.((((((........)))))).))))))))))))))))).)))))))).....    reads      mm         sample
    ...........uguucuuccaugguaaccu..............................................    1          0          seq
    ...........uguucuuccaugguaaccug.............................................    1          0          seq
    ...........uguucuuccaugguaaccugU............................................    1          1          seq
    ...........uguucCuccaugguaaccuga............................................    2          1          seq
    ...........uguucuuccaugguaaccugG............................................    1          1          seq
    ...........uguucuuccaugguaaccuga............................................    64         0          seq
    ...........uguuAuccaugguaaccuga.............................................    1          1          seq
    ...........uguucuuccaugguaaccCga............................................    2          1          seq
    ...........uguucuuccaugguaaccugac...........................................    2          0          seq
    ...........uguucuuccaugguaaccugaU...........................................    6          1          seq
```

miRBase precursor : novel-nve-miR-46
Total read count : 5578
novel-nve-miR-46_guide read count : 2789
novel-nve-miR-47-1_star read count : 2782
novel-nve-miR-46_star read count : 3
remaining reads : 4

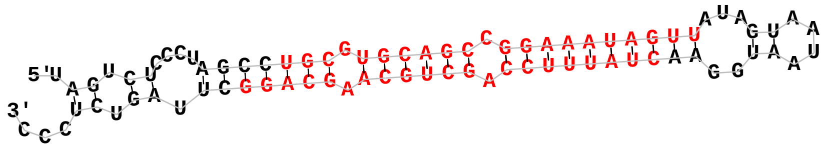

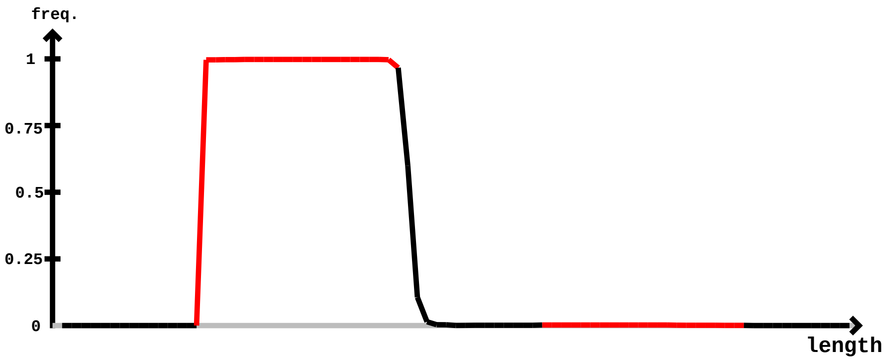


```
                                                               novel-nve-miR-46_star
                   novel-nve-miR-47-1_star
                   novel-nve-miR-46_guide
5'- uagucuccuagccugcgugcagccggaaauaguuauaguaauaauggaacuauuuccagcugcaagcaggcuuagucuccc   -3'    exp
    .((.((....(((((((.((((((.((((((((((...((....))..)))))))))).)))))).))))))).)).))..    reads      mm         sample
    .............ugcgugcagccggaaauag.................................................    2          0          seq
    .............ugcgugUagccggaaauag.................................................    1          1          seq
    .............ugcgugcagccggaaauaAu................................................    3          1          seq
    .............ugcgugcagccggaaauagu................................................    66         0          seq
    .............ugcgugcagccggaaauagC................................................    5          1          seq
    .............uAcgugcagccggaaauagu................................................    1          1          seq
    .............ugcgugcagccggaGauagu................................................    1          1          seq
    .............ugcgugcagccggaaaCagu................................................    5          1          seq
    .............ugcgugcagccggaaauagA................................................    1          1          seq
    .............ugcgugcagccggaaauagG................................................    2          1          seq
    .............ugcgugcagGcggaaauaguu...............................................    2          1          seq
    .............Agcgugcagccggaaauaguu...............................................    6          1          seq
    .............ugcgAgcagccggaaauaguu...............................................    1          1          seq
    .............ugcCugcagccggaaauaguu...............................................    2          1          seq
    .............ugcgugcGgccggaaauaguu...............................................    1          1          seq
    .............ugcgugcagccggaaauaguC...............................................    138        1          seq
    .............ugcgugcagccggaaauUguu...............................................    1          1          seq
    .............ugcgugcagccggaCauaguu...............................................    1          1          seq
    .............ugcgugcagccggGaauaguu...............................................    5          1          seq
    .............ugcgugcagccggaaGuaguu...............................................    3          1          seq
    .............ugAgugcagccggaaauaguu...............................................    1          1          seq
    .............uUcgugcagccggaaauaguu...............................................    3          1          seq
    .............ugcgugcagccggaaauGguu...............................................    1          1          seq
    .............ugcgugUagccggaaauaguu...............................................    4          1          seq
    .............ugcgugcagccggaaaAaguu...............................................    1          1          seq
    .............ugcgugcagccggaaaCaguu...............................................    1          1          seq
    .............ugcgugcagccggaaauaguu...............................................    805        0          seq
    .............ugcgugcagccggaaauaAuu...............................................    2          1          seq
    .............ugcAugcagccggaaauaguu...............................................    2          1          seq
    .............ugcUugcagccggaaauaguu...............................................    1          1          seq
    .............ugcguCcagccggaaauaguu...............................................    1          1          seq
    .............ugcgugcagccggaaauagGu...............................................    1          1          seq
    .............Cgcgugcagccggaaauaguu...............................................    2          1          seq
```

novel-nve-miR-46_star

novel-nve-miR-47-1_star

novel-nve-miR-48_star

uagucucccuagccugcgugcagccggaaauaguuauaguaauaauggaacuauuuccagcugcaagcaggcuuagucuccc

| | | | |
|---|---|---|---|
| ..............ugcgugcagccggaaauagAu............................................... | 3 | 1 | seq |
| ..............uAcgugcagccggaaauaguu............................................... | 1 | 1 | seq |
| ..............ugcgugcagcGggaaauaguu............................................... | 1 | 1 | seq |
| ..............ugcgugcagccUgaaauaguu............................................... | 3 | 1 | seq |
| ..............ugcgCgcagccggaaauaguu............................................... | 4 | 1 | seq |
| ..............ugcgugcagccggaaUuaguu............................................... | 1 | 1 | seq |
| ..............ugcgugcagccggaGauaguu............................................... | 7 | 1 | seq |
| ..............Ggcgugcagccggaaauaguu............................................... | 2 | 1 | seq |
| ..............ugcgugcagccggaaauagCu............................................... | 2 | 1 | seq |
| ..............ugGgugcagccggaaauaguu............................................... | 1 | 1 | seq |
| ..............ugcguAcagccggaaauaguu............................................... | 2 | 1 | seq |
| ..............ugcgugcagccggaUauaguu............................................... | 1 | 1 | seq |
| ..............ugcgugcagccgAaaauaguu............................................... | 2 | 1 | seq |
| ..............ugcgugcagccggaaauaguA............................................... | 12 | 1 | seq |
| ..............ugcgugGagccggaaauaguua.............................................. | 1 | 1 | seq |
| ..............ugcgugcagccggaaauaguuC.............................................. | 74 | 1 | seq |
| ..............ugcgugcagccggaaaAaguua.............................................. | 1 | 1 | seq |
| ..............ugcgugcagccggaaUuaguua.............................................. | 1 | 1 | seq |
| ..............ugcgugcagUcggaaauaguua.............................................. | 4 | 1 | seq |
| ..............ugcgugcagccggaaauGguua.............................................. | 2 | 1 | seq |
| ..............ugcgugcUgccggaaauaguua.............................................. | 2 | 1 | seq |
| ..............ugcgugcagccggaaauaguua.............................................. | 1106 | 0 | seq |
| ..............ugcgugcagccgCaaauaguua.............................................. | 2 | 1 | seq |
| ..............ugcgugcagccUgaaauaguua.............................................. | 1 | 1 | seq |
| ..............Ggcgugcagccggaaauaguua.............................................. | 1 | 1 | seq |
| ..............ugcgugAagccggaaauaguua.............................................. | 1 | 1 | seq |
| ..............ugcgugcagcGggaaauaguua.............................................. | 1 | 1 | seq |
| ..............ugcgugcagccggaaaCaguua.............................................. | 1 | 1 | seq |
| ..............Agcgugcagccggaaauaguua.............................................. | 10 | 1 | seq |
| ..............ugcgugcagccggaaauUguua.............................................. | 1 | 1 | seq |
| ..............ugcgugcagccggaUauaguua.............................................. | 1 | 1 | seq |
| ..............ugcgCgcagccggaaauaguua.............................................. | 8 | 1 | seq |
| ..............ugcgAgcagccggaaauaguua.............................................. | 5 | 1 | seq |
| ..............ugcgugcagcAggaaauaguua.............................................. | 2 | 1 | seq |
| ..............uAcgugcagccggaaauaguua.............................................. | 2 | 1 | seq |
| ..............ugcgugcagccggaaauagCua.............................................. | 2 | 1 | seq |
| ..............ugcgugcagccggaGauaguua.............................................. | 7 | 1 | seq |
| ..............ugcgugcagccggaaauaguCa.............................................. | 4 | 1 | seq |
| ..............ugcgugcagccggaaauaguuU.............................................. | 80 | 1 | seq |
| ..............ugcgugcagccAgaaauaguua.............................................. | 1 | 1 | seq |
| ..............ugcgugcaAccggaaauaguua.............................................. | 2 | 1 | seq |
| ..............ugcgugcGgccggaaauaguua.............................................. | 6 | 1 | seq |
| ..............ugcgugUagccggaaauaguua.............................................. | 2 | 1 | seq |
| ..............ugcgugcagccggaaauaguGa.............................................. | 1 | 1 | seq |
| ..............ugcgugcagccggaaauaguuG.............................................. | 2 | 1 | seq |
| ..............ugUgugcagccggaaauaguua.............................................. | 7 | 1 | seq |
| ..............ugcgugcagccggaaauagAua.............................................. | 2 | 1 | seq |
| ..............ugcgugcagccggGaauaguua.............................................. | 5 | 1 | seq |
| ..............ugcgugcagccgUaaauaguua.............................................. | 1 | 1 | seq |
| ..............ugcgugcagccgAaaauaguua.............................................. | 1 | 1 | seq |
| ..............Cgcgugcagccggaaauaguua.............................................. | 8 | 1 | seq |
| ..............ugcAugcagccggaaauaguua.............................................. | 3 | 1 | seq |
| ..............ugcgugcagccggaaauaAuua.............................................. | 2 | 1 | seq |
| ..............ugcgugcagcUggaaauaguua.............................................. | 4 | 1 | seq |
| ..............ugcgugcagccggaaauagGua.............................................. | 1 | 1 | seq |
| ..............ugcgugcagccggaaGuaguua.............................................. | 10 | 1 | seq |
| ..............ugcgugcagccggaaauaguuCu............................................. | 17 | 1 | seq |
| ..............ugcgugUagccggaaauaguuau............................................. | 1 | 1 | seq |
| ..............ugcgugcagccggaaauaguuUu............................................. | 5 | 1 | seq |
| ..............ugcgugcagccggaaauagCuau............................................. | 1 | 1 | seq |
| ..............ugcgAgcagccggaaauaguuau............................................. | 1 | 1 | seq |
| ..............ugcguCcagccggaaauaguuau............................................. | 1 | 1 | seq |
| ..............ugcgugcagccAgaaauaguuau............................................. | 2 | 1 | seq |
| ..............Cgcgugcagccggaaauaguuau............................................. | 1 | 1 | seq |
| ..............ugcgugcagccggaaauaguuaG............................................. | 3 | 1 | seq |
| ..............ugUgugcagccggaaauaguuau............................................. | 1 | 1 | seq |
| ..............ugcgugcagccggaaauUguuau............................................. | 1 | 1 | seq |
| ..............ugcgugcagccUgaaauaguuau............................................. | 1 | 1 | seq |

```
                                                  novel-nve-miR-46_star
              novel-nve-miR-47-1_star
              novel-nve-miR-46_guide
uagucucccuagcc[illegible]uaguaauaauggaacuauuuccagcugcaagcaggcuuagucuccc

..............ugcgugcagccggaaauaguuaC.............................................   54    1    seq
..............ugcguAcagccggaaauaguuau.............................................   1     1    seq
..............ugcgugcagccggaaauaguuau.............................................   143   0    seq
..............ugcgugcagccggaaauGguuau.............................................   1     1    seq
..............ugcgugcagccggaaauaguuaA.............................................   19    1    seq
..............ugcgugcagccggGaauaguuau.............................................   1     1    seq
..............ugcgugcagccggaaauaguuaCa............................................   4     1    seq
..............ugcgugcagccggaaauaguuauC............................................   3     1    seq
..............ugcgugcagccggaaauaguuaua............................................   7     0    seq
..............ugcgugcagccggaaauaguuaAa............................................   2     1    seq
..............ugcgugcagccggaaauCguuaua............................................   1     1    seq
..............ugcgugcagccggaaauaguuauU............................................   14    1    seq
..............ugcgugcagccggaaauaguuauaC...........................................   1     1    seq
..............ugcgugcagccggaaauaguuauaAu..........................................   4     1    seq
..............ugcgugcagccggaaauaguuauaCu..........................................   3     1    seq
..............ugcgugcagccggaaauaguuauaUua.........................................   1     1    seq
................cgugcagccggaaauaguuaC.............................................   1     1    seq
................cgugcagccggaaauaguuau.............................................   1     0    seq
..................ugcagccggaaauagAua..............................................   2     1    seq
..........................................auaauggaacuauuuccGgcug..................   2     1    seq
...........................................uaauggaacuauuuccGgcugc.................   1     1    seq
..................................................acuauuuccagcugcaCgc..............   1     1    seq
..................................................acuauuuccagcugcaCgcagg...........   2     1    seq
```

| | | |
|---|---|---|
| Provisional ID | | novel-nve-miR-47-1 |
| Score total | : | 850.4 |
| Score for star read(s) | : | 3.9 |
| Score for read counts | : | 839.8 |
| Score for mfe | : | 2.1 |
| Score for randfold | : | 1.6 |
| Score for cons. seed | : | 3 |
| Total read count | : | 1659 |
| Mature read count | : | 1654 |
| Loop read count | : | 0 |
| Star read count | : | 5 |

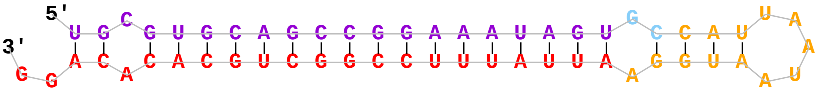


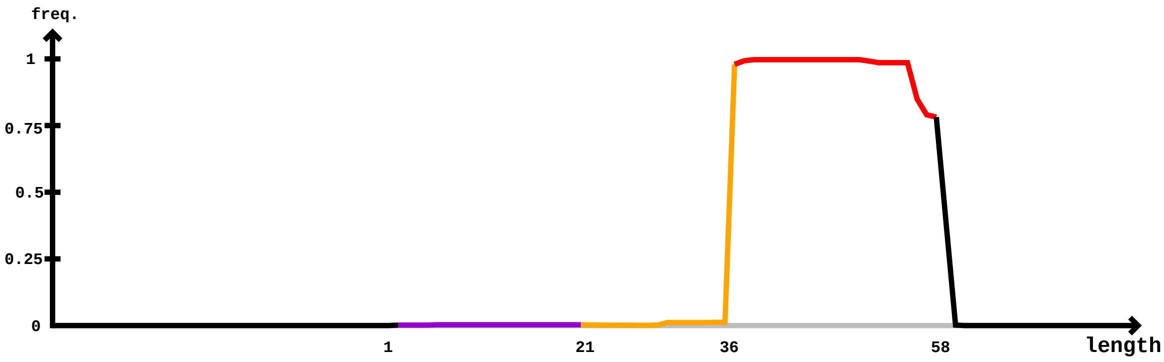


**Star** **Mature**

```
5'- ucucaacauaacacaguuucaguuuauuuuagccugcgugcagccggaaauagugccauuaauaauggaauuauuuccggcugcacacaggcuucauuaauaguuaaauaaa   -3'   obs
    ucucaacauaacacaguuucaguuuauuuuagccugcgugcagccggaaauagugccauuaauaauggaauuauuuccggcugcacacaggcuucauuaauaguuaaauaaa         exp
    .......................(((((..((((((.(((((((((((((((((.((((.....)))).))))))))))))))))).)))))).....))))).........   reads   mm   sample
    ..................................ugcgugcagccggaaauagu..........................................................   1       0    seq
    ..................................ugcgugcagccggaaauagG..........................................................   1       1    seq
    ..................................ugUgugcagccggaaauagugc........................................................   1       1    seq
    ......................................ugcagccggaaauaguCccau.....................................................   1       1    seq
    ......................................ugcagccggaaauagugccauua...................................................   1       0    seq
    ..............................................................auaauggaaCuauuuccggcug............................   3       1    seq
    ...............................................................uaauggaaCuauuuccggcug............................   6       1    seq
    ...............................................................uaauggaaCuauuuccggcugc...........................   10      1    seq
    ....................................................................Aaauuauuuccggcugcacacagg....................   1       1    seq
    ......................................................................auuauuuccggcugcacaA.......................   8       1    seq
    ......................................................................auuauuuccggcugcaNac.......................   1       1    seq
    ......................................................................auuauuuccggcuUcacac.......................   1       1    seq
    ......................................................................auuauAuccggcugcacac.......................   1       1    seq
    ......................................................................auuauuuccggcugcacaU.......................   66      1    seq
    ......................................................................auuauuuccggcugcacac.......................   145     0    seq
    ......................................................................auuauuuAcggcugcacac.......................   1       1    seq
    ......................................................................Guuauuuccggcugcacac.......................   3       1    seq
    ......................................................................auuauuCccggcugcacac.......................   1       1    seq
    ......................................................................auuauuuccggcAgcacac.......................   1       1    seq
    ......................................................................auuauuuccggcugcGcac.......................   1       1    seq
    ......................................................................auuauuuccggcugcacacU......................   4       1    seq
    ......................................................................auuauuuccggcugcacacG......................   2       1    seq
    ......................................................................aAuauuuccggcugcacaca......................   1       1    seq
    ......................................................................auuauuuccggcCgcacaca......................   1       1    seq
    ......................................................................Guuauuuccggcugcacaca......................   1       1    seq
    ......................................................................auuauuuGcggcugcacaca......................   1       1    seq
    ......................................................................auCauuuccggcugcacaca......................   1       1    seq
    ......................................................................auuauuuccggcugcacaca......................   86      0    seq
    ......................................................................auuauuCccggcugcacaca......................   2       1    seq
    ......................................................................auuauuuccggcugcacacaA.....................   3       1    seq
    ......................................................................auuauuuccggcugcacacag.....................   10      0    seq
    ......................................................................auuauuuccggcugcacacaC.....................   1       1    seq
    ......................................................................auuauuuccggcugcacacaAg....................   1       1    seq
```

Star Mature

| | | | |
|---|---|---|---|
| ucucaacauaacacaguuucaguuuauuuuagccugcgugcagccggaaauagugccauuaauaauggaauuauuuccggcugcacacaggcuucauuaaauaguuaaauaaa | | | |
| .....................................................................auCauuuccggcugcacacagg...................... | 4 | 1 | seq |
| .....................................................................aAuauuuccggcugcacacagg...................... | 6 | 1 | seq |
| .....................................................................auuauuuccggcugcacacCgg...................... | 4 | 1 | seq |
| .....................................................................auuauuuccggcugcacacagU...................... | 44 | 1 | seq |
| .....................................................................auuauuuccggcuAcacacagg...................... | 1 | 1 | seq |
| .....................................................................auAauuuccggcugcacacagg...................... | 1 | 1 | seq |
| .....................................................................auuauuuccAgcugcacacagg...................... | 1 | 1 | seq |
| .....................................................................auuauuuccggcugcUcacagg...................... | 4 | 1 | seq |
| .....................................................................auuauuuccggcugUacacagg...................... | 3 | 1 | seq |
| .....................................................................Guuauuuccggcugcacacagg...................... | 14 | 1 | seq |
| .....................................................................auuauuuccggcCgcacacagg...................... | 3 | 1 | seq |
| .....................................................................auuauuucAggcugcacacagg...................... | 3 | 1 | seq |
| .....................................................................auuauuuccggUugcacacagg...................... | 2 | 1 | seq |
| .....................................................................auuauuuccggcugcacacUgg...................... | 3 | 1 | seq |
| .....................................................................auuaCuuccggcugcacacagg...................... | 1 | 1 | seq |
| .....................................................................auuauuuccggcugcacacagC...................... | 10 | 1 | seq |
| .....................................................................auuauuuccggcugcacacGgg...................... | 7 | 1 | seq |
| .....................................................................auuGuuuccggcugcacacagg...................... | 1 | 1 | seq |
| .....................................................................auuauCuccggcugcacacagg...................... | 2 | 1 | seq |
| .....................................................................auuauuuccggcGgcacacagg...................... | 1 | 1 | seq |
| .....................................................................auuauuuccggcugcacaUagg...................... | 1 | 1 | seq |
| .....................................................................auuauuuccggcugcaUacagg...................... | 6 | 1 | seq |
| .....................................................................auuauuuccggcugcacacagg...................... | 963 | 0 | seq |
| .....................................................................auuCuuuccggcugcacacagg...................... | 1 | 1 | seq |
| .....................................................................auuauuAccggcugcacacagg...................... | 2 | 1 | seq |
| .....................................................................auuauuuccggcugcacacagA...................... | 145 | 1 | seq |
| .....................................................................auuauuCccggcugcacacagg...................... | 5 | 1 | seq |
| .....................................................................auuauuuccggcugcacUcagg...................... | 2 | 1 | seq |
| .....................................................................auuauuuccggcAgcacacagg...................... | 1 | 1 | seq |
| .....................................................................auuauAuccggcugcacacagg...................... | 2 | 1 | seq |
| .....................................................................auuauuuccggcugcacacaUg...................... | 1 | 1 | seq |
| .....................................................................auuauuuccgAcugcacacagg...................... | 1 | 1 | seq |
| .....................................................................aGuauuuccggcugcacacagg...................... | 1 | 1 | seq |
| .....................................................................auuauuuccggcugcGcacagg...................... | 4 | 1 | seq |
| .....................................................................auuauuuccggAugcacacagg...................... | 2 | 1 | seq |
| .....................................................................auuaAuuccggcugcacacagg...................... | 2 | 1 | seq |
| .....................................................................Uuuauuuccggcugcacacagg...................... | 2 | 1 | seq |
| .....................................................................auuauuuccggcugcacacaggA..................... | 1 | 1 | seq |
| .....................................................................auuauuuccggcugcacacaggU..................... | 22 | 1 | seq |
| .....................................................................auuauuuccggcugcacacaggc..................... | 2 | 0 | seq |
| ......................................................................uuauuuccggcugcacacagg...................... | 7 | 0 | seq |
| ......................................................................uuauuuccggcugcacacagA...................... | 1 | 1 | seq |
| ......................................................................uuauuuccggcugcacacaggc..................... | 10 | 0 | seq |
| ......................................................................uuauuuccggcugcacacaggU..................... | 4 | 1 | seq |
| .......................................................................uauuuccggcugcacacagg...................... | 1 | 0 | seq |
| .......................................................................Aauuuccggcugcacacagg...................... | 1 | 1 | seq |
| .......................................................................uauuuccggcugcacacaggc..................... | 2 | 0 | seq |
| .......................................................................uauuuccggcugcacacaggcA.................... | 1 | 1 | seq |
| .......................................................................uauuuccggcugcacacaggcu.................... | 2 | 0 | seq |

| | | |
|---|---|---|
| Provisional ID | : | Novel-nve-miR-47-2 |
| Score total | : | 957.3 |
| Score for star read(s) | : | 3.9 |
| Score for read counts | : | 946.4 |
| Score for mfe | : | 2.4 |
| Score for randfold | : | 1.6 |
| Score for cons. seed | : | 3 |
| Total read count | : | 1868 |
| Mature read count | : | 1654 |
| Loop read count | : | 0 |
| Star read count | : | 214 |

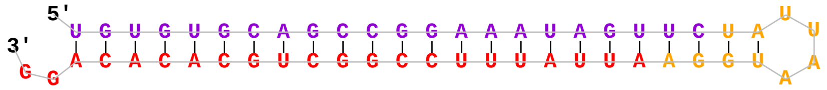


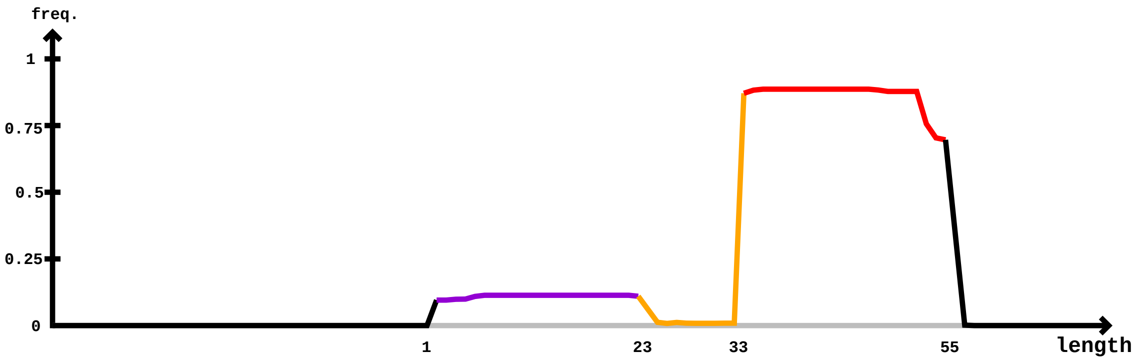


Star Mature

```
5'- ugucgaaacgcuacaaaaagccauggcaggucuaggccugugugcagccggaaauaguucuauuaauggaauuauuuccggcugcacacaggcuaagcaacgcuaaga   -3'   obs
    ugucgaaacgcuacaaaaagccauggcaggucuaggccugugugcagccggaaauaguucuauuaauggaauuauuuccggcugcacacaggcuaagcaacgcuaaga         exp
    .........(((......)))..((((..(.((.((((((((((((((((((((((((((((....)))))))))))))))))))))))))))).))).)...))))...    reads   mm   sample
    ......................................ugugugcagccggaaauGguu.................................................    1       1    seq
    ......................................ugugugcagccggaaauaguC.................................................    2       1    seq
    ......................................ugugugcagccggaaauaguu.................................................    4       0    seq
    ......................................ugugugcagccggaaauaguGc................................................    1       1    seq
    ......................................Agugugcagccggaaauaguuc................................................    2       1    seq
    ......................................ugugugcGgccggaaauaguuc................................................    4       1    seq
    ......................................ugugugcagcAggaaauaguuc................................................    1       1    seq
    ......................................ugugugcagccggaaauaguCc................................................    1       1    seq
    ......................................Cgugugcagccggaaauaguuc................................................    1       1    seq
    ......................................ugugugcagccggaaauaguuA................................................    7       1    seq
    ......................................ugugugcagccggaaauagCuc................................................    1       1    seq
    ......................................ugugugcagccggaUauaguuc................................................    1       1    seq
    ......................................ugugugcagcUggaaauaguuc................................................    1       1    seq
    ......................................ugugugcagcGggaaauaguuc................................................    1       1    seq
    ......................................ugugugcagccggaaauaguuc................................................    81      0    seq
    ......................................ugugugcagccggaaauaguuU................................................    14      1    seq
    ......................................Cgugugcagccggaaauaguucu...............................................    1       1    seq
    ......................................ugugugcagAcggaaauaguucu...............................................    1       1    seq
    ......................................ugugugcagccggaaGuaguucu...............................................    1       1    seq
    ......................................ugugugcagccggaaauaguucC...............................................    4       1    seq
    ......................................ugugugcagccggaaauaguucu...............................................    39      0    seq
    ......................................ugugugcagccggaaauaguucA...............................................    5       1    seq
    ......................................ugugugcagccggaaauaguucG...............................................    2       1    seq
    ......................................ugugugcagccggaaauaAuucua..............................................    1       1    seq
    ......................................ugugugcagccggaaauaguucuC..............................................    2       1    seq
    ......................................ugugugcagccggaaauaguucuU..............................................    1       1    seq
    ........................................ugugcagccggaaauaguucu...............................................    2       0    seq
    ........................................ugugcagccggaaauaguucua..............................................    3       0    seq
    ........................................ugugcagccggaaauaguucuau.............................................    1       0    seq
    .........................................gugcagccggaaauaguucA...............................................    1       1    seq
    ..........................................ugcagccggaaauaguuc................................................    2       0    seq
    ..........................................ugcagccggaaauaguucA...............................................    1       1    seq
    ..........................................ugcagccggaaauaguucu...............................................    9       0    seq
```

```
                                              Star                              Mature
ugucgaaacgcuacaaaaagccauggcaggucuaggccugugugcagccggaaauaguucuauuaauggaauuauuuccggcugcacacaggcuaagcaacgcuaaga
..........................................ugcagccggaaauaguucua..............................................	1	0	seq
..........................................ugcagccggaaauaguucuaC.............................................	1	1	seq
..........................................ugcagccggaaauaguucuau.............................................	1	0	seq
..........................................ugcagccggaaauaguucuauC............................................	2	1	seq
..........................................ugcagccggaaauaguucuauu............................................	1	0	seq
..........................................ugcagccggaaGuaguucuauua...........................................	1	1	seq
...........................................gcagccggaaauaguucuaC.............................................	1	1	seq
...........................................gcagccggaaauaguucuau.............................................	5	0	seq
...........................................gcagccggaaauaguucuauu............................................	2	0	seq
...............................................................uaauggaaCuauuuccggcug........................	6	1	seq
...............................................................uaauggaaCuauuuccggcugc.......................	10	1	seq
....................................................................Aaauuauuuccggcugcacacagg................	1	1	seq
......................................................................auuauuAcggcugcacac....................	1	1	seq
......................................................................auuauuuccggcugcGcac...................	1	1	seq
......................................................................auuauuuccggcuUcacac...................	1	1	seq
......................................................................auuauuCccggcugcacac...................	1	1	seq
......................................................................auuauuuccggcugcacaA...................	8	1	seq
......................................................................auuauAuccggcugcacac...................	1	1	seq
......................................................................auuauuuccggcugcacaU...................	66	1	seq
......................................................................Guuauuuccggcugcacac...................	3	1	seq
......................................................................auuauuuccggcugcaNac...................	1	1	seq
......................................................................auuauuuccggcugcacac...................	145	0	seq
......................................................................auuauuuccggcAgcacac...................	1	1	seq
......................................................................auuauuuccggcugcacacG..................	2	1	seq
......................................................................auuauuuccggcugcacaca..................	86	0	seq
......................................................................auuauuuccggcCgcacaca..................	1	1	seq
......................................................................auuauuuGcggcugcacaca..................	1	1	seq
......................................................................Guuauuuccggcugcacaca..................	1	1	seq
......................................................................auuauuCccggcugcacaca..................	2	1	seq
......................................................................auCauuuccggcugcacaca..................	1	1	seq
......................................................................auuauuuccggcugcacacU..................	4	1	seq
......................................................................aAuauuuccggcugcacaca..................	1	1	seq
......................................................................auuauuuccggcugcacacaC.................	1	1	seq
......................................................................auuauuuccggcugcacacag.................	10	0	seq
......................................................................auuauuuccggcugcacacaA.................	3	1	seq
......................................................................auuauuuccggcugcacacagg................	963	0	seq
......................................................................auuauuuccggcugcacacagU................	44	1	seq
......................................................................auuauAuccggcugcacacagg................	2	1	seq
......................................................................auuauCuccggcugcacacagg................	2	1	seq
......................................................................auuauuuccggUugcacacagg................	2	1	seq
......................................................................auuauuuccggcugcacacCgg................	4	1	seq
......................................................................auuauuuccggcugcaUacagg................	6	1	seq
......................................................................auAauuuccggcugcacacagg................	1	1	seq
......................................................................auuaAuuccggcugcacacagg................	2	1	seq
......................................................................auuauuuccggcugcacacaUg................	1	1	seq
......................................................................auuauuuccggcugcGcacagg................	4	1	seq
......................................................................auuauuuccggcugcacaUagg................	1	1	seq
......................................................................auuaCuuccggcugcacacagg................	1	1	seq
......................................................................auuauuuccAgcugcacacagg................	1	1	seq
......................................................................auuauuuccggcugcacacaAg................	1	1	seq
......................................................................auuauuuccggcGgcacacagg................	1	1	seq
......................................................................auuauuuccgAcugcacacagg................	1	1	seq
......................................................................auCauuuccggcugcacacagg................	4	1	seq
......................................................................auuauuuccggcuAcacacagg................	1	1	seq
......................................................................auuauuuccggAugcacacagg................	2	1	seq
......................................................................auuauuucAggcugcacacagg................	3	1	seq
......................................................................auuauuuccggcugcacacagA................	145	1	seq
......................................................................auuauuCccggcugcacacagg................	5	1	seq
......................................................................auuauuuccggcugcacUcagg................	2	1	seq
......................................................................auuCuuuccggcugcacacagg................	1	1	seq
......................................................................aGuauuuccggcugcacacagg................	1	1	seq
......................................................................auuauuuccggcugcacacagC................	10	1	seq
......................................................................auuauuuccggcAgcacacagg................	1	1	seq
......................................................................auuauuuccggcugcacacGgg................	7	1	seq
......................................................................Uuuauuuccggcugcacacagg................	2	1	seq
......................................................................aAuauuuccggcugcacacagg................	6	1	seq
......................................................................auuauuuccggcugUacacagg................	3	1	seq
......................................................................auuGuuuccggcugcacacagg................	1	1	seq
......................................................................auuauuuccggcugcacacUgg................	3	1	seq
......................................................................Guuauuuccggcugcacacagg................	14	1	seq
```

```
                                      Star                            Mature
ugucgaaacgcuacaaaaagccauggcaggucuaggccugugugcagccggaaauaguucuauuaauggaauuauuuccggcugcacacaggcuaagcaacgcuaaga
......................................................................auuauuAccggcugcacacagg................   2    1    seq
......................................................................auuauuuccggcugcUcacagg................   4    1    seq
......................................................................auuauuuccggcCgcacacagg................   3    1    seq
......................................................................auuauuuccggcugcacacaggU...............   22   1    seq
......................................................................auuauuuccggcugcacacaggA...............   1    1    seq
......................................................................auuauuuccggcugcacacaggc...............   2    0    seq
.......................................................................uuauuuccggcugcacacagg................   7    0    seq
.......................................................................uuauuuccggcugcacacagA................   1    1    seq
.......................................................................uuauuuccggcugcacacaggc...............   10   0    seq
.......................................................................uuauuuccggcugcacacaggU...............   4    1    seq
........................................................................uauuuccggcugcacacagg................   1    0    seq
........................................................................Aauuuccggcugcacacagg................   1    1    seq
........................................................................uauuuccggcugcacacaggc...............   2    0    seq
........................................................................uauuuccggcugcacacaggcA..............   1    1    seq
........................................................................uauuuccggcugcacacaggcu..............   2    0    seq
```

miRBase precursor : novel-nve-miR-48-1
Total read count : 2176
novel-nve-miR-48-1_guide read count : 2175
novel-nve-miR-48-1_star read count : 1
remaining reads : 0

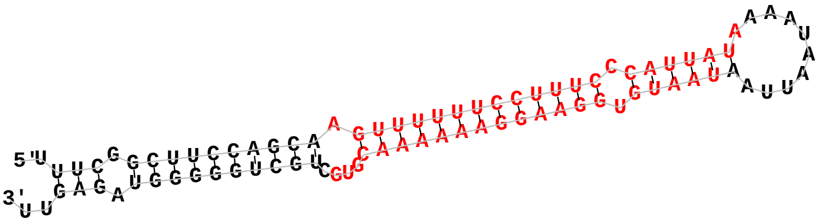

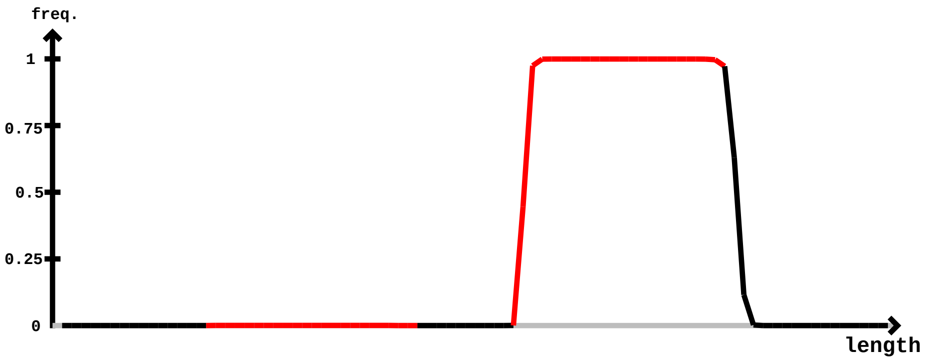


novel-nve-miR-48-1_star

novel-nve-miR-48-1_guide

| 5'- uuucggcuuccagcaaguuuuuuccuuucccauuauaaaauaauuaauaauguggaaggaaaaaacgugcugcuggggguagaguu -3' | exp | | |
|---|---|---|---|
| .(((.((((((((((.(((((((((((((.((((((..........)))))).)))))))))))))....)))))))))).))).. | reads | mm | sample |
| ...............aguuuuuuccuuucccauC.................................................... | 1 | 1 | seq |
| ..............................................auaauguggaaggaaaaaacgug................. | 1 | 0 | seq |
| ...............................................uaauguggaaggaaaaaacg................... | 1 | 0 | seq |
| ...............................................uaauguggaaggaaaaaacgG.................. | 4 | 1 | seq |
| ...............................................Gaauguggaaggaaaaaacgu.................. | 1 | 1 | seq |
| ...............................................uaauguggaaggaaaaaacgC.................. | 6 | 1 | seq |
| ...............................................uaauguggaaggaaaaGacgu.................. | 1 | 1 | seq |
| ...............................................uaauguggaaggaaaaaacgA.................. | 3 | 1 | seq |
| ...............................................uaauguggaaggaaaaaacgu.................. | 17 | 0 | seq |
| ...............................................uaauguggaaggaaaaaaUgug................. | 1 | 1 | seq |
| ...............................................uaauguggaagAaaaaaacgug................. | 1 | 1 | seq |
| ...............................................uaauguggaaggaaaaaGcgug................. | 3 | 1 | seq |
| ...............................................uaauguggaaggaaaaaacguU................. | 18 | 1 | seq |
| ...............................................uGauguggaaggaaaaaacgug................. | 1 | 1 | seq |
| ...............................................uaauguggaUggaaaaaacgug................. | 2 | 1 | seq |
| ...............................................uaauguggaaggaaaaaacAug................. | 1 | 1 | seq |
| ...............................................Aaauguggaaggaaaaaacgug................. | 9 | 1 | seq |
| ...............................................uaauguggaaggaaaaaacguC................. | 1 | 1 | seq |
| ...............................................uaauguggaaAgaaaaaacgug................. | 1 | 1 | seq |
| ...............................................uaauguggaaggGaaaaacgug................. | 3 | 1 | seq |
| ...............................................uaauguggaGggaaaaaacgug................. | 2 | 1 | seq |
| ...............................................uaGuguggaaggaaaaaacgug................. | 3 | 1 | seq |
| ...............................................uaauguAgaaggaaaaaacgug................. | 1 | 1 | seq |
| ...............................................uaauguggaaggaaGaaacgug................. | 3 | 1 | seq |
| ...............................................uaauguggaaggaaaaGacgug................. | 1 | 1 | seq |
| ...............................................uaauguggaaggaaaaaacgGg................. | 2 | 1 | seq |
| ...............................................uaauguggaaggaaaaaacgug................. | 498 | 0 | seq |
| ...............................................uaaCguggaaggaaaaaacgug................. | 1 | 1 | seq |
| ...............................................uaauguggaaggaaaaaacguA................. | 94 | 1 | seq |
| ...............................................uaauguggGaggaaaaaacgug................. | 6 | 1 | seq |
| ...............................................uaauguggaaggaaaaaacgCg................. | 9 | 1 | seq |
| ...............................................uaauguggaaggaaaGaacgug................. | 2 | 1 | seq |
| ...............................................uaauguggaaggaGaaaacgug................. | 3 | 1 | seq |
| ...............................................uaauguggaaggaaUaaacgug................. | 1 | 1 | seq |

```
               novel-nve-miR-48-1_star
                                               novel-nve-miR-48-1_guide
uuucggcuuccagcaaguuuuuuccuuucccauuauaaaauaauuaauaauguggaaggaaaaaacgugcugcuggggguagaguu

...............................................uaauguggaaggaaaUaacgug.................	1	1	seq
...............................................uaauguggaaggaaaaaaAgugc................	1	1	seq
...............................................uaauguggaaggaaaaaacguCc................	1	1	seq
...............................................Aaauguggaaggaaaaaacgugc................	1	1	seq
...............................................uaauguggaaggaaaaaacguUc................	1	1	seq
...............................................uaauguggaaggaaaaaacgugA................	7	1	seq
...............................................uaauguggaaggaaaaaacgCgc................	1	1	seq
...............................................uaauguggaaggGaaaaacgugc................	1	1	seq
...............................................uaaAguggaaggaaaaaacgugc................	1	1	seq
...............................................uaauguggaaggaGaaaacgugc................	1	1	seq
...............................................Caauguggaaggaaaaaacgugc................	2	1	seq
...............................................uaauguggaaUgaaaaaacgugc................	1	1	seq
...............................................uaauguggaaggaaaGaacgugc................	2	1	seq
...............................................uUauguggaaggaaaaaacgugc................	1	1	seq
...............................................uaGuguggaaggaaaaaacgugc................	1	1	seq
...............................................uaauguggaaggaaaaaacgugG................	3	1	seq
...............................................uGauguggaaggaaaaaacgugc................	1	1	seq
...............................................uaauguggaaggaaaaaacgugU................	79	1	seq
...............................................uaaCguggaaggaaaaaacgugc................	2	1	seq
...............................................uaauguggaaggaaaaaacgugc................	122	0	seq
...............................................uaaugAggaaggaaaaaacgugc................	1	1	seq
...............................................uaauguggaaggaaaaaacgugAu...............	1	1	seq
...............................................uaauguggaaggaaaaaacgugcA...............	5	1	seq
...............................................uaauguggaaggaaaaaacgugcC...............	5	1	seq
...............................................uaauguggaaggaaaaaGcgugcu...............	2	1	seq
...............................................uaauguggaaggaaaaaacgugcu...............	22	0	seq
...............................................uaauguggaaggaaaaaacgugcuA..............	1	1	seq
...............................................uaauguggaaggaaaaaacgugcuC..............	1	1	seq
................................................aauguggaaggaaaaaac....................	1	0	seq
................................................aauguggaaggaaaaaacA...................	2	1	seq
................................................aauguggaaggaaaaaacg...................	2	0	seq
................................................aauguggaaggaaaaaacgu..................	11	0	seq
................................................aauguggaaggaaaaaacgC..................	6	1	seq
................................................aauguggaaggaaaaaacgG..................	2	1	seq
................................................aauguggaGggaaaaaacgu..................	1	1	seq
................................................aauguggaaggaaaaaaAgug.................	1	1	seq
................................................aauguggaaggaaaaaacgCg.................	1	1	seq
................................................aauguggaaggaaaaaacguA.................	8	1	seq
................................................aauguggaaggaaaaaacguC.................	2	1	seq
................................................aauguggaaggaaGaaacgug.................	1	1	seq
................................................aauguggaaggaaaaaacgGg.................	1	1	seq
................................................aauguggaaggaaaaaacgug.................	46	0	seq
................................................aauguggaaggaaaGaacgug.................	1	1	seq
................................................aauguggaaggaaaaaaUgug.................	1	1	seq
................................................aauguggaaggaaaaaacguU.................	2	1	seq
................................................aaugugUaaggaaaaaacgug.................	1	1	seq
................................................aauguggaagUaaaaaacgugc................	1	1	seq
................................................aauguggaaggaaaaaacgugG................	4	1	seq
................................................aauguggaaggaaaaaacgugc................	506	0	seq
................................................aaGguggaaggaaaaaacgugc................	1	1	seq
................................................aauguggaaggaUaaaacgugc................	1	1	seq
................................................aauguggaaggaaaaaGcgugc................	1	1	seq
................................................aauguggaaggaaaaaacAugc................	1	1	seq
................................................aauguggaaggaaaaaacgAgc................	1	1	seq
................................................aauguggaaggaaaaaCcgugc................	1	1	seq
................................................aauguggaaggGaaaaacgugc................	2	1	seq
................................................aauguggaGggaaaaaacgugc................	3	1	seq
................................................aauguggaaggaaaaaacgugU................	288	1	seq
................................................aCuguggaaggaaaaaacgugc................	1	1	seq
................................................Gauguggaaggaaaaaacgugc................	3	1	seq
................................................aauguggaaggaaaaaacgugA................	48	1	seq
................................................aaCguggaaggaaaaaacgugc................	1	1	seq
................................................aauguggaaggaaaaaacgCgc................	1	1	seq
................................................aauguggGaggaaaaaacgugc................	3	1	seq
................................................aauguggaaggaaaaaacguAc................	1	1	seq
................................................aauguggaagAaaaaaacgugc................	2	1	seq
................................................aauguCgaaggaaaaaacgugc................	2	1	seq
................................................aaugAggaaggaaaaaacgugc................	4	1	seq
................................................aauguggaaggaaaGaacgugc................	1	1	seq
```

novel-nve-miR-48-1_star

novel-nve-miR-48-1_guide

uuucggcuuccagcaaguuuuuuccuuucccauuauaaaauaauuaauaauguggaaggaaaaaacgugcugcuggggguagaguu

| | | | |
|---|---|---|---|
| ................................................aGuguggaaggaaaaaacgugc................ | 1 | 1 | seq |
| ................................................aauguggaaCgaaaaaacgugc................ | 1 | 1 | seq |
| ................................................aauguggaaggaaaaaacguCc................ | 1 | 1 | seq |
| ................................................aGuguggaaggaaaaaacgugcu............... | 1 | 1 | seq |
| ................................................aauguggaaggaaaaaacgugcA............... | 15 | 1 | seq |
| ................................................aauguAgaaggaaaaaacgugcu............... | 1 | 1 | seq |
| ................................................aaAguggaaggaaaaaacgugcu............... | 2 | 1 | seq |
| ................................................aauguggaaggaaaaaacgugGu............... | 1 | 1 | seq |
| ................................................aauguggaaggaaaaaacgAgcu............... | 1 | 1 | seq |
| ................................................aauguggaUggaaaaaacgugcu............... | 1 | 1 | seq |
| ................................................aauguggaaggaaaaaacgugcC............... | 30 | 1 | seq |
| ................................................aauguggGaggaaaaaacgugcu............... | 2 | 1 | seq |
| ................................................aauguggaaggaaaaaacgugcu............... | 113 | 0 | seq |
| ................................................aauguggaaggaaaaaacgugcG............... | 5 | 1 | seq |
| ................................................aauguggaaggaaaaaacgCgcu............... | 1 | 1 | seq |
| ................................................aauguggaaggaaaaaacgugAu............... | 3 | 1 | seq |
| ................................................aauguggaaggaGaaaacgugcu............... | 2 | 1 | seq |
| ................................................aauguggaagUaaaaaacgugcu............... | 1 | 1 | seq |
| ................................................aaugugUaaggaaaaaacgugcu............... | 1 | 1 | seq |
| ................................................aauguggaaggaaaaaacgugcuU.............. | 1 | 1 | seq |
| .................................................auguggaaggaaaaaacgu.................. | 1 | 0 | seq |
| .................................................auguggaaggaaaaaacgug................. | 7 | 0 | seq |
| .................................................auguggaaggaaaaaacguA................. | 1 | 1 | seq |
| .................................................auguggaaggaaaaaacguU................. | 2 | 1 | seq |
| .................................................auguggaaggaaaaaacguC................. | 1 | 1 | seq |
| .................................................auguggaaggaaaaaacgugA................ | 1 | 1 | seq |
| .................................................auguggaaggaaaaaacgGgc................ | 1 | 1 | seq |
| .................................................auguggaaggaaaaaacgugU................ | 2 | 1 | seq |
| .................................................aCguggaaggaaaaaacgugc................ | 1 | 1 | seq |
| .................................................auguggaaggaaaaaacgugc................ | 3 | 0 | seq |
| .................................................aAguggaaggaaaaaacgugc................ | 1 | 1 | seq |
| .................................................Cuguggaaggaaaaaacgugcu............... | 1 | 1 | seq |
| .................................................auguggaaggaaaaaacgugcG............... | 1 | 1 | seq |
| .................................................auguggaaggaaaaaacgugcu............... | 21 | 0 | seq |
| .................................................auguggaaggaaaaaacgugcC............... | 4 | 1 | seq |
| .................................................auguggaaggaaaaaacgugcA............... | 2 | 1 | seq |
| .................................................auguggaagAaaaaaacgugcu............... | 2 | 1 | seq |
| .................................................auguggaaggaaaaaacgugcuU.............. | 2 | 1 | seq |
| ..................................................uguggaaggaaaaaacgug................. | 1 | 0 | seq |

| | | |
|---|---|---|
| Provisional ID | | novel-nve-miR-48-2 |
| Score total | : | 2264.8 |
| Score for star read(s) | : | 3.9 |
| Score for read counts | : | 2258.1 |
| Score for mfe | : | 1.8 |
| Score for randfold | : | 1.6 |
| Score for cons. seed | : | -0.6 |
| Total read count | : | 4441 |
| Mature read count | : | 4402 |
| Loop read count | : | 0 |
| Star read count | : | 39 |

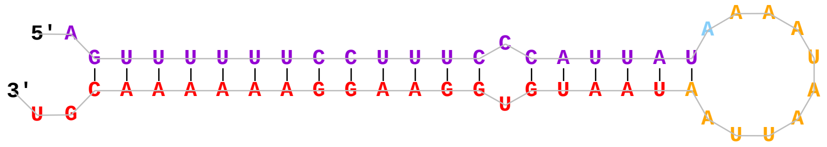


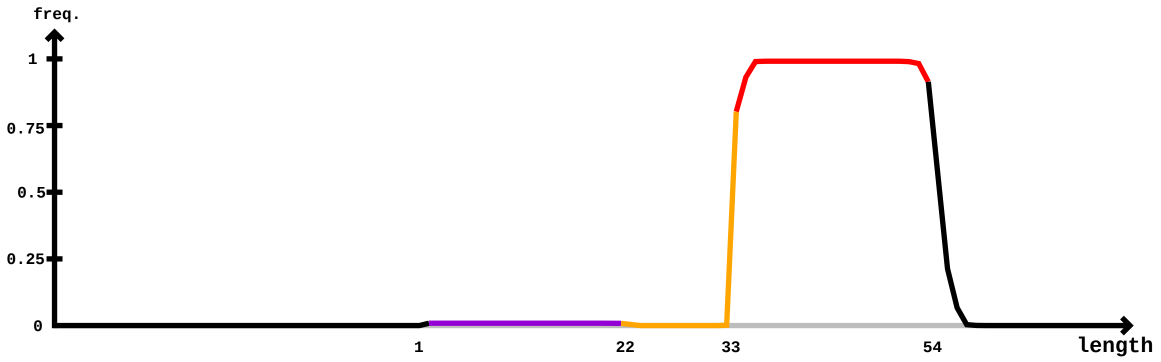


Star Mature

```
5'- caacuggccguauucgccuuguuuucggcuuccagcaaguuuuuuccuuucccauuauaaaauaauuaauaauguggaaggaaaaaacgugcugcuggggguagaguuac  -3'   obs
    caacuggccguauucgccuuguuuucggcuuccagcaaguuuuuuccuuucccauuauaaaauaauuaauaauguggaaggaaaaaacgugcugcuggggguagaguuac         exp
    .(((.(((.......)))..)))(((.((((((((((.(((((((((((((.((((((..........)))))).)))))))))))))....)))))))))).)))....   reads    mm    sample
    .....................................aguuuuuuccuuucccauC......................................................   1        1     seq
    .....................................aguuuuuuccuuucccauuG.....................................................   1        1     seq
    .....................................aguuuuuuccuuucccauuau....................................................   27       0     seq
    .....................................aguuuCuuccuuucccauuau....................................................   1        1     seq
    .....................................aguuuuuuccuuucccauuaC....................................................   5        1     seq
    .....................................aguuuuuuccuuucccaCuau....................................................   1        1     seq
    .....................................aCuuuuuuccuuucccauuau....................................................   1        1     seq
    .....................................aguuuuuuccuuucccauuaua...................................................   2        0     seq
    ....................................................................auaauguggaaggaaaaaac......................   1        0     seq
    ....................................................................auaauguggaaggaaaaaacU.....................   1        1     seq
    ....................................................................auaauguggaaggaaaaaacgu....................   1        0     seq
    ....................................................................auaauguggaaggaaaaaacguC...................   1        1     seq
    ....................................................................auaauguggaaggaaaaaacgugAu.................   1        1     seq
    .....................................................................uaauguggaaggaaaaaU.......................   1        1     seq
    .....................................................................uaauguggaaggaaaaaa.......................   5        0     seq
    .....................................................................uaauguggaaggaaaaaC.......................   1        1     seq
    .....................................................................uaauguggaaggaaaaaCc......................   1        1     seq
    .....................................................................uaauguggaaggaaaaaac......................   24       0     seq
    .....................................................................uaauguggaaCgaaaaaac......................   1        1     seq
    .....................................................................uaauguggaaggaaaaaaA......................   2        1     seq
    .....................................................................uaauguCgaaggaaaaaacg.....................   2        1     seq
    .....................................................................uaauguggaaggGaaaaacg.....................   1        1     seq
    .....................................................................uaauguggaaggaaaaaacU.....................   8        1     seq
    .....................................................................Caauguggaaggaaaaaacg.....................   1        1     seq
    .....................................................................uaaAguggaaggaaaaaacg.....................   2        1     seq
    .....................................................................uaauCuggaaggaaaaaacg.....................   1        1     seq
    .....................................................................uaauguggaaggaaaaaacC.....................   3        1     seq
    .....................................................................uaauguggaaggaaaaaacg.....................   227      0     seq
    .....................................................................Aaauguggaaggaaaaaacg.....................   3        1     seq
    .....................................................................uaaugGggaaggaaaaaacg.....................   2        1     seq
    .....................................................................uaauguggaaggaaaaaGcg.....................   1        1     seq
    .....................................................................uaauguggaaUgaaaaaacg.....................   1        1     seq
    .....................................................................uaauguggaaggaaaaGacg.....................   1        1     seq
```

Star                    Mature

| | | | |
|---|---|---|---|
| caacuggccguauucgccuuguuuucggcuuccagcaaguuuuuuccuuucccauuauaaaauaauuaauaauguggaaggaaaaaacgugcugcuggggguagaguuac | | | |
| .....................................................................uaauguggaaggaGaaaacg..................... | 1 | 1 | seq |
| .....................................................................uaauguggaaggaaaaaacA..................... | 30 | 1 | seq |
| .....................................................................uaauguggaagAaaaaaacg..................... | 1 | 1 | seq |
| .....................................................................uaauAuggaaggaaaaaacgu.................... | 2 | 1 | seq |
| .....................................................................uaauguggaagAaaaaaacgu.................... | 7 | 1 | seq |
| .....................................................................uaauguggaaggaaaaUacgu.................... | 1 | 1 | seq |
| .....................................................................uaaugGggaaggaaaaaacgu.................... | 1 | 1 | seq |
| .....................................................................uaauguggUaggaaaaaacgu.................... | 1 | 1 | seq |
| .....................................................................uaGuguggaaggaaaaaacgu.................... | 4 | 1 | seq |
| .....................................................................uaauguggaaggaaaaGacgu.................... | 4 | 1 | seq |
| .....................................................................uaauguggaaggaaaaaacCu.................... | 1 | 1 | seq |
| .....................................................................uaauguggaaggaaaGaacgu.................... | 5 | 1 | seq |
| .....................................................................uaaugCggaaggaaaaaacgu.................... | 4 | 1 | seq |
| .....................................................................uCauguggaaggaaaaaacgu.................... | 1 | 1 | seq |
| .....................................................................uaauguggaaggaaaaaaAgu.................... | 2 | 1 | seq |
| .....................................................................uaaugAggaaggaaaaaacgu.................... | 6 | 1 | seq |
| .....................................................................Aaauguggaaggaaaaaacgu.................... | 22 | 1 | seq |
| .....................................................................uaauguggaaggaaaaaacgG.................... | 42 | 1 | seq |
| .....................................................................uaauguggaaggaaaaaGcgu.................... | 4 | 1 | seq |
| .....................................................................uaauguggaagUaaaaaacgu.................... | 2 | 1 | seq |
| .....................................................................uaauguggaaggaaUaaacgu.................... | 3 | 1 | seq |
| .....................................................................uaauguggaaggaGaaaacgu.................... | 3 | 1 | seq |
| .....................................................................uaauguggaaggGaaaaacgu.................... | 5 | 1 | seq |
| .....................................................................uaauguggaaggaaaaaacAu.................... | 7 | 1 | seq |
| .....................................................................uaauguggaaggaUaaaacgu.................... | 2 | 1 | seq |
| .....................................................................uaauguggaaggaaaaaacgu.................... | 1397 | 0 | seq |
| .....................................................................uaauUuggaaggaaaaaacgu.................... | 2 | 1 | seq |
| .....................................................................uaauguggaaUgaaaaaacgu.................... | 1 | 1 | seq |
| .....................................................................uaauguggaaggaaaaCacgu.................... | 2 | 1 | seq |
| .....................................................................uaauguggaaggUaaaaacgu.................... | 2 | 1 | seq |
| .....................................................................Caauguggaaggaaaaaacgu.................... | 4 | 1 | seq |
| .....................................................................uGauguggaaggaaaaaacgu.................... | 2 | 1 | seq |
| .....................................................................uaauguggaaggaaaaaacgA.................... | 39 | 1 | seq |
| .....................................................................uaaCguggaaggaaaaaacgu.................... | 5 | 1 | seq |
| .....................................................................uaaAguggaaggaaaaaacgu.................... | 4 | 1 | seq |
| .....................................................................uaauguggaaggaaaaaCcgu.................... | 4 | 1 | seq |
| .....................................................................uaauguggaGggaaaaaacgu.................... | 8 | 1 | seq |
| .....................................................................uaauguggaaggaaaaaacUu.................... | 1 | 1 | seq |
| .....................................................................uaauguCgaaggaaaaaacgu.................... | 1 | 1 | seq |
| .....................................................................uaauguggGaggaaaaaacgu.................... | 6 | 1 | seq |
| .....................................................................Gaauguggaaggaaaaaacgu.................... | 4 | 1 | seq |
| .....................................................................uaaugugAaaggaaaaaacgu.................... | 2 | 1 | seq |
| .....................................................................uaauguggaaggaaGaaacgu.................... | 6 | 1 | seq |
| .....................................................................uaauguggaCggaaaaaacgu.................... | 1 | 1 | seq |
| .....................................................................uaauguggaaggGaaaaacgug................... | 3 | 1 | seq |
| .....................................................................Caauguggaaggaaaaaacgug................... | 2 | 1 | seq |
| .....................................................................uaauguggaagAaaaaaacgug................... | 6 | 1 | seq |
| .....................................................................uaaugAggaaggaaaaaacgug................... | 1 | 1 | seq |
| .....................................................................uaauguggaaggaaaaaacUug................... | 1 | 1 | seq |
| .....................................................................uaauguggUaggaaaaaacgug................... | 1 | 1 | seq |
| .....................................................................uaauguggaaggaaaaaacAug................... | 5 | 1 | seq |
| .....................................................................uaauguggaaggaaaaaacgAg................... | 1 | 1 | seq |
| .....................................................................uaauCuggaaggaaaaaacgug................... | 1 | 1 | seq |
| .....................................................................uaauguggaaAgaaaaaacgug................... | 1 | 1 | seq |
| .....................................................................uaauguUgaaggaaaaaacgug................... | 1 | 1 | seq |
| .....................................................................uaauguggaaggaaaaaacCug................... | 1 | 1 | seq |
| .....................................................................Aaauguggaaggaaaaaacgug................... | 11 | 1 | seq |
| .....................................................................uaaugugUaaggaaaaaacgug................... | 1 | 1 | seq |
| .....................................................................uaauguggaaggaGaaaacgug................... | 3 | 1 | seq |
| .....................................................................uaauguggaaggaaaaaCcgug................... | 3 | 1 | seq |
| .....................................................................uaaCguggaaggaaaaaacgug................... | 2 | 1 | seq |
| .....................................................................uaauguggaaggaaaaGacgug................... | 3 | 1 | seq |
| .....................................................................uaauguggaaggaaaaaacguC................... | 18 | 1 | seq |
| .....................................................................uaauguggaaggaaaaaacguU................... | 74 | 1 | seq |
| .....................................................................uaauguggaaggaaaGaacgug................... | 5 | 1 | seq |
| .....................................................................uGauguggaaggaaaaaacgug................... | 3 | 1 | seq |
| .....................................................................uaauguggaaggaaaaaacgGg................... | 6 | 1 | seq |
| .....................................................................uaauguggaaggaaaaaGcgug................... | 3 | 1 | seq |
| .....................................................................uaaugCggaaggaaaaaacgug................... | 1 | 1 | seq |
| .....................................................................uaauguggGaggaaaaaacgug................... | 3 | 1 | seq |

**Star** **Mature**

| | | | |
|---|---|---|---|
| caacuggccguauucgccuuguuuucggcuuccagcaaguuuuuuccuuucccauuauaaaauaauuaauaauguggaaggaaaaaacgugcugcuggggguagaguuac | | | |
| .....................................................................uaauguggaaUgaaaaaacgug................... | 1 | 1 | seq |
| .....................................................................uaauguAgaaggaaaaaacgug................... | 1 | 1 | seq |
| .....................................................................uaauguggaaggaaaaaUcgug................... | 1 | 1 | seq |
| .....................................................................uaauguggaaggaaaaaacgug................... | 797 | 0 | seq |
| .....................................................................uaauguggaaggaaaaaacguA................... | 122 | 1 | seq |
| .....................................................................uaauguggaGggaaaaaacgug................... | 2 | 1 | seq |
| .....................................................................uaauguggaaggUaaaaacgug................... | 1 | 1 | seq |
| .....................................................................uaauguggaaggaGaaaacgugc.................. | 1 | 1 | seq |
| .....................................................................uaauguggaaggaaaaaacgugU.................. | 124 | 1 | seq |
| .....................................................................uaauguAgaaggaaaaaacgugc.................. | 1 | 1 | seq |
| .....................................................................uaauguggaaggaaaaaacgugA.................. | 28 | 1 | seq |
| .....................................................................uaauguggGaggaaaaaacgugc.................. | 1 | 1 | seq |
| .....................................................................uaauguggaaggaaaaaacAugc.................. | 1 | 1 | seq |
| .....................................................................uaauguggaaggaaaaCacgugc.................. | 1 | 1 | seq |
| .....................................................................uaaugAggaaggaaaaaacgugc.................. | 1 | 1 | seq |
| .....................................................................uaauguggaaggaaaaaacgugG.................. | 7 | 1 | seq |
| .....................................................................uaauguggaaggaaaaaacgugc.................. | 224 | 0 | seq |
| .....................................................................uaauguggaaggUaaaaacgugc.................. | 1 | 1 | seq |
| .....................................................................uaauguggaGggaaaaaacgugc.................. | 1 | 1 | seq |
| .....................................................................uaauguggaaggGaaaaacgugc.................. | 1 | 1 | seq |
| .....................................................................uaauguggaaggaaaaaacguAc.................. | 1 | 1 | seq |
| .....................................................................uaauguggaaggaaaaaacgAgc.................. | 1 | 1 | seq |
| .....................................................................uaauguggUaggaaaaaacgugc.................. | 1 | 1 | seq |
| .....................................................................Caauguggaaggaaaaaacgugc.................. | 1 | 1 | seq |
| .....................................................................Aaauguggaaggaaaaaacgugc.................. | 2 | 1 | seq |
| .....................................................................uaaCguggaaggaaaaaacgugc.................. | 1 | 1 | seq |
| .....................................................................uaGuguggaaggaaaaaacgugc.................. | 1 | 1 | seq |
| .....................................................................uaauguggaagAaaaaaacgugc.................. | 1 | 1 | seq |
| .....................................................................uaauguggaaggaaaGaacgugc.................. | 1 | 1 | seq |
| .....................................................................uaauguggaaggaaaaaacgugcu................. | 53 | 0 | seq |
| .....................................................................uaauguggaaggaaaaaacgugcG................. | 4 | 1 | seq |
| .....................................................................uaaugAggaaggaaaaaacgugcu................. | 1 | 1 | seq |
| .....................................................................uaauguggaaggaaaaaacgugcC................. | 13 | 1 | seq |
| .....................................................................uaauguggaaggaaaaaacgugUu................. | 3 | 1 | seq |
| .....................................................................uUauguggaaggaaaaaacgugcu................. | 1 | 1 | seq |
| .....................................................................uaauguggaaggaaGaaacgugcu................. | 1 | 1 | seq |
| .....................................................................uaauguggaaggaaaaaacgugAu................. | 36 | 1 | seq |
| .....................................................................uaauguggaaggaaaaaacgugcA................. | 11 | 1 | seq |
| .....................................................................uaauguggaaggaaaGaacgugcu................. | 1 | 1 | seq |
| .....................................................................uaauguggaaggaaaaaacgugcuA................ | 2 | 1 | seq |
| .....................................................................uaauguggaaggaaaaaacgugAug................ | 1 | 1 | seq |
| .....................................................................uaauguggaaggaaaaaacgugcuU................ | 4 | 1 | seq |
| .....................................................................uaauguggaaggaaaaaacgugcug................ | 1 | 0 | seq |
| .....................................................................uaauguggaaggaaaaaacgugcuUc............... | 1 | 1 | seq |
| ......................................................................Uauguggaaggaaaaaac...................... | 1 | 1 | seq |
| ......................................................................aauguggaaggaaaaaac...................... | 2 | 0 | seq |
| ......................................................................aauguggaaggaaaaaacg..................... | 12 | 0 | seq |
| ......................................................................Gauguggaaggaaaaaacg..................... | 1 | 1 | seq |
| ......................................................................aauguggaaggaaaaaacA..................... | 2 | 1 | seq |
| ......................................................................aauguggaagAaaaaaacg..................... | 1 | 1 | seq |
| ......................................................................aauguggaaggaaaaaacgu.................... | 89 | 0 | seq |
| ......................................................................aauguggaaggaaaaaGcgu.................... | 1 | 1 | seq |
| ......................................................................aauguggaaggaaGaaacgu.................... | 1 | 1 | seq |
| ......................................................................aGuguggaaggaaaaaacgu.................... | 2 | 1 | seq |
| ......................................................................aauCuggaaggaaaaaacgu.................... | 1 | 1 | seq |
| ......................................................................Uauguggaaggaaaaaacgu.................... | 1 | 1 | seq |
| ......................................................................aauguggaaggaaaaaacgA.................... | 5 | 1 | seq |
| ......................................................................aaugAggaaggaaaaaacgu.................... | 1 | 1 | seq |
| ......................................................................aauguggaaggaaaaaacgG.................... | 1 | 1 | seq |
| ......................................................................aauguggaGggaaaaaacgu.................... | 1 | 1 | seq |
| ......................................................................aauguggaaggGaaaaacgu.................... | 1 | 1 | seq |
| ......................................................................aUuguggaaggaaaaaacgu.................... | 1 | 1 | seq |
| ......................................................................aauguggaUggaaaaaacgug................... | 1 | 1 | seq |
| ......................................................................Uauguggaaggaaaaaacgug................... | 1 | 1 | seq |
| ......................................................................aauguAgaaggaaaaaacgug................... | 1 | 1 | seq |
| ......................................................................aauguggaaggaaaaaaAgug................... | 1 | 1 | seq |
| ......................................................................aauguggaaggaaaaaacguU................... | 3 | 1 | seq |
| ......................................................................aauguggaaggaaaaaacgug................... | 131 | 0 | seq |
| ......................................................................aauguggGaggaaaaaacgug................... | 2 | 1 | seq |
| ......................................................................aaugugUaaggaaaaaacgug................... | 1 | 1 | seq |

```
                                    Star                          Mature
caacuggccguauucgccuuguuuucggcuuccagcaaguuuuuuccuuucccauuauaaaauaauuaauaauguggaaggaaaaaacgugcugcuggggguagaguuac
......................................................................aauguggaagAaaaaaacgug...................    2    1    seq
......................................................................aauguggaaggaaaaGacgug...................    2    1    seq
......................................................................aauguggaGggaaaaaacgug...................    1    1    seq
......................................................................aauguggaaggaaaaaacguA...................    9    1    seq
......................................................................aauguggaaggaaaaaacgugA..................    12   1    seq
......................................................................aauguggaaggaaaaaacgugU..................    41   1    seq
......................................................................aauguggaaggaaaGaacgugc..................    1    1    seq
......................................................................aauguAgaaggaaaaaacgugc..................    1    1    seq
......................................................................aauguggaaggaaaaaacgugc..................    131  0    seq
......................................................................aauguggaaggaaaaaGcgugc..................    1    1    seq
......................................................................aauguggaaggaaaaaacguCc..................    1    1    seq
......................................................................aauguggaaggaaGaaacgugc..................    1    1    seq
......................................................................aGuguggaaggaaaaaacgugc..................    1    1    seq
......................................................................aaugAggaaggaaaaaacgugc..................    1    1    seq
......................................................................aauguggaaggGaaaaacgugc..................    1    1    seq
......................................................................aauguggaaggaaaaaacgugG..................    3    1    seq
......................................................................aauguggaaggaaaaaCcgugc..................    2    1    seq
......................................................................aaAguggaaggaaaaaacgugc..................    1    1    seq
......................................................................aauguggaaggaaaaaacgugcC.................    14   1    seq
......................................................................aaAguggaaggaaaaaacgugcu.................    1    1    seq
......................................................................aauguggaaggaaaaaacgugcu.................    46   0    seq
......................................................................aauguggaaggaaaaGacgugcu.................    1    1    seq
......................................................................aauguggaaggaaaaaacgugAu.................    4    1    seq
......................................................................aauguggaaggaaaaaacgugcG.................    12   1    seq
......................................................................aaCguggaaggaaaaaacgugcu.................    1    1    seq
......................................................................Gauguggaaggaaaaaacgugcu.................    1    1    seq
......................................................................aauguggaagAaaaaaacgugcu.................    1    1    seq
......................................................................aauguggaaggaaaaaacgugcA.................    11   1    seq
......................................................................aauguggaaggaaaaaacgugcuA................    1    1    seq
......................................................................aauguggaaggaaaaaacgugcuUc...............    1    1    seq
.......................................................................auguggaaggaaaaaacg.....................    1    0    seq
.......................................................................auguggaaggaaaaGacgu....................    1    1    seq
.......................................................................auguggaaggaaGaaacgu....................    1    1    seq
.......................................................................auCuggaaggaaaaaacgu....................    1    1    seq
.......................................................................auguggaaggaaaGaacgu....................    2    1    seq
.......................................................................auguggaaggaaaaaacgu....................    29   0    seq
.......................................................................auguggaaggaaaaaacgA....................    2    1    seq
.......................................................................auguggaaggUaaaaacgug...................    1    1    seq
.......................................................................auguggaaggaaaaaacgug...................    81   0    seq
.......................................................................Guguggaaggaaaaaacgug...................    1    1    seq
.......................................................................auguggaaggaaGaaacgug...................    1    1    seq
.......................................................................auguggaaggaaaaaaCug...................    1    1    seq
.......................................................................augCggaaggaaaaaacgug...................    1    1    seq
.......................................................................auguggaaggaaaaaacguU...................    3    1    seq
.......................................................................auAuggaaggaaaaaacgug...................    1    1    seq
.......................................................................auguggaaggaaaaaacgGg...................    1    1    seq
.......................................................................augugCaaggaaaaaacgug...................    1    1    seq
.......................................................................auguggaaggaaaaaacguC...................    1    1    seq
.......................................................................auguggaaggaaaaaacguA...................    14   1    seq
.......................................................................auguggaaggaaaaGacgugc..................    1    1    seq
.......................................................................auguggaaggaaaaaacgugU..................    10   1    seq
.......................................................................auguggaaggaaaaaacgugc..................    33   0    seq
.......................................................................auguggaaggaaaaaacgugA..................    4    1    seq
.......................................................................auguggaaggaaaaaacgugG..................    1    1    seq
.......................................................................auguggaaggaaaaaacgugAu.................    4    1    seq
.......................................................................auguggaaggaaaaaacgugcu.................    36   0    seq
.......................................................................auguggaaggaaaaaacgugcA.................    6    1    seq
.......................................................................auguggaaggaaaaaacgugcC.................    11   1    seq
.......................................................................auguggaaggaaaaaacAugcu.................    1    1    seq
.......................................................................auguggaaggaaaaaacgugcG.................    4    1    seq
.......................................................................Guguggaaggaaaaaacgugcu.................    1    1    seq
.......................................................................auguggaaggaaaaaacgugcuU................    3    1    seq
.......................................................................auguggaaggaaaaaacgugcuA................    1    1    seq
........................................................................uguggaaggaaaaaacgug...................    3    0    seq
........................................................................uguggUaggaaaaaacgug...................    1    1    seq
........................................................................uguggaaggaaaaaacgugU..................    1    1    seq
........................................................................uguggaaggaaaaaacgugcu.................    1    0    seq
```

miRBase precursor : novel-nve-miR-49
Total read count : 4251
novel-nve-miR-49_guide read count : 4035
novel-nve-miR-49_star read count : 103
remaining reads : 113

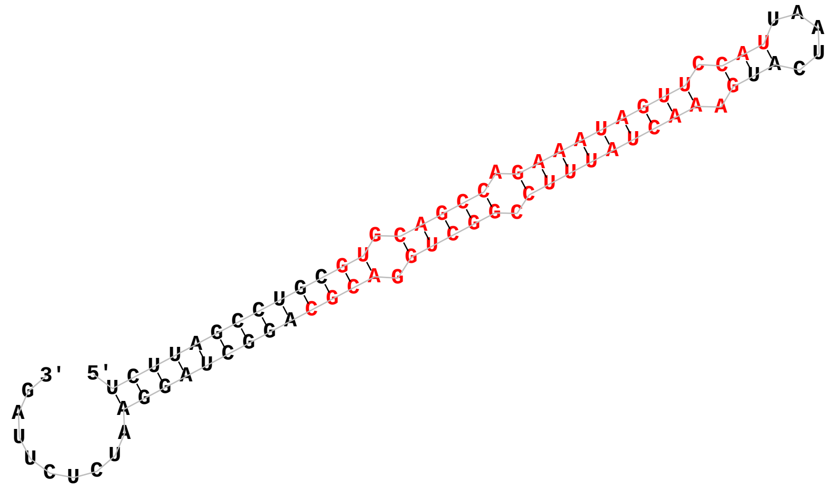

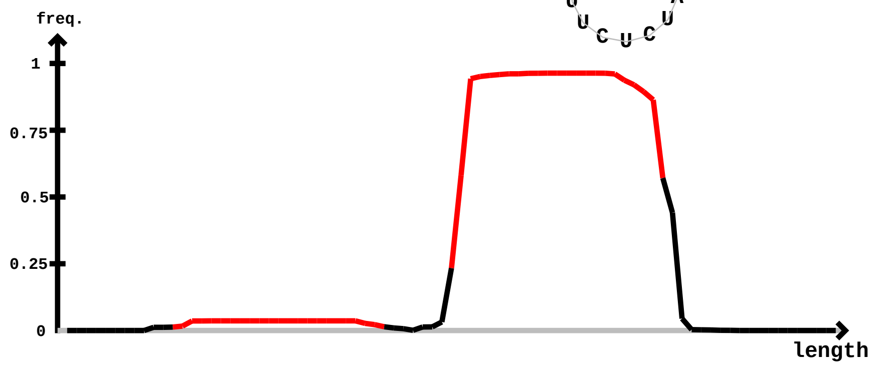


```
                 novel-nve-miR-49_star
                                      novel-nve-miR-49_guide
5'- ucuuagccugcgugcagccagaaauaguuccauuaaucaugaaacuauuuccggcuggacgcaggcuaggaaucucuuag  -3'    exp
    ((((((((((((.(((((.(((((((((.(((.....))).))))))))).))))).))))))))))))...........     reads      mm         sample
    ........ugcgugcagccGgaaauaguuc..................................................     40         1          seq
    ........ugcgugcagccGgaaauaguucc.................................................     11         1          seq
    ..........cgugcagccGgaaauaguucca................................................     3          1          seq
    ..........cgugcagccGgaaauaguuccau..............................................      1          1          seq
    ...........gugcagccGgaaauaguucc.................................................     5          1          seq
    ...........gugcagccGgaaauaguucca................................................     4          1          seq
    ...........gugcagccagaaauaguuccU................................................     1          1          seq
    ...........gugcagccGgaaauaguuccau..............................................      4          1          seq
    ...........gugcagccGgaaauaguuccauu.............................................      1          1          seq
    ............ugcagccGgaaauaguucc.................................................     4          1          seq
    ............ugcagccGgaaauaguucca................................................     26         1          seq
    ............ugcagccagGaauaguucca................................................     1          1          seq
    ............ugcagccGgaaauaguuccau...............................................     14         1          seq
    ............ugcagccGgaaauaguuccauu..............................................     11         1          seq
    ............ugcagccGgaaauaguuccauua.............................................     26         1          seq
    ..............cagccGgaaauaguuccauua.............................................     2          1          seq
    ................................uuaaucaugaaacuauuuccggcuggacg...................     1          0          seq
    ...................................aucaugaaacuauuuccggcu........................     2          0          seq
    ...................................aucaugaaacuauuuccggcuggacg...................     1          0          seq
    ....................................ucaugaaacuauuuccggU.........................     1          1          seq
    ....................................ucaugaaacuauuuccggcA........................     2          1          seq
    ....................................ucaugaaacuauuuccggcu........................     6          0          seq
    ....................................ucaugaaacuauuuccggcC........................     1          1          seq
    ....................................ucaugaaacuauuuccggcug.......................     9          0          seq
    ....................................ucaugaaaUuauuuccggcug.......................     1          1          seq
    ....................................ucaugaaacuauuuccggcuggaU....................     2          1          seq
    ....................................ucaugaaacuauuuccggcuggac....................     4          0          seq
    ....................................ucaugaaacuauuuccggcuggacg...................     17         0          seq
    ....................................ucaugaaacuauuuccggcuggacU...................     6          1          seq
    ....................................ucaugaaacuauuuccggcuggacA...................     3          1          seq
    .....................................caugaaacuauuuccggcuggacU...................     2          1          seq
    ......................................augaaacuauuuccggcu........................     1          0          seq
    ......................................augaaacuauuuccggcug.......................     9          0          seq
    ......................................augaaacuauuucUggcug.......................     1          1          seq
```

```
           novel-nve-miR-49_star
                                        novel-nve-miR-49_guide
ucuuagccugcgugcagccagaaauaguuccauuaaucaugaaacuauuuccggcuggacgcaggcuaggaaucucuuag

......................................augaaacuauuuccggcuA.......................    3      1      seq
......................................augaaacuGuuuccggcug.......................    1      1      seq
......................................augaaacuauuuccggcGg.......................    1      1      seq
......................................augaaacuauuuccggcugg......................    4      0      seq
......................................augaaacuGuuuccggcugg......................    1      1      seq
......................................augaaacuauuuccggcugU......................    1      1      seq
......................................augaaacuaCuuccggcugg......................    1      1      seq
......................................augaaacuauuuccggcugga.....................    4      0      seq
......................................augaaacuauuuccggcuggG.....................    1      1      seq
......................................augaaacuauuuccggcuggaU....................    2      1      seq
......................................augaaacuauuuccggcuggac....................    3      0      seq
......................................augaaacuauuuccggcuggacA...................    5      1      seq
......................................augaaacuauuuccggcuggacg...................    17     0      seq
......................................augaaacuauuuccggcuggacU...................    20     1      seq
......................................augaaacuauuuccggcuggacC...................    2      1      seq
......................................augaaacuauuAccggcuggacg...................    1      1      seq
.......................................ugaaacuauuuccggcuA.......................    4      1      seq
.......................................ugaaacuauuuccggcug.......................    63     0      seq
.......................................ugaaacuauuuAcggcug.......................    1      1      seq
.......................................ugaaacuauuuUcggcug.......................    4      1      seq
.......................................ugaaacuauuuccggcuU.......................    1      1      seq
.......................................ugaaacuCuuuccggcug.......................    1      1      seq
.......................................ugaaacuauuuccggcuC.......................    1      1      seq
.......................................ugaaacuauuuccAgcugg......................    1      1      seq
.......................................ugaaacuauuuccggcugA......................    7      1      seq
.......................................ugaaacuauuuUcggcugg......................    1      1      seq
.......................................ugaGacuauuuccggcugg......................    1      1      seq
.......................................ugaaGcuauuuccggcugg......................    1      1      seq
.......................................Agaaacuauuuccggcugg......................    4      1      seq
.......................................ugaaacuauuuccggcugU......................    3      1      seq
.......................................ugaaacuauuuccggcuAg......................    1      1      seq
.......................................ugaaacuauuuccggcugg......................    39     0      seq
.......................................ugaaacuauuuccggcugUa.....................    1      1      seq
.......................................ugaaacuauuuUcggcugga.....................    1      1      seq
.......................................ugaaCcuauuuccggcugga.....................    1      1      seq
.......................................ugaaacuauuCccggcugga.....................    1      1      seq
.......................................ugaaacuauuuccggcuggG.....................    4      1      seq
.......................................ugaaacuauuuccAgcugga.....................    1      1      seq
.......................................Agaaacuauuuccggcugga.....................    3      1      seq
.......................................ugaaacuauuuccggcugga.....................    66     0      seq
.......................................ugaaacuauuuccgUcuggac....................    1      1      seq
.......................................ugaaacuauuuUcggcuggac....................    1      1      seq
.......................................ugaaUcuauuuccggcuggac....................    1      1      seq
.......................................ugaaacuauuuccggcuggaA....................    1      1      seq
.......................................Agaaacuauuuccggcuggac....................    1      1      seq
.......................................ugaaacuauuuccggcAggac....................    1      1      seq
.......................................ugaaacuauuuccggcuggac....................    31     0      seq
.......................................ugaaacuauuuccggcuggaU....................    28     1      seq
.......................................ugaaacuauuuccggcuggacA...................    49     1      seq
.......................................ugaaacuauuCccggcuggacg...................    1      1      seq
.......................................ugaaaUuauuuccggcuggacg...................    3      1      seq
.......................................ugaaacuauuucAggcuggacg...................    1      1      seq
.......................................Cgaaacuauuuccggcuggacg...................    2      1      seq
.......................................ugaaacuauuuccggcuggacg...................    274    0      seq
.......................................ugaaacuauuuccggcuAgacg...................    1      1      seq
.......................................ugaaacuauuuccggcuggaUg...................    1      1      seq
.......................................uAaaacuauuuccggcuggacg...................    1      1      seq
.......................................ugaaacCauuuccggcuggacg...................    1      1      seq
.......................................ugaaGcuauuuccggcuggacg...................    2      1      seq
.......................................ugaaacuauuuccggcuggacU...................    107    1      seq
.......................................ugaaacuauuuUcggcuggacg...................    2      1      seq
.......................................ugaaacuauuuAcggcuggacg...................    1      1      seq
.......................................ugaaacuauuuccggcuggacC...................    15     1      seq
.......................................ugCaacuauuuccggcuggacg...................    1      1      seq
.......................................ugaaacuauCuccggcuggacg...................    2      1      seq
.......................................uCaaacuauuuccggcuggacg...................    2      1      seq
.......................................ugaaacuauuuccggcuggaAg...................    1      1      seq
.......................................ugaaacuauAuccggcuggacg...................    1      1      seq
.......................................Agaaacuauuuccggcuggacg...................    3      1      seq
```

```
                     novel-nve-miR-49_star
                                                        novel-nve-miR-49_guide
ucuuagccugcgugcagccagaaauaguuccauuaaucaugaaacuauuuccggcuggacgcaggcuaggaaucucuuag

.......................................ugaaacuauuuccggcuggacgU..................    41    1    seq
.......................................ugaaacuauuuccggcuggacgA..................    13    1    seq
.......................................ugaaacuauuuccggcuggacgc..................    5     0    seq
.......................................Agaaacuauuuccggcuggacgc..................    1     1    seq
.......................................ugaaacuauuuccggcuggacgUa.................    2     1    seq
.......................................ugaaacuauuuccggcuggacgcG.................    2     1    seq
.......................................ugaaacuauuuccggcuggacgcC.................    7     1    seq
.......................................ugaaacuauuuccggcuggacgcU.................    34    1    seq
.......................................ugaaacuauuuccggcuggacgca.................    4     0    seq
.......................................ugaaacuauuuccggcuggacgAa.................    2     1    seq
........................................gaaacuauuuccggcugg......................    6     0    seq
........................................gaaacuauuuccggcugA......................    2     1    seq
........................................gaaacuauuuccggcugga.....................    19    0    seq
........................................gaaacuauuucAggcugga.....................    1     1    seq
........................................gGaacuauuuccggcugga.....................    3     1    seq
........................................gaaacuauuuccggcuggac....................    20    0    seq
........................................gaaacuauuuccggcuggaU....................    6     1    seq
........................................Caaacuauuuccggcuggacg...................    1     1    seq
........................................gaaacuauuuccggcuggGcg...................    1     1    seq
........................................gaaacuauuuccggcuggacU...................    177   1    seq
........................................gaaacAauuuccggcuggacg...................    1     1    seq
........................................gaaacuauuucUggcuggacg...................    1     1    seq
........................................gaaGcuauuuccggcuggacg...................    2     1    seq
........................................gaaacCauuuccggcuggacg...................    1     1    seq
........................................gaaacuauuuccggcuggacA...................    66    1    seq
........................................gaaacuauuuccggcuggacg...................    165   0    seq
........................................gaaacuauuuUcggcuggacg...................    1     1    seq
........................................gaaacuauuAccggcuggacg...................    1     1    seq
........................................gGaacuauuuccggcuggacg...................    2     1    seq
........................................gaaacuauuuccggcAggacg...................    1     1    seq
........................................gaaacuaCuuccggcuggacg...................    1     1    seq
........................................Uaaacuauuuccggcuggacg...................    1     1    seq
........................................gaaacuGuuuccggcuggacg...................    2     1    seq
........................................gaaacuauuuccggcuggacC...................    47    1    seq
........................................gaaacuauAuccggcuggacg...................    1     1    seq
........................................gaaacuauuuccgAcuggacg...................    1     1    seq
........................................gaaacuauuuccggcuggacgG..................    4     1    seq
........................................gaaacuauuuccggcuggacUc..................    1     1    seq
........................................gaaacuauuuccggcCggacgc..................    1     1    seq
........................................gaaacuauuuccggcuggacgA..................    14    1    seq
........................................gaaacuauuuccggcuggacgU..................    51    1    seq
........................................gaaacuauuuccggcuggacgc..................    89    0    seq
........................................gaaacGauuuccggcuggacgc..................    1     1    seq
........................................gaaacuauuuccggcuggCcgc..................    1     1    seq
........................................gaaacCauuuccggcuggacgc..................    1     1    seq
........................................gaaacuauuuccggcuggacgcC.................    93    1    seq
........................................gaaacuauuuccggcuggacgAa.................    1     1    seq
........................................gaGacuauuuccggcuggacgca.................    2     1    seq
........................................gaaacuauuuccggcuggacgca.................    79    0    seq
........................................gaaacuauuuccggcuggacgcU.................    562   1    seq
........................................gaaacuauuuccAgcuggacgca.................    1     1    seq
........................................gaaUcuauuuccggcuggacgca.................    1     1    seq
........................................gGaacuauuuccggcuggacgca.................    1     1    seq
........................................gaaacuauuuccggcuggGcgca.................    2     1    seq
........................................gaaacuauuucUggcuggacgca.................    1     1    seq
........................................gaaacuauuuccggcCggacgca.................    1     1    seq
........................................gaaacCauuuccggcuggacgca.................    1     1    seq
........................................gaaacuauuuccggcuggacgcG.................    24    1    seq
........................................gaaacuauuuccggcuggacgcaC................    4     1    seq
........................................gaaacuauuuccggcuggacgcag................    1     0    seq
........................................gaaacuauuuccggcuggacgcaA................    1     1    seq
........................................gaaacuauuuccggcuggacgcaU................    7     1    seq
........................................gaaacuauuuccggcuggacgcUg................    2     1    seq
.........................................aaacuauuuccggcugga.....................    3     0    seq
.........................................aaacuauuuccggcuggaA....................    1     1    seq
.........................................aaacuauuCccggcuggac....................    2     1    seq
.........................................aaacuauuuccggcuggac....................    20    0    seq
.........................................aaacuauuuccggcuggaU....................    5     1    seq
.........................................aaacuauuuccggcuggGcg...................    2     1    seq
```

```
           novel-nve-miR-49_star
                                             novel-nve-miR-49_guide
ucuuagccugcgugcagccagaaauaguuccauuaaucaugaaacuauuuccggcuggacgcaggcuaggaaucucuuag

.........................................aaacuauuuccggcuggacg...................    154    0    seq
.........................................aaacuauuuccggcCggacg...................    1      1    seq
.........................................aaacuauuuccggcuggacA...................    34     1    seq
.........................................aaacAauuuccggcuggacg...................    1      1    seq
.........................................aaacuauuuccUgcuggacg...................    2      1    seq
.........................................aaacuauuucAggcuggacg...................    1      1    seq
.........................................aaacuauuuccggcuggacU...................    9      1    seq
.........................................aaacuauuuAcggcuggacg...................    1      1    seq
.........................................aaaAuauuuccggcuggacg...................    1      1    seq
.........................................aaacuauuucUggcuggacg...................    1      1    seq
.........................................aaaUuauuuccggcuggacg...................    2      1    seq
.........................................aaacuauuuccggcuggacC...................    2      1    seq
.........................................aaGcuauuuccggcuggacg...................    1      1    seq
.........................................aaacuauuuccgAcuggacg...................    1      1    seq
.........................................aaacuauuuccggcuggacUc..................    1      1    seq
.........................................aaacuauuuccggcCggacgc..................    2      1    seq
.........................................aaacuauuuccAgcuggacgc..................    1      1    seq
.........................................aaaUuauuuccggcuggacgc..................    2      1    seq
.........................................aaacuauuuccggcuggacgA..................    16     1    seq
.........................................aaacuauuuccggAuggacgc..................    2      1    seq
.........................................Gaacuauuuccggcuggacgc..................    1      1    seq
.........................................aaacuauuuccggcuggacgG..................    1      1    seq
.........................................aaacuauuuccggcuggGcgc..................    1      1    seq
.........................................aaacCauuuccggcuggacgc..................    1      1    seq
.........................................Caacuauuuccggcuggacgc..................    1      1    seq
.........................................aaacuGuuuccggcuggacgc..................    1      1    seq
.........................................aaacuauuuccggcuggacAc..................    1      1    seq
.........................................aaacuaCuuccggcuggacgc..................    1      1    seq
.........................................aaacuauuuccggcuAgacgc..................    1      1    seq
.........................................aaacuauuuccggcuggacgU..................    75     1    seq
.........................................aaacuauuuAcggcuggacgc..................    1      1    seq
.........................................aaacAauuuccggcuggacgc..................    1      1    seq
.........................................aaacuauuuccggcuggacgc..................    193    0    seq
.........................................aaacuauuuccggcuggacgcC.................    108    1    seq
.........................................aaacuauuuccggcuggacgUa.................    1      1    seq
.........................................aaacuauuCccggcuggacgca.................    1      1    seq
.........................................aaacuauuuccggcuggacgcG.................    19     1    seq
.........................................aaacuauuuccggcuggacgca.................    106    0    seq
.........................................aaacuauuuccggUuggacgca.................    2      1    seq
.........................................aaaUuauuuccggcuggacgca.................    1      1    seq
.........................................aaacuauuucUggcuggacgca.................    1      1    seq
.........................................aGacuauuuccggcuggacgca.................    1      1    seq
.........................................aaacuauuuccggcuggacgcU.................    593    1    seq
.........................................aaacuauuuccUgcuggacgca.................    1      1    seq
.........................................aaacuauuAccggcuggacgca.................    1      1    seq
.........................................aaacuauuuccggcuggacAca.................    1      1    seq
.........................................aaacuauuuccggcuggaAgca.................    1      1    seq
.........................................aaacuaCuuccggcuggacgca.................    1      1    seq
.........................................aaacuauuuccggcuggacgcUg................    4      1    seq
.........................................aaacuauuuccggcuggacgcaA................    8      1    seq
.........................................aaacuauuuccggcuggacgcaC................    21     1    seq
.........................................aaacuauuuccggcuggacgcaU................    117    1    seq
.........................................aaacuauuuccggcuggacgcaUg...............    2      1    seq
..........................................aacuauuuccggcuggacg...................    3      0    seq
..........................................aacuauuuccggcuggacA...................    1      1    seq
..........................................aacuauuuccggcugCacg...................    5      1    seq
..........................................aacuauuuccggcugCacgc..................    11     1    seq
..........................................aacuauuuccggcuggacgc..................    2      0    seq
..........................................aacuauuuccggcuggacgU..................    2      1    seq
..........................................aacuauuuccggcuggacgA..................    1      1    seq
..........................................aacuauuuccggcuggacgcG.................    1      1    seq
..........................................aacuauuuccggcugCacgca.................    1      1    seq
..........................................aacuauuuccggcuggacgca.................    2      0    seq
..........................................aacuauuuccggcuggacgcU.................    6      1    seq
..........................................aacuauuuccggcuggacgcaU................    1      1    seq
...........................................acuauuuccggcugCacg...................    3      1    seq
...........................................acuauuuccggcuggacC...................    1      1    seq
...........................................acuauuuccggcugCacgc..................    4      1    seq
...........................................acuauuuccggcugUacgc..................    1      1    seq
```

```
           novel-nve-miR-49_star
                            novel-nve-miR-49_guide
ucuuagccugcgugcagccagaaauaguuccauuaaucaugaaacuauuuccggcuggacgcaggcuaggaaucucuuag

...........................................acuauuuccggcuggacgc..................    1     0     seq
...........................................acuauuuccggcugCacgca.................    7     1     seq
............................................cuauuuccggcuggacgc..................    2     0     seq
............................................cuauuuccggcuggacgcU.................    2     1     seq
............................................cuauuuccggcuggacgcC.................    1     1     seq
............................................cuauuuccggcuggacgcG.................    1     1     seq
............................................cuauuuccggcuggacgcaC................    1     1     seq
............................................cuauuuccggcugCacgcag................    1     1     seq
............................................cuauuuccggcuggacgcaU................    4     1     seq
............................................cuauuuccggcugCacgcagg...............    1     1     seq
.............................................uauuuccggcugCacgca.................    4     1     seq
.............................................uauuuccggcuggacgcU.................    3     1     seq
.............................................uauuuccggcuggacgcC.................    1     1     seq
.............................................uauuuccggcugCacgcag................    4     1     seq
...............................................uuuccggcugCacgcaggc..............    2     1     seq
...............................................uAuccggcuggacgcaggcu.............    1     1     seq
...............................................uuuccggcugCacgcaggcu.............    4     1     seq
...............................................uuuccggcugCacgcaggcua............    1     1     seq
................................................uuccggcuggacgcaggcuaU...........    1     1     seq
.................................................uccggcuggacgcaggcuaA...........    2     1     seq
```

| | | |
|---|---|---|
| Provisional ID | | novel-nve-miR-51 |
| Score total | : | 45.5 |
| Score for star read(s) | : | 3.9 |
| Score for read counts | : | 37.3 |
| Score for mfe | : | -0.3 |
| Score for randfold | : | 1.6 |
| Score for cons. seed | : | 3 |
| Total read count | : | 85 |
| Mature read count | : | 82 |
| Loop read count | : | 0 |
| Star read count | : | 3 |

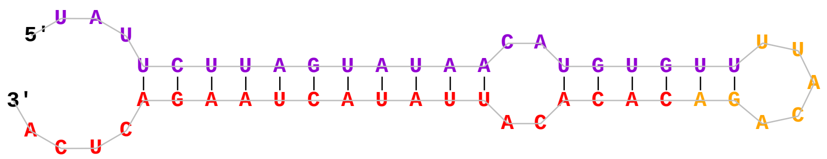

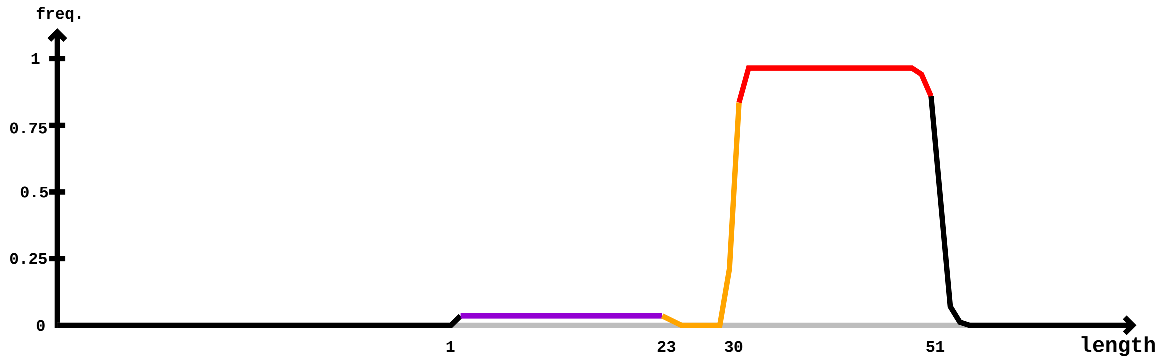


Star Mature

```
5'- gugugcuuuuaaaguccacauuuauaauuaagucuuacaauauucuuaguauaacauguguuuuacagacacacauuauacuaagacucaaaauggcucaauuugaaggu  -3'   obs
    gugugcuuuuaaaguccacauuuauaauuaagucuuacaauauucuuaguauaacauguguuuuacagacacacauuauacuaagacucaaaauggcucaauuugaaggu         exp
    (((..((.....))..))).((((.((((.((((.........(((((((((((..((((((.....))))))..)))))))))))........)))).))))))))...   reads   mm   sample
    ........................................uauucuuaguauaacauguguu................................................   3       0    seq
    ....................................................................acacacauuauacuaagacG......................   1       1    seq
    ....................................................................acacacauuauacuaagacuA.....................   1       1    seq
    ....................................................................acacacauuauacuaagacuc.....................   3       0    seq
    ....................................................................acacacauuauacuaagacucC....................   1       1    seq
    ....................................................................acacacauuauacuaagacucU....................   3       1    seq
    ....................................................................acacacauuauacuaagacucG....................   4       1    seq
    ....................................................................acacacauuauacuaagacuca....................   1       0    seq
    ....................................................................acacacauuauacuaagacucaG...................   1       1    seq
    ....................................................................acacacauuauacuaagacucaa...................   3       0    seq
    .....................................................................cacacauuauacuaagacu......................   1       0    seq
    .....................................................................cacacauuauacuaagacuc.....................   2       0    seq
    .....................................................................cacacauuauacuaagacuU.....................   1       1    seq
    .....................................................................Uacacauuauacuaagacuca....................   1       1    seq
    .....................................................................cacacauuauacuaagacucG....................   15      1    seq
    .....................................................................cacacauuauacuaagacucC....................   2       1    seq
    .....................................................................cacacauuauacuaagacucU....................   7       1    seq
    .....................................................................cacacauuauacuaagacuca....................   12      0    seq
    .....................................................................cacacauuauacuaagacucaU...................   4       1    seq
    .....................................................................cacacauuauacuaagacucaG...................   1       1    seq
    .....................................................................cacacauuauacuaagacucaa...................   5       0    seq
    .....................................................................caUacauuauacuaagacucaa...................   1       1    seq
    .....................................................................cacacauuauacuaagacucUa...................   1       1    seq
    ......................................................................acacauuauacuaagacucG....................   1       1    seq
    ......................................................................acacauuauacuaagacuca....................   2       0    seq
    ......................................................................acacauuauacuaagacucaa...................   1       0    seq
    ......................................................................acacauuauacuaagacucaG...................   1       1    seq
    ......................................................................acacauuauacuaagacucaaG..................   1       1    seq
    ......................................................................acacauuauacuaagacucaaU..................   2       1    seq
    ......................................................................acacauuauacuaagacucaaa..................   2       0    seq
    ......................................................................acacauuauacuaagacucaaaU.................   1       1    seq
```

miRBase precursor : novel-nve-miR-53
Total read count : 230
novel-nve-miR-53_guide read count : 218
novel-nve-miR-53_star read count : 12
remaining reads : 0

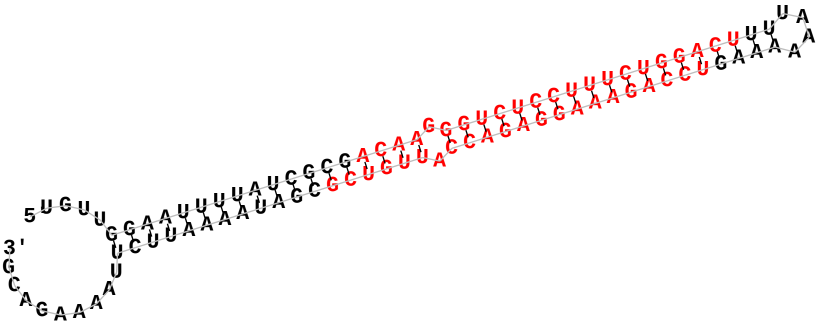

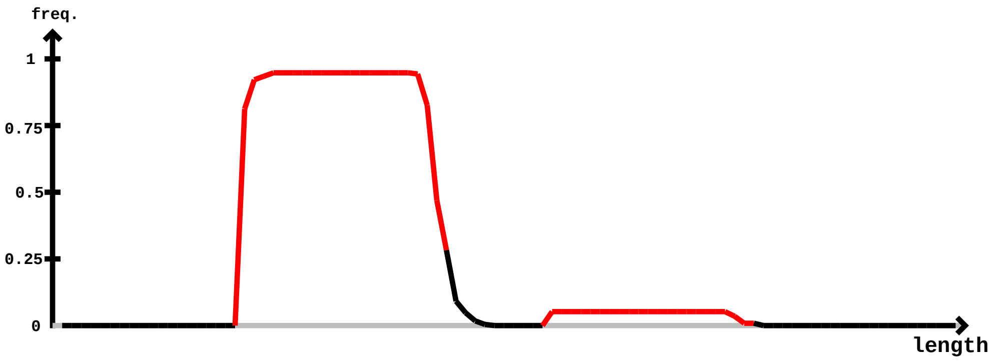


novel-nve-miR-53_star

novel-nve-miR-53_guide

```
5'- uguuggaauuuuaucgcgacaagggucuccuuucuggacuuuuaaaaaaguccagaaaggagaccauugucgcgauaaaauucuuaaaagacg   -3'    exp
    .(((((((((((((((((((((.(((((((((((((((((((....))))))))))))))))))).)))))))))))))))))).....))).    reads   mm   sample
    ..................acaagggucuccuuucug.........................................................    1       0    seq
    ..................acaagggucuccuuucugA........................................................    2       1    seq
    ..................acaagggucuccuuucugU........................................................    2       1    seq
    ..................acaagggucuccuuucugg........................................................    23      0    seq
    ..................acaagggucuccuuucCgga.......................................................    1       1    seq
    ..................acaagggucuccuuucuggU.......................................................    1       1    seq
    ..................acaagggucCccuuucugga.......................................................    1       1    seq
    ..................acaagggucuccuuucugga.......................................................    59      0    seq
    ..................Ccaagggucuccuuucugga.......................................................    1       1    seq
    ..................acaagggucuUcuuucugga.......................................................    1       1    seq
    ..................acaagggucuccCuucugga.......................................................    2       1    seq
    ..................acaagggCcuccuuucugga.......................................................    1       1    seq
    ..................acaaggguUuccuuucugga.......................................................    1       1    seq
    ..................acaagggucuccuuucuggG.......................................................    2       1    seq
    ..................acaagggucuccuuucuggaA......................................................    2       1    seq
    ..................aUaagggucuccuuucuggac......................................................    1       1    seq
    ..................acaagggucuccuuucuggaU......................................................    12      1    seq
    ..................acaagggucuccuuucuggac......................................................    23      0    seq
    ..................acaagggucAccuuucuggac......................................................    1       1    seq
    ..................acaagggucuccuuucuggacC.....................................................    4       1    seq
    ..................acaagggucuccuuucuggacA.....................................................    2       1    seq
    ..................acaagggucuccuuucuggacG.....................................................    1       1    seq
    ..................acaaggguUuccuuucuggacu.....................................................    1       1    seq
    ..................acaagggucuccuGucuggacu.....................................................    1       1    seq
    ..................acaagggucuccuuucuggaAu.....................................................    1       1    seq
    ..................acaagggucuccuuucuggacu.....................................................    26      0    seq
    ..................acaagggucuccuuucuggacuA....................................................    4       1    seq
    ..................acaagggucuccuuucuggaUuu....................................................    2       1    seq
    ..................acaagggucuccuuucuggacuAu...................................................    3       1    seq
    ..................acaagggucuccuuucuggaUuuu...................................................    1       1    seq
    ..................acaagggucuccuuucuggacuuu...................................................    1       0    seq
    ..................acaagggucuccuuucuggacuuAu..................................................    1       1    seq
    ..................acaagggucuccuuucuggacuuuu..................................................    2       0    seq
    ...................caagggucuccuuucuggG.......................................................    1       1    seq
```

```
                                            novel-nve-miR-53_star
                  novel-nve-miR-53_guide
uguuggaauuuuaucgcgacaagggucuccuuucuggacuuuuaaaaaaguccagaaaggagaccauugucgcgauaaaauucuuaaaagacg

...................caagggucuccuuucugga.......................................................	9	0	seq
...................caagggucuccuuucuggaU......................................................	1	1	seq
...................caagggucuccuuucuggac......................................................	2	0	seq
...................caagggucuccuuucuggacu.....................................................	6	0	seq
...................caagggucuccuuucuCgacu.....................................................	1	1	seq
...................caagggucuccuuucuggaUu.....................................................	1	1	seq
...................caagggucuccuuucuggacuu....................................................	3	0	seq
...................caagggucuccuuucuggacuuu...................................................	1	0	seq
....................aagggucuccuuucugga.......................................................	2	0	seq
....................aagggucuccuuucuggacuuC...................................................	1	1	seq
.....................agggucuccuuucuggaU......................................................	1	1	seq
.....................agggucuccuuucuggacuu....................................................	1	0	seq
.....................agggucuccuuucuggacuuuuU.................................................	1	1	seq
..................................................uccagaaaggagaccauug........................	4	0	seq
..................................................uccagaaaggagaccauugG.......................	1	1	seq
..................................................uccagaaaggagGccauugu.......................	1	1	seq
..................................................uccagaaaggagaccauugu.......................	3	0	seq
..................................................uccagaaaggagaccauugC.......................	1	1	seq
..................................................uccagaaaggagaccauugucA.....................	1	1	seq
..................................................uccagaaaggagaccauugucg.....................	1	0	seq
```

miRBase precursor : novel-nve-miR-54
Total read count : 2306
novel-nve-miR-54_guide read count : 2295
novel-nve-miR-54_star read count : 8
remaining reads : 3

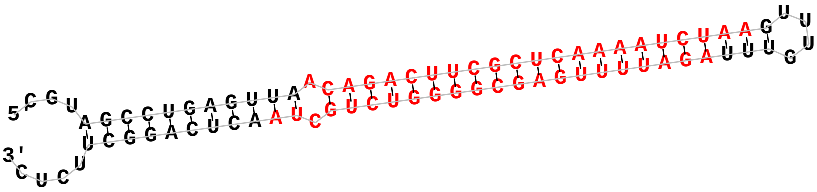


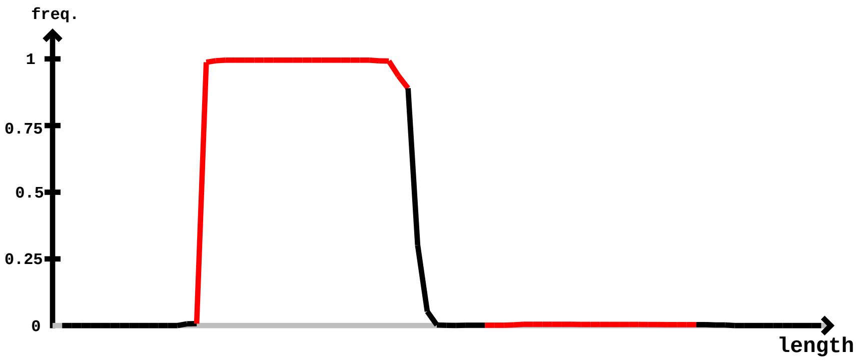


```
                                             novel-nve-miR-54_star
                   novel-nve-miR-54_guide
5'- cguagccugaguuaacagacuucgcucaaaaucuaaguuuguuuagauuuugagcggggucugcuaacucaggcuucuc   -3'    exp
    .(.((((((((((.((((((((((((((((((((((....)))))))))))))))))))))).)))))))))).)..   reads     mm        sample
    ............uaacagacuuUgcucaaaau...............................................   2         1         seq
    ............uaacagacuucgcucaaaauc..............................................   1         0         seq
    ............uaacagacuuUgcucaaaaucu.............................................   2         1         seq
    ............uaacGgacuucgcucaaaaucu.............................................   1         1         seq
    ............uaacagacuucgcucaaaaucu.............................................   7         0         seq
    ............uaacagacuucgcucaaaaucA.............................................   1         1         seq
    ............uaacagacuucgcucaaaaucC.............................................   1         1         seq
    .............aacagacuucgcucaaaaucC.............................................   1         1         seq
    .............aacagacuucgcucaaaaucuaa...........................................   1         0         seq
    ..............acagacuucgcucaaaau...............................................   3         0         seq
    ..............acagacuucgcucaaaaC...............................................   1         1         seq
    ..............acagacuucgcucaaaauc..............................................   1         0         seq
    ..............acagacuucgcucaaaaucC.............................................   24        1         seq
    ..............acagacuucgcucaaaaucA.............................................   1         1         seq
    ..............acagacuucgcucaaaauUu.............................................   1         1         seq
    ..............acagacuGcgcucaaaaucu.............................................   1         1         seq
    ..............acagacuCcgcucaaaaucu.............................................   1         1         seq
    ..............acagaAuucgcucaaaaucu.............................................   1         1         seq
    ..............acGgacuucgcucaaaaucu.............................................   1         1         seq
    ..............acagacuucgcucaaaaucu.............................................   81        0         seq
    ..............acagacuucgcucaaaaucG.............................................   4         1         seq
    ..............acagacuucgcucaaGaucua............................................   1         1         seq
    ..............Gcagacuucgcucaaaaucua............................................   1         1         seq
    ..............acagacuCcgcucaaaaucua............................................   2         1         seq
    ..............acagacuucgcucaaaaucuG............................................   1         1         seq
    ..............acagacuuUgcucaaaaucua............................................   1         1         seq
    ..............acagacuuGgcucaaaaucua............................................   1         1         seq
    ..............acagacuucgcucaaaaucua............................................   100       0         seq
    ..............acagacCucgcucaaaaucua............................................   1         1         seq
    ..............acagacuucgcucaaGaucuaa...........................................   2         1         seq
    ..............acagacuGcgcucaaaaucuaa...........................................   1         1         seq
    ..............acaCacuucgcucaaaaucuaa...........................................   1         1         seq
    ..............Gcagacuucgcucaaaaucuaa...........................................   7         1         seq
    ..............acagacuucgAucaaaaucuaa...........................................   5         1         seq
```

novel-nve-miR-54_star

novel-nve-miR-54_guide

cguagccugaguuaacagacuucgcucaaaaucuaaguuuguuuagauuuugagcggggucugcuaacucaggcuucuc

| | | | |
|---|---|---|---|
| ..............acagaAuucgcucaaaaucuaa........................................... | 4 | 1 | seq |
| ..............acagUcuucgcucaaaaucuaa........................................... | 1 | 1 | seq |
| ..............aAagacuucgcucaaaaucuaa........................................... | 2 | 1 | seq |
| ..............acagGcuucgcucaaaaucuaa........................................... | 3 | 1 | seq |
| ..............acagacuucgcuAaaaaucuaa........................................... | 4 | 1 | seq |
| ..............acagacuucgcucaaaaucuaU........................................... | 51 | 1 | seq |
| ..............acagacuucgcucaaUaucuaa........................................... | 1 | 1 | seq |
| ..............acagacuucgcucaaaaucuaG........................................... | 20 | 1 | seq |
| ..............acagacuucgUucaaaaucuaa........................................... | 1 | 1 | seq |
| ..............Ccagacuucgcucaaaaucuaa........................................... | 1 | 1 | seq |
| ..............acagacuucgcucGaaaucuaa........................................... | 3 | 1 | seq |
| ..............acagacuucgcucaGaaucuaa........................................... | 3 | 1 | seq |
| ..............acagacuucgcucaaaGucuaa........................................... | 4 | 1 | seq |
| ..............acagacuuUgcucaaaaucuaa........................................... | 4 | 1 | seq |
| ..............acagacuucgcucaaaaucuaC........................................... | 41 | 1 | seq |
| ..............acagacuucCcucaaaaucuaa........................................... | 4 | 1 | seq |
| ..............aUagacuucgcucaaaaucuaa........................................... | 3 | 1 | seq |
| ..............acagacGucgcucaaaaucuaa........................................... | 1 | 1 | seq |
| ..............Ucagacuucgcucaaaaucuaa........................................... | 1 | 1 | seq |
| ..............acaUacuucgcucaaaaucuaa........................................... | 2 | 1 | seq |
| ..............acagacuucgcucaaaauUuaa........................................... | 8 | 1 | seq |
| ..............acagacuucUcucaaaaucuaa........................................... | 2 | 1 | seq |
| ..............acGgacuucgcucaaaaucuaa........................................... | 3 | 1 | seq |
| ..............acagacuCcgcucaaaaucuaa........................................... | 7 | 1 | seq |
| ..............acagacuucAcucaaaaucuaa........................................... | 2 | 1 | seq |
| ..............acagacuucgcCcaaaaucuaa........................................... | 5 | 1 | seq |
| ..............acagacAucgcucaaaaucuaa........................................... | 4 | 1 | seq |
| ..............acagCcuucgcucaaaaucuaa........................................... | 1 | 1 | seq |
| ..............acagacuucgcucaaaaCcuaa........................................... | 2 | 1 | seq |
| ..............acUgacuucgcucaaaaucuaa........................................... | 1 | 1 | seq |
| ..............acagacuucgcucaaaaucuGa........................................... | 4 | 1 | seq |
| ..............acagacuucgcucaaaaucuaa........................................... | 1140 | 0 | seq |
| ..............acagacCucgcucaaaaucuaa........................................... | 4 | 1 | seq |
| ..............acagacuucgcucaaaaucuaag.......................................... | 1 | 0 | seq |
| ..............acagacuucgcucaaaaucuaCg.......................................... | 2 | 1 | seq |
| ..............acagacuucgcucaaaaucuaaA.......................................... | 46 | 1 | seq |
| ..............acagacuucgcucaaaaucuaaC.......................................... | 177 | 1 | seq |
| ..............acagacuucgcucaaaaucuaaU.......................................... | 338 | 1 | seq |
| ..............acagacuucgcucaaaaucuaaUu......................................... | 37 | 1 | seq |
| ..............acagacuucgcucaaaaucuaagu......................................... | 1 | 0 | seq |
| ..............acagacuucgcucaaaaucuaaAu......................................... | 6 | 1 | seq |
| ..............acagacuucgcucaaaaucuaaCu......................................... | 67 | 1 | seq |
| ..............acagacuucgcucaaaaucuaaUuu........................................ | 3 | 1 | seq |
| ..............acagacuucgcucaaaaucuaaUuuu....................................... | 1 | 1 | seq |
| ...............cagacuucgcucaaaaucuaC........................................... | 1 | 1 | seq |
| ...............cagacuucgcucaaaaucuaa........................................... | 1 | 0 | seq |
| ...............cagacuucgcucaaaaucuaaC.......................................... | 1 | 1 | seq |
| ...............cagacuucgcucaaaaucuaaU.......................................... | 6 | 1 | seq |
| ...............cagacuucgcucaaaaucuaaCu......................................... | 4 | 1 | seq |
| ................agacuucgcucaaaaucuaaC.......................................... | 3 | 1 | seq |
| ................agacuucgcucaaaaucuaaUu......................................... | 1 | 1 | seq |
| ................agacuucgcucaaaaucuaaCu......................................... | 1 | 1 | seq |
| ...................................aguuuguuuagauuGuga.......................... | 1 | 1 | seq |
| .........................................uuuagauuuugagcggggu................... | 1 | 0 | seq |
| .........................................uuuagauuuugagcgggguUu................. | 1 | 1 | seq |
| ..............................................auuuugagcggggucugUuaa............ | 3 | 1 | seq |
| ...............................................uuuugagcggggucugcuaacu.......... | 1 | 0 | seq |
| ...............................................uuuugagcggggucugcuaacG.......... | 1 | 1 | seq |
| ...............................................uuuugagcggggucugUuaacu.......... | 3 | 1 | seq |

miRBase precursor : novel-nve-miR-55
Total read count : 1714
novel-nve-miR-55_guide read count : 1502
novel-nve-miR-55_star read count : 211
remaining reads : 1

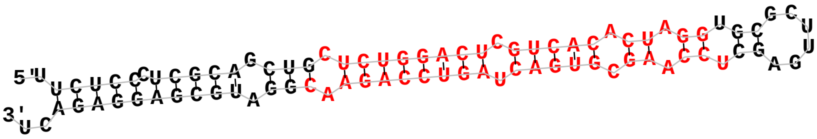

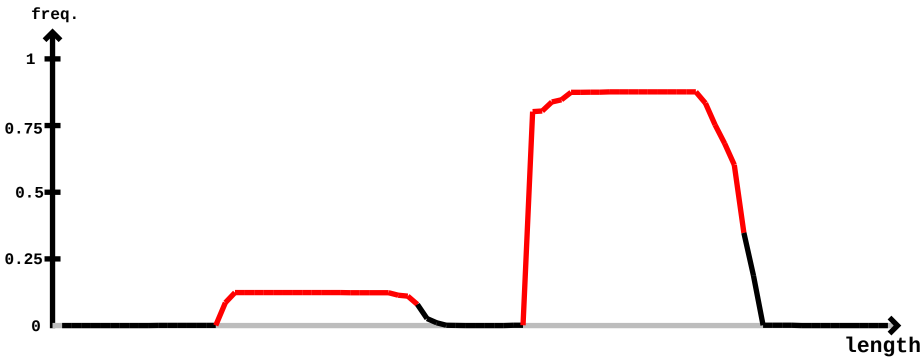


```
                                                     novel-nve-miR-55_guide
                    novel-nve-miR-55_star
5'- uucucccucgcagcugcucuggacucgucacacuaggugcgcuugagcuccaagcgugacuaguccagaacggaugcgaggagacu  -3'   exp
    .(((((.(((((.(((.((((((((.(((((.((.((.((......)).)).)).))))).)))))))).))).))))))))))..    reads     mm     sample
    .........gcagcCgcucuggacucguc.........................................................   1         1      seq
    ................cucuggacucgucacacC....................................................   3         1      seq
    ................cucuggacucgucacacA....................................................   1         1      seq
    ................cucuggacucgucacacu....................................................   10        0      seq
    ................cucuggacucgucacacG....................................................   2         1      seq
    ................cucuggacucgucacacuU...................................................   1         1      seq
    ................cucuggacucgucacacua...................................................   2         0      seq
    ................cucugAacucgucacacuag..................................................   1         1      seq
    ................cucuggacucgucacacuag..................................................   37        0      seq
    ................cucuggacucgucGcacuag..................................................   1         1      seq
    ................cucCggacucgucacacuag..................................................   1         1      seq
    ................cucuggacucgucacacuaU..................................................   1         1      seq
    ................Uucuggacucgucacacuag..................................................   2         1      seq
    ................cucuggacucgCcacacuag..................................................   1         1      seq
    ................cucuggacucgucacacuaA..................................................   4         1      seq
    ................cucuCgacucgucacacuag..................................................   1         1      seq
    ................cucuggacucgucaUacuagg.................................................   1         1      seq
    ................cuAuggacucgucacacuagg.................................................   1         1      seq
    ................cucuggacucgucacacuagA.................................................   1         1      seq
    ................cAcuggacucgucacacuagg.................................................   1         1      seq
    ................cucuggacucgucacacuagg.................................................   59        0      seq
    ................cucuggacucgucacacuagU.................................................   1         1      seq
    ................cucuggacuAgucacacuagg.................................................   1         1      seq
    ................cucugAacucgucacacuaggu................................................   1         1      seq
    ................cucuggacucgucacacuaggu................................................   4         0      seq
    ................cucuggacucgucacacuaggC................................................   7         1      seq
    ................cucuggacucgucacacuaggCg...............................................   1         1      seq
    .................uUuggacucgucacacua...................................................   2         1      seq
    .................ucuggacucgucacacuG...................................................   1         1      seq
    .................ucuggacucgucacacuag..................................................   3         0      seq
    .................ucuggacucgucacacCag..................................................   1         1      seq
    .................ucugUacucgucacacuagg.................................................   1         1      seq
    .................ucuggacucgucacacuagg.................................................   18        0      seq
    .................ucuggacucgucacacuagA.................................................   3         1      seq
```

novel-nve-miR-55_guide

novel-nve-miR-55_star

uucucccucgcagcugcucuggacucgucacacuaggugcgcuugagcuccaagcgugacuaguccagaacggaugcgaggagacu

| | | | |
|---|---|---|---|
| .................Ccuggacucgucacacuagg................................................. | 1 | 1 | seq |
| .................ucuggacucgucacacuagU................................................. | 2 | 1 | seq |
| .................ucuCgacucgucacacuagg................................................. | 1 | 1 | seq |
| .................ucuggacucgucacacuaggC................................................ | 11 | 1 | seq |
| .................ucuggacucgucacacuaggu................................................ | 4 | 0 | seq |
| .................ucuggacucgucacacuaggCg............................................... | 12 | 1 | seq |
| .................ucuggacucgucacacuaggug............................................... | 3 | 0 | seq |
| .................ucuggacucgucacacuaggCgc.............................................. | 1 | 1 | seq |
| .................ucuggacucgucacacuaggugcg............................................. | 1 | 0 | seq |
| ..............................................gcuccaagcgugacuaguccC................... | 1 | 1 | seq |
| ..............................................gcCccaagcgugacuagucca................... | 1 | 1 | seq |
| ................................................uccaagUgugacuagucc.................... | 2 | 1 | seq |
| ................................................uccaagcgugacuagucc.................... | 31 | 0 | seq |
| ................................................uccaagcgugacuagucU.................... | 28 | 1 | seq |
| ................................................uccaagcgugacuagucA.................... | 7 | 1 | seq |
| ................................................Cccaagcgugacuagucc.................... | 2 | 1 | seq |
| ................................................Gccaagcgugacuagucc.................... | 2 | 1 | seq |
| ................................................uccaagcgugacuagucG.................... | 1 | 1 | seq |
| ................................................uccaagcgugacuaguccG................... | 4 | 1 | seq |
| ................................................uccaagcgugacuaguccU................... | 3 | 1 | seq |
| ................................................Cccaagcgugacuagucca................... | 2 | 1 | seq |
| ................................................Accaagcgugacuagucca................... | 3 | 1 | seq |
| ................................................uUcaagcgugacuagucca................... | 1 | 1 | seq |
| ................................................uccaagcgugacuaguccC................... | 5 | 1 | seq |
| ................................................ucUaagcgugacuagucca................... | 1 | 1 | seq |
| ................................................uccaagcgugacuagucUa................... | 1 | 1 | seq |
| ................................................uccaagcgugacuaguUca................... | 2 | 1 | seq |
| ................................................uccaagcgugacuagucca................... | 115 | 0 | seq |
| ................................................uccaagcgugacuaguccaU.................. | 3 | 1 | seq |
| ................................................uccaagcgugacuaguccaC.................. | 1 | 1 | seq |
| ................................................uccaagcgugGcuaguccag.................. | 2 | 1 | seq |
| ................................................uccaagcCugacuaguccag.................. | 1 | 1 | seq |
| ................................................Accaagcgugacuaguccag.................. | 2 | 1 | seq |
| ................................................Cccaagcgugacuaguccag.................. | 1 | 1 | seq |
| ................................................uccaagcgugacCaguccag.................. | 1 | 1 | seq |
| ................................................uccaagcgugacuaguccGg.................. | 1 | 1 | seq |
| ................................................uccaagcgugacuGguccag.................. | 1 | 1 | seq |
| ................................................uccaagcgugacuagucAag.................. | 1 | 1 | seq |
| ................................................uccaagcgugacuaUuccag.................. | 2 | 1 | seq |
| ................................................uccaagcgugacuaguccag.................. | 86 | 0 | seq |
| ................................................uccaGgcgugacuaguccag.................. | 1 | 1 | seq |
| ................................................uccaagcgCgacuaguccag.................. | 1 | 1 | seq |
| ................................................uccaagcgugacuagCccag.................. | 1 | 1 | seq |
| ................................................uccaagcgugacuaguccaA.................. | 10 | 1 | seq |
| ................................................Accaagcgugacuaguccaga................. | 3 | 1 | seq |
| ................................................uccaagcgugacuaguccaga................. | 119 | 0 | seq |
| ................................................Gccaagcgugacuaguccaga................. | 1 | 1 | seq |
| ................................................uccaagcgugacuaguccagG................. | 5 | 1 | seq |
| ................................................uccaagcgugacuaguccagU................. | 1 | 1 | seq |
| ................................................uccaagcgugacuaUuccaga................. | 1 | 1 | seq |
| ................................................uccaagcguAacuaguccaga................. | 1 | 1 | seq |
| ................................................uccaagcgugacuaguccagaa................ | 375 | 0 | seq |
| ................................................uccaagcgugacuaguccagGa................ | 1 | 1 | seq |
| ................................................uccaagcgugacuaguccagUa................ | 1 | 1 | seq |
| ................................................uccaagcgugacuaguccagaG................ | 7 | 1 | seq |
| ................................................uccGagcgugacuaguccagaa................ | 1 | 1 | seq |
| ................................................uccaagcgugacuagucAagaa................ | 2 | 1 | seq |
| ................................................Cccaagcgugacuaguccagaa................ | 2 | 1 | seq |
| ................................................uccaaCcgugacuaguccagaa................ | 1 | 1 | seq |
| ................................................uccaagcgugacuaguccaUaa................ | 1 | 1 | seq |
| ................................................uccaagcgugacuaguccCgaa................ | 1 | 1 | seq |
| ................................................uccaagcgugacuaAuccagaa................ | 1 | 1 | seq |
| ................................................uccaagcgugCcuaguccagaa................ | 1 | 1 | seq |
| ................................................Accaagcgugacuaguccagaa................ | 7 | 1 | seq |
| ................................................uccaagcCugacuaguccagaa................ | 1 | 1 | seq |
| ................................................uGcaagcgugacuaguccagaa................ | 1 | 1 | seq |
| ................................................uccaagcgugacuaguccagaC................ | 2 | 1 | seq |
| ................................................ucUaagcgugacuaguccagaa................ | 3 | 1 | seq |
| ................................................uccaagcgugacuaUuccagaa................ | 1 | 1 | seq |

```
                                                novel-nve-miR-55_guide
                novel-nve-miR-55_star
uucucccucgcagcugcucuggacucgucacacuagguugcgcuugagcuccaagcgugacuaguccagaacggaugcgaggagacu

................................................uccaagcgCgacuaguccagaa................    1    1    seq
................................................uccaGgcgugacuaguccagaa................    1    1    seq
................................................uccaagcgugacAaguccagaa................    2    1    seq
................................................uccaagcgugGcuaguccagaa................    5    1    seq
................................................uccaagcgugacuaguccagaU................    9    1    seq
................................................Cccaagcgugacuaguccagaac...............    1    1    seq
................................................uccaagcgugacuaguccagaaU...............    134  1    seq
................................................uccaagUgugacuaguccagaac...............    1    1    seq
................................................uccaagcgugacuaguccagaaA...............    17   1    seq
................................................uccaagcgugacuaguccGgaac...............    1    1    seq
................................................uccaagcgugacuagAccagaac...............    1    1    seq
................................................uccaagcgugacuaguccagaac...............    98   0    seq
................................................uccaagcgAgacuaguccagaac...............    1    1    seq
................................................uccaagcgugacuaguccagaaG...............    3    1    seq
................................................uccaagcgugGcuaguccagaac...............    2    1    seq
................................................Accaagcgugacuaguccagaac...............    3    1    seq
................................................uccaagcgugacuaguccagaacU..............    175  1    seq
................................................uccaagcgugacuaguccagaacA..............    21   1    seq
................................................ucAaagcgugacuaguccagaacg..............    1    1    seq
................................................uccaagcgugacuagucUagaacg..............    1    1    seq
................................................uccaagcgugacuaguccagaacg..............    10   0    seq
................................................uccaagcgugacuaguccagaacC..............    19   1    seq
................................................uUcaagcgugacuaguccagaacg..............    1    1    seq
.................................................ccaagcgugacuaguccaU..................    2    1    seq
.................................................ccaagcgugacuaguccaga.................    1    0    seq
.................................................ccaagcgugacuaguccagaaG...............    1    1    seq
..................................................caagcgugacuaguccag..................    3    0    seq
..................................................caagcgugacuaguccaga.................    3    0    seq
..................................................caagcgugacuaguccUgaa................    3    1    seq
..................................................caagcgugacuaguccagaa................    7    0    seq
..................................................caagcgugacuaguccCgaac...............    1    1    seq
..................................................caagcgugacuaguccagaaU...............    2    1    seq
..................................................caagcgugacuaguccagaac...............    4    0    seq
..................................................caagcgugacuaguccagaacU..............    7    1    seq
..................................................caGgcgugacuaguccagaacg..............    1    1    seq
..................................................caagcgugacuaguccagaacA..............    4    1    seq
..................................................caagcgugacuaguccagaacC..............    1    1    seq
..................................................caagcgugGcuaguccagaacg..............    1    1    seq
..................................................Uaagcgugacuaguccagaacg..............    1    1    seq
..................................................caagcgugacuaguccagaacg..............    19   0    seq
..................................................caagcgugacuaguccUgaacg..............    1    1    seq
...................................................aagcgugacuaguccaga.................    2    0    seq
...................................................aagcgugacuaguccaUaacg..............    1    1    seq
...................................................aagcgugacuaguccagaacg..............    9    0    seq
...................................................aagcgugacuaguccagaacU..............    1    1    seq
...................................................aagcgugacuaguccagaacA..............    1    1    seq
....................................................agcgugacuaguccagaaU...............    1    1    seq
....................................................agcgugacuaguccagaac...............    4    0    seq
....................................................Ggcgugacuaguccagaacg..............    1    1    seq
....................................................agcgugacuagCccagaacg..............    1    1    seq
....................................................agcgugacuaguccagaacA..............    8    1    seq
....................................................agcgugacuaguccagaacC..............    1    1    seq
....................................................agcgugacuaguccagaacU..............    11   1    seq
....................................................agcgugacuaguccagaacg..............    20   0    seq
....................................................agcgugacuaguccagaUcg..............    1    1    seq
......................................................cgugacuaguccagaacg..............    1    0    seq
........................................................ugacuaguccagaacggauA..........    1    1    seq
........................................................ugacuaguccagaacggaug..........    1    0    seq
```

miRBase precursor : novel-nve-miR-56
Total read count : 184
novel-nve-miR-56_guide read count : 160
novel-nve-miR-56_star read count : 24
remaining reads : 0

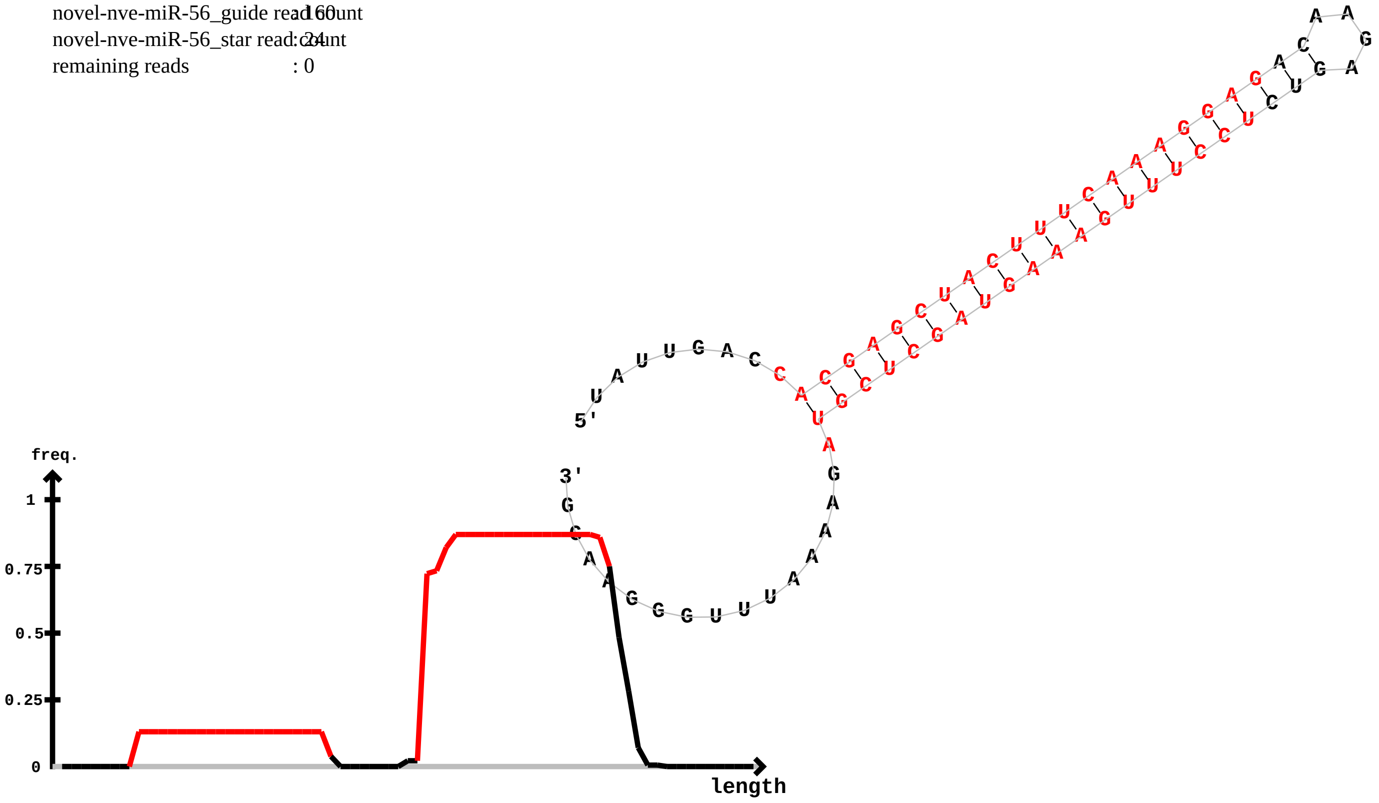


```
                   novel-nve-miR-56_star
                                          novel-nve-miR-56_guide
5'- uauugaccacgagcuacuuucaaaggagacaagagucuccuuugaaaguagcucguagaaaauuugggaagg  -3'    exp
    ........((((((((((((((((((((((....))))))))))))))))))))))................    reads      mm         sample
    .......cacgagcuacuuucaaagga.............................................    15         0          seq
    .......cacgUgcuacuuucaaagga.............................................    1          1          seq
    .......cacgagcuacuuCcaaagga.............................................    1          1          seq
    .......cacgagcuacuuucaaaggag............................................    1          0          seq
    .......cacgagcuacuuucaaaggaC............................................    2          1          seq
    .......cacgagcuacuuucaaaggaU............................................    4          1          seq
    ...................................ucuccuuugaaaguagcucg.................    1          0          seq
    ...................................ucuccuuugaaaguagcucgua...............    2          0          seq
    ...................................ucuccuuugaaaguagcucguag..............    1          0          seq
    .....................................uccuuugaaaguagcucg.................    1          0          seq
    .....................................uccuuugaaaguagcucgu................    17         0          seq
    .....................................uccuuugaaaguagcucgC................    1          1          seq
    .....................................uccuuugaaaguagcucgA................    2          1          seq
    .....................................uccuuugaaaguGgcucgua...............    1          1          seq
    .....................................uccuuugaaaguagcucguG...............    12         1          seq
    .....................................uccuuugaaaguagcucgua...............    33         0          seq
    .....................................uccuuugaaaguagcucguU...............    1          1          seq
    .....................................uccuuugaaaguagcucguaU..............    1          1          seq
    .....................................uccuuugaaaguaUcucguag..............    1          1          seq
    .....................................uccuuugaaaguagcucguag..............    8          0          seq
    .....................................uccuuugaaaguagcucguGg..............    10         1          seq
    .....................................ucAuuugaaaguagcucguaga.............    2          1          seq
    .....................................uccuuugaaaguagcucguGga.............    1          1          seq
    .....................................uccuuugaaaguagcucguaga.............    27         0          seq
    .....................................uccuCugaaaguagcucguaga.............    1          1          seq
    .....................................uccuuugaaaguagcucguagaa............    10         0          seq
    ......................................Ucuuugaaaguagcucguag..............    1          1          seq
    ......................................ccuuugaaaguagcucguaga.............    1          0          seq
    .......................................cuuugaaaguagcucguag..............    2          0          seq
    .......................................cuuugaaaguagcucguGg..............    7          1          seq
    .......................................cuuugaaaguagcucguaga.............    7          0          seq
    ........................................uuugaaaguagcucguGg..............    6          1          seq
    ........................................uuugaaaguagcucguagaa............    1          0          seq
    ........................................uuugaaaguagcucguagaU............    1          1          seq
```

novel-nve-miR-56_star

novel-nve-miR-56_guide

uauugaccacgagcuacuuucaaaggagacaagaguccuuugaaaguagcucguagaaaauuugggaagg

........................................uuugaaaguagcucguagaaaa.......... 1 0 seq

miRBase precursor : novel-nve-miR-57
Total read count : 289
novel-nve-miR-57_guide read count : 288
novel-nve-miR-57_star read count : 1
remaining reads : 0

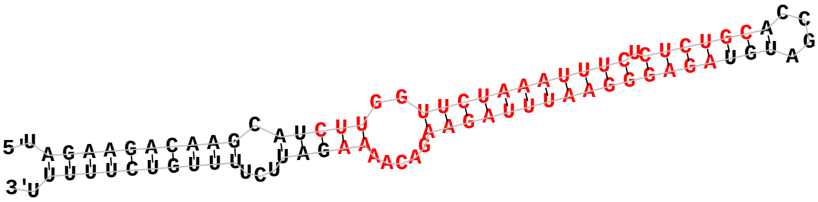

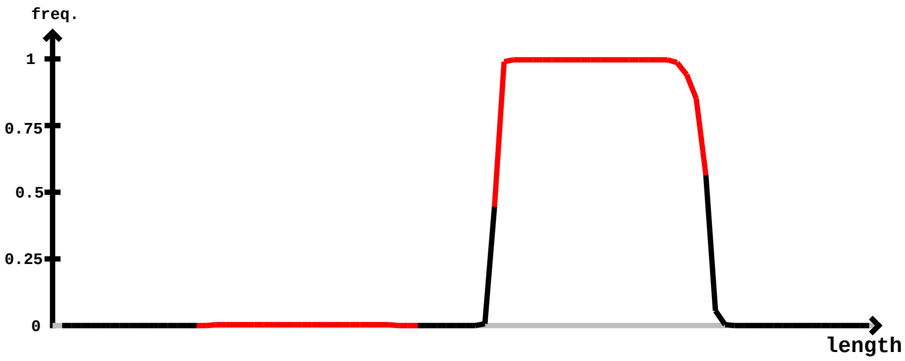


novel-nve-miR-57_star

novel-nve-miR-57_guide

```
5'- uagaagacaagcaucuugguucuaaauuucucucugcaccgauguagagggaauuuagaagacaaaagauucuuuugucuuuuu  -3'   exp
    .((((((((((.(((((..(((((((((((.(((((((....)))))))))))))))))).....)))))...))))))))))).    reads     mm     sample
    ...............uugguucuaaauuucucuA..................................................    1         1      seq
    ...........................................guagagggaauuuagaagacaa...................    1         0      seq
    ...........................................guagagggaauuuagaagacaaaa.................    1         0      seq
    ............................................uagagggaauuuagaagac.....................    1         0      seq
    ............................................uagagggaaCuuagaagaca....................    1         1      seq
    ............................................uagagggaauuuagaagaca....................    11        0      seq
    ............................................uagagggaauuuagaagacaU...................    1         1      seq
    ............................................uagagggaauuuagaagacaC...................    1         1      seq
    ............................................uagagggaauuuagaagacaa...................    13        0      seq
    ............................................uGgagggaauuuagaagacaa...................    1         1      seq
    ............................................uagaggUaauuuagaagacaa...................    1         1      seq
    ............................................uagagggaauuuagaagacaaC..................    3         1      seq
    ............................................uagagggaauuuaAaagacaaa..................    1         1      seq
    ............................................uagagggaauuuagaagacaaU..................    6         1      seq
    ............................................uagagggaauuuagaagacaGa..................    1         1      seq
    ............................................uagagggaauuuagaagacaaa..................    39        0      seq
    ............................................uagagggaauuuagaagacaaaC.................    2         1      seq
    ............................................uagagggaauuuagaagacaaaU.................    9         1      seq
    ............................................uagagggaauuuagaagacaaaa.................    29        0      seq
    ............................................uagagggaauuuagaagacaaaaU................    6         1      seq
    ............................................uagagggaauuuagaagacaaaaA................    1         1      seq
    .............................................agagggaauuuagaagac.....................    1         0      seq
    .............................................agagggUauuuagaagac.....................    1         1      seq
    .............................................agagggaauuuagaagaca....................    1         0      seq
    .............................................agagggaauuuagaagacaa...................    4         0      seq
    .............................................agagggaauuuaUaagacaa...................    2         1      seq
    .............................................agagggaauuuagaagacaaU..................    1         1      seq
    .............................................agagggaauuuagaagacaaa..................    27        0      seq
    .............................................agagggaaCuuagaagacaaa..................    1         1      seq
    .............................................Ggagggaauuuagaagacaaa..................    1         1      seq
    .............................................agagggaauuuagaaUacaaa..................    1         1      seq
    .............................................agGgggaauuuagaagacaaa..................    1         1      seq
    .............................................agagggaauuuagaagacaGa..................    1         1      seq
    .............................................agagggaauuuagaagacaaaa.................    80        0      seq
```

```
              novel-nve-miR-57_star
                                             novel-nve-miR-57_guide
uagaagacaagcaucuugguucuaaauuucucucugcaccgauguagagggaauuuagaagacaaaagauucuuuugucuuuuu

.............................................agaggCaauuuagaagacaaaa.................    1    1    seq
.............................................agagggaauuuagaagacaaCa.................    1    1    seq
.............................................agagggaauuuagGagacaaaa.................    1    1    seq
.............................................agagggaauuuagaagacGaaa.................    1    1    seq
.............................................agagggaauuuagaagacaaaU.................    10   1    seq
.............................................agagggGauuuagaagacaaaa.................    1    1    seq
.............................................Ggagggaauuuagaagacaaaa.................    2    1    seq
.............................................agagggaauuuagaagacaaaC.................    9    1    seq
.............................................agagggaauuuagaagacaaaaU................    8    1    seq
.............................................agagggaauuuagaagacaaaagG...............    1    1    seq
..............................................gagggaauuuagaagacaa...................    2    0    seq
```

miRBase precursor : novel-nve-miR-58
Total read count : 373
novel-nve-miR-58_guide read count : 371
novel-nve-miR-58_star read count : 2
remaining reads : 0

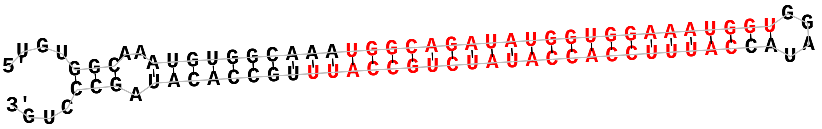


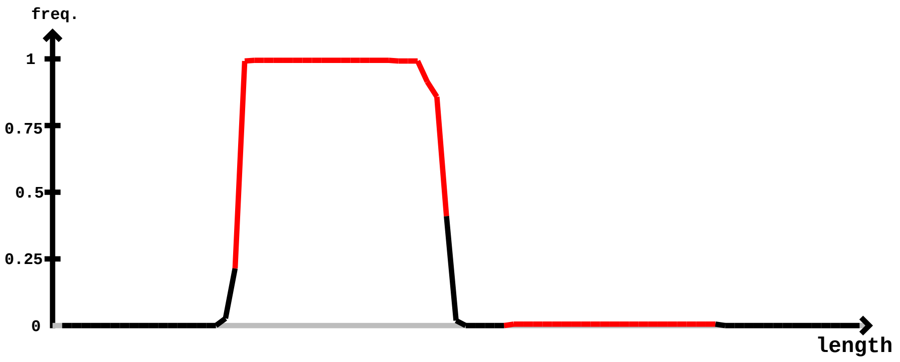


novel-nve-miR-58_guide
novel-nve-miR-58_star

| 5'- uguggcaaauguggcaaauggcagauaugguggaaaugguggauaccauuuccaccauaucugccauuugccacauagcccug -3' | exp | | |
|---|---|---|---|
| ...(((..((((((((((((((((((((((((((((((((....)))))))))))))))))))))))))))))))).)))... | reads | mm | sample |
| ................aauggcagauauggugga................................................. | 1 | 0 | seq |
| ................aauggcagauaugguggaaau.............................................. | 1 | 0 | seq |
| ................aauggcagauaugguggaaaug............................................. | 3 | 0 | seq |
| ................aauggcagauaugguggaaaugA............................................ | 1 | 1 | seq |
| ................aauggcagauaugguggaaaugg............................................ | 4 | 0 | seq |
| .................auggcagauaugguggaaau.............................................. | 10 | 0 | seq |
| .................auggcagauaugguggaaaug............................................. | 2 | 0 | seq |
| .................Guggcagauaugguggaaaugg............................................ | 1 | 1 | seq |
| .................auggcagauaugguggaaaugg............................................ | 47 | 0 | seq |
| .................auggAagauaugguggaaaugg............................................ | 1 | 1 | seq |
| .................auggUagauaugguggaaaugg............................................ | 1 | 1 | seq |
| .................auggcagauaugguggGaaugg............................................ | 2 | 1 | seq |
| .................auggcagauaugguggaaaugA............................................ | 3 | 1 | seq |
| .................auggcagauaCgguggaaaugg............................................ | 1 | 1 | seq |
| .................auggcagauaugguggaaGugg............................................ | 1 | 1 | seq |
| .................auggcagauaugguggaaauggu........................................... | 1 | 0 | seq |
| ..................uggcagauaugguggaaaC.............................................. | 2 | 1 | seq |
| ..................uggUagauaugguggaaau.............................................. | 1 | 1 | seq |
| ..................uggcagaGaugguggaaau.............................................. | 1 | 1 | seq |
| ..................uggcagauaugguggaaau.............................................. | 14 | 0 | seq |
| ..................uggcagauaugguggaaaug............................................. | 13 | 0 | seq |
| ..................uggcagauaugguggaaauA............................................. | 3 | 1 | seq |
| ..................uggcagauauUguggaaaugg............................................ | 1 | 1 | seq |
| ..................uggcagauaugguggaaaugC............................................ | 2 | 1 | seq |
| ..................ugUcagauaugguggaaaugg............................................ | 1 | 1 | seq |
| ..................uggcagUuaugguggaaaugg............................................ | 1 | 1 | seq |
| ..................uggcagauauggCggaaaugg............................................ | 1 | 1 | seq |
| ..................uggcagauaugguggaaaugA............................................ | 11 | 1 | seq |
| ..................uggcagauaugguggaaaugU............................................ | 4 | 1 | seq |
| ..................uggcagauaugguCgaaaugg............................................ | 2 | 1 | seq |
| ..................uUgcagauaugguggaaaugg............................................ | 1 | 1 | seq |
| ..................uggcagauaugguggaaaugg............................................ | 78 | 0 | seq |
| ..................uggcagaCaugguggaaaugg............................................ | 2 | 1 | seq |
| ..................uggcaAauaugguggaaaugg............................................ | 1 | 1 | seq |

```
                  novel-nve-miR-58_guide
                                              novel-nve-miR-58_star
uguggcaaauguggcaaauggcagauaugguggaaaugguggauaccauuuccaccauaucugccauuugccacauagcccug

..................uggcagGuaugguggaaauggu...........................................    1      1      seq
..................uggcagauaugguggaaauggu...........................................    116    0      seq
..................Gggcagauaugguggaaauggu...........................................    2      1      seq
..................uggcagauaugguggaaUuggu...........................................    1      1      seq
..................uCgcagauaugguggaaauggu...........................................    1      1      seq
..................uggcagauaugguggaaauggG...........................................    2      1      seq
..................Aggcagauaugguggaaauggu...........................................    1      1      seq
..................uggcagauaugguggaaauggA...........................................    10     1      seq
..................uggcagauaugguggaaauggC...........................................    10     1      seq
..................uggcagauaugguggaaaugguC..........................................    1      1      seq
..................uggcagauaugguggaaaugguU..........................................    6      1      seq
...................Ugcagauaugguggaaauggu...........................................    1      1      seq
..............................................cauuuccaccauaucugccaCu...............    2      1      seq
```

miRBase precursor : novel-nve-miR-59
Total read count : 1378
novel-nve-miR-59_guide read count : 1332
novel-nve-miR-59_star read count : 22
remaining reads : 24

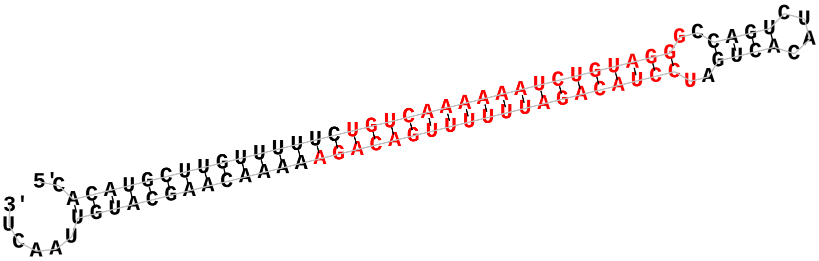

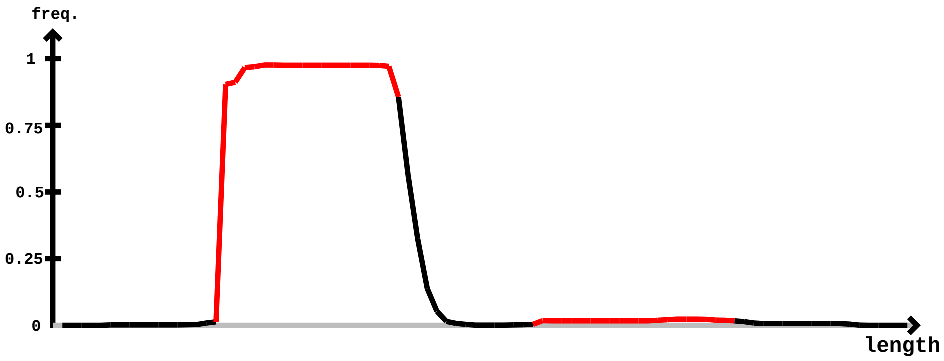


```
                    novel-nve-miR-59_guide
                                                 novel-nve-miR-59_star
5'- cacaugcuuguuuuucugucaaaaaaucuguagggccagucuacacugauccuacagauuuuuugacagaaaaacaagcauguuaacu   -3'   exp
    .(((((((((((((((((((((((((((((((((..((((....))))..))))))))))))))))))))))))))))))))).....    reads   mm   sample
    ....ugcuuguuuuucugucaa..................................................................    2       0    seq
    ............uuucugucaaaaaaucugua........................................................    1       0    seq
    .............uucugucaaaaaaucuguagg......................................................    1       0    seq
    ..............ucugucaaaaaaucuguag.......................................................    4       0    seq
    ..............ucugucaaaaaaucuguaA.......................................................    1       1    seq
    ..............ucugucaaaaaaucuguagg......................................................    3       0    seq
    ...............cugucaaaaaaucuguagg......................................................    3       0    seq
    ...............cugucaaaaaaucuguagA......................................................    2       1    seq
    ...............cugucaaaaaaucuguaggg.....................................................    1       0    seq
    ................ugucaaaaaaucuguagA......................................................    12      1    seq
    ................Ggucaaaaaaucuguagg......................................................    1       1    seq
    ................ugucaaaaaaucuguUgg......................................................    1       1    seq
    ................ugucaaaaaaCcuguagg......................................................    1       1    seq
    ................ugucaaaGaaucuguagg......................................................    5       1    seq
    ................ugAcaaaaaaucuguagg......................................................    1       1    seq
    ................ugucaaaaaaucuguGgg......................................................    2       1    seq
    ................ugucGaaaaaucuguagg......................................................    1       1    seq
    ................ugucaaaaaauUuguagg......................................................    1       1    seq
    ................ugucaaaaaaucuguagg......................................................    117     0    seq
    ................uAucaaaaaaucuguagg......................................................    1       1    seq
    ................ugucaaaaaaucuguagU......................................................    3       1    seq
    ................ugucaaaaGaucuguagg......................................................    2       1    seq
    ................Agucaaaaaaucuguagg......................................................    1       1    seq
    ................ugucaaaGaaucuguaggg.....................................................    9       1    seq
    ................ugucaGaaaaucuguaggg.....................................................    2       1    seq
    ................ugucaaGaaaucuguaggg.....................................................    2       1    seq
    ................uguUaaaaaaucuguaggg.....................................................    4       1    seq
    ................ugucaaaaaaucCguaggg.....................................................    4       1    seq
    ................ugucGaaaaaucuguaggg.....................................................    2       1    seq
    ................Ggucaaaaaaucuguaggg.....................................................    2       1    seq
    ................ugucaaaaaaucuguGggg.....................................................    1       1    seq
    ................ugucaaaUaaucuguaggg.....................................................    1       1    seq
    ................ugucaaaaaaucuguaggU.....................................................    16      1    seq
    ................ugucaaaaaauUuguaggg.....................................................    3       1    seq
```

```
                novel-nve-miR-59_guide
                                                 novel-nve-miR-59_star
cacaugcuuguuuuucugucaaaaaaucuguagggccagucuacacugauccuacagauuuuuugacagaaaaacaagcauguuaacu

................ugucaaaaaaucuguaggg.....................................................    305    0    seq
................ugucaaaaaaCcuguaggg.....................................................    2      1    seq
................Agucaaaaaaucuguaggg.....................................................    4      1    seq
................ugucCaaaaaucuguaggg.....................................................    2      1    seq
................ugucaaaaaGucuguaggg.....................................................    2      1    seq
................ugucaaaaaaucuguaggC.....................................................    1      1    seq
................ugucaaaCaaucuguaggg.....................................................    1      1    seq
................ugucaaaaaaucuUuaggg.....................................................    2      1    seq
................ugucaaaaGaucuguaggg.....................................................    5      1    seq
................Cgucaaaaaaucuguaggg.....................................................    1      1    seq
................ugucUaaaaaucuguaggg.....................................................    1      1    seq
................ugucaaaaaaucuguagCg.....................................................    1      1    seq
................ugucaaaaaaucuguaggA.....................................................    29     1    seq
................ugucaaaaaaucCguagggc....................................................    1      1    seq
................ugucaaaaaaucugGagggc....................................................    1      1    seq
................Agucaaaaaaucuguagggc....................................................    4      1    seq
................ugucaaaaaGucuguagggc....................................................    2      1    seq
................ugucaaaaaaucuguaAggc....................................................    1      1    seq
................ugucaUaaaaucuguagggc....................................................    1      1    seq
................ugucaaaaaaucuguagggU....................................................    62     1    seq
................ugucaaaaaaucuguagggA....................................................    21     1    seq
................uguUaaaaaaucuguagggc....................................................    1      1    seq
................ugAcaaaaaaucuguagggc....................................................    1      1    seq
................ugucaaaaGaucuguagggc....................................................    6      1    seq
................Cgucaaaaaaucuguagggc....................................................    2      1    seq
................ugucaaaaaaucuguagggc....................................................    190    0    seq
................ugucaaaaaauAuguagggc....................................................    1      1    seq
................ugucaaaaaaucuguaUggc....................................................    1      1    seq
................ugucaaaGaaucuguagggc....................................................    5      1    seq
................ugucaaaUaaucuguagggc....................................................    1      1    seq
................ugucaaaaaaucuguagggG....................................................    5      1    seq
................ugucaGaaaaucuguagggc....................................................    2      1    seq
................ugucaaaaaauUuguagggc....................................................    1      1    seq
................ugucaaaaGaucuguagggcc...................................................    2      1    seq
................ugCcaaaaaaucuguagggcc...................................................    1      1    seq
................ugucaaaaaaucuguGgggcc...................................................    1      1    seq
................ugucaaaaaauAuguagggcc...................................................    1      1    seq
................ugucaaaaaaucuguagggcc...................................................    117    0    seq
................uAucaaaaaaucuguagggcc...................................................    1      1    seq
................ugucaaaaaaucuguagggcG...................................................    2      1    seq
................ugucaaGaaaucuguagggcc...................................................    1      1    seq
................ugucaaaaaaucuguagggcA...................................................    27     1    seq
................ugucaGaaaaucuguagggcc...................................................    1      1    seq
................ugucaaaaaaucuguagggcU...................................................    70     1    seq
................Agucaaaaaaucuguagggcc...................................................    2      1    seq
................ugucaaaaaaucuguagggccU..................................................    26     1    seq
................ugucaaaaaaucuguagggcUa..................................................    1      1    seq
................ugAcaaaaaaucuguagggcca..................................................    1      1    seq
................Agucaaaaaaucuguagggcca..................................................    3      1    seq
................ugucaaaaaaucuguagggUca..................................................    2      1    seq
................ugucaaUaaaucuguagggcca..................................................    1      1    seq
................ugCcaaaaaaucuguagggcca..................................................    3      1    seq
................ugucaaaaaaucuguagggccC..................................................    9      1    seq
................ugucaaaaaaucuguaggUcca..................................................    1      1    seq
................ugucaaaaaaucuguagggccG..................................................    3      1    seq
................ugucaaaaaaucuguagggcca..................................................    57     0    seq
................ugucaaaaaaucuguagggccaU.................................................    8      1    seq
................ugucaaaaaaucuguagggccag.................................................    16     0    seq
................ugucaaaaaGucuguagggccag.................................................    2      1    seq
................ugucaaaaaaucuguagggccaC.................................................    2      1    seq
................ugucaaaaaaucuguagggccaCu................................................    1      1    seq
................ugucaaaaaaucuguagggccaUu................................................    5      1    seq
.................gAcaaaaaaucuguagggc....................................................    1      1    seq
.................gucaaaaaaucuguagggcc...................................................    2      0    seq
.................gucaaaaaaucuguagggcU...................................................    2      1    seq
.................gucaaaaaaucCguagggcca..................................................    1      1    seq
.................gucaaaaaaucuguagggcca..................................................    2      0    seq
.................gucaaaaaaucAguagggcca..................................................    1      1    seq
.................gucaaaaaaucuguagggccU..................................................    1      1    seq
```

```
                novel-nve-miR-59_guide
                                                 novel-nve-miR-59_star
cacaugcuuguuuuucugucaaaaaaucuguagggccagucuacacugauccuacagauuuuuugacagaaaacaagcauguuaacu

.................gucaaaaaaucuguagggccag.................................................    1    0    seq
..................ucaaaaaaucuguagggU....................................................    11   1    seq
..................Gcaaaaaaucuguagggc....................................................    1    1    seq
..................ucaaaaaaucuguagggc....................................................    8    0    seq
..................ucaaaaaaucuguagggcc...................................................    17   0    seq
..................ucaaaaaaucuguaggAcc...................................................    1    1    seq
..................ucaaaaaaucuguagggcA...................................................    1    1    seq
..................ucaaaaaaucuguagggcU...................................................    6    1    seq
..................ucaaaaaaucuguagggcca..................................................    3    0    seq
..................ucaaaGaaucuguagggccag.................................................    1    1    seq
..................ucaaaaaaucuguagggccag.................................................    15   0    seq
..................Acaaaaaaucuguagggccag.................................................    1    1    seq
..................ucaaaaaaucuguagggccaA.................................................    1    1    seq
..................ucaaaaaaucuguagggccaAu................................................    1    1    seq
..................ucaaaGaaucuguagggccagu................................................    3    1    seq
..................ucaaaaaaucuguagggccaguc...............................................    3    0    seq
..................ucaaaaaaucuguagggccagucu..............................................    3    0    seq
...................caaaaaaucuguagggcc...................................................    2    0    seq
...................caaaaaaucuguagggcca..................................................    1    0    seq
...................caaaaaaucuguagggccag.................................................    1    0    seq
....................aaaaaaucuguagggcca..................................................    1    0    seq
....................aaaaaaucuguagggccag.................................................    5    0    seq
....................aaaaaaucuguagggccaguU...............................................    2    1    seq
....................aaaaaaucuguagggccagucu..............................................    1    0    seq
......................aaaaucuguagggccagucuacacugau......................................    1    0    seq
..............................................ugauccuacagauuuuuuga......................    1    0    seq
...............................................gauccuacagauuuuuugacag...................    1    0    seq
................................................auccuacagauuuuuugac.....................    2    0    seq
.................................................uccuacagauuuuuugac.....................    2    0    seq
.................................................uccuacagauuuuuugaca....................    1    0    seq
.................................................uccuacagauuuuuugacag...................    1    0    seq
.................................................uccuacagauuuuuugacaA...................    1    1    seq
.................................................ucUuacagauuuuuugacaga..................    1    1    seq
.................................................uccuacagauuuuuugacagC..................    1    1    seq
.................................................uccuacagauuuuuugacaCa..................    2    1    seq
.................................................uccuacagauuuuuugacagCa.................    1    1    seq
.................................................uccuacagauuuuuugacaUaa.................    1    1    seq
.................................................uccuacagauuuuuugacagaC.................    1    1    seq
.................................................uccuacagauuuuuugacagaa.................    4    0    seq
.................................................uccuacagauuuuuugacagaaU................    2    1    seq
.................................................uccuacagauuuuuugacagaaa................    1    0    seq
.............................................................uuugacagaaaaacaagcaug......    1    0    seq
.............................................................uuugacagaaaaacaagcGug......    1    1    seq
.............................................................uuugacagaaaaacaagcauA......    1    1    seq
..............................................................uugacagaaaaacaagcaug......    2    0    seq
..............................................................uugacagaaaaacaagcaugG.....    1    1    seq
...............................................................ugacagaaaaacaagcau.......    3    0    seq
```

miRBase precursor : novel-nve-miR-62
Total read count : 182
novel-nve-miR-62_guide read count : 175
novel-nve-miR-62_star read count : 7
remaining reads : 0

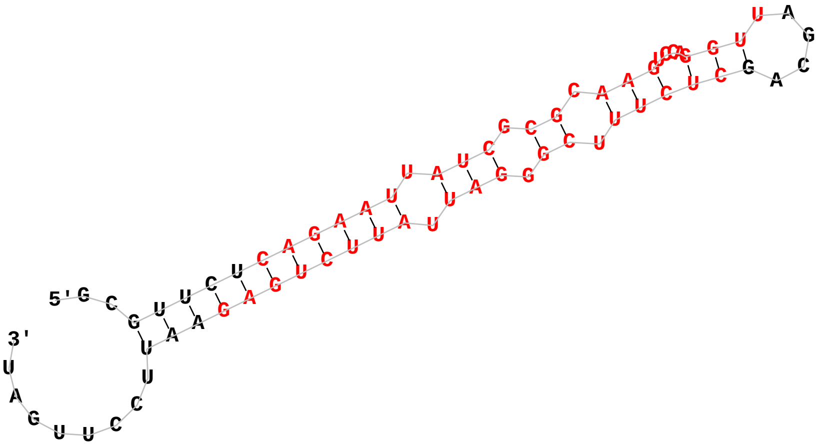

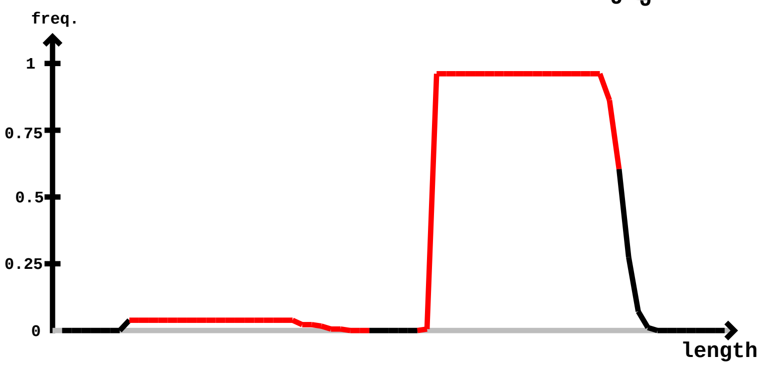


novel-nve-miR-62_guide

novel-nve-miR-62_star

```
5'- gcguucucagaauuaucgcgcaaguccagguuagcagcucuuucgggauuauucugagaauuccuugau   -3'    exp
    ..((((((((((.(((.((.(((....(((.....)))))).)).))).))))))))))..........    reads      mm         sample
    ......ucagaauuaucgcgcaag.............................................    3          0          seq
    ......ucagaauuaucgcgcaaguc...........................................    1          0          seq
    ......ucagaauuaucgcgcaagucc..........................................    1          0          seq
    ......ucagaauuaucgcgcaagucU..........................................    1          1          seq
    ......ucagaauuaucgcgcaaguccaC........................................    1          1          seq
    .....................................cucuuucgggauuauucugag...........    1          0          seq
    ......................................uAuuucgggauuauucug.............    1          1          seq
    ......................................ucuuucgggauuauucuA.............    1          1          seq
    ......................................Acuuucgggauuauucug.............    1          1          seq
    ......................................ucuuucgggauuauucug.............    15         0          seq
    ......................................ucuuucgggauuauucuga............    45         0          seq
    ......................................ucuuucgggauuauucugU............    1          1          seq
    ......................................Ccuuucgggauuauucuga............    1          1          seq
    ......................................Acuuucgggauuauucugag...........    2          1          seq
    ......................................ucuuucgggauuauucugUg...........    1          1          seq
    ......................................ucuuucgggauuauucugaC...........    1          1          seq
    ......................................ucuuucgggauuauucugaA...........    3          1          seq
    ......................................ucuuucgggauuauucugag...........    51         0          seq
    ......................................ucuuucgggauuauucugaU...........    1          1          seq
    ......................................ucuuucgggauuGuucugaga..........    1          1          seq
    ......................................ucuuucgggauuauucugaga..........    34         0          seq
    ......................................ucuuucgggauuauucugagG..........    1          1          seq
    ......................................ucuuucgggGuuauucugaga..........    1          1          seq
    ......................................ucuuucgggauuauucugagaU.........    2          1          seq
    ......................................ucuuucgggauuauucugagaC.........    1          1          seq
    ......................................ucCuucgggauuauucugagaa.........    1          1          seq
    ......................................ucuuucgggauuauucugagaa.........    6          0          seq
    ......................................ucuuucgggauuauucugagaG.........    1          1          seq
    ......................................ucuuucgggauuauucugagaCu........    1          1          seq
    ......................................ucuuucgggauuauucugagaUu........    1          1          seq
```

miRBase precursor : novel-nve-miR-63
Total read count : 190
novel-nve-miR-63_guide read count : 186
novel-nve-miR-63_star read count : 0
remaining reads : 4

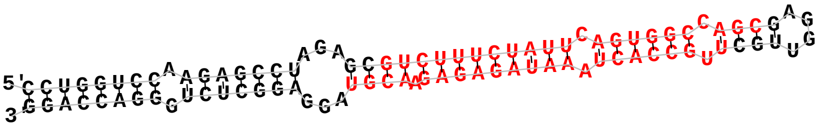

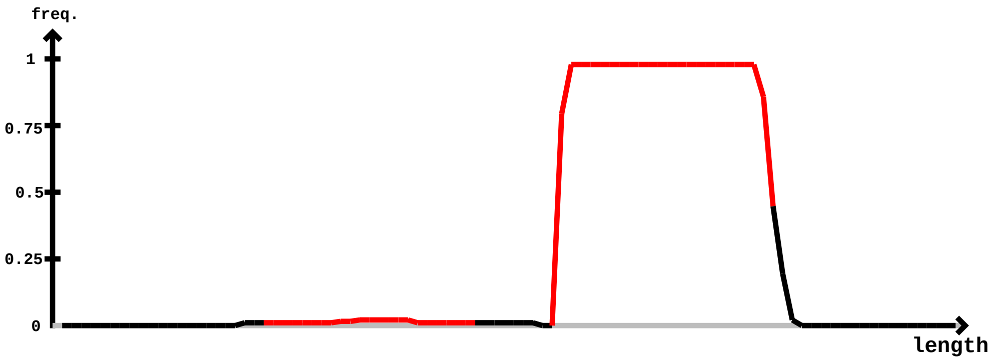


```
                         novel-nve-miR-63_star
                                                novel-nve-miR-63_guide
5'- ccugguccaagagccuagagcgucuuucuauucaguggccagcgagguugcuugccacuaaauagagagaacguaggaggcucugggaccagg   -3'    exp
    ((((((((.(((((((...(((((((((((((.((((((.((((....)))).)))))).))))))))).))))...))))))).))))))))   reads   mm   sample
    ..................agcgucuCucuauucagu.........................................................   2       1    seq
    ............................uauuUaguggccagcgagguu............................................   1       1    seq
    ..............................uuUaguggccagcgagguu............................................   1       1    seq
    ...................................................uugccacuaaauagagGgaac.....................   1       1    seq
    ...................................................uugccacuaaauagagagaac.....................   18      0    seq
    ...................................................uugccacuaaauagagagaaU.....................   4       1    seq
    ...................................................uugccacuaaauagagagaacC....................   7       1    seq
    ...................................................uugccacuaaauagagagaacg....................   14      0    seq
    ...................................................uugccacuaaauagagagaacA....................   5       1    seq
    ...................................................uugccacuaaauagagagaacU....................   51      1    seq
    ...................................................uugcUacuaaauagagagaacg....................   1       1    seq
    ...................................................uugccacuaaauagagagaacgu...................   23      0    seq
    ...................................................uugccacuaaauagagGacgu...................   1       1    seq
    ...................................................uugccacuaaauagagagaacgC...................   1       1    seq
    ...................................................uugccacuaaauagagUgaacgu...................   1       1    seq
    ...................................................uugccacuaaGuagagagaacgu...................   1       1    seq
    ...................................................uugccacuaaauagagagaacUu...................   3       1    seq
    ...................................................uugccacuaaauagagagaacgA...................   1       1    seq
    ...................................................uugccacuaaauagagagaacgua..................   6       0    seq
    ...................................................uugccacuaaauagagagaacguU..................   9       1    seq
    ...................................................uugccacuaaauagagagaacguaU.................   1       1    seq
    ...................................................uugccacuaaauagagagaacguaC.................   1       1    seq
    ...................................................uugccacuaaauagagagaacguag.................   2       0    seq
    ....................................................ugccacuaaauagagagaacgu...................   10      0    seq
    ....................................................ugccacuaaauagagagaacgC...................   2       1    seq
    ....................................................ugccacuaaauagagagaacUu...................   4       1    seq
    ....................................................ugccacuaaauagagagaacgA...................   1       1    seq
    ....................................................ugccacuaaauagagagaacgua..................   9       0    seq
    ....................................................ugccacuaaauagagagaacguU..................   7       1    seq
    ....................................................ugccacuaaauagagagaacguC..................   1       1    seq
    ....................................................uAccacuaaauagagagaacgua..................   1       1    seq
```

miRBase precursor : novel-nve-miR-66
Total read count : 334
novel-nve-miR-66_guide read count : 265
novel-nve-miR-66_star read count : 69
remaining reads : 0

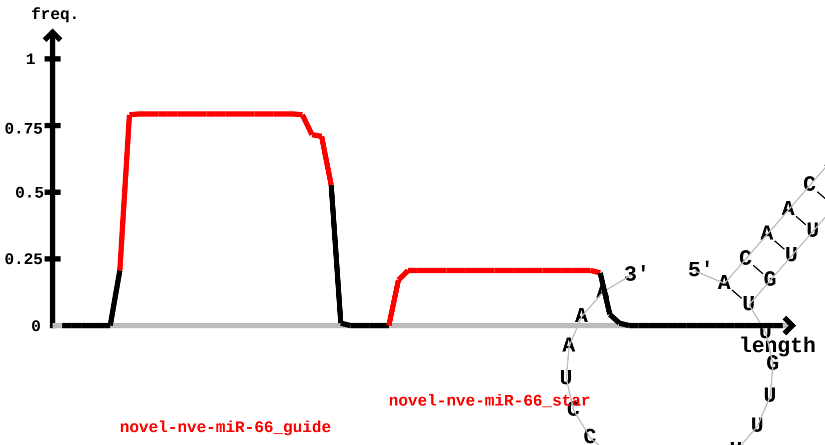


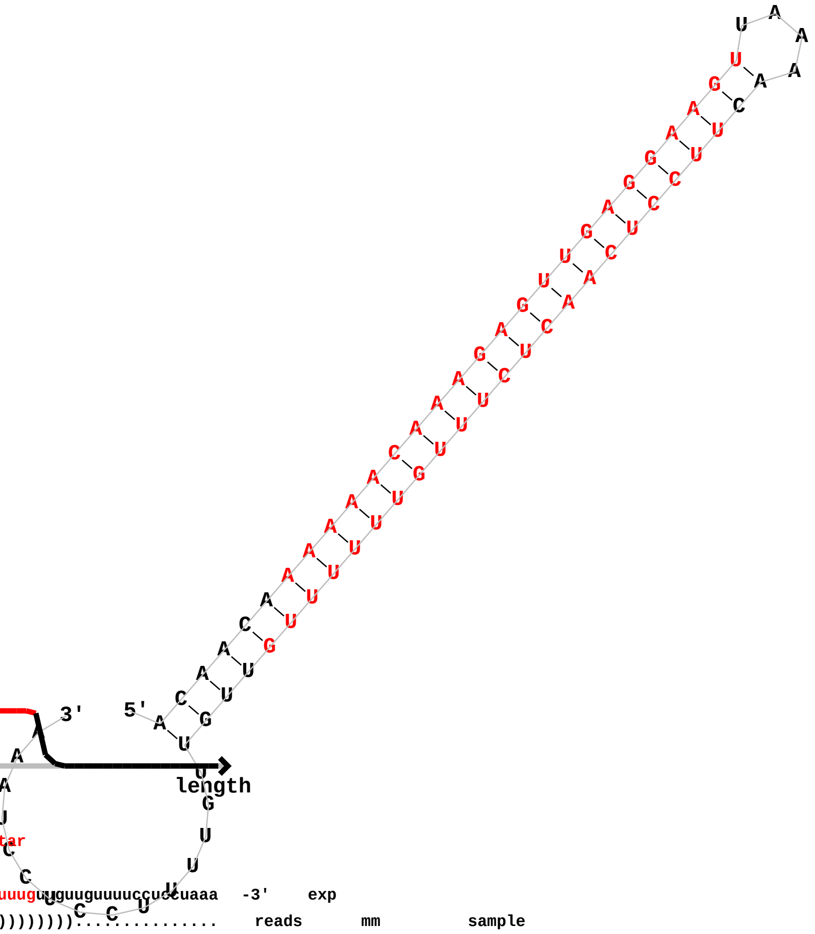


5'- acacaaaaaaacaaagaguugaggaaguuaaaacuuccucaacucuuuguuuuuuguuguuguuuuccuccuaaa -3' exp

| ((((((((((((((((((((((((((((....))))))))))))))))))))))))))))................ | reads | mm | sample |
|---|---|---|---|
| .....aaaaaacaaagaguugagga................................................... | 4 | 0 | seq |
| .....aGaaaacaaagaguugagga................................................... | 1 | 1 | seq |
| .....aaaaaacaaagaguugaggaa.................................................. | 1 | 0 | seq |
| .....aaaaaacaaagaguugaggaaU................................................. | 11 | 1 | seq |
| .....aaaaaacaaagaguugGggaag................................................. | 1 | 1 | seq |
| .....aaaaaacaaagaguugaggaag................................................. | 17 | 0 | seq |
| .....aaaaaacaaagaguugaggaaA................................................. | 2 | 1 | seq |
| .....aaaaaacaaagaguugaggaaC................................................. | 5 | 1 | seq |
| .....aaaaaacaaagGguugaggaag................................................. | 1 | 1 | seq |
| .....aaaaaacaaagaguugaggaGg................................................. | 1 | 1 | seq |
| .....aaaaaacaaagaguugaggaaUu................................................ | 4 | 1 | seq |
| .....aaaaaacaaagUguugaggaagu................................................ | 1 | 1 | seq |
| .....aaaaaacaGagaguugaggaagu................................................ | 2 | 1 | seq |
| .....aGaaaacaaagaguugaggaagu................................................ | 1 | 1 | seq |
| .....aaaaaacaaagaguugaggaagC................................................ | 6 | 1 | seq |
| .....aaaaaacaaagaguugaggaagA................................................ | 1 | 1 | seq |
| .....aaaaaacaaagaguugaggaagu................................................ | 7 | 0 | seq |
| .....aaaaaacaaagaguugaggaaguu............................................... | 2 | 0 | seq |
| .....aaaaaacaaagaguugaggaaguC............................................... | 1 | 1 | seq |
| ......aaaaacaaagaguugagA.................................................... | 1 | 1 | seq |
| ......aaaaacaaagagCugagga................................................... | 1 | 1 | seq |
| ......aaaaacaaagaguugagga................................................... | 18 | 0 | seq |
| ......aaaaacaaagaguugaggG................................................... | 1 | 1 | seq |
| ......aaaaacaaagaguugaggaa.................................................. | 1 | 0 | seq |
| ......aaaaacaaagaguugaggaag................................................. | 21 | 0 | seq |
| ......aaaaacaaagaguugaggaaA................................................. | 1 | 1 | seq |
| ......aaaaacaaagagGugaggaag................................................. | 1 | 1 | seq |
| ......aaaaacaaagaguCgaggaagu................................................ | 1 | 1 | seq |
| ......aaaaacaaagaguugaggaagA................................................ | 4 | 1 | seq |
| ......aaaaacaaagaguugaUgaagu................................................ | 1 | 1 | seq |
| ......aaaaacaaagaguugaggaagC................................................ | 17 | 1 | seq |
| ......aGaaacaaagaguugaggaagu................................................ | 2 | 1 | seq |
| ......aaaaacaaGgaguugaggaagu................................................ | 1 | 1 | seq |
| ......aaaaacaaagaguugaggaagG................................................ | 1 | 1 | seq |

```
                                  novel-nve-miR-66_star
      novel-nve-miR-66_guide
acaacaaaaaacaaagaguugaggaaguuaaaacuuccucaacucuuuguuuuuuguuguuguuuuccuccuaaa

......aaaaGcaaagaguugaggaagu...............................................    2          1          seq
......aaaaacaaagaguugGggaagu...............................................    2          1          seq
......aaaaacaaagaguugaggaGgu...............................................    2          1          seq
......aaaaacaaagaguuUaggaagu...............................................    1          1          seq
......aaaaacaaagaguuCaggaagu...............................................    1          1          seq
......aaaaacUaagaguugaggaagu...............................................    1          1          seq
......aaaaacaaagaguugaggaagu...............................................    110        0          seq
......aaaaacaGagaguugaggaagu...............................................    4          1          seq
.......aaaacaaagaguugaggaagu...............................................    1          0          seq
..................................uuccucaacucuuuguuuuuA....................    1          1          seq
..................................uuccucaacucuuuguuuuuuA...................    9          1          seq
..................................uuccuUaacucuuuguuuuuug...................    1          1          seq
..................................uuccucaacucCuuguuuuuug...................    1          1          seq
..................................uuccucaacucuuuguuuuuCg...................    1          1          seq
..................................uuccucaacucuuuguuuuuug...................    36         0          seq
..................................uuccucaacucuuuguuCuuug...................    1          1          seq
..................................uuccucGacucuuuguuuuuugu..................    3          1          seq
..................................uuccucaacucuuuguuuuuugu..................    1          0          seq
..................................uuccucaacucuuuguuuuuuguA.................    3          1          seq
...................................uccucaacucuuuguuuuuu....................    2          0          seq
...................................uccucaacucuuuguuuuuug...................    3          0          seq
...................................uccucaacucuuuguuuuuugu..................    4          0          seq
...................................uccucaacucuuuguuuuuugC..................    2          1          seq
...................................uccucaacucuuuguuuuuugA..................    1          1          seq
```

miRBase precursor : novel-nve-miR-68
Total read count : 1570
novel-nve-miR-68_guide read count : 21
novel-nve-miR-68_star read count : 1549
remaining reads : 0

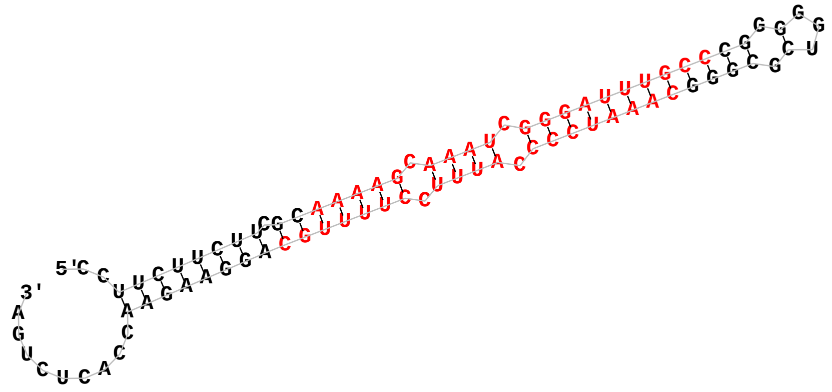

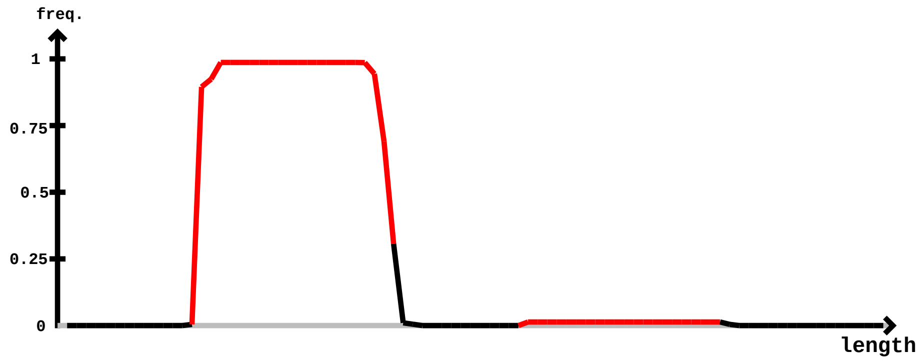


```
                                                      novel-nve-miR-68_guide
                 novel-nve-miR-68_star
5'- ccuucuucuucgcaaaagcaaaucgggauuugcccgggggucgcgggcaaauccccauuuccuuuugcaggaagaaccacucuga   -3'    exp
    ..((((((((.(((((((.((((.((((((((((((.(...).)))))))))))).)))).)))))))))))))))).........    reads      mm         sample
    ............caaaagGaaaucgggauu.......................................................    1          1          seq
    ............caaaagGaaaucgggauuugc....................................................    1          1          seq
    ............caaaagGaaaucgggauuugcc...................................................    5          1          seq
    .............aaaagGaaaucgggauuu.......................................................    66         1          seq
    .............aaaagcaaaucgggauuug.....................................................    2          0          seq
    .............aaaagGaaaucgggauuug.....................................................    386        1          seq
    .............aaaagcaaaucgggauuugc....................................................    1          0          seq
    .............aaaagGaaaucgggauuugc....................................................    522        1          seq
    .............aaaagAaaaucgggauuugcc...................................................    1          1          seq
    .............aaaagGaaaucgggauuugcc...................................................    412        1          seq
    .............aaaagcaaaucgggauuugcc...................................................    1          0          seq
    .............aaaagGaaaucgggauuugccc..................................................    7          1          seq
    ..............aaagGaaaucgggauuug.....................................................    9          1          seq
    ..............aaagGaaaucgggauuugc....................................................    19         1          seq
    ..............aaagGaaaucgggauuugcc...................................................    16         1          seq
    ..............aaagGaaaucgggauuugcccg.................................................    2          1          seq
    ...............aagAaaaucgggauuugc....................................................    3          1          seq
    ...............aagGaaaucgggauuugc....................................................    58         1          seq
    ...............aagGaaaucgggauuugcc...................................................    29         1          seq
    ...............aagAaaaucgggauuugcc...................................................    1          1          seq
    ...............aagGaaaucgggauuugccc..................................................    1          1          seq
    ...............aagGaaaucgggauuugcccg.................................................    6          1          seq
    ...............................................caaauccccauuuccuuuugU.................    3          1          seq
    ...............................................caaauccccauuuccuuuugc.................    9          0          seq
    ...............................................caaauccccauuuccCuuugc.................    1          1          seq
    ...............................................caaauccccauuuccuuuugA.................    1          1          seq
    ...............................................caaauccccauuucUuuuugca................    1          1          seq
    ...............................................caaauccccauuuccuuuugcU................    3          1          seq
    ...............................................caaauccccauuuccuuuugca................    3          0          seq
```

miRBase precursor : novel-nve-miR-69
Total read count : 468
novel-nve-miR-69_guide read count : 468
novel-nve-miR-69_star read count : 0
remaining reads : 0

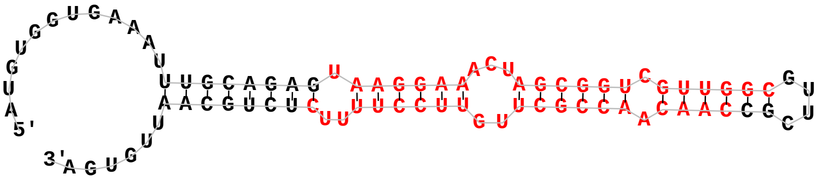

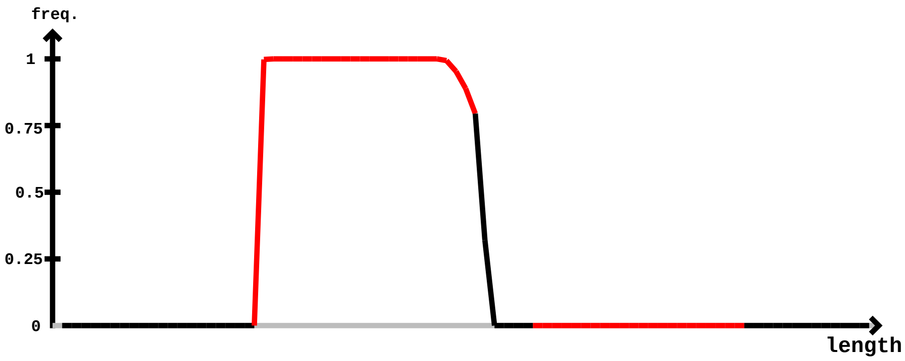


novel-nve-miR-69_guide

novel-nve-miR-69_star

```
5'- auguggugaaauuugcagaguaaggaaacuagcggucguuggcguucgccaacaaccgcuuguuccuuuucucugcaauuguga  -3'    exp
    ............((((((((.((((((...((((((.((((((....)))))).))))))..))))))..))))))))......    reads      mm        sample
    ....................uaaggaaacuagcggucgu.............................................    3          0         seq
    ....................uaaggaaacuagcggucguu............................................    16         0         seq
    ....................uaaggaaacuagcggucguC............................................    3          1         seq
    ....................uaaggaaacuagcggucguuU...........................................    1          1         seq
    ....................uaaggaaacuagcggucguug...........................................    25         0         seq
    ....................uaaggaaacuagcggucguuA...........................................    2          1         seq
    ....................uaaggaaacuagcAgucguug...........................................    2          1         seq
    ....................uaaggaaacuagcggucguugg..........................................    35         0         seq
    ....................uaaggaaacuagcggucguugA..........................................    4          1         seq
    ....................uaaggaaacuagcggucguugU..........................................    2          1         seq
    ....................uaaggaaacuagcggucguuAg..........................................    1          1         seq
    ....................uaaggaaacuagcggucguGgg..........................................    1          1         seq
    ....................uaaggaaacuagcggucguugC..........................................    1          1         seq
    ....................uaaggaaacuagcggucguuAgc.........................................    2          1         seq
    ....................uaaggaaacuGgcggucguuggc.........................................    1          1         seq
    ....................uaaggaaacuagcggucguuggU.........................................    56         1         seq
    ....................uaaggaaacuagcggucguuggG.........................................    1          1         seq
    ....................uaaggaaacuagcggucguuggA.........................................    9          1         seq
    ....................uaaggUaacuagcggucguuggc.........................................    1          1         seq
    ....................Aaaggaaacuagcggucguuggc.........................................    1          1         seq
    ....................uaaggaaacuagcggucguuggc.........................................    138        0         seq
    ....................uaaggGaacuagcggucguuggc.........................................    4          1         seq
    ....................uGaggaaacuagcggucguuggc.........................................    2          1         seq
    ....................uaaggaaaUuagcggucguuggc.........................................    1          1         seq
    ....................Gaaggaaacuagcggucguuggc.........................................    1          1         seq
    ....................uaaggaaacuagcgAucguuggc.........................................    1          1         seq
    ....................uaGggaaacuagcggucguuggc.........................................    1          1         seq
    ....................Caaggaaacuagcggucguuggc.........................................    2          1         seq
    ....................uaaggaaacuagcggucguuggcg........................................    6          0         seq
    ....................uaaggaaacuagcggucguuggcU........................................    109        1         seq
    ....................uaaggaaacuagcggucguuggcA........................................    20         1         seq
    ....................uaaggaaacuagcggucguuggcC........................................    14         1         seq
    ....................uGaggaaacuagcggucguuggcg........................................    1          1         seq
    .....................aaggaaacuagcggucguuggc.........................................    1          0         seq
```

novel-nve-miR-69_guide

novel-nve-miR-69_star

augguggugaaauuugcagaguaaggaaacuagcggucguuggcguucgccaacaaccgcuuguuccuuuucucugcaauuguga

| | | |
|---|---|---|
| Provisional ID | | novel-nve-miR-71 |
| Score total | : | 5.7 |
| Score for star read(s) | : | -1.3 |
| Score for read counts | : | 0 |
| Score for mfe | : | 2.4 |
| Score for randfold | : | 1.6 |
| Score for cons. seed | : | 3 |
| Total read count | : | 1082 |
| Mature read count | : | 1080 |
| Loop read count | : | 0 |
| Star read count | : | 2 |

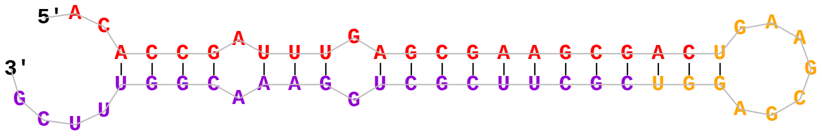


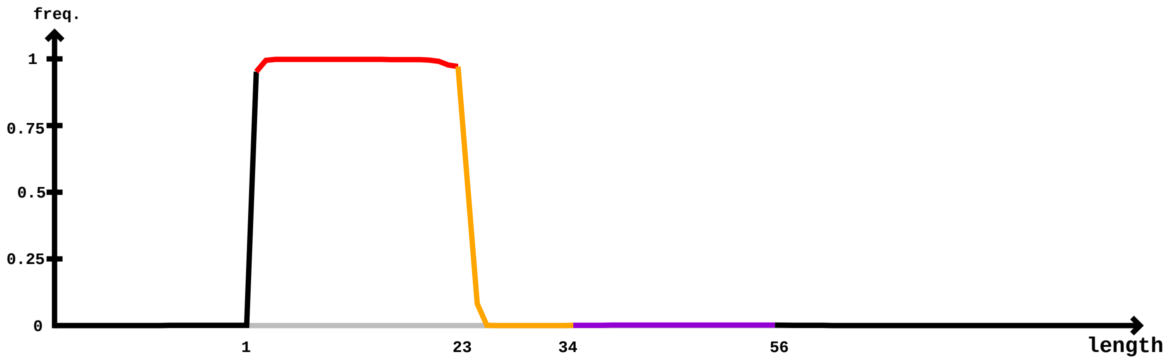


**Mature** **Star**

```
5'- ggcuccgagcuucgcggcgacaccgauuugagcgaagcgacugaagcgaggucgcuucgcuggaaacgguuucgccucggagccucauuaagaaccuucgauuuggucacu  -3'   obs
    ggcuccgagcuucgcggcgacaccgauuugagcgaagcgacugaagcgaggucgcuucgcuggaaacgguuucgccucggagccucauuaagaaccuucgauuuggucacu        exp
    (((((((.......(((((.((((.(((.(((((((((((.......))))))))))).))).)))).))))))))))))).......((.((........))))...   reads   mm   sample
    ..........uucgcggcgacaGcgauuugagc..............................................................................   1       1    seq
    ...................acaccgauuugagcgaag..........................................................................   2       0    seq
    ...................acaccgauuugagcgaagc.........................................................................   3       0    seq
    ...................acaccgauuugagcgaagU.........................................................................   1       1    seq
    ...................Gcaccgauuugagcgaagcg........................................................................   1       1    seq
    ...................acaccgauuugagcgaagcA........................................................................   1       1    seq
    ...................acaccgauuugaAcgaagcg........................................................................   1       1    seq
    ...................acaccgauuugagcgaagcg........................................................................   12      0    seq
    ...................acaccgauuugagcgaagcga.......................................................................   2       0    seq
    ...................Ccaccgauuugagcgaagcgac......................................................................   1       1    seq
    ...................acaccgauuugaCcgaagcgac......................................................................   1       1    seq
    ...................Ucaccgauuugagcgaagcgac......................................................................   1       1    seq
    ...................acaccgauuugagcgGagcgac......................................................................   1       1    seq
    ...................acaAcgauuugagcgaagcgac......................................................................   2       1    seq
    ...................acGccgauuugagcgaagcgac......................................................................   4       1    seq
    ...................aAaccgauuugagcgaagcgac......................................................................   1       1    seq
    ...................acaccgaAuugagcgaagcgac......................................................................   1       1    seq
    ...................acaccgUuuugagcgaagcgac......................................................................   1       1    seq
    ...................acaccgauuugagcgaagcgaA......................................................................   1       1    seq
    ...................acaccgGuuugagcgaagcgac......................................................................   1       1    seq
    ...................Gcaccgauuugagcgaagcgac......................................................................   8       1    seq
    ...................aUaccgauuugagcgaagcgac......................................................................   1       1    seq
    ...................acaccgaCuugagcgaagcgac......................................................................   2       1    seq
    ...................acaccgauuugagcgaagcgac......................................................................   507     0    seq
    ...................acaccgauuugagcgaagcgGc......................................................................   2       1    seq
    ...................acaccAauuugagcgaagcgac......................................................................   5       1    seq
    ...................acaccgauAugagcgaagcgac......................................................................   1       1    seq
    ...................acaccgauuugGgcgaagcgac......................................................................   2       1    seq
    ...................acaccgauuugagcgaagUgac......................................................................   9       1    seq
    ...................acaccgauCugagcgaagcgac......................................................................   2       1    seq
    ...................acaccgauuugagUgaagcgac......................................................................   1       1    seq
    ...................acaccgauuCgagcgaagcgac......................................................................   1       1    seq
    ...................acaccgauuugagcgaagcgaU......................................................................   28      1    seq
```

```
                   Mature                          Star
ggcuccgagcuucgcggcgacaccgauuugagcgaagcgacugaagcgaggucgcuucgcuggaaacgguuucgccucggagccucauuaagaaccuucgauuuggucacu
...................acaccAauuugagcgaagcgacu.....................................................................	1	1	seq
...................acaccgauuugagcgaagcgacu.....................................................................	250	0	seq
...................acaccgauuugGgcgaagcgacu.....................................................................	1	1	seq
...................acaccgauuugagcgaagcgaAu.....................................................................	1	1	seq
...................acaccgauuugagcgaagcgacA.....................................................................	15	1	seq
...................acaccgaAuugagcgaagcgacu.....................................................................	1	1	seq
...................acaccgauuugagcgGagcgacu.....................................................................	1	1	seq
...................acGccgauuugagcgaagcgacu.....................................................................	2	1	seq
...................acaccgauuugagcgaagcgaUu.....................................................................	1	1	seq
...................acaccgauuugagcgaagcgacC.....................................................................	41	1	seq
...................Gcaccgauuugagcgaagcgacu.....................................................................	7	1	seq
...................acacAgauuugagcgaagcgacu.....................................................................	1	1	seq
...................acaccgaCuugagcgaagcgacu.....................................................................	1	1	seq
...................acaccgauCugagcgaagcgacu.....................................................................	1	1	seq
...................Ccaccgauuugagcgaagcgacu.....................................................................	1	1	seq
...................acaccgauuugagUgaagcgacu.....................................................................	4	1	seq
...................acaccgauuCgagcgaagcgacu.....................................................................	1	1	seq
...................acaccgauuugagcgaagcgacG.....................................................................	1	1	seq
...................acaccgauuugagcgaagcgGcu.....................................................................	4	1	seq
...................acaccgauuugagcgaagcgacuA....................................................................	2	1	seq
...................acaccgauuugagcgaagcgacuU....................................................................	76	1	seq
...................acaccgauuugagcgaagcgacug....................................................................	2	0	seq
...................acaccgauuugagcgaagcgacuC....................................................................	7	1	seq
...................acaccgauuugagcgaagcgacuUa...................................................................	1	1	seq
....................caccgauuugagcgaagc.........................................................................	1	0	seq
....................caccgauuugagcgaagcga.......................................................................	4	0	seq
....................caccgauuugagcgaagcgac......................................................................	16	0	seq
....................caccgauuugagcgaagcgaU......................................................................	1	1	seq
....................caccgauuugagcgaagcgacu.....................................................................	22	0	seq
....................caccgauuugagcgaagcgacC.....................................................................	2	1	seq
.....................accgauuugagcgaagcgacu.....................................................................	1	0	seq
.....................accgauuugagcgaGgcgacu.....................................................................	1	1	seq
.....................accgauuugagcgaagcgacG.....................................................................	1	1	seq
.....................accgauuugagcgaagcgacuA....................................................................	1	1	seq
....................................................cgcuucgcuggaaacgguuucg.....................................	1	0	seq
........................................................ucgcuggaaacgguuucgccucC................................	1	1	seq
```

miRBase precursor : novel-nve-miR-72
Total read count : 69
novel-nve-miR-72_guide read count : 62
novel-nve-miR-72_star read count : 7
remaining reads : 0

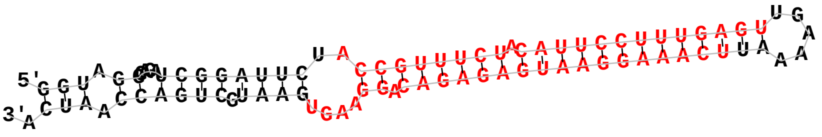

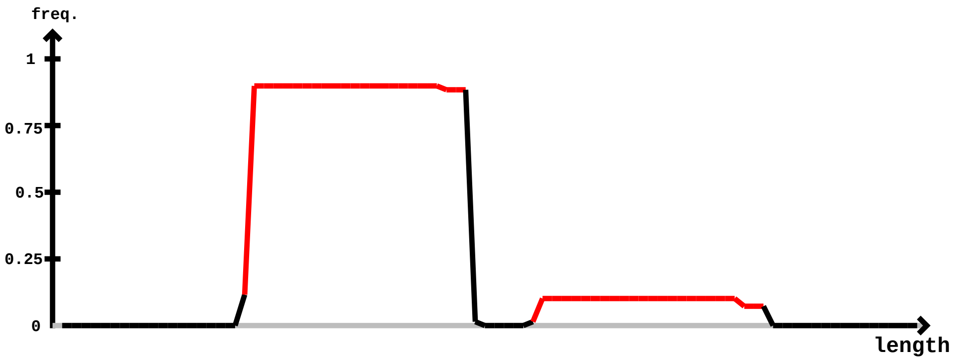


novel-nve-miR-72_guide

novel-nve-miR-72_star

```
5'- gguagggagaucggauucuaccguuucuacauuccuuugaguugaaaauucaaaggaaugagagacaggaagugaaugcugaccaauca   -3'    exp
    (((.((....((((((((..((((((((.(((((((((((((.....))))))))))))))))))).))....)))).)))))).))).      reads      mm        sample
    ..................uaccguuucuacauuccuuug..................................................      1          0         seq
    ..................uaccguuucuacauuccuuugagu...............................................      5          0         seq
    ..................uaccguuucuacauuccuuugagC...............................................      2          1         seq
    ...................Gccguuucuacauuccuuugagu...............................................      1          1         seq
    ...................accguuucuacauuccuuugagA...............................................      1          1         seq
    ...................accguuucuacauuccuuugagu...............................................      32         0         seq
    ...................accguuucuacauuccCuugagu...............................................      3          1         seq
    ...................accguuucuaAauuccuuugagu...............................................      1          1         seq
    ...................aUcguuucuacauuccuuugagu...............................................      1          1         seq
    ...................accguuucuacauuccuuugagG...............................................      1          1         seq
    ...................accguuucuacauuccuuugagC...............................................      12         1         seq
    ...................accguuucuacauuccuuugGgu...............................................      1          1         seq
    ...................accguuucuacauuccuuugaguG..............................................      1          1         seq
    ................................................uucaaaggaaugagagacaggU...................      1          1         seq
    .................................................ucaaaggaaugagagacaggU...................      1          1         seq
    .................................................ucaaaggaaugagagacaggaagG................      1          1         seq
    .................................................ucaaaggaaugagagacaggaaUu................      1          1         seq
    .................................................ucaaaggaaugagagacaggaagu................      3          0         seq
```

miRBase precursor : novel-nve-miR-73
Total read count : 270
novel-nve-miR-73_guide read count : 259
novel-nve-miR-73_star read count : 6
remaining reads : 5

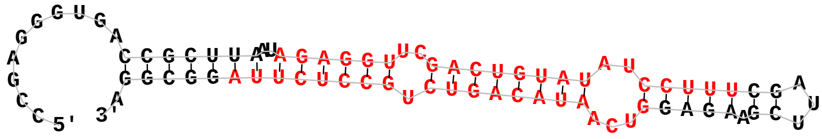

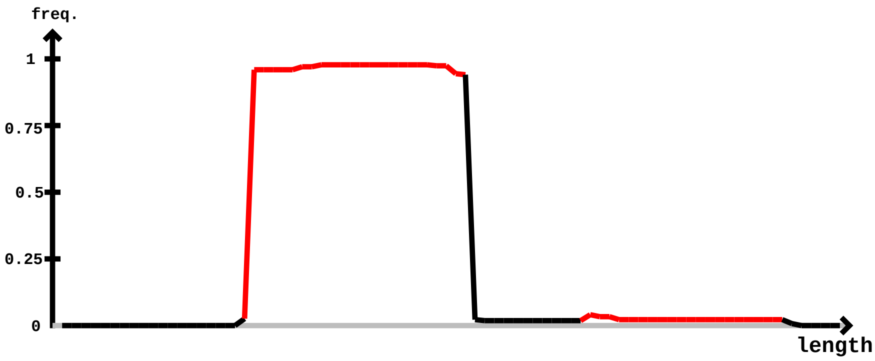


```
                                                       novel-nve-miR-73_star
                          novel-nve-miR-73_guide
5'- ccgagggugaccgcuuaauagagguucgacuguauauccuuucgauucgaagaggucaauacagucugccucuuaggcgga  -3'   exp
    ..........(((((((..((((((..((((((((..(((((((...)).)))))...)))))))).))))))))))))).   reads   mm   sample
    ..................uagagguucgacuguauauccu.........................................   1       0    seq
    ..................uagagguucgacuguauauccC.........................................   1       1    seq
    ..................uagagguucgacuguauauccuuu.......................................   5       0    seq
    ...................agagguucgacuguauauc...........................................   1       0    seq
    ...................agagguucgacuguauauccA.........................................   1       1    seq
    ...................agagguucgacuguauauccu.........................................   5       0    seq
    ...................agagguucgacuguauauccuu........................................   1       0    seq
    ...................agagguucgGcuguauauccuuu.......................................   2       1    seq
    ...................agagguucgacAguauauccuuu.......................................   2       1    seq
    ...................agagguucgacuguauaucUuuu.......................................   2       1    seq
    ...................agagguucgacuguauauccuuA.......................................   6       1    seq
    ...................agagguCcgacuguauauccuuu.......................................   1       1    seq
    ...................agagguucgacuguauauccuuu.......................................   195     0    seq
    ...................agagAuucgacuguauauccuuu.......................................   1       1    seq
    ...................agagguuUgacuguauauccuuu.......................................   3       1    seq
    ...................agagguucgacuguauaCccuuu.......................................   1       1    seq
    ...................agagguucgacuguauauccuuG.......................................   1       1    seq
    ...................Ggagguucgacuguauauccuuu.......................................   4       1    seq
    ...................agagguucgacuguauauccuCu.......................................   1       1    seq
    ...................agGgguucgacuguauauccuuu.......................................   1       1    seq
    ...................agagguucUacuguauauccuuu.......................................   1       1    seq
    ...................agaggCucgacuguauauccuuu.......................................   1       1    seq
    ...................agagguucgacuguauauccuuC.......................................   20      1    seq
    ...................agaggGucgacuguauauccuuu.......................................   1       1    seq
    ...................agagguucgacuguauauccuuuU......................................   1       1    seq
    ........................uucgacuguauauccuuucgauucgaagagguc........................   3       0    seq
    ..........................cgacuguauauccuuucgauucgaagagU..........................   1       1    seq
    ..........................cgacuguauauccuuCcgauucgaagagg..........................   1       1    seq
    ......................................................gucaauacagucugccucuuU......   3       1    seq
    ......................................................gucaauacagucugccucuua......   1       0    seq
    ......................................................gucaauacagucugccucuuaU.....   2       1    seq
```

miRBase precursor : novel-nve-miR-75
Total read count : 246
novel-nve-miR-75_guide read count : 245
novel-nve-miR-75_star read count : 1
remaining reads : 0

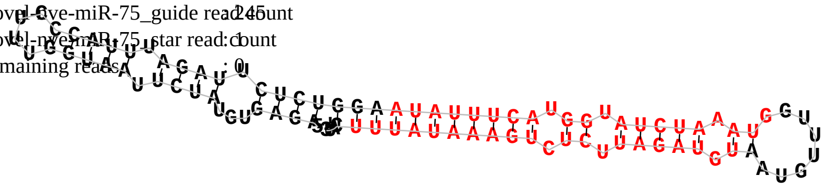

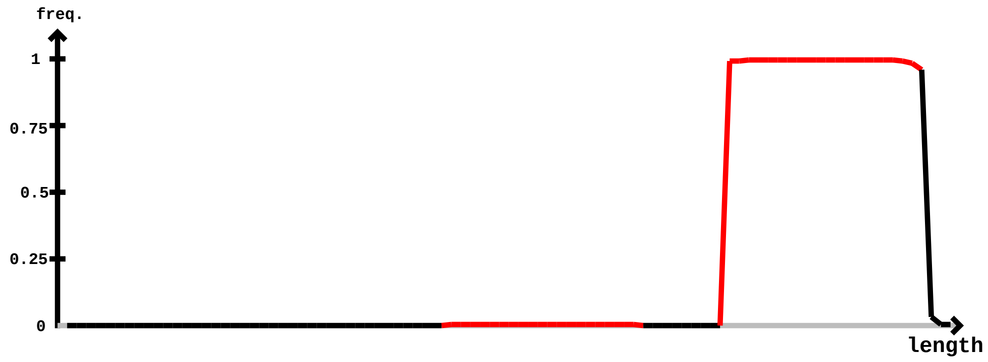


```
                                          novel-nve-miR-75_star
                                                                                novel-nve-miR-75_guide
5'- agaguguaucuuaaugguuucccauuuagauucucuggaauauuucaugguaucuaaaugguuuguaauguagauucucugaaauauuucau  -3'   exp
    ((((...((((.(((((....))))).)))).))))(((((((((((.((.(((((.((........)).))))).)).)))))))))))..   reads   mm   sample
    .......................................auauuucaugguaucuaaau.................................   1       0    seq
    ....................................................................uguagauucucugaaaua......   1       0    seq
    ....................................................................uguagauucucugaaauaG.....   1       1    seq
    ....................................................................uguagauucucugaaauau.....   1       0    seq
    ....................................................................uguagauucucugaaauauu....   5       0    seq
    ....................................................................uguagauucucugaaauauC....   1       1    seq
    ....................................................................Aguagauucucugaaauauuu...   1       1    seq
    ....................................................................uguagauucucugaaauauuu...   188     0    seq
    ....................................................................ugCagauucucugaaauauuu...   1       1    seq
    ....................................................................uguagauuAucugaaauauuu...   1       1    seq
    ....................................................................uguagGuucucugaaauauuu...   1       1    seq
    ....................................................................uguagauucucugaaauauCu...   2       1    seq
    ....................................................................uguagauucuUugaaauauuu...   1       1    seq
    ....................................................................uguagauucAcugaaauauuu...   1       1    seq
    ....................................................................uguagauucucugaaGuauuu...   3       1    seq
    ....................................................................uguagauucucugaUauauuu...   1       1    seq
    ....................................................................uguagauCcucugaaauauuu...   1       1    seq
    ....................................................................uguagauucucugaaauauuA...   2       1    seq
    ....................................................................uguagauucucugaaauauuG...   1       1    seq
    ....................................................................uguagauucucugaaauauuC...   24      1    seq
    ....................................................................uguagauucucugaaauauuuA..   4       1    seq
    ....................................................................uguagauucucugaaauauuuU..   2       1    seq
    ....................................................................uguagauucucugaaauauuuc..   1       0    seq
    ......................................................................uagauucucugaaauauuucau   1       0    seq
```

miRBase precursor : novel-nve-miR-76
Total read count : 69
novel-nve-miR-76_guide read count : 68
novel-nve-miR-76_star read count : 1
remaining reads : 0

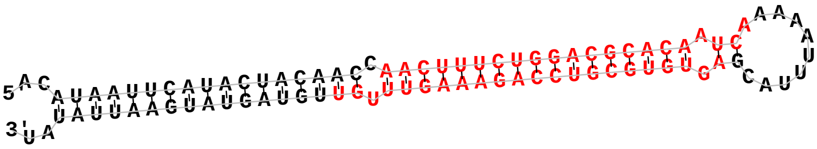

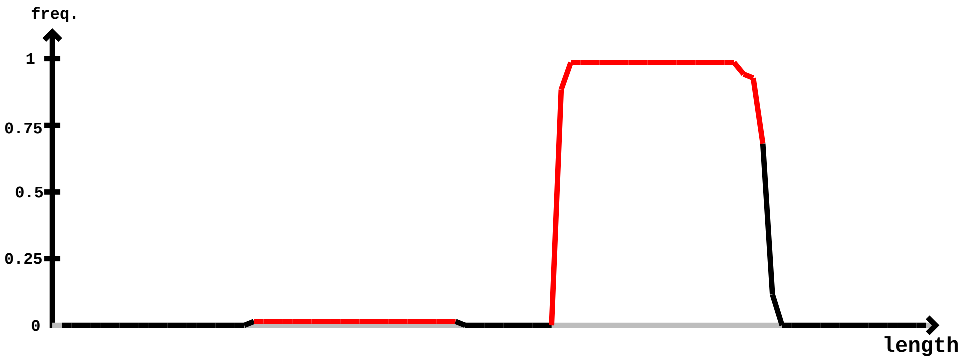


```
                                                    novel-nve-miR-76_guide
                      novel-nve-miR-76_star
5'- acauaauucauacuacaaccaacuuucuggacgcacaaucaaaaauuuacgagugugcguccagaaaguuuguuguaguaugaauuauau   -3'   exp
    ..(((((((((((((((((.(((((((((((((((((.((..........)).))))))))))))))))).)))))))))))))))))..   reads      mm        sample
    ..................caacuuucuggacgcacaauca..................................................   1          0         seq
    ..................................................aUugugcguccagaaaguuug...................   1          1         seq
    ..................................................agugugcguccagaaaguuug...................   15         0         seq
    ..................................................agugugcAuccagaaaguuug...................   1          1         seq
    ..................................................agugugcguccagaaGguuugu..................   1          1         seq
    ..................................................agugugcguccagaaaguuugG..................   1          1         seq
    ..................................................agugugcguccagaaaguuugC..................   4          1         seq
    ..................................................agugugcguccUgaaaguuugu..................   1          1         seq
    ..................................................agugugcgucUagaaaguuugu..................   1          1         seq
    ..................................................agugugcguccagaaaguuugA..................   1          1         seq
    ..................................................agugugcguccagaaaguuugu..................   29         0         seq
    ..................................................agCgugcguccagaaaguuugu..................   1          1         seq
    ..................................................agugugcguccagaaaguuuguu.................   5          0         seq
    ...................................................Uugugcguccagaaaguu.....................   3          1         seq
    ...................................................Uugugcguccagaaaguuu....................   1          1         seq
    ...................................................gugugcguccagaaaguuuguC.................   1          1         seq
    ...................................................gugugcguccagaaaguuuguA.................   2          1         seq
```

miRBase precursor : novel-nve-miR-79
Total read count : 298
novel-nve-miR-79_guide read count : 294
novel-nve-miR-79_star read count : 3
remaining reads : 1

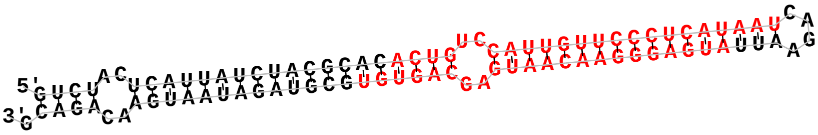


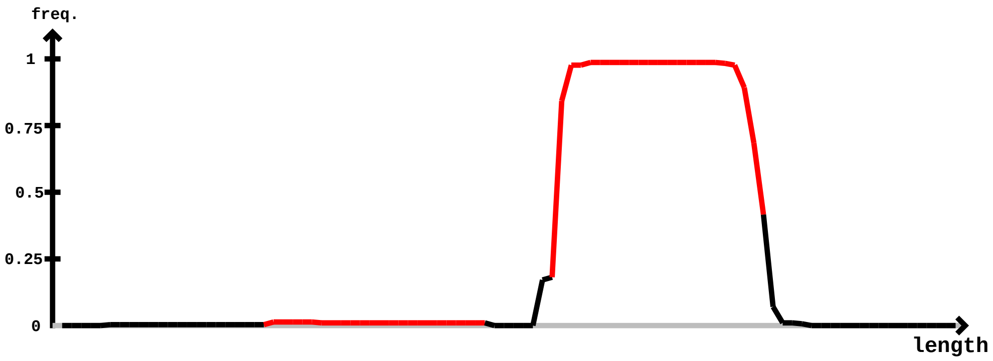


novel-nve-miR-79_star
novel-nve-miR-79_guide

```
5'- gucuacucauuaucuacgcacacuguccauuguucccucauaaucagaauuaugagggaacaaugagcagugugcguagauaaugaacagacg   -3'    exp
    ((((..((((((((((((((((((..(((((((((((((((((....)))))))))))))))))..))))))))))))))))))..)))).   reads     mm        sample
    ....acucauuaucuacgcacacugu...................................................................   1         0         seq
    .....................acuguccauuguucccucauaau.................................................   3         0         seq
    .................................................uuaugagggaacaaugagc.........................   1         0         seq
    .................................................uuaugagggaacaaugagcC........................   1         1         seq
    .................................................uuaugagggaacaaugagcGg.......................   1         1         seq
    .................................................uuaugagggaacaaugagcag.......................   4         0         seq
    .................................................uuaugagggaacaaugagcagu......................   33        0         seq
    .................................................uuaugagggaacaaugagcagC......................   2         1         seq
    .................................................uCaugagggaacaaugagcagu......................   1         1         seq
    .................................................uAaugagggaacaaugagcagu......................   1         1         seq
    .................................................uuaugGgggaacaaugagcagu......................   1         1         seq
    .................................................uuaugagggaaAaaugagcagu......................   1         1         seq
    .................................................uuaugagggaacaaugagcagA......................   2         1         seq
    .................................................uuaugagggaacaaugagcagug.....................   1         0         seq
    .................................................uuaugagggaacaaugagcaguU.....................   2         1         seq
    ..................................................Aaugagggaacaaugagcagu......................   1         1         seq
    ..................................................uaugagggaacaaugagcagug.....................   2         0         seq
    ...................................................augaggUaacaaugagca........................   1         1         seq
    ...................................................augaCggaacaaugagcag.......................   1         1         seq
    ...................................................augagggaacaaugagcag.......................   13        0         seq
    ...................................................augagggaacaaugagcaA.......................   1         1         seq
    ...................................................augagggaacaaugagcGgu......................   1         1         seq
    ...................................................augagggaacaaugagcagC......................   2         1         seq
    ...................................................augagggaaGaaugagcagu......................   1         1         seq
    ...................................................augagggaacaaugagcagu......................   12        0         seq
    ...................................................augagggaacaaugagUagu......................   1         1         seq
    ...................................................augagggaacaaugagUagug.....................   2         1         seq
    ...................................................augagggaaUaaugagcagug.....................   1         1         seq
    ...................................................augagggaacaaugagcaguA.....................   5         1         seq
    ...................................................augagggaacaaCgagcagug.....................   1         1         seq
    ...................................................augagggGacaaugagcagug.....................   1         1         seq
    ...................................................augagggaacaaugGgcagug.....................   1         1         seq
    ...................................................augagggaacaaugagcagug.....................   48        0         seq
    ...................................................augagUgaacaaugagcagug.....................   1         1         seq
```

```
                     novel-nve-miR-79_star
                                                   novel-nve-miR-79_guide
gucuacucauuaucuacgcacacuguccauuguucccucauaaucagaauuaugagggaacaaugagcagugugcguagauaaugaacagacg

...................................................augagggaacaaugagcaguU.....................    2    1    seq
...................................................auCagggaacaaugagcagug.....................    1    1    seq
...................................................augagggaacaaugagcagGg.....................    1    1    seq
...................................................augagggaacaaugagcUgugu....................    1    1    seq
...................................................augagggaacaaugagcagugu....................    71   0    seq
...................................................augagggaacaaAgagcagugu....................    1    1    seq
...................................................auAagggaacaaugagcagugu....................    1    1    seq
...................................................augagggaacaaugagcagugC....................    9    1    seq
...................................................augagggaacGaugagcagugu....................    1    1    seq
...................................................augagggaacaGugagcagugu....................    1    1    seq
...................................................augagggaacaaugagcagGgu....................    1    1    seq
...................................................augagggaacaaugagcagugA....................    1    1    seq
...................................................augagggaacaaugagcagugG....................    1    1    seq
...................................................augagggaacaaugagcagugug...................    1    0    seq
...................................................augagggaacaaugagcaguguC...................    2    1    seq
...................................................augGgggaacaaugagcagugug...................    1    1    seq
...................................................augagggaacaaugagcaguguU...................    8    1    seq
....................................................ugagggaacaaugagcag.......................    5    0    seq
....................................................ugagggaacaaugagcagu......................    3    0    seq
....................................................ugagggaacaaugagcagug.....................    10   0    seq
....................................................ugagggaacaaugagcaguA.....................    1    1    seq
....................................................ugagggaacaaugagcagugu....................    15   0    seq
....................................................Agagggaacaaugagcagugug...................    1    1    seq
....................................................ugagggaacaaugagcaguguU...................    1    1    seq
....................................................ugagggaacaaugagcagugug...................    3    0    seq
....................................................ugagggaacaaugagcaguguC...................    1    1    seq
......................................................agggaacaaugagcagugugcA.................    1    1    seq
......................................................agggaacaaugagcagugugcgC................    1    1    seq
......................................................agggaacaaugagcagugugcgu................    1    0    seq
```

miRBase precursor : novel-nve-miR-83
Total read count : 312
novel-nve-miR-83_guide read count : 213
novel-nve-miR-83_star read count : 98
remaining reads : 1

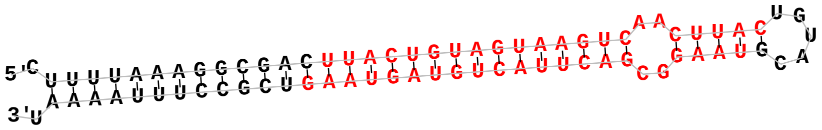

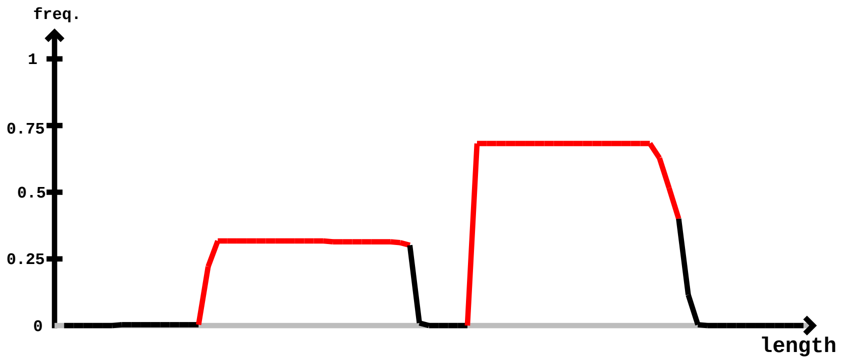


```
                    novel-nve-miR-83_star
                                        novel-nve-miR-83_guide
5'- cuuuuaaaggcgacuuacuguaguaagucaacuuacuguacguaaggcgacuuacuguaguaagucgccuuuaaaau  -3'   exp
    .((((((((((((((((((((((((((((..(((((.....)))))..)))))))))))))))))))))))))))))).   reads   mm   sample
    .....aaaggcgacuuacuguaguaag..................................................   1       0    seq
    ..............uuacuguaguaagucaacuu...........................................   1       0    seq
    ..............uuacuguaguaagucaacuua..........................................   1       0    seq
    ..............uuacuguagAaagucaacuuac.........................................   1       1    seq
    ..............uuacuguaguaagucaacuuaA.........................................   1       1    seq
    ..............uuacuguaguaagucaacuuaU.........................................   3       1    seq
    ..............uuacuguaguGagucaacuuac.........................................   1       1    seq
    ..............uuacuguaguaagucaacuuac.........................................   59      0    seq
    ..............uAacuguaguaagucaacuuacu........................................   1       1    seq
    ...............uacuguaguaagucaacuua..........................................   1       0    seq
    ...............Cacuguaguaagucaacuua..........................................   1       1    seq
    ...............uacuguaguaagucaacuuaU.........................................   2       1    seq
    ...............Aacuguaguaagucaacuuac.........................................   1       1    seq
    ...............uacuguaguaagucaCcuuac.........................................   1       1    seq
    ...............uacuguaguaagucaacuuac.........................................   22      0    seq
    ...............uacuguaguaagucaacuuacA........................................   1       1    seq
    ...............uacuguaguaagucaacuuacu........................................   1       0    seq
    ..........................................uaaggcgacuuacuguagC................   3       1    seq
    ..........................................uaaggcgacuuacuguagu................   12      0    seq
    ..........................................uaaggcgaUuuacuguagu................   1       1    seq
    ..........................................uaaggcgacuuacuguagG................   1       1    seq
    ..........................................uaaggcgacuuaAuguagua...............   1       1    seq
    ..........................................uaaggcgacuuacuguGgua...............   1       1    seq
    ..........................................uaaggcgGcuuacuguagua...............   1       1    seq
    ..........................................uaaggcgacuuacuguaguG...............   2       1    seq
    ..........................................uaaggcgacuuacuguagCa...............   1       1    seq
    ..........................................uaaggcgacuuacuguagua...............   29      0    seq
    ..........................................uaaUgcgacuuacuguaguaa..............   1       1    seq
    ..........................................uaaggcgacCuacuguaguaa..............   1       1    seq
    ..........................................uaaggcgacuuacuguaguaa..............   33      0    seq
    ..........................................uaaggcgacuuacuguaguaG..............   1       1    seq
    ..........................................uaaggcgacuuacuguaguaag.............   33      0    seq
    ..........................................uaaggAgacuuacuguaguaag.............   1       1    seq
    ..........................................uaaggcgacuuacuguaguaaA.............   6       1    seq
```

```
              novel-nve-miR-83_star
                                          novel-nve-miR-83_guide
cuuuuaaaggcgacuuacuguaguaagucaacuuacuguacguaaggcgacuuacuguaguaagucgccuuuaaaau

..........................................uaaggUgacuuacuguaguaag.............    1     1     seq
..........................................uaaggcgacuuacuguaguaaU.............    35    1     seq
..........................................uaaggcgacuuacCguaguaag.............    1     1     seq
..........................................uaaggcgacuuacuguaguaaC.............    12    1     seq
..........................................uaaggcgacuuacuguaguaagA............    3     1     seq
..........................................uaaggcgacuuacuguaguaaUu............    14    1     seq
..........................................uaaggcgacuuacuguaguaaAu............    1     1     seq
..........................................uaaggcgacuuacuguaguaagu............    10    0     seq
..........................................uaaggcgacuuacuguaguaaCu............    7     1     seq
..........................................uaaggcgacuuacuguaguaaguU...........    1     1     seq
```

miRBase precursor : novel-nve-miR-84
Total read count : 118
novel-nve-miR-84_guide read count : 90
novel-nve-miR-84_star read count : 24
remaining reads : 4

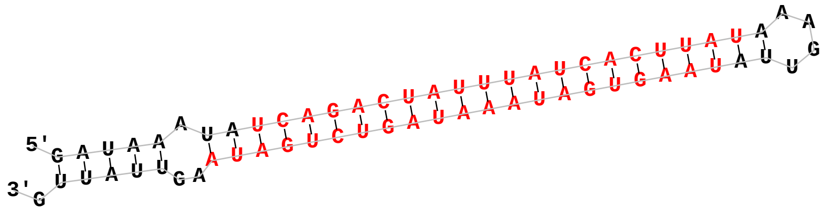


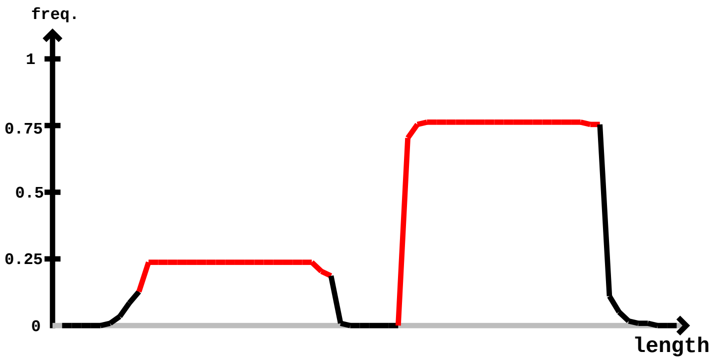


```
                                  novel-nve-miR-84_guide
          novel-nve-miR-84_star
5'- gauaaauaucagacuauuuaucacuuauaaaguuauaagugauaaaauagucugauaaguuauug  -3'   exp
    (((((.((((((((((((((((((((((....))))))))))))))))))))))..))))).   reads     mm     sample
    ....aauaucagacuauuuaucacuu.......................................   1       0      seq
    .....auaucagacuauuuaAcacuuau.....................................   1       1      seq
    .....Uuaucagacuauuuaucacuuau.....................................   1       1      seq
    .....auaucagacuauuuaucacuuauU....................................   1       1      seq
    ......uaucagacuauuuaucacuuau.....................................   6       0      seq
    .......auAagacuauuuaucacuu.......................................   1       1      seq
    .......aucagacuauuuaucacuu.......................................   1       0      seq
    .......aucagacuauuuaucacuuU......................................   1       1      seq
    .......aucagacuauuuaucacuuau.....................................   2       0      seq
    ........ucagacuauuuaucacuu.......................................   1       0      seq
    ........ucagacuauuuaucacuuU......................................   1       1      seq
    ........ucagacuauuuaucacuuaA.....................................   1       1      seq
    ........ucagacuauuuaucacuuaC.....................................   1       1      seq
    ........ucagacuauuuaucacuuau.....................................   9       0      seq
    ...................................uaagugauaaaauagucugC..........   1       1      seq
    ...................................uaagugauaaaauagucugauC........   4       1      seq
    ...................................uaagugauaaaauagucugauU........   51      1      seq
    ...................................uaagugauaaaauagucugaua........   14      0      seq
    ...................................uaagugauaaaauagucugauG........   1       1      seq
    ...................................uaagugauaaaauagucugauaU.......   1       1      seq
    ...................................uaagugauaaaauagucugauaa.......   4       0      seq
    ...................................uaagugauaaaauagucugauCa.......   2       1      seq
    ...................................uaagugauaaaauagucugauaaA......   1       1      seq
    ...................................uaagugauaaaauagucugauaag......   3       0      seq
    ...................................uaagugauaaaauagucugauaaguuU...   1       1      seq
    ....................................aagugauaaaauagucugauU........   2       1      seq
    ....................................aagugauaaaauagucugaua........   4       0      seq
    .....................................agugauaaaauagucugauaagu.....   1       0      seq
```

miRBase precursor : novel-nve-miR-85
Total read count : 391
novel-nve-miR-85_guide read count : 350
novel-nve-miR-85_star read count : 28
remaining reads : 13

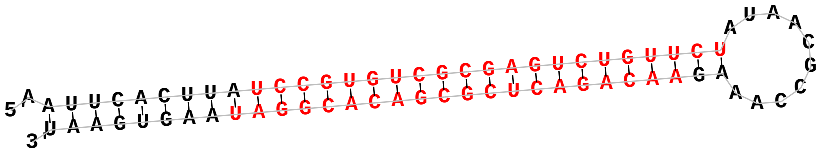

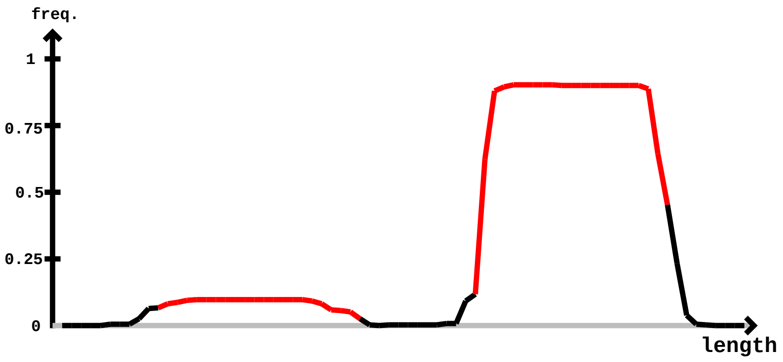


novel-nve-miR-85_star
novel-nve-miR-85_guide

```
5'- aauucacuuauccgugucgcgagucuguucuauaacgccaaagaacagacucgcgacacggauaagugaau   -3'    exp
    .((((((((((((((((((((((((((((((..........))))))))))))))))))))))))))))))    reads      mm        sample
    ....caUuuauccgugucgcgaguc..............................................    2          1         seq
    .......uuauccgugucgcgagucu.............................................    1          0         seq
    .......uuauccgugucgcgagucC.............................................    1          1         seq
    .......uuauccgugucgcgagucug............................................    3          0         seq
    .......uuauccgugucgUgagucug............................................    1          1         seq
    .......uuauccgugucgcgaCucug............................................    1          1         seq
    .......uuauccgugucgcgagucuguucu........................................    1          0         seq
    ........uauccgugucgcgagucu.............................................    2          0         seq
    ........uUuccgugucgcgagucug............................................    2          1         seq
    ........uauccguguAgcgagucug............................................    1          1         seq
    ........uauccgugucgcgagucuA............................................    1          1         seq
    ........uauccguguAgcgagucugu...........................................    1          1         seq
    ........uauccguguAgcgagucuguuc.........................................    6          1         seq
    ........uauccgugucgcgagucuguuc.........................................    1          0         seq
    ........uauccguguAgcgagucuguucu........................................    1          1         seq
    .........auccgugucgcgagucuguuA.........................................    1          1         seq
    ..........uccgugucgcgagucuguu..........................................    2          0         seq
    ..........uccgugucgcgagucuguCcu........................................    1          1         seq
    ..........Accgugucgcgagucuguucu........................................    1          1         seq
    ..........uccgugucgcgagucuguucu........................................    2          0         seq
    ...........ccgugucgcgagucuguuc.........................................    2          0         seq
    ............cgugucgcgagucuguucu........................................    1          0         seq
    ............cgugucgcAagucuguucu........................................    2          1         seq
    .............gugucgcgagucuguucuU.......................................    1          1         seq
    .................................aacgcUaaagaacagacu....................    1          1         seq
    .......................................aaagaacagacucgcgacacU...........    1          1         seq
    .......................................aaagaacagacucgcgacacC...........    1          1         seq
    .........................................agaacagacuUgcgacacg...........    1          1         seq
    .........................................agaacagacucgcgacacA...........    1          1         seq
    .........................................agaacagacucgcgacacg...........    1          0         seq
    .........................................agaacagacucgcgacacgg..........    15         0         seq
    .........................................agaacagacucgcgacacgU..........    1          1         seq
    .........................................agaacagacucgcgacacgA..........    4          1         seq
    .........................................agaacGgacucgcgacacgg..........    1          1         seq
```

novel-nve-miR-85_star

novel-nve-miR-85_guide

aauucacuuauccgugucgcgagucuguucuauaacgccaaagaacagacucgcgacacggauaagugaau

.........................................agaacagacucgAgacacgg.......... 1 1 seq
.........................................agaacagacucgcgacacggC......... 1 1 seq
.........................................agaacagacucgcgacacggU......... 2 1 seq
.........................................Ugaacagacucgcgacacgga......... 1 1 seq
.........................................agaacagacucgcgacacggG......... 1 1 seq
.........................................agaacagacucgcgacacgga......... 3 0 seq
..........................................gaacagacuUgcgacacgg.......... 1 1 seq
..........................................Aaacagacucgcgacacgg.......... 1 1 seq
..........................................gaacagacucgcgacacgg.......... 1 0 seq
..........................................gaacagacucgcgacacgC.......... 1 1 seq
..........................................gaacagacucgcgacacgga......... 2 0 seq
..........................................gaacagacucgcgacacggU......... 2 1 seq
..........................................gaacagacucgcgacacggau........ 1 0 seq
..........................................gaacagacucgcgacacggUu........ 1 1 seq
...........................................Gacagacucgcgacacgg.......... 1 1 seq
...........................................aacagaUucgcgacacgg.......... 1 1 seq
...........................................aacagacucgcgacacgU.......... 6 1 seq
...........................................aacagacucgcAacacgg.......... 1 1 seq
...........................................aacagacucgcgacacgg.......... 52 0 seq
...........................................aacagacucgcgacacCg.......... 1 1 seq
...........................................aacagacucgcgacacgA.......... 4 1 seq
...........................................aacagGcucgcgacacgg.......... 1 1 seq
...........................................aacagacCcgcgacacgg.......... 1 1 seq
...........................................aacGgacucgcgacacgg.......... 1 1 seq
...........................................aacagacucgcgacacggC......... 1 1 seq
...........................................Cacagacucgcgacacgga......... 1 1 seq
...........................................aacaAacucgcgacacgga......... 1 1 seq
...........................................aacagacucgcgacacggU......... 10 1 seq
...........................................aacagacucgcgacacgga......... 31 0 seq
...........................................aacagacucgcgacacggG......... 3 1 seq
...........................................aacagacucgcgacacggaA........ 2 1 seq
...........................................Gacagacucgcgacacggau........ 1 1 seq
...........................................aacagacucgUgacacggau........ 1 1 seq
...........................................aacagGcucgcgacacggau........ 1 1 seq
...........................................aacaAacucgcgacacggau........ 1 1 seq
...........................................aacagacucgcgacacggaC........ 3 1 seq
...........................................aacagacucgcgacacggUu........ 4 1 seq
...........................................aacagacucgcgacacggau........ 35 0 seq
...........................................aacagacuUgcgacacggau........ 1 1 seq
...........................................aacagacucgcgacacggauC....... 3 1 seq
...........................................aacagacucgcgacacggaCa....... 1 1 seq
...........................................aacagacuUgcgacacggaua....... 2 1 seq
...........................................aaUagacucgcgacacggaua....... 1 1 seq
...........................................aacagacucgcgacacggauU....... 24 1 seq
...........................................aacagacucgcgacacggaua....... 2 0 seq
............................................acagacAcgcgacacgga......... 1 1 seq
............................................aUagacucgcgacacgga......... 1 1 seq
............................................acagacucgcgacacgga......... 12 0 seq
............................................acagacucgcgacacggU......... 1 1 seq
............................................acagacucgcgacaUgga......... 1 1 seq
............................................acagacucgcgacGcggau........ 1 1 seq
............................................acagacucgcgacacggaC........ 1 1 seq
............................................acagacucgcgUcacggau........ 1 1 seq
............................................acagacuAgcgacacggau........ 1 1 seq
............................................acagacuUgcgacacggau........ 3 1 seq
............................................acagacucgcgacacggaG........ 2 1 seq
............................................acagacucgcgacacggCu........ 2 1 seq
............................................acagacucgcgacacggau........ 22 0 seq
............................................acagacucgcgacacAgaua....... 1 1 seq
............................................acagaUucgcgacacggaua....... 1 1 seq
............................................acagacucgcgacacggauU....... 11 1 seq
............................................acagacucgcgacacggauG....... 1 1 seq
............................................acagacucgcgacacggaua....... 24 0 seq
............................................acagacucgcgGcacggaua....... 1 1 seq
............................................acagacucgcgacacggauaa...... 4 0 seq
............................................acagacucgcgacacggauaU...... 6 1 seq
............................................acagacucgcgacacggauaG...... 1 1 seq
............................................acagacucgcgacacggauaUg..... 1 1 seq
.............................................cagacucgcgacacggau........ 2 0 seq

```
          novel-nve-miR-85_star
                                           novel-nve-miR-85_guide
aauucacuuauccgugucgcgagucuguucuauaacgccaaagaacagacucgcgacacggauaagugaau

.............................................cagacucgcgacacggaua.......    1           0           seq
.............................................cagacucgcgacacggauU.......    1           1           seq
.............................................Uagacucgcgacacggauaa......    1           1           seq
.............................................cagacucgcgacacggauUa......    1           1           seq
..............................................agacucgcgacGcggaua.......    2           1           seq
..............................................agacucgcgacacggauaaCu....    1           1           seq
```

miRBase precursor : novel-nve-miR-86
Total read count : 154
novel-nve-miR-86_guide read count : 126
novel-nve-miR-86_star read count : 21
remaining reads : 7

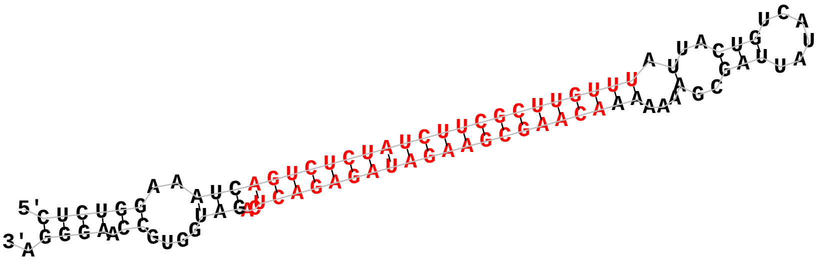

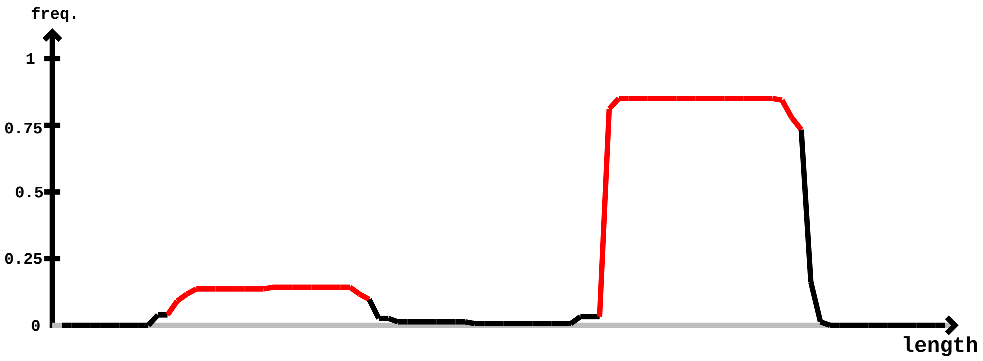


novel-nve-miR-86_guide
novel-nve-miR-86_star

| | | | |
|---|---|---|---|
| 5'- cucuggaaaucagucucuaucuucgcuuguuuauuacugucauauuagcgaaaaaaacaagcgaagauagagacucagauggugccaaggga -3' | exp | | |
| (((((..(((((((((((((((((((((((((.(..(((......)))..)...)))))))))))))))))))))..)))....)).)))). | reads | mm | sample |
| .........uUagucucuaucuucgcuugu.............................................................. | 1 | 1 | seq |
| .........ucagucucuaucuucgcuugu.............................................................. | 2 | 0 | seq |
| .........ucagucucuaucuucgcuuguu............................................................. | 3 | 0 | seq |
| ...........agucucuaucuucgcuugu.............................................................. | 1 | 0 | seq |
| ...........agucucuaucuucgcuuguuC............................................................ | 1 | 1 | seq |
| ...........agucucuaucuucgcCuguuu............................................................ | 1 | 1 | seq |
| ...........agucucuaucuucgcuuguuu............................................................ | 5 | 0 | seq |
| ............gucucuaucuucgcuuguuu............................................................ | 3 | 0 | seq |
| ............gucucuaucuucgcuuguCu............................................................ | 1 | 1 | seq |
| .............ucucuaucuucgcuuguuua........................................................... | 1 | 0 | seq |
| .............ucucuaucuucgcuuguuuau.......................................................... | 1 | 0 | seq |
| .............ucucuaucuucgcuuguuuaC.......................................................... | 1 | 1 | seq |
| .....................uucgcuuguuuauuacuguca.................................................. | 1 | 0 | seq |
| .................................uuacugucauauuagcgaaU....................................... | 1 | 1 | seq |
| .....................................................aaaacaagcgaagauagagac.................. | 1 | 0 | seq |
| .....................................................Uaaacaagcgaagauagagacu................. | 3 | 1 | seq |
| .....................................................aaaacaagcgaagauagagacu................. | 1 | 0 | seq |
| ........................................................acaagcgaagauagagacu................. | 3 | 0 | seq |
| ........................................................acaagcgaagauagagacG................. | 1 | 1 | seq |
| ........................................................acaagcgaagauagagacC................. | 1 | 1 | seq |
| ........................................................acaagcgaagGuagagacu................. | 1 | 1 | seq |
| ........................................................acaagcgaagauagagacuG................ | 1 | 1 | seq |
| ........................................................acaagcgaagauagagacuU................ | 3 | 1 | seq |
| ........................................................acaagcgaagauagagacuc................ | 2 | 0 | seq |
| ........................................................acaagcgaagauagagacGc................ | 1 | 1 | seq |
| ........................................................acaagcgaagauagagacuGa............... | 4 | 1 | seq |
| ........................................................acaagcgaagauagagacucU............... | 41 | 1 | seq |
| ........................................................acaagcgaagauagGgacuca............... | 1 | 1 | seq |
| ........................................................acaagcgaagauagagacuca............... | 28 | 0 | seq |
| ........................................................acaagcgaagauaCagacuca............... | 1 | 1 | seq |
| ........................................................acaagcgaagauagagacucC............... | 9 | 1 | seq |
| ........................................................acaagcgaagauagagacucG............... | 3 | 1 | seq |
| ........................................................acaagcgaagauagagacucaU.............. | 7 | 1 | seq |
| ........................................................acaagcgaagauagagacucaA.............. | 3 | 1 | seq |

```
                                                        novel-nve-miR-86_guide
           novel-nve-miR-86_star
cucuggaaaucagucucuaucuucgcuuguuuauuacugucauauuagcgaaaaaaacaagcgaagauagagacucagauggugccaaggga

........................................................acaagcgGagauagagacucag..............    1           1           seq
........................................................acaagcgaagauagagacucaC..............    2           1           seq
........................................................acaagcgaagauagagacucag..............    7           0           seq
.........................................................caagcgaagauagagacuca...............    1           0           seq
.........................................................caagcgaagauagagacucaU..............    1           1           seq
.........................................................caagcgaagauagagacucag..............    2           0           seq
.........................................................caagcgaagauagagacucUga.............    2           1           seq
```

| | | |
|---|---|---|
| Provisional ID | | novel-nve-miR-87 |
| Score total | : | 72.5 |
| Score for star read(s) | : | 3.9 |
| Score for read counts | : | 64.9 |
| Score for mfe | : | 2.7 |
| Score for randfold | : | 1.6 |
| Score for cons. seed | : | -0.6 |
| Total read count | : | 139 |
| Mature read count | : | 107 |
| Loop read count | : | 0 |
| Star read count | : | 32 |

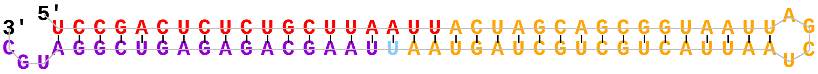


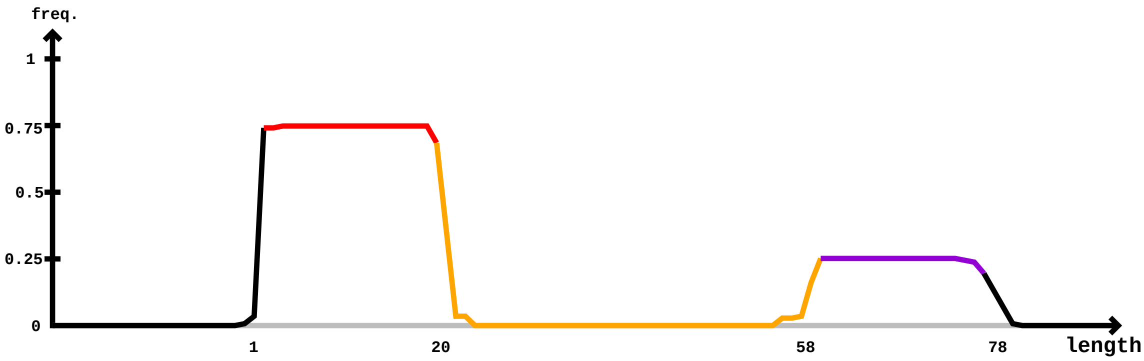


**Mature** **Star**

```
5'- cuacgcuuucuauugaaacauccgacucucugcuuaauuacuagcagcgguaauuagcuaauuacugcugcuaguaauuaagcagagagucggaugcuucaacaacaau   -3'   obs
    cuacgcuuucuauugaaacauccgacucucugcuuaauuacuagcagcgguaauuagcuaauuacugcugcuaguaauuaagcagagagucggaugcuucaacaacaau         exp
    ............(((((.(((((((((((((((((((((((((((((((((((((....))))))))))))))))))))))))))))))))))))).))))).......   reads   mm   sample
    ..................cauccgacucucugcuuaau.......................................................................   1   0   seq
    ...................auccgacucucugcuuaauu......................................................................   4   0   seq
    ....................uccgacucucugcuuaau.......................................................................   6   0   seq
    ....................uccgacucucugcuuaaC.......................................................................   2   1   seq
    ....................uccgacucucugcuCaauu......................................................................   1   1   seq
    ....................uccgacucucugcuuaauC......................................................................   11   1   seq
    ....................uccgacucucugcuuaaCu......................................................................   2   1   seq
    ....................uccgacucucugcuuaauu......................................................................   66   0   seq
    ....................uccgacucucugcuuaauua.....................................................................   3   0   seq
    ....................uccgacucucugcuuaauuU.....................................................................   6   1   seq
    ....................Accgacucucugcuuaauuacu...................................................................   1   1   seq
    ....................uccgacucucugcuuaauuacu...................................................................   2   0   seq
    ....................uccgacucucugcuuaauuacC...................................................................   1   1   seq
    ......................cgacucucugcuuaauuacu...................................................................   1   0   seq
    ..........................................................................uaauuaagcagagagucgg................   1   0   seq
    ..........................................................................uaauuaagcagagagucggU...............   1   1   seq
    ..........................................................................uaauuaagcagagagucggau..............   1   0   seq
    ..........................................................................uaauuaagcagagagucggauA.............   1   1   seq
    ............................................................................auuaagcagagagucggau..............   1   0   seq
    .............................................................................uuaagcagagagucggau..............   3   0   seq
    .............................................................................uuaagcagagagucggaA..............   1   1   seq
    .............................................................................uuaagcagagagucggauA.............   1   1   seq
    .............................................................................uuaagcagagagucggauU.............   5   1   seq
    .............................................................................uuaagcagagagucggaug.............   4   0   seq
    .............................................................................uuaagcagagagucggaugA............   1   1   seq
    .............................................................................uuaagcagagagucggauAc............   1   1   seq
    .............................................................................uuaagcagagagucggaugc............   1   0   seq
    .............................................................................uuaagcagagagucggaugcC...........   1   1   seq
    ..............................................................................uaagcagagagucAgaug.............   1   1   seq
    ..............................................................................uaagcagagagucggaugA............   3   1   seq
    ..............................................................................uaagcagagUgucggaugc............   1   1   seq
    ..............................................................................uaagcagagagucggaugc............   5   0   seq
    ..............................................................................uaagcagagagucggauUc............   1   1   seq
```

Mature Star

```
cuacgcuuucuauugaaacauccgacucucugcuuaauuacuagcagcgguaauuagcuaauuacugcugcuaguaauuaagcagagagucggaugcuucaacaacaau
............................................................................uaagcagaUagucggaugc...........    1    1    seq
............................................................................uaagcagagagucggaugcGu.........    1    1    seq
```

miRBase precursor : novel-nve-miR-88
Total read count : 180
novel-nve-miR-88_guide read count : 162
novel-nve-miR-88_star read count : 18
remaining reads : 0

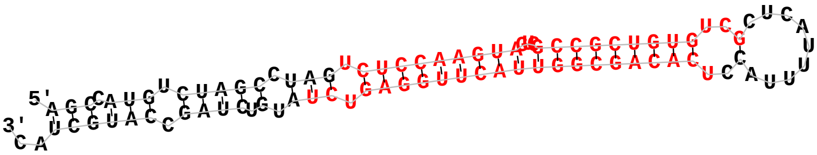

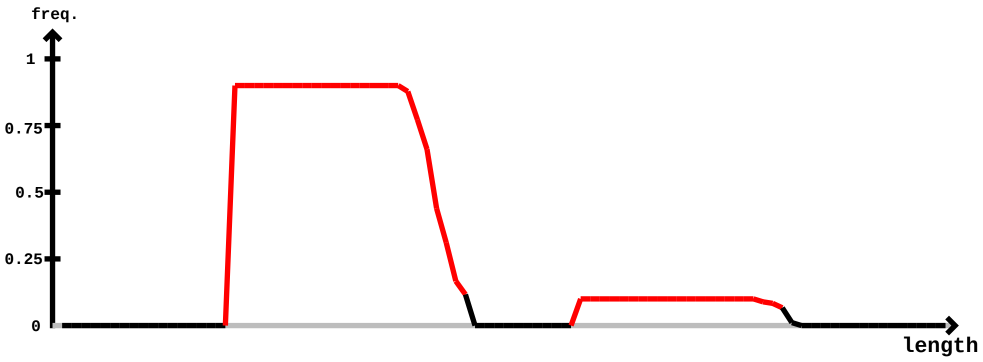


```
                                                                        novel-nve-miR-88_star
                      novel-nve-miR-88_guide
5'- agccaugucuagccuagucuccaaguagucgccgcugugucgcucauuuuaccucacagcgguuacuuggagucuaugucuagccaugcuac   -3'    exp
    (((.(((.(((((.(((.(((((((((...((((((((..(.........)..))))))))))))))))).))).).)))).))))))..    reads     mm       sample
    .................ucuccaaguagucgccgU.........................................................    2         1        seq
    .................ucuccaaguagucgccgc.........................................................    1         0        seq
    .................ucuccaaguagucgcUgc.........................................................    1         1        seq
    .................ucuccaaguagucgccgcC........................................................    2         1        seq
    .................ucuccaaguagucgccgcG........................................................    2         1        seq
    .................ucuccaaguagucgccgcA........................................................    3         1        seq
    .................ucuccaaguagucgccgcu........................................................    12        0        seq
    .................ucuccaaguagucgccgcug.......................................................    17        0        seq
    .................Acuccaaguagucgccgcug.......................................................    1         1        seq
    .................ucuccaaguagucgccgcuA.......................................................    2         1        seq
    .................ucuccaaguagucgccgcugA......................................................    1         1        seq
    .................ucuccaaguagucgccgcugG......................................................    1         1        seq
    .................ucuccaaguagucgcAgcugu......................................................    1         1        seq
    .................ucuccaaguagucgccgcugC......................................................    6         1        seq
    .................ucuccaaguagucgAcgcugu......................................................    1         1        seq
    .................ucuccaaguagucgccgcugu......................................................    29        0        seq
    .................Gcuccaaguagucgccgcugu......................................................    1         1        seq
    .................ucuccaaguagucgccgcuguC.....................................................    1         1        seq
    .................ucuccaaguagucgccgcuguU.....................................................    3         1        seq
    .................ucuccaaguagucgccgcugug.....................................................    16        0        seq
    .................ucuccaaguagucgccgcuguA.....................................................    3         1        seq
    .................ucuccaaguagucgccgcuguUu....................................................    1         1        seq
    .................ucuccaaguagucgccgcugugC....................................................    3         1        seq
    .................ucuccaaguagucgccgcugugA....................................................    1         1        seq
    .................ucuccaaguagucgccgcuguAu....................................................    1         1        seq
    .................Acuccaaguagucgccgcugugu....................................................    1         1        seq
    .................ucuccaaguagucgccgcugugu....................................................    19        0        seq
    .................ucuccaaguagucgccgcuguguG...................................................    1         1        seq
    .................ucuccaaguagucgccgcuguguU...................................................    3         1        seq
    .................ucuccaaguagucgccgcuguguc...................................................    4         0        seq
    .................ucuccaaguagucgGcgcuguguc...................................................    1         1        seq
    .................ucuccaagAagucgccgcugugucg..................................................    1         1        seq
    .................ucuccaaguagucgccgcugugucU..................................................    4         1        seq
    .................ucuccaaguagucgccgcugugucg..................................................    13        0        seq
```

```
                                                     novel-nve-miR-88_star
                 novel-nve-miR-88_guide
agccaugucuagccuagucuccaaguagucgccgcugugucgcucauuuuaccucacagcgguuacuuggagucuaugucuagccaugcuac

.................ucuccaaguagucgccgcugugucA..................................................    3    1    seq
.....................................................ucacagcgguuacuuggaU....................    1    1    seq
.....................................................ucacagcgguuacuuggag....................    1    0    seq
.....................................................ucacagcgguuacuuggagu...................    1    0    seq
.....................................................ucacagcgguuacuuggaguU..................    1    1    seq
.....................................................ucacagcgguuacuuggaguc..................    2    0    seq
.....................................................ucacagcgguuacuuggagucu.................    9    0    seq
.....................................................ucacagcgguuacuuggagucA.................    1    1    seq
.....................................................ucacagcgguuacuuggagucuU................    2    1    seq
```

miRBase precursor : novel-nve-miR-89
Total read count : 1076
novel-nve-miR-89_guide read count : 988
novel-nve-miR-89_star read count : 82
remaining reads : 6

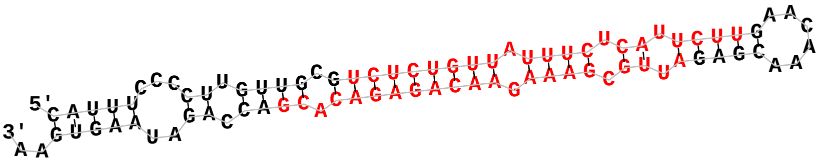

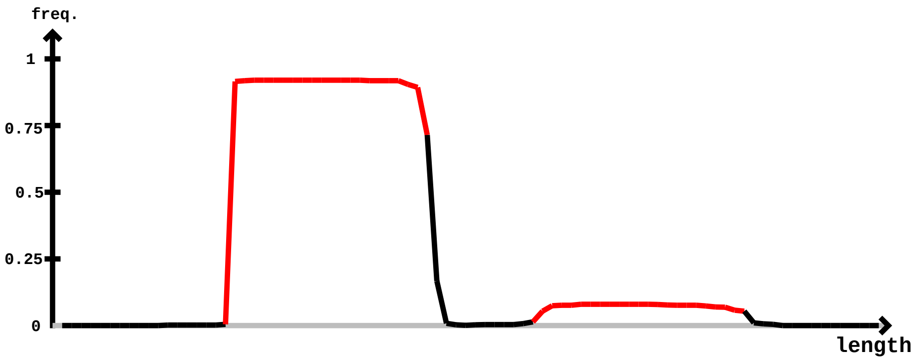


```
                novel-nve-miR-89_guide
                                         novel-nve-miR-89_star
5'- cauuuccccuuguugcgucucuguuauuucucauucuugaacaaacgagauugcgaaagaacagagacacgaccagauaaguguaa  -3'    exp
    (((((...((.((((.(((((((((.((((.((.(((((......))))).)).)))).))))))))).)))).))..)))))..   reads      mm         sample
    ..........uguugcgucucuguuauuucu.....................................................   2          0          seq
    ................gucucuguuauuucucauucuu..............................................   1          0          seq
    ................gucucuguuauuucucauucuG..............................................   1          1          seq
    ................Cucucuguuauuucucauucuu..............................................   1          1          seq
    .................ucucuguuauuucucauC.................................................   4          1          seq
    .................ucucuguuauuuGucauu.................................................   1          1          seq
    .................ucucuguuauuucucauu.................................................   10         0          seq
    .................ucucuguuauuucucauuU................................................   2          1          seq
    .................ucucuguuauuucucauuc................................................   10         0          seq
    .................ucucugCuauuucucauucu...............................................   1          1          seq
    .................ucucuguuauuucucGuucu...............................................   1          1          seq
    .................ucucuguuauuucucauucA...............................................   6          1          seq
    .................ucucuguuauuucCcauucu...............................................   2          1          seq
    .................ucucuguuauuucucauucC...............................................   41         1          seq
    .................ucucuguuauuuAucauucu...............................................   1          1          seq
    .................ucucuguAauuucucauucu...............................................   1          1          seq
    .................ucucuguCauuucucauucu...............................................   1          1          seq
    .................ucucCguuauuucucauucu...............................................   1          1          seq
    .................ucucuguuauuucucauucu...............................................   131        0          seq
    .................Ccucuguuauuucucauucu...............................................   1          1          seq
    .................ucucuguuauuucuAauucu...............................................   1          1          seq
    .................ucucuguuauCucucauucu...............................................   2          1          seq
    .................ucAcuguuauuucucauucu...............................................   1          1          seq
    .................ucucuguuauuucucauucG...............................................   1          1          seq
    .................ucucuguuauuucucauucuC..............................................   83         1          seq
    .................ucucuguuaGuucucauucuu..............................................   1          1          seq
    .................ucucuguuaCuucucauucuu..............................................   3          1          seq
    .................ucuAuguuauuucucauucuu..............................................   1          1          seq
    .................ucucuguuGuuucucauucuu..............................................   2          1          seq
    .................Ccucuguuauuucucauucuu..............................................   3          1          seq
    .................ucucuguuCuuucucauucuu..............................................   1          1          seq
    .................ucCcuguuauuucucauucuu..............................................   1          1          seq
    .................ucucuCuuauuucucauucuu..............................................   1          1          seq
    .................uUucuguuauuucucauucuu..............................................   2          1          seq
```

```
                 novel-nve-miR-89_guide
                                                 novel-nve-miR-89_star
cauuuccccuuguugcgucucuguuauuucucauucuugaacaaacgagauugcgaaagaacagagacacgaccagauaagugaa

.................ucAcuguuauuucucauucuu...............................................   1     1     seq
.................ucucuguuauuCcucauucuu...............................................   2     1     seq
.................ucucuguuauCucucauucuu...............................................   1     1     seq
.................ucucCguuauuucucauucuu...............................................   1     1     seq
.................ucucugCuauuucucauucuu...............................................   1     1     seq
.................ucucuguuauuucucGuucuu...............................................   3     1     seq
.................ucucuguuauuucucauucuG...............................................   3     1     seq
.................ucucuUuuauuucucauucuu...............................................   1     1     seq
.................ucucugAuauuucucauucuu...............................................   1     1     seq
.................ucucuguuauuucuUauucuu...............................................   1     1     seq
.................ucucGguuauuucucauucuu...............................................   1     1     seq
.................Acucuguuauuucucauucuu...............................................   3     1     seq
.................uAucuguuauuucucauucuu...............................................   1     1     seq
.................ucucuguuauuAcucauucuu...............................................   2     1     seq
.................ucucuguuauuucAcauucuu...............................................   1     1     seq
.................ucucuguuauuucucauucuu...............................................   450   0     seq
.................ucucuguuauuucucauuAuu...............................................   2     1     seq
.................ucucuguuauuucucauucCu...............................................   1     1     seq
.................ucucuguuauuucucauucuA...............................................   8     1     seq
.................ucucuguuauuucucaCucuu...............................................   1     1     seq
.................ucucuguuauuucucauucuug..............................................   129   0     seq
.................ucucuguuauuucAcauucuug..............................................   1     1     seq
.................ucucuguuauuCcucauucuug..............................................   1     1     seq
.................ucucuguuauuucucauucuuA..............................................   9     1     seq
.................Acucuguuauuucucauucuug..............................................   1     1     seq
.................ucucuguuauuucucauucuuC..............................................   3     1     seq
.................ucucuguuauuucucauCcuug..............................................   1     1     seq
.................ucucuguuauuucucauucuuU..............................................   21    1     seq
.................ucucuguuauuUucauucuug..............................................   1     1     seq
.................Ccucuguuauuucucauucuug..............................................   2     1     seq
.................ucucugAuauuucucauucuug..............................................   1     1     seq
.................ucucCguuauuucucauucuug..............................................   1     1     seq
.................ucucuguuauuucucauucuuUa.............................................   2     1     seq
.................ucucuguuauuucucauucuugC.............................................   1     1     seq
.................ucucuguuauuucucauucuugU.............................................   2     1     seq
.................ucucuguuauuucucauucuugaU............................................   1     1     seq
.................ucucuguuauuucucauucuugaC............................................   1     1     seq
..................cucuguuauuucucauucuu...............................................   3     0     seq
...................ucuguuauuucucauucuuUa.............................................   1     1     seq
...................ucugCuauuucucauucuugaa............................................   1     1     seq
.........................................caaacgagaCugcgaaagaa........................   1     1     seq
..........................................aaacgagaCugcgaaagaac.......................   2     1     seq
...........................................aacgagaCugcgaaagaaca......................   1     1     seq
...............................................agauugcgaaagaacagag...................   2     0     seq
...............................................agauugcgaaagaacagaga..................   1     0     seq
...............................................Ggauugcgaaagaacagagaca................   1     1     seq
................................................gauugcgaaagaacagag...................   1     0     seq
................................................gauugcgaaagaacagagacacA..............   3     1     seq
................................................gauugcgaaagaacagagacacU..............   3     1     seq
.................................................auugcgaaagaacagaga..................   3     0     seq
.................................................auugcgaaagaacagagac.................   1     0     seq
.................................................auugcgaaagaacagagaca................   7     0     seq
.................................................auugcgaaagaacagagacac...............   1     0     seq
.................................................auugcgaaagaacagagCcac...............   1     1     seq
.................................................auugcgaaagaacagagacaU...............   1     1     seq
.................................................auugcgaaagaacagagacacC..............   5     1     seq
.................................................auugcgaaagaacagagacacA..............   5     1     seq
.................................................auugcgaaagaacagagacacU..............   19    1     seq
..................................................uugcgaaagaacagagaca................   4     0     seq
..................................................uugcgaaagaacagagGcac...............   1     1     seq
..................................................uugcgaaagaacagagacacA..............   1     1     seq
..................................................uugcgaaagaacagagacacC..............   1     1     seq
..................................................uugcgaaagaacagagacacU..............   8     1     seq
..................................................uugcgaaagaacagagacacga.............   1     0     seq
..................................................uugcgaaagaGcagagacacga.............   1     1     seq
..................................................uugcgaaagaacagagacacgacc...........   1     0     seq
..................................................uugcgaaagaacagagacacgacG...........   1     1     seq
..................................................uugcgaaagaacagagacacgacU...........   3     1     seq
...................................................ugcgaaagaacagagacacC..............   1     1     seq
```

```
                 novel-nve-miR-89_guide
                                                 novel-nve-miR-89_star
cauuuccccuuguugcgucucuguuauuucucauucuugaacaaacgagauugcgaaagaacagagacacgaccagauaagugaa

...................................................ugcgaaagaacagagacacU..............    1          1          seq
.....................................................cgaaagaacagagacGcga.............    2          1          seq
.....................................................cgaaagaacagagacGcgac............    2          1          seq
```

miRBase precursor : novel-nve-miR-91
Total read count : 174
novel-nve-miR-91_guide read count : 159
novel-nve-miR-91_star read count : 15
remaining reads : 0

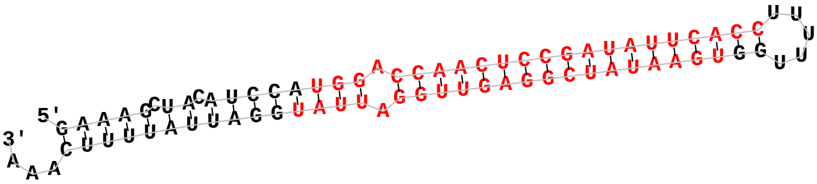

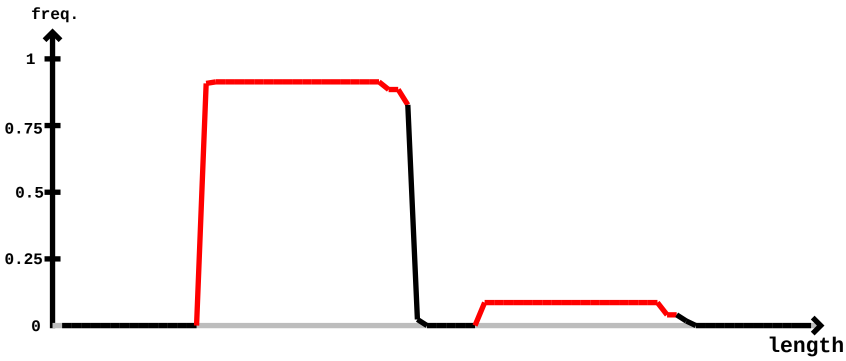


```
                 novel-nve-miR-91_guide
                                         novel-nve-miR-91_star
5'- gaaagcuacauccauggaccaacuccgauauucaccuuuuuggugaauaucggaguuggauuauggauuauuuucaaa   -3'    exp
    (((((.((.((((((((.((((((((((((((((((.....)))))))))))))))))).)))))))))))))))...   reads     mm      sample
    ..............Aggaccaacuccgauauuc.............................................   1         1       seq
    ..............uggaccaacuccgauauuc.............................................   4         0       seq
    ..............uggaccaacuccgauauucaU...........................................   1         1       seq
    ..............uggaccaacuccgauauucac...........................................   9         0       seq
    ..............uggaccaacuccgauauucacA..........................................   13        1       seq
    ..............uggaccaacuccgauauucacU..........................................   37        1       seq
    ..............uggacAaacuccgauauucacc..........................................   1         1       seq
    ..............uggaccaacuccgauauucacc..........................................   86        0       seq
    ..............uggaccaacuccgauauCcacc..........................................   2         1       seq
    ..............uggaccaacuccgauauucaccu.........................................   2         0       seq
    ..............uggaccaacuccgauauucaccC.........................................   1         1       seq
    ..............uggaccaacuccgauauucacAu.........................................   1         1       seq
    ...............Ugaccaacuccgauauucacc..........................................   1         1       seq
    ...........................................ugaauaucggaguuggauu................   8         0       seq
    ...........................................ugaauaucggaguuggauuaA..............   3         1       seq
    ...........................................ugaauaucggaguuggauuaG..............   1         1       seq
    ...........................................ugaauaucggaguuggauuaug.............   3         0       seq
```

miRBase precursor : novel-nve-miR-92
Total read count : 241
novel-nve-miR-92_guide read count : 233
novel-nve-miR-92_star read count : 6
remaining reads : 2

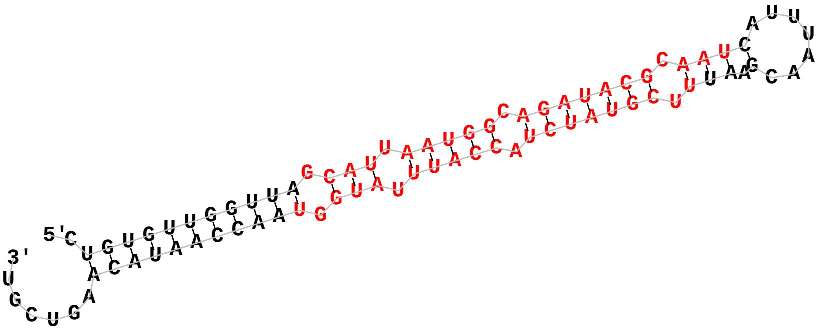

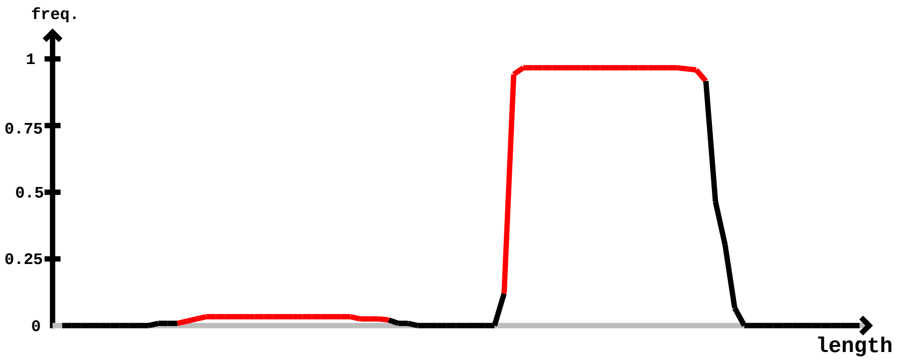


novel-nve-miR-92_star

novel-nve-miR-92_guide

```
5'- cuguguugguuagcauuaauggcagauacgcaaucauuuaacgaauuucguaucuaccauuuaugguaaccaauacaagucgu   -3'    exp
    .((((((((((.(((.(((((.(((((((.((((........).))).))))))).))))).))).)))))))))).......    reads      mm        sample
    .........uuagcauuaauggcagauacg.....................................................    2          0         seq
    ............gcauuaauggcagauacgcaau.................................................    1          0         seq
    ............gcauuaauggcagauacgcaaA.................................................    1          1         seq
    .............cauuaauggcagauacgcaa..................................................    1          0         seq
    .............cauuaauggcagauacgcaaC.................................................    1          1         seq
    ..............auuaauggcagauacgcaaucG...............................................    1          1         seq
    ..............auuaauggcagauacgcaauca...............................................    1          0         seq
    .............................................uuucguaucuaccauuuau...................    1          0         seq
    .............................................uuucguaucuaccauuuaug..................    1          0         seq
    .............................................uuucguaucuaccauuuaugg.................    2          0         seq
    .............................................uuucguaucuaccauuuauggu................    12         0         seq
    .............................................uuucguaucuGccauuuauggu................    1          1         seq
    .............................................uuucguaucuaccauuuauggC................    7          1         seq
    .............................................uuucguaucuaccauuuauggG................    1          1         seq
    .............................................uuucguaucuaccauuuaugguC...............    1          1         seq
    .............................................uuucguaucuaccauuuaugguU...............    2          1         seq
    .............................................uuucguaucuaccauuuaugguUa..............    1          1         seq
    ..............................................uucguaucuaccauuuaugg.................    7          0         seq
    ..............................................uucguaucuaccauuuaugA.................    1          1         seq
    ..............................................uucguaucuGccauuuauggu................    8          1         seq
    ..............................................uucguaucAaccauuuauggu................    1          1         seq
    ..............................................uucguaucuaccauuuauggu................    65         0         seq
    ..............................................uucguaucuaccauuuauggA................    1          1         seq
    ..............................................uucguaucuaccauuuauggC................    11         1         seq
    ..............................................uucguGucuaccauuuauggu................    1          1         seq
    ..............................................uucguaucuGccauuuauggua...............    2          1         seq
    ..............................................uucguaucuaccauuuaugguU...............    19         1         seq
    ..............................................Aucguaucuaccauuuauggua...............    1          1         seq
    ..............................................uucguaucuaccauuuauggua...............    10         0         seq
    ..............................................uucguaucuaccauuuaugguC...............    4          1         seq
    ..............................................Aucguaucuaccauuuaugguaa..............    1          1         seq
    ..............................................uucguaucuaccauuuaugguaa..............    27         0         seq
    ..............................................uucguaucuaccauuuaugguaC..............    5          1         seq
    ..............................................uucguaucuGccauuuaugguaa..............    4          1         seq
```

```
            novel-nve-miR-92_star
                                              novel-nve-miR-92_guide
cuguguugguuagcauuaauggcagauacgcaaucauuuaacgaauuucguaucuaccauuuaugguaaccaauacaagucgu

..............................................uucguaucuaccauuuaugguaU..............    14    1    seq
..............................................uucguaucuaccauuuaugguaaA.............    2     1    seq
..............................................uucguaucuaccauuuaugguaac.............    2     0    seq
..............................................uucguaucuaccauuuaugguaaU.............    12    1    seq
...............................................ucguaucuaccauuuauggu................    1     0    seq
...............................................ucguaucuaccauuuaugguaa..............    5     0    seq
```

miRBase precursor : novel-nve-miR-94
Total read count : 79
novel-nve-miR-94_guide read count : 79
novel-nve-miR-94_star read count : 0
remaining reads : 0

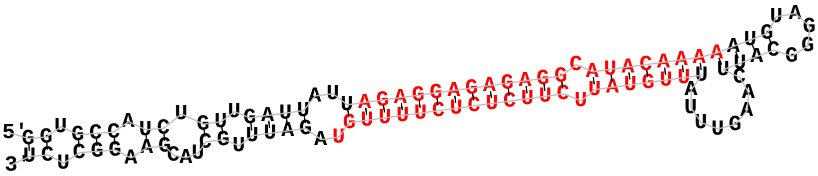

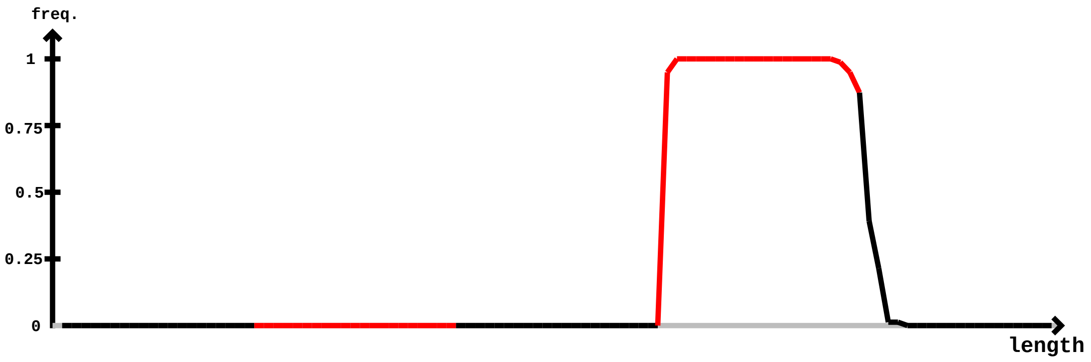


```
                                      novel-nve-miR-94_star
                                                                          novel-nve-miR-94_guide
5'- ggugccaucuguugauuauuagaggagagaggcauacaaaaauguagggcauucaaguuuauuuguauucuucucucuuuuguagauuugcuacgaaggcucu   -3'    exp
    ((.(((.((.((.((((..((((((((((((.((((((((((.....)))).......))))))).)))))))))))))..)))).))...)).))).))   reads      mm          sample
    ..............................................................uuguauucuucucucuuC.......................   1          1           seq
    ..............................................................uuguauucuucucucuuuu......................   3          0           seq
    ..............................................................uuguauucuucucucuuuug.....................   5          0           seq
    ..............................................................uuguauucuucucucuuuuA.....................   1          1           seq
    ..............................................................uuguauucuucucucuuuugC....................   9          1           seq
    ..............................................................uuguauucuucucucuuuugG....................   1          1           seq
    ..............................................................uAguauucuucucucuuuugu....................   1          1           seq
    ..............................................................uuguauucuucucucuuuugu....................   25         0           seq
    ..............................................................uuguauucuucucAcuuuugu....................   1          1           seq
    ..............................................................uuguauucuucucucuuuugA....................   1          1           seq
    ..............................................................uuguauucuucucucuuuuguC...................   1          1           seq
    ..............................................................uuguauucuucucucuuuCgua...................   1          1           seq
    ..............................................................uuguauucuucucucuuuugua...................   4          0           seq
    ..............................................................uuguauucuucucucuuuuguU...................   6          1           seq
    ..............................................................uuguauucuucucucuuuuguaA..................   2          1           seq
    ..............................................................uuguauucuucucucuuuuguaC..................   1          1           seq
    ..............................................................uuguauucuucucucuuuuguag..................   2          0           seq
    ..............................................................uuguauucuucucucuuuuguaU..................   9          1           seq
    ..............................................................uuguauucuucucucuuuuguaUau................   1          1           seq
    ...............................................................uguauucuucucucuuuuguU...................   1          1           seq
    ...............................................................uguauucuucucucuuuugua...................   1          0           seq
    ...............................................................uguauucuucucucuuuuguaU..................   2          1           seq
```

miRBase precursor : novel-nve-miR-96
Total read count : 2519
novel-nve-miR-96_guide read count : 2517
novel-nve-miR-96_star read count : 2
remaining reads : 0

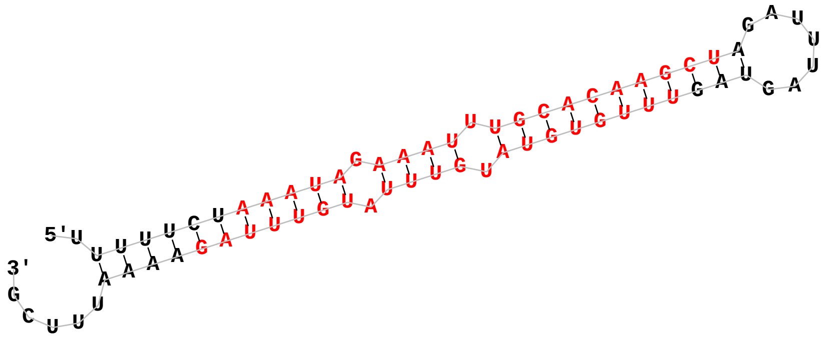

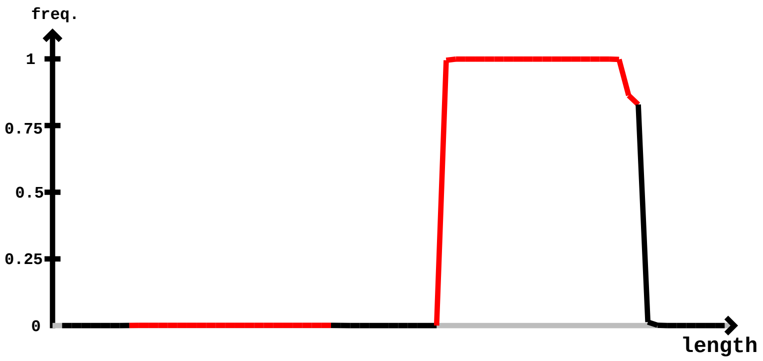


```
                                        novel-nve-miR-96_guide
          novel-nve-miR-96_star
5'- uuuuucuaaauagaaauuugcacaagcuagauuuaguaguuuguguauguuuauguuuagaaaauuucg   -3'    exp
    .((((((((((.((((.((((((((((.......)))))))))).)))).)))))))))).........    reads    mm    sample
    ......uaaauagaaauuugcacaagcu.........................................    1        0     seq
    .......aaauagaaauuugcacaagcuU........................................    1        1     seq
    .......................................uuuguguauguuuCuguu............    3        1     seq
    .......................................uuuguguauguuuauguuu...........    3        0     seq
    .......................................uuuguguauguuuauguuC...........    1        1     seq
    .......................................uuuguguauguuuCuguuu...........    336      1     seq
    .......................................uuuguguauguuuCuguuua..........    86       1     seq
    .......................................uuuguguauguuuUuguuuag.........    1        1     seq
    .......................................uuuguguauAuuuauguuuag.........    1        1     seq
    .......................................uuuguguauguuuCuguuuag.........    2025     1     seq
    .......................................uuuguguauguuuauguuuaA.........    1        1     seq
    .......................................uuuguguauguuuauguuuag.........    16       0     seq
    .......................................uuuguguauguuuauguuuaga........    1        0     seq
    .......................................uuuguguauguuuCuguuuaga........    29       1     seq
    .......................................uuuguguauguuuCuguuuagaa.......    2        1     seq
    .......................................uuuguguauguuuauguuuagUa.......    1        1     seq
    ........................................uuguguauguuuCuguuuag.........    11       1     seq
```

miRBase precursor : novel-nve-miR-102-1
Total read count : 229
novel-nve-miR-102-1_guide read count : 207
novel-nve-miR-102-1_star read count : 21
remaining reads : 1

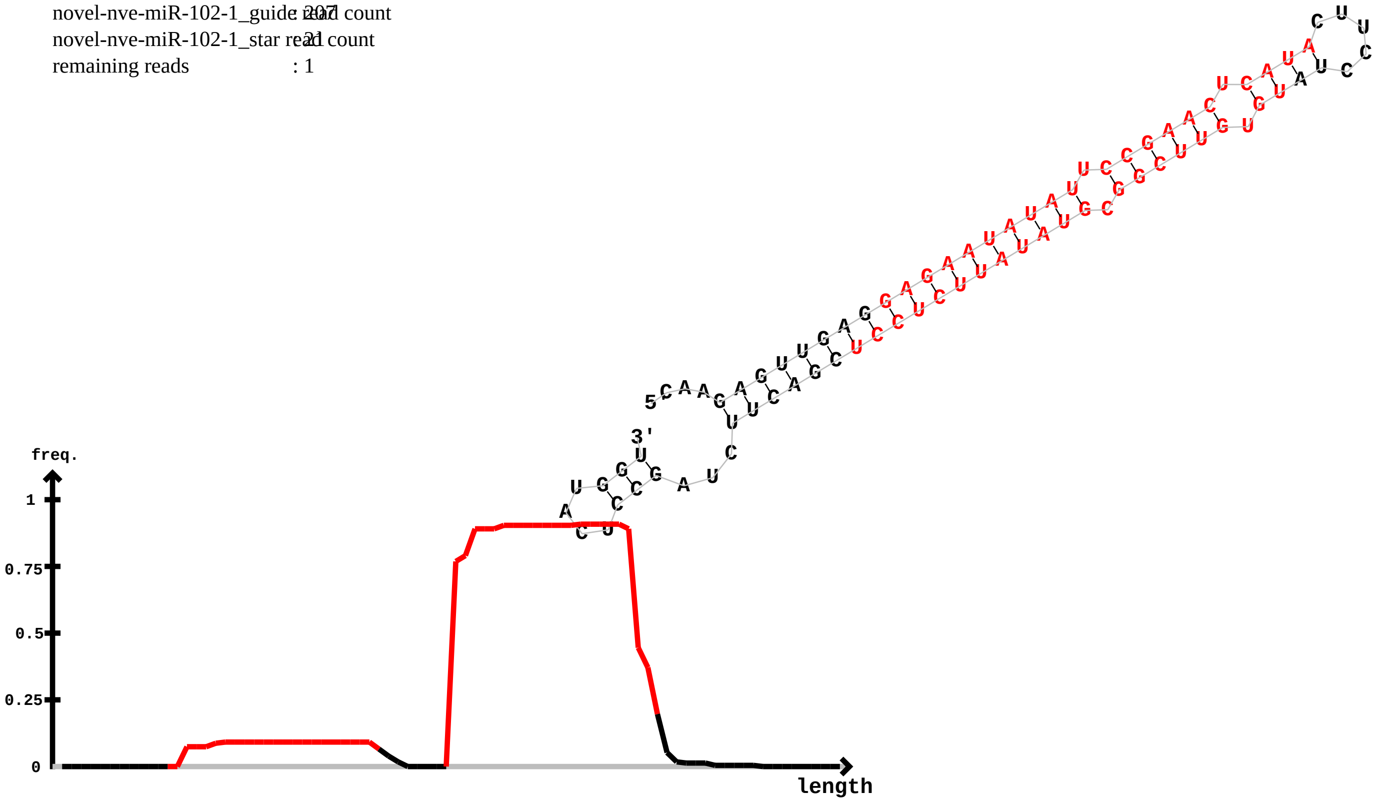


```
                                    novel-nve-miR-102-1_guide
              novel-nve-miR-102-1_star
5'- caagaguugaggagaauauauuccgaacucauacuuccuauguguucggcguauauucuccucgacuucuagccucauggu   -3'    exp
    ...((((((((((((((((((.((((((.((((.....)))).)))))).))))))))))))))))))...(((....)))    reads      mm         sample
    ............agaauauauuccgaacucaC.................................................    1          1          seq
    ............agaauauauuccgaacucau.................................................    4          0          seq
    ............agaGuauauuccgaacucau.................................................    1          1          seq
    ............agaauauauuccgaacucaua................................................    4          0          seq
    ............agaauauauuccgaacucauU................................................    2          1          seq
    ............agaauauauuccgaacucauaU...............................................    4          1          seq
    ............Ggaauauauuccgaacucauac...............................................    1          1          seq
    ...............auauauuccgaacucauacu..............................................    2          0          seq
    ...............auauauuccgaacucauacC..............................................    1          1          seq
    ................uauauuccgaacucauaUu..............................................    1          1          seq
    ........................................uguguucggcguauauuc.......................    3          0          seq
    ........................................uguguucggcguauauuU.......................    1          1          seq
    ........................................uguguucggcguauauucC......................    15         1          seq
    ........................................uguguucggcguauauucA......................    1          1          seq
    ........................................uguguucggcguauauucG......................    13         1          seq
    ........................................ugCguucggcguauauucu......................    2          1          seq
    ........................................uguguucggcguauauucu......................    65         0          seq
    ........................................Gguguucggcguauauucu......................    1          1          seq
    ........................................uguguucggcguauauuUu......................    2          1          seq
    ........................................uguguucggcguauUuucu......................    1          1          seq
    ........................................uguguucggcguauauucAc.....................    1          1          seq
    ........................................uguguucggcguauauucuc.....................    8          0          seq
    ........................................uguguucggcguauauucuU.....................    2          1          seq
    ........................................uguguucggcguauauucucG....................    2          1          seq
    ........................................uguguucggcguauauucucc....................    17         0          seq
    ........................................uguguucggcguauauucucA....................    5          1          seq
    ........................................uguguucggcguauauucucU....................    9          1          seq
    ........................................uguguucggcguUuauucucc....................    1          1          seq
    ........................................uguguucggcguauauucuAc....................    2          1          seq
    ........................................uguguucggcguauauucuccG...................    3          1          seq
    ........................................ugugCucggcguauauucuccu...................    1          1          seq
    ........................................uguguucggcguauauCcuccu...................    1          1          seq
    ........................................uguguucggcguauauucuccC...................    2          1          seq
    ........................................uguguucggcguauauucuccA...................    2          1          seq
```

novel-nve-miR-102-1_guide

novel-nve-miR-102-1_star

caagaguugaggagaauauauuccgaacucauacuuccuauguguucggcguauauucuccucgacuucuagccucauggu

| | | | |
|---|---|---|---|
| ........................................uguguucggcguauauucuccu................... | 15 | 0 | seq |
| ........................................uguguucggcguauauucuccuU.................. | 1 | 1 | seq |
| .........................................guguucggcguauauucu...................... | 2 | 0 | seq |
| .........................................guguucggcguauauucucc.................... | 1 | 0 | seq |
| .........................................guguucggcAuauauucucc.................... | 1 | 1 | seq |
| .........................................guguucggcguauauucuccuU.................. | 1 | 1 | seq |
| ..........................................uguucggcguauauucuc..................... | 3 | 0 | seq |
| ..........................................uguucggcguauauucuU..................... | 2 | 1 | seq |
| ..........................................uguucggcguauauucuA..................... | 1 | 1 | seq |
| ..........................................uguucggcguauauucucU.................... | 2 | 1 | seq |
| ..........................................uguucggcguauauucuccA................... | 3 | 1 | seq |
| ..........................................uguucggcguauauucuccu................... | 2 | 0 | seq |
| ..........................................uguucggcguauauucuccC................... | 2 | 1 | seq |
| ..........................................uguucggcguauauucuccG................... | 1 | 1 | seq |
| ..........................................uguucggcguauauucuAcu................... | 1 | 1 | seq |
| ..........................................uguucggcguauauucuccuc.................. | 1 | 0 | seq |
| ..........................................uguucggcguauauucuccuU.................. | 5 | 1 | seq |
| .............................................ucggcAuauauucuccucg................. | 1 | 1 | seq |
| .............................................ucggcguauauucuccucgacu.............. | 2 | 0 | seq |
| .....................................................uauucuccucgacuucuCg......... | 1 | 1 | seq |

miRBase precursor : novel-nve-miR-103-1
Total read count : 2972
novel-nve-miR-103-1_guide read count : 2932
novel-nve-miR-103-1_star read count : 19
remaining reads : 21

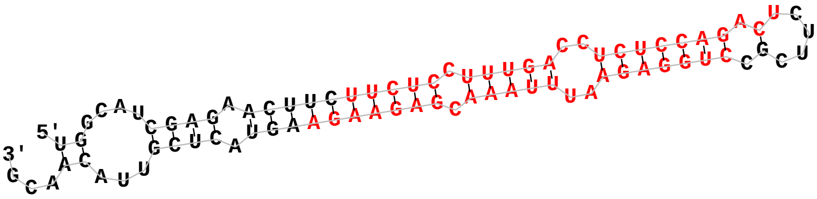

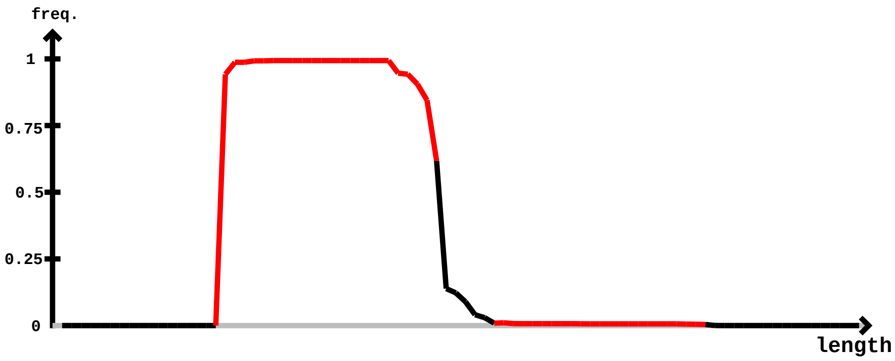


novel-nve-miR-103-1_guide

novel-nve-miR-103-1_star

5'- uggcaucgagaacuucuucuccuuugaccucuccagacucuucgccuggagaauuuaaacgagaagaaguacucguuacaacg -3' exp

| ((....((((.((((((((((.(((((..(((((((.(.....).)))))))..))))).)))))))))).)))...))... | reads | mm | sample |
|---|---|---|---|
| ................uuUuccuuugaccucucc................................................. | 1 | 1 | seq |
| ................uuGuccuuugaccucucc................................................. | 1 | 1 | seq |
| ................uucuccuuugaccucucA................................................. | 13 | 1 | seq |
| ................uCcuccuuugaccucucc................................................. | 1 | 1 | seq |
| ................uucuccuuugaccucCcc................................................. | 1 | 1 | seq |
| ................uucuccuuugacUucucc................................................. | 28 | 1 | seq |
| ................uucuccuAugaccucucc................................................. | 2 | 1 | seq |
| ................uucuccuuugaccucucU................................................. | 23 | 1 | seq |
| ................uucuccuCugaccucucc................................................. | 1 | 1 | seq |
| ................Cucuccuuugaccucucc................................................. | 1 | 1 | seq |
| ................uucuccuuugaUcucucc................................................. | 6 | 1 | seq |
| ................uucuccuuugaccucucc................................................. | 58 | 0 | seq |
| ................uucuccuuugCccucucc................................................. | 2 | 1 | seq |
| ................uucuccuuugGccucucc................................................. | 2 | 1 | seq |
| ................uucuccuuugaccucuUc................................................. | 1 | 1 | seq |
| ................uucuccuuugaccuUucca................................................ | 1 | 1 | seq |
| ................uucuccuuugaccucuccC................................................ | 1 | 1 | seq |
| ................uucuccuuugaccucucca................................................ | 9 | 0 | seq |
| ................uucuccuuugaccucucUag............................................... | 3 | 1 | seq |
| ................uucucUuuugaccucuccag............................................... | 1 | 1 | seq |
| ................uucuccuuugaUcucuccag............................................... | 9 | 1 | seq |
| ................uucuccuuugacUucuccag............................................... | 11 | 1 | seq |
| ................uucuUcuuugaccucuccag............................................... | 1 | 1 | seq |
| ................uucuccuuAgaccucuccag............................................... | 1 | 1 | seq |
| ................uucuccuuuCaccucuccag............................................... | 1 | 1 | seq |
| ................uuUuccuuugaccucuccag............................................... | 1 | 1 | seq |
| ................uucuccuuugaccucuccaU............................................... | 4 | 1 | seq |
| ................uucuccuuugacAucuccag............................................... | 1 | 1 | seq |
| ................uucuccuuugaccucuUcag............................................... | 2 | 1 | seq |
| ................uucuccuuugaccucuccaA............................................... | 9 | 1 | seq |
| ................uucuccuuugaccucuccaC............................................... | 1 | 1 | seq |
| ................uAcuccuuugaccucuccag............................................... | 1 | 1 | seq |
| ................uucuccuuugaccucuccag............................................... | 63 | 0 | seq |
| ................uucuccuuugaccucuccaga.............................................. | 92 | 0 | seq |

```
                novel-nve-miR-103-1_guide
                                             novel-nve-miR-103-1_star
uggcaucgagaacuucuucuccuuugaccucuccagacucuucgccuggagaauuuaaacgagaagaaguacucguuacaacg

................uucuccuuugaccucCccaga..............................................   1     1   seq
................uucuccuuugaccCcuccaga..............................................   1     1   seq
................uucuUcuuugaccucuccaga..............................................   3     1   seq
................uucuccuuugaccucuUcaga..............................................   2     1   seq
................uucucUuuugaccucuccaga..............................................   1     1   seq
................uucuccuuugacAucuccaga..............................................   1     1   seq
................uucuccGuugaccucuccaga..............................................   1     1   seq
................uucuccuuugaccucuccagU..............................................   2     1   seq
................uucuccuuugacUucuccaga..............................................   60    1   seq
................uucuccuAugaccucuccaga..............................................   1     1   seq
................uucuccuuugaUcucuccaga..............................................   5     1   seq
................uucuccCuugaccucuccaga..............................................   1     1   seq
................uucuccuuugGccucuccaga..............................................   2     1   seq
................uucuccuuugaccucuccagG..............................................   5     1   seq
................uucuccuuugaccucuGcagac.............................................   1     1   seq
................uucuccuuugaccucucUagac.............................................   3     1   seq
................uucuccuuCgaccucuccagac.............................................   1     1   seq
................uucuccuAugaccucuccagac.............................................   1     1   seq
................uucuccuuugaccucuccagac.............................................   363   0   seq
................uucucUuuugaccucuccagac.............................................   1     1   seq
................uucCccuuugaccucuccagac.............................................   2     1   seq
................Cucuccuuugaccucuccagac.............................................   4     1   seq
................uucuccuuugaccucuccagaA.............................................   11    1   seq
................uucuccuuuCaccucuccagac.............................................   1     1   seq
................uCcuccuuugaccucuccagac.............................................   2     1   seq
................uucuccuuugaccucuccagaU.............................................   66    1   seq
................uucuccuuugaccucuccGgac.............................................   1     1   seq
................uuUuccuuugaccucuccagac.............................................   4     1   seq
................uucuccuuugacUucuccagac.............................................   111   1   seq
................uucuAcuuugaccucuccagac.............................................   1     1   seq
................uucuccuuugaccCcuccagac.............................................   3     1   seq
................uucuccuuugaccucuccagGc.............................................   5     1   seq
................uucuccuuugaccucucAagac.............................................   2     1   seq
................uucuccuuugaUcucuccagac.............................................   49    1   seq
................uucuccuCugaccucuccagac.............................................   1     1   seq
................uucuccuuugCccucuccagac.............................................   1     1   seq
................uuAuccuuugaccucuccagac.............................................   1     1   seq
................uucuccuuugaAcucuccagac.............................................   1     1   seq
................uAcuccuuugaccucuccagac.............................................   1     1   seq
................uucuUcuuugaccucuccagac.............................................   9     1   seq
................uucuccuuugaccucuUcagac.............................................   9     1   seq
................uucucAuuugaccucuccagac.............................................   1     1   seq
................Aucuccuuugaccucuccagac.............................................   3     1   seq
................uucuccuuugaccucCccagac.............................................   4     1   seq
................Gucuccuuugaccucuccagac.............................................   1     1   seq
................uucuccCuugaccucuccagac.............................................   2     1   seq
................uucuccuuugaccuUuccagac.............................................   1     1   seq
................uGcuccuuugaccucuccagacu............................................   1     1   seq
................Aucuccuuugaccucuccagacu............................................   3     1   seq
................uucuccuuugaccucuccagacu............................................   754   0   seq
................uucuccuuugaccucuccagacG............................................   22    1   seq
................uucuccuuugaccucucGagacu............................................   4     1   seq
................uAcuccuuugaccucuccagacu............................................   4     1   seq
................uucuccuuugaUcucuccagacu............................................   43    1   seq
................uucuccuuugacUucuccagacu............................................   206   1   seq
................uucuccuuugaccucuccUgacu............................................   1     1   seq
................uucuccuuugaccucuccGgacu............................................   7     1   seq
................Gucuccuuugaccucuccagacu............................................   2     1   seq
................uucuccAuugaccucuccagacu............................................   2     1   seq
................uucuccuuugaccucuccagacC............................................   119   1   seq
................uucuccuuugaccucuccagaGu............................................   1     1   seq
................uucuccuuCgaccucuccagacu............................................   6     1   seq
................uucucUuuugaccucuccagacu............................................   3     1   seq
................uucuccuuugaccucuccagaUu............................................   2     1   seq
................uucuccuuugaAcucuccagacu............................................   1     1   seq
................uucuccuuugGccucuccagacu............................................   1     1   seq
................uucuccuuugaccCcuccagacu............................................   7     1   seq
................uucuccuuugaGcucuccagacu............................................   1     1   seq
................uucuccuAugaccucuccagacu............................................   2     1   seq
```

```
                novel-nve-miR-103-1_guide
                                             novel-nve-miR-103-1_star
uggcaucgagaacuucuucuccuuugaccucuccagacucuucgccuggagaauuuaaacgagaagaaguacucguuacaacg

................uucCccuuugaccucuccagacu............................................	1	1	seq
................uuUuccuuugaccucuccagacu............................................	5	1	seq
................uucuccuuugaccucuUcagacu............................................	20	1	seq
................uucuccuuugacAucuccagacu............................................	1	1	seq
................uucuccuCugaccucuccagacu............................................	2	1	seq
................uucuccuuugaccucuGcagacu............................................	1	1	seq
................Cucuccuuugaccucuccagacu............................................	4	1	seq
................uucuccuuugaccucuccCgacu............................................	1	1	seq
................uCcuccuuugaccucuccagacu............................................	4	1	seq
................uucuccGuugaccucuccagacu............................................	1	1	seq
................uucuccuuugaccucucAagacu............................................	2	1	seq
................uucuccuuugaccucCccagacu............................................	3	1	seq
................uucuccuuugaccucuccagaAu............................................	1	1	seq
................uucuUcuuugaccucuccagacu............................................	7	1	seq
................uucuccuuugaccucuccagacA............................................	93	1	seq
................uucuccuuugaccucuccagGcu............................................	1	1	seq
................uucuccuuAgaccucuccagacu............................................	1	1	seq
................uucuccuuugaccuGuccagacu............................................	1	1	seq
................uucuccuuugaccucuccagCcu............................................	1	1	seq
................uucuccuuugaccucucUagacu............................................	20	1	seq
................uucuccCuugaccucuccagacu............................................	4	1	seq
................uucucAuuugaccucuccagacu............................................	4	1	seq
................uucuccuuugaccuUuccagacu............................................	7	1	seq
................uucuccuuugacUucuccagacuc...........................................	2	1	seq
................uuGuccuuugaccucuccagacuc...........................................	1	1	seq
................uucuccuuugaccucucGagacuc...........................................	1	1	seq
................uucuccuuugaccucuccagacuU...........................................	15	1	seq
................uucuccuuugaccucuccagacuA...........................................	10	1	seq
................uucuccuuugaccucuccagacuG...........................................	1	1	seq
................uucuccuuugaccucuccagacuc...........................................	9	0	seq
................uucuccuuugaccucuccagacuAu..........................................	37	1	seq
................uucuccuuugaAcucuccagacucu..........................................	1	1	seq
................uucuccuuugaccucuccagacucC..........................................	2	1	seq
................uucuccuuugaccucuccagacucu..........................................	23	0	seq
................uucuccuuugaccucuccagacuUu..........................................	2	1	seq
................uucuccuuugaUcucuccagacucu..........................................	2	1	seq
................uucuccuuugaccucuUcagacucu..........................................	1	1	seq
................uucuccuuugacUucuccagacucu..........................................	3	1	seq
................uCcuccuuugaccucuccagacucuu.........................................	1	1	seq
................uucuAcuuugaccucuccagacucuu.........................................	1	1	seq
................uucuccuuugaccucuccagacucuC.........................................	14	1	seq
................uucuccuuugacUucuccagacucuu.........................................	9	1	seq
................uucuccuuugaccucuccagacuAuu.........................................	1	1	seq
................uucuccuuugaccucuccagacucuu.........................................	80	0	seq
................uucuccuCugaccucuccagacucuu.........................................	1	1	seq
................uucuccuuugaccucuccagacuUuu.........................................	6	1	seq
................uucuccuuugaccucuccagacucGu.........................................	1	1	seq
................uucuccuuugaccucuGcagacucuu.........................................	1	1	seq
................Cucuccuuugaccucuccagacucuu.........................................	1	1	seq
................uucuccuuugaccucuccagacucuA.........................................	1	1	seq
................Aucuccuuugaccucuccagacucuu.........................................	2	1	seq
................uuUuccuuugaccucuccagacucuu.........................................	1	1	seq
................uucuccuuugaccucuccGgacucuu.........................................	1	1	seq
................uucuccuuugaUcucuccagacucuu.........................................	5	1	seq
................uucuccuuugacUucuccagacucuuc........................................	4	1	seq
................uucuccuuugaccucuccagacucuuU........................................	7	1	seq
................uucuccuuugaccucuccagacucuuA........................................	2	1	seq
................uucuccuuugaccucuccagacuUuuc........................................	1	1	seq
................uucuccuuugaccucuccagacucuuc........................................	8	0	seq
................uucuccuuugaccucuccagacucuucU.......................................	3	1	seq
................uucuccuuugacUucuccagacucuucg.......................................	5	1	seq
................uucuccuuugaUcucuccagacucuucg.......................................	2	1	seq
................uucuccuuugaccucuccagacucuuAg.......................................	1	1	seq
................uucuccuAugaccucuccagacucuucg.......................................	1	1	seq
................uucuccuuugaccucuccagacucuuUg.......................................	1	1	seq
................uucuccuuugaccucucAagacucuucg.......................................	1	1	seq
................uuGuccuuugaccucuccagacucuucg.......................................	1	1	seq
................uucuccuuugacAucuccagacucuucg.......................................	1	1	seq
................uucuccuuugaccucuccagacucuucA.......................................	8	1	seq
```

```
                novel-nve-miR-103-1_guide
                                             novel-nve-miR-103-1_star
uggcaucgagaacuucuucuccuuugaccucuccagacucuucgccuggagaauuuaaacgagaagaaguacucguuacaacg

................uucuccuuugGccucuccagacucuucg.......................................    1     1     seq
................uucuccuuugaccucuccagacucuucg.......................................    19    0     seq
................uucuccuuugacUucuccagacucuucgc......................................    1     1     seq
................uucuccuuugaccucuccagacucuucgc......................................    2     0     seq
................uucuccuuugaccucuccagacucuucgA......................................    1     1     seq
................uucuccuuugaccucuccagacucuucgU......................................    1     1     seq
................uucuccuuugaccucuccagacucuucgcU.....................................    8     1     seq
................uucuccuuugaccucuccagacucuucgcA.....................................    1     1     seq
................uucuccuuugaccucuccagacucuucgcUu....................................    1     1     seq
.................ucuccuuugaUcucuccag...............................................    1     1     seq
.................ucuccuuugaccucuccagac.............................................    4     0     seq
.................uUuccuuugaccucuccagac.............................................    3     1     seq
.................ucuccuuugacUucuccagacu............................................    7     1     seq
.................Acuccuuugaccucuccagacu............................................    1     1     seq
.................ucuccuuugaccucuUcagacu............................................    1     1     seq
.................uUuccuuugaccucuccagacu............................................    3     1     seq
.................ucuccuuugaccucucUagacu............................................    2     1     seq
.................ucuccuuugaccucuccagacA............................................    2     1     seq
.................ucuccuuugaccGcuccagacu............................................    1     1     seq
.................ucuccuuugaccucuccagacu............................................    15    0     seq
.................ucuccuuugaccucuccagacC............................................    4     1     seq
.................ucucUuuugaccucuccagacu............................................    5     1     seq
.................ucuccuuugaccucuccUgacu............................................    1     1     seq
.................ucuccuuugaccucuccagacuU...........................................    2     1     seq
.................ucuccuuugaccucuccagacuG...........................................    1     1     seq
.................ucuccuuugaccucuccagacuA...........................................    3     1     seq
.................ucuccuuugaccucuccagacuc...........................................    3     0     seq
.................ucuccuuugaccucuccagacucA..........................................    1     1     seq
.................ucuccuuugacUucuccagacucu..........................................    1     1     seq
.................ucuccuuugaccuUuccagacucu..........................................    2     1     seq
.................ucCccuuugaccucuccagacucu..........................................    1     1     seq
.................ucuccuuugaccucuccagacucC..........................................    5     1     seq
.................uUuccuuugaccucuccagacucu..........................................    1     1     seq
.................ucuccuuugaccucuccagacucu..........................................    10    0     seq
.................ucuccuuugaUcucuccagacucu..........................................    4     1     seq
.................ucuccuuugacUucuccagacucuu.........................................    2     1     seq
.................ucuccuuugaccucuccagacucuu.........................................    9     0     seq
.................ucuccuuugaccucuccagacucuG.........................................    1     1     seq
.................ucuccuuugaUcucuccagacucuu.........................................    4     1     seq
.................ucuccuuugaUcucuccagacucuuc........................................    2     1     seq
.................ucuccuuAgaccucuccagacucuuc........................................    1     1     seq
.................ucuccuuugaccucuccagacucuuc........................................    7     0     seq
.................ucuccuuugaccucuccagacucuuU........................................    1     1     seq
.................ucuccuuugaccCcuccagacucuucg.......................................    1     1     seq
.................ucuccuuugaUcucuccagacucuucg.......................................    7     1     seq
.................ucuccuuugaccucuccagacucuucA.......................................    1     1     seq
.................ucuccuuugaccucuccagacucuucC.......................................    1     1     seq
.................ucuccuuugaccucuccagacucuucg.......................................    5     0     seq
.................uUuccuuugaccucuccagacucuucg.......................................    1     1     seq
.................ucuccuuCgaccucuccagacucuucg.......................................    1     1     seq
.................ucuccuuugaccucuccagacucuucU.......................................    4     1     seq
.................ucuccuuugaccucuccagacucuucgUcuggagaa..............................    1     1     seq
.................ucuccuuugaccucuccagacucuucgccuggagaau.............................    1     0     seq
...................uccuuugaccuUuccaga..............................................    1     1     seq
...................uccuuugaccucuccagac.............................................    2     0     seq
...................uccuuugaccucuccagaU.............................................    1     1     seq
...................uccuuugacUucuccagac.............................................    1     1     seq
...................uccuuugaccucuccagacu............................................    3     0     seq
...................uccuuugaccucuccagacucu..........................................    1     0     seq
...................uccuuugaccucuccagacucA..........................................    1     1     seq
...................uccuuugaccucuccagacucuucg.......................................    2     0     seq
...................uccuuugaccucuccagacucuucgU......................................    4     1     seq
.....................cuuugaccucuccagacucuu.........................................    3     0     seq
............................................ccuggagaauuuaaacgaga...................    2     0     seq
............................................ccuggagaauuuaaacgagaa..................    1     0     seq
............................................ccuggagaauuuaaacgagaag.................    2     0     seq
............................................ccuggagGauuuaaacgagaag.................    1     1     seq
.............................................cuggagaauuuaaacgaga...................    1     0     seq
.............................................cuggagaauuuaaacgagGaga................    1     1     seq
```

```
                novel-nve-miR-103-1_guide
                                             novel-nve-miR-103-1_star
uggcaucgagaacuucuucuccuuugaccucuccagacucuucgccuggagaauuuaaacgagaagaaguacucguuacaacg

.............................................cuggagaGuuuaaacgagaaga................    1          1          seq
.............................................cuggagaauuuaaacgagaagG................    1          1          seq
.............................................cuggagaauuuaaacgagaaga................    7          0          seq
.............................................cuggagaauuuaaacgagaagU................    1          1          seq
.............................................cuggagaauuuaaacgagaagaC...............    1          1          seq
```

### Starless novel *Nematostella* miRNAs

miRBase precursor : novel-nve-miR-34
Total read count : 2101
novel-nve-miR-34_guide read count : 2101
novel-nve-miR-34_star read count : 0
remaining reads : 0

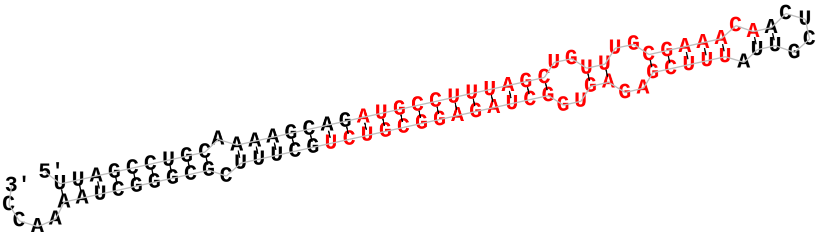

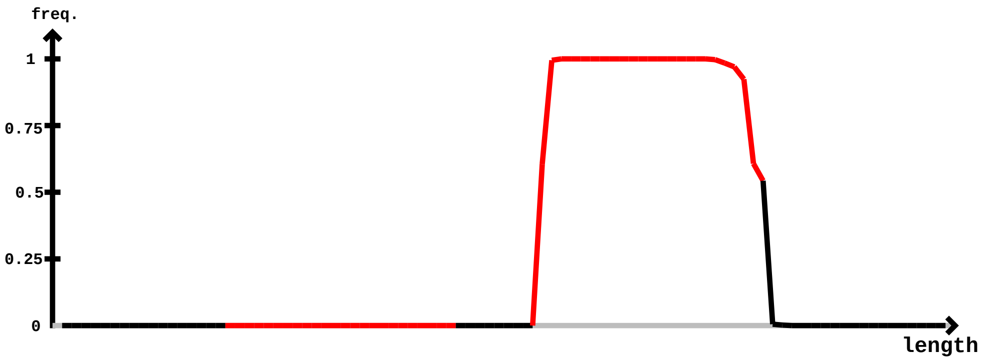


```
                                                                  novel-nve-miR-34_guide
                       novel-nve-miR-34_star
5'- uuagccugcaaaagcagaugccuuuagcuguuugcgaaacaacucguuauuucgagaguggcuagaggcgucugcuuucgcgggcuaaaacc   -3'    exp
    ((((((((.(((((((((((((((((((..((..(((((.((....)).)))))..))..)))))))))))))))))).)))))))))....     reads       mm        sample
    ..................................................uuucgagaguggcuagaA..........................     1           1         seq
    ..................................................uuucgagaguggcuagag..........................     4           0         seq
    ..................................................uuCcgagaguggcuagag..........................     1           1         seq
    ..................................................uuucgagaguggcuagagg.........................     11          0         seq
    ..................................................uuucgagaguggcuagagA.........................     1           1         seq
    ..................................................uuucgagaguggcuGgagg.........................     2           1         seq
    ..................................................uuucgagaguggcuagaggU........................     5           1         seq
    ..................................................uuucgagaguggcuagaggc........................     11          0         seq
    ..................................................uuucgagaguggcuagaggA........................     1           1         seq
    ..................................................uuucgagaguggcuagaggcA.......................     10          1         seq
    ..................................................Cuucgagaguggcuagaggcg.......................     2           1         seq
    ..................................................uuucUagaguggcuagaggcg.......................     1           1         seq
    ..................................................uuucgagaguggcuagaggcg.......................     35          0         seq
    ..................................................uuucgagaguggcuagaggcgA......................     10          1         seq
    ..................................................uuucgaAaguggcuagaggcgu......................     1           1         seq
    ..................................................uuucgagaguggcuagaggcgG......................     6           1         seq
    ..................................................uGucgagaguggcuagaggcgu......................     1           1         seq
    ..................................................uuucgagaguggcuagaggcgC......................     45          1         seq
    ..................................................uuuUgagaguggcuagaggcgu......................     2           1         seq
    ..................................................uuucgagGguggcuagaggcgu......................     3           1         seq
    ..................................................uuucgagagugUcuagaggcgu......................     3           1         seq
    ..................................................uuucgagaguggcuGgaggcgu......................     3           1         seq
    ..................................................Auucgagaguggcuagaggcgu......................     1           1         seq
    ..................................................uuucgagaguggcuagGggcgu......................     5           1         seq
    ..................................................uuuAgagaguggcuagaggcgu......................     1           1         seq
    ..................................................uAucgagaguggcuagaggcgu......................     3           1         seq
    ..................................................uuucgagaguggUuagaggcgu......................     1           1         seq
    ..................................................uuucgagaguggcuagaggcgu......................     275         0         seq
    ..................................................uuucgagaguggcuagagUcgu......................     1           1         seq
    ..................................................uuucgagagugAcuagaggcgu......................     2           1         seq
    ..................................................uuucAagaguggcuagaggcgu......................     1           1         seq
    ..................................................Cuucgagaguggcuagaggcgu......................     1           1         seq
    ..................................................uCucgagaguggcuagaggcgu......................     1           1         seq
    ..................................................uuCcgagaguggcuagaggcgu......................     3           1         seq
```

```
                                                 novel-nve-miR-34_guide
                 novel-nve-miR-34_star
uuagccugcaaaagcagaugccuuuagcuguuugcgaaacaacucguuauuucgagaguggcuagaggcgucugcuuucgcgggcuaaaacc

.................................................uuucgagaguggcuagaggUgu.....................   2     1     seq
.................................................uuucgagaguggcCagaggcgu.....................   1     1     seq
.................................................uuucgagaguggcuagaggcguA....................   2     1     seq
.................................................Cuucgagaguggcuagaggcguc....................   1     1     seq
.................................................uuAcgagaguggcuagaggcguc....................   1     1     seq
.................................................uuCcgagaguggcuagaggcguc....................   1     1     seq
.................................................uuucgagaguggcuagaggcguc....................   45    0     seq
.................................................uuucgagaguggcuagaggcguU....................   23    1     seq
.................................................uuucgagaguggUuagaggcguc....................   1     1     seq
.................................................uuucgagaguggcuagaggcgucA...................   61    1     seq
.................................................Cuucgagaguggcuagaggcgucu...................   2     1     seq
.................................................uuucgagaguggcuagaggcgucu...................   513   0     seq
.................................................uuucAagaguggcuagaggcgucu...................   1     1     seq
.................................................uCucgagaguggcuagaggcgucu...................   4     1     seq
.................................................uuucgagaguggcCagaggcgucu...................   2     1     seq
.................................................uuucgagaguggcuagaggcgucG...................   10    1     seq
.................................................uuucgUgaguggcuagaggcgucu...................   1     1     seq
.................................................uuucgagGguggcuagaggcgucu...................   5     1     seq
.................................................uuucgagaguAgcuagaggcgucu...................   1     1     seq
.................................................uuucgagaguggcuagGggcgucu...................   1     1     seq
.................................................Auucgagaguggcuagaggcgucu...................   7     1     seq
.................................................uuucgagaguggcuagaAgcgucu...................   3     1     seq
.................................................uuucgagaguggcuGgaggcgucu...................   1     1     seq
.................................................uuucgagaguggcGagaggcgucu...................   1     1     seq
.................................................uuucgagaguggcuCgaggcgucu...................   1     1     seq
.................................................uuucgagaguggcuagaggcgucC...................   84    1     seq
.................................................uAucgagaguggcuagaggcgucu...................   4     1     seq
.................................................uuucgagagAggcuagaggcgucu...................   1     1     seq
.................................................uuucgagaguggcuagCggcgucu...................   1     1     seq
.................................................uuCcgagaguggcuagaggcgucu...................   3     1     seq
.................................................uuucUagaguggcuagaggcgucu...................   1     1     seq
.................................................uuucgagaguggcuagaggcguUu...................   9     1     seq
.................................................uuucgagagCggcuagaggcgucu...................   5     1     seq
.................................................uuucgagaguggcuagaggcgCcu...................   3     1     seq
.................................................uuucgGgaguggcuagaggcgucu...................   8     1     seq
.................................................uuucgagaguggcuagaggcgucuU..................   4     1     seq
.................................................uuucgagaguggcuagaggcgucuUc.................   4     1     seq
..................................................uuUgagaguggcuagagg........................   1     1     seq
..................................................uucgagaguggcuagagA........................   3     1     seq
..................................................uucgagaguggcuGgagg........................   1     1     seq
..................................................uucgagaguggcuagagg........................   8     0     seq
..................................................uucgagaguggcuagaggU.......................   6     1     seq
..................................................uucgagaguggcuagaggc.......................   6     0     seq
..................................................uucgagaguggcuagaggG.......................   1     1     seq
..................................................Aucgagaguggcuagaggcg......................   1     1     seq
..................................................Cucgagaguggcuagaggcg......................   1     1     seq
..................................................uucgagaguggcuagaCgcg......................   1     1     seq
..................................................uucgagaguggcuagaggcU......................   2     1     seq
..................................................uucgagaguggUuagaggcg......................   1     1     seq
..................................................uucgagaguggcuagaggcA......................   6     1     seq
..................................................uucgagaguggcuagaggcg......................   36    0     seq
..................................................uucgaAaguggcuagaggcg......................   1     1     seq
..................................................uCcgagaguggcuagaggcgu.....................   1     1     seq
..................................................uucgaguAgcuagaggcgu.....................   2     1     seq
..................................................uucgagaguggcuagaggAgu.....................   1     1     seq
..................................................uucgagaCuggcuagaggcgu.....................   1     1     seq
..................................................uucgagaguggUuagaggcgu.....................   3     1     seq
..................................................uAcgagaguggcuagaggcgu.....................   1     1     seq
..................................................uucgagaguggcuagaggcgu.....................   219   0     seq
..................................................uucgGgaguggcuagaggcgu.....................   1     1     seq
..................................................uucgagaguggcuGgaggcgu.....................   4     1     seq
..................................................Aucgagaguggcuagaggcgu.....................   3     1     seq
..................................................uucgagaAuggcuagaggcgu.....................   1     1     seq
..................................................uuAgagaguggcuagaggcgu.....................   1     1     seq
..................................................uucgagaguggcuagaggcgG.....................   3     1     seq
..................................................uucgagaguggAuagaggcgu.....................   1     1     seq
..................................................uucgagaguggcuagaggcgC.....................   42    1     seq
..................................................uucgagGguggcuagaggcgu.....................   1     1     seq
..................................................uucgagaguggcuagaggcCu.....................   1     1     seq
```

```
                                                   novel-nve-miR-34_guide
                 novel-nve-miR-34_star
uuagccugcaaaagcagaugccuuuagcuguuugcgaaacaacucguuauuucgagaguggcuagaggcgucugcuuucgcgggcuaaaacc

..................................................uucgagaguggcuagaggcgA.....................    5      1      seq
..................................................uucgagagugAcuagaggcguc....................    1      1      seq
..................................................uucgagaguggcuagaggcAuc....................    1      1      seq
..................................................uucgagaguggcuagaggcguA....................    3      1      seq
..................................................uucgagaguggcuagaggcguc....................    38     0      seq
..................................................uucgGgaguggcuagaggcguc....................    2      1      seq
..................................................uucgagaguggcuagaggcguU....................    13     1      seq
..................................................uuUgagaguggcuagaggcguc....................    1      1      seq
..................................................uucgagaguggcuagaggUguc....................    1      1      seq
..................................................uucgagaguggcuagaggcgucG...................    9      1      seq
..................................................uucgagaguggcuagaggcgCcu...................    1      1      seq
..................................................uucgagaguggcuagUggcgucu...................    1      1      seq
..................................................uucgagaguggcuagaggcgGcu...................    1      1      seq
..................................................uucgagaguggcuUgaggcgucu...................    1      1      seq
..................................................uucgagaguggcCagaggcgucu...................    2      1      seq
..................................................uucgagaguggcuagaggcgucC...................    50     1      seq
..................................................uucgagaguggcuagaggcgucA...................    28     1      seq
..................................................uucgagaguggcuagaggcguUu...................    6      1      seq
..................................................uucgagaguggcuagGggcgucu...................    2      1      seq
..................................................uCcgagaguggcuagaggcgucu...................    1      1      seq
..................................................uucgagagAggcuagaggcgucu...................    1      1      seq
..................................................Cucgagaguggcuagaggcgucu...................    1      1      seq
..................................................uucgGgaguggcuagaggcgucu...................    1      1      seq
..................................................uucgagaguggcuagaggcgucu...................    281    0      seq
..................................................uucgagaguggcuCgaggcgucu...................    1      1      seq
..................................................Gucgagaguggcuagaggcgucu...................    2      1      seq
..................................................uucgagaguggcuagaggcgucuA..................    3      1      seq
...................................................uAgagaguggcuagaggcgu.....................    1      1      seq
...................................................uUgagaguggcuagaggcgu.....................    1      1      seq
...................................................uUgagaguggcuagaggcguc....................    1      1      seq
...................................................ucgagaguggcuagaggcgucA...................    1      1      seq
...................................................uUgagaguggcuagaggcgucu...................    1      1      seq
...................................................ucgagaguggcuagaggcgucC...................    1      1      seq
...................................................ucgagaguggcuagaggcgucu...................    5      0      seq
```

| | | |
|---|---|---|
| Provisional ID | | novel-nve-miR-35 |
| Score total | : | 6.1 |
| Score for star read(s) | : | -1.3 |
| Score for read counts | : | 0 |
| Score for mfe | : | 2.8 |
| Score for randfold | : | 1.6 |
| Score for cons. seed | : | 3 |
| Total read count | : | 535 |
| Mature read count | : | 535 |
| Loop read count | : | 0 |
| Star read count | : | 0 |

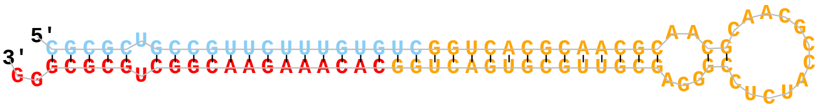


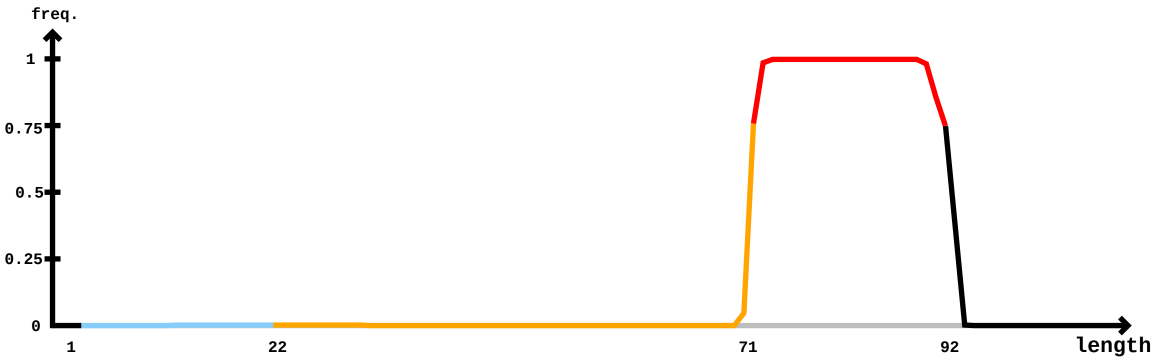


**Star** **Mature**

```
5'- ccgcgcugccguucuuugugucggucacgcaacgcaacgcaacgccaucucccgggagcguugcgugacuggcacaaagaacggcugcgcgggagacuacacaggggucgg   -3'    exp
    (((((.(((((((((((((((((((((((((((((..((............))...))))))))))))))))))))))))))))).)))))..((((.......))))..   reads     mm        sample
    ...........uucuuugugucggucacgca...............................................................................   1         0         seq
    ......................................................................gcacaaagaacggcugcgcU....................   1         1         seq
    ......................................................................gcacaaagaacggcugcgcg....................   4         0         seq
    ......................................................................gcacaaagaacggcugcgcA....................   1         1         seq
    ......................................................................Ccacaaagaacggcugcgcgg...................   1         1         seq
    ......................................................................Ucacaaagaacggcugcgcgg...................   1         1         seq
    ......................................................................gcacaaagaacggcugcgcgA...................   1         1         seq
    ......................................................................gcacaaagaacggcugcgcgg...................   1         0         seq
    ......................................................................Ccacaaagaacggcugcgcggg..................   1         1         seq
    ......................................................................gcacaaagaacggcugcgcggg..................   8         0         seq
    ......................................................................Ucacaaagaacggcugcgcggg..................   2         1         seq
    ......................................................................gcacaaagaacggcugcgcgCg..................   1         1         seq
    ......................................................................gcacaaagaacggcugcgcggA..................   1         1         seq
    ......................................................................gcaUaaagaacggcugcgcggg..................   1         1         seq
    ......................................................................Acacaaagaacggcugcgcgggag................   1         1         seq
    .......................................................................cacaaagaacggcugcgU.....................   1         1         seq
    .......................................................................cacaaagaacggcugcgc.....................   8         0         seq
    .......................................................................cacaaagaacggcugUgcg....................   2         1         seq
    .......................................................................cacaaagaacggcugcgcC....................   1         1         seq
    .......................................................................cacaaGgaacggcugcgcg....................   1         1         seq
    .......................................................................cacaaagaacggcugcgcA....................   4         1         seq
    .......................................................................cacaaagaCcggcugcgcg....................   1         1         seq
    .......................................................................cacaaagaacggcCgcgcg....................   1         1         seq
    .......................................................................cacaaagaacggcugcgcg....................   42        0         seq
    .......................................................................cacaaagCacggcugcgcgg...................   1         1         seq
    .......................................................................cacaaagaacggcugcgcgg...................   34        0         seq
    .......................................................................cacaaagaacggcugUgcgg...................   1         1         seq
    .......................................................................cacaaagaacggcugcgcgA...................   5         1         seq
    .......................................................................cacaaagaacggUugcgcggg..................   2         1         seq
    .......................................................................cacaaagaacggcugcgcggA..................   19        1         seq
    .......................................................................cacaaagaacggcugcgcgUg..................   1         1         seq
    .......................................................................cacaaagaacggcugUgcggg..................   1         1         seq
    .......................................................................cacaaagaUcggcugcgcggg..................   1         1         seq
    .......................................................................cacaaagaacggcugcgcggC..................   4         1         seq
```

```
 Star                                                                  Mature
ccgcgcugccguucuuugugucggucacgcaacgcaacgcaacgccaucuccgggagcguugcgugacuggcacaaagaacggcugcgcgggagacuacacaggggucgg
.......................................................................cUcaaagaacggcugcgcggg..................  2    1    seq
.......................................................................cacGaagaacggcugcgcggg..................  2    1    seq
.......................................................................cacaaagaCcggcugcgcggg..................  1    1    seq
.......................................................................caGaaagaacggcugcgcggg..................  1    1    seq
.......................................................................cGcaaagaacggcugcgcggg..................  1    1    seq
.......................................................................cacaaagaacggcugcgcggU..................  6    1    seq
.......................................................................cacaaagaacggcugcgcAgg..................  1    1    seq
.......................................................................cacaaagaacggcugcgcggg..................  205  0    seq
.......................................................................cacaaagaacgAcugcgcggg..................  1    1    seq
.......................................................................cacaaagaacggcugcCcggg..................  1    1    seq
.......................................................................cacaaagaacUgcugcgcggg..................  1    1    seq
.......................................................................cacaaagaacggcuCcgcggg..................  1    1    seq
.......................................................................Aacaaagaacggcugcgcggg..................  1    1    seq
.......................................................................cacaaagaacggcuAcgcggga.................  1    1    seq
.......................................................................cacaaagaacggcugcgcgggC.................  3    1    seq
.......................................................................cacaaagaacggcugcgcgggU.................  10   1    seq
.......................................................................cacaaagaacggcugcgcgggG.................  1    1    seq
.......................................................................cacaaagaacggcugcgcggga.................  11   0    seq
.......................................................................cacaaagaGcggcugcgcggga.................  1    1    seq
........................................................................acaaagaacggcugcgcg....................  6    0    seq
........................................................................acaaagaacggcugcgcC....................  1    1    seq
........................................................................acaaagaacggcugcgcA....................  2    1    seq
........................................................................acaaagaacggcugcgcgg...................  9    0    seq
........................................................................aUaaagaacggcugcgcgg...................  1    1    seq
........................................................................acaaagaacggcugcgcgA...................  2    1    seq
........................................................................acaaagaacUgcugcgcgg...................  1    1    seq
........................................................................acaCagaacggcugcgcggg..................  1    1    seq
........................................................................acaaagaacggcugcgcggA..................  1    1    seq
........................................................................acaaagaacggcugcgcggg..................  17   0    seq
........................................................................acGaagaacggcugcgcggg..................  1    1    seq
........................................................................acaaagaaAggcugcgcggga.................  1    1    seq
........................................................................acaaagaacggcugcgcggga.................  20   0    seq
........................................................................acaaagaacggcugcgcgggU.................  50   1    seq
........................................................................Ccaaagaacggcugcgcggga.................  1    1    seq
........................................................................acaaagaacggcugcgcgggC.................  8    1    seq
.........................................................................caaagaacggcugcgcggg..................  2    0    seq
.........................................................................caaagaacggcugcgcggU..................  1    1    seq
.........................................................................caaagaacggcugcgcgggG.................  1    1    seq
.........................................................................caaagaacggcugcgcgggU.................  3    1    seq
```

miRBase precursor : novel-nve-miR-36-1
Total read count : 2592
novel-nve-miR-36-1_guide read count : 2592
novel-nve-miR-36-1_star read count : 0
remaining reads : 0

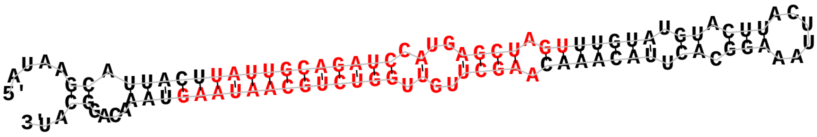

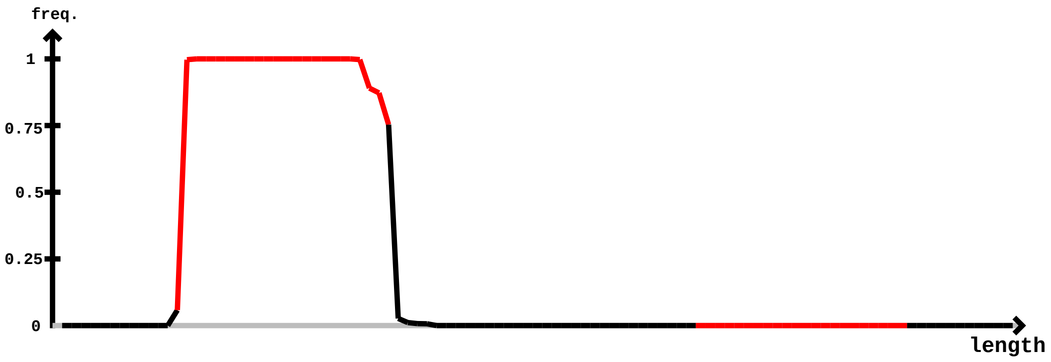


```
                  novel-nve-miR-36-1_guide
                                                                          novel-nve-miR-36-1_star
5'- auaagcauuacuuauugcagauccaugagcuaguuuguauguacuuacuuaaaggcacuuacaaacaagcuuguuggucugcaauaaguaaacaggcau  -3'   exp
    ....((.((((((((((((((((.(..((((.(((((((.((.(((......))).)).))))))).))))..).))))))))))))))))....))..    reads   mm   sample
    ...........uuauugcagUuccaugagcu....................................................................    1       1    seq
    ...........uuauugcagauccaugagcC....................................................................    4       1    seq
    ...........uuauugcagauccaugagcu....................................................................    6       0    seq
    ...........uuauugcagauccaugagcua...................................................................    5       0    seq
    ...........uuauugcagauccaugagcuaA..................................................................    3       1    seq
    ...........uuauugcagauccaugagcuag..................................................................    8       0    seq
    ...........uuauugcagauccaCgagcuag..................................................................    1       1    seq
    ...........uuauugcagauccaugagcuGg..................................................................    1       1    seq
    ...........uuauugcagauccCugagcuagu.................................................................    1       1    seq
    ...........uuauugcagauccaugaCcuagu.................................................................    1       1    seq
    ...........Cuauugcagauccaugagcuagu.................................................................    1       1    seq
    ...........Auauugcagauccaugagcuagu.................................................................    1       1    seq
    ...........uuauugcagauccaugagcuagu.................................................................    84      0    seq
    ...........uuauugcagauccaugagcuagC.................................................................    16      1    seq
    ...........uuauugcagauccaCgagcuagu.................................................................    1       1    seq
    ...........uuauuAcagauccaugagcuagu.................................................................    1       1    seq
    ...........uuauugcagauccaugagcuGgu.................................................................    2       1    seq
    ...........uuauugcagauccaugagcuagA.................................................................    1       1    seq
    ...........uuauGgcagauccaugagcuagu.................................................................    1       1    seq
    ...........uuauugcagauccaugaAcuagu.................................................................    1       1    seq
    ...........uuauugUagauccaugagcuagu.................................................................    1       1    seq
    ...........uuauugcagauccaugagcuaguu................................................................    4       0    seq
    ...........uuauugcagauccaugagcuaguuu...............................................................    2       0    seq
    ...........uuauugcagauccaugagcuaguuA...............................................................    2       1    seq
    ...........uuauugcagauccaugagcuaguuuAu.............................................................    1       1    seq
    ............uauugcagauccaugagc.....................................................................    5       0    seq
    ............uauugcagauccaugagU.....................................................................    2       1    seq
    ............uauugcagaucAaugagcu....................................................................    1       1    seq
    ............Aauugcagauccaugagcu....................................................................    3       1    seq
    ............uauugcagauccGugagcu....................................................................    1       1    seq
    ............uauugcagauccaCgagcu....................................................................    3       1    seq
    ............uaCugcagauccaugagcu....................................................................    2       1    seq
    ............uauugcagauccaugagcG....................................................................    4       1    seq
    ............uauCgcagauccaugagcu....................................................................    2       1    seq
```

```
            novel-nve-miR-36-1_guide
                                                                  novel-nve-miR-36-1_star
auaagcauuacuuauugcagauccaugagcuaguuuguauguacuuacuuaaaggcacuuacaaacaagcuuguuggucugcaauaaguaaacaggcau

............uauugcGgauccaugagcu....................................................................    3      1      seq
............uauugcagauccaugagcA....................................................................    14     1      seq
............uauugcagauccaugagcu....................................................................    201    0      seq
............uauugcagauccaugagcC....................................................................    20     1      seq
............uauugcagauccaugGgcu....................................................................    1      1      seq
............uauugcaUauccaugagcu....................................................................    1      1      seq
............uaAugcagauccaugagcu....................................................................    1      1      seq
............uauugcagGuccaugagcu....................................................................    2      1      seq
............uauugUagauccaugagcu....................................................................    3      1      seq
............uauugcagaucUaugagcu....................................................................    4      1      seq
............uauugcagauccaugagcGa...................................................................    1      1      seq
............uauugcagGuccaugagcua...................................................................    1      1      seq
............uauugcagauccaugagcuG...................................................................    1      1      seq
............Aauugcagauccaugagcua...................................................................    1      1      seq
............uauugcagaAccaugagcua...................................................................    1      1      seq
............uauugcagauccaugagcua...................................................................    35     0      seq
............uauugcagauccaugagUua...................................................................    1      1      seq
............uauugcagauccauUagcua...................................................................    1      1      seq
............Aauugcagauccaugagcuag..................................................................    3      1      seq
............uauugcagauccaugagcuaC..................................................................    1      1      seq
............uauugcagauccaugagcuaU..................................................................    4      1      seq
............uauugcUgauccaugagcuag..................................................................    1      1      seq
............uauugcagauccaugGgcuag..................................................................    1      1      seq
............uauugcagauccauAagcuag..................................................................    2      1      seq
............uaAugcagauccaugagcuag..................................................................    1      1      seq
............uauugcagauAcaugagcuag..................................................................    1      1      seq
............uauugcagauccaugagcGag..................................................................    1      1      seq
............uauugcagauccaugagcuGg..................................................................    4      1      seq
............uauugcagauccaugagcuaA..................................................................    20     1      seq
............uauugcagauccauUagcuag..................................................................    1      1      seq
............uauAgcagauccaugagcuag..................................................................    2      1      seq
............uauugcagGuccaugagcuag..................................................................    2      1      seq
............uauugcGgauccaugagcuag..................................................................    1      1      seq
............Cauugcagauccaugagcuag..................................................................    3      1      seq
............uauugcagauccaugagcuag..................................................................    240    0      seq
............uauuAcagauccaugagcuag..................................................................    2      1      seq
............uauugcagaucAaugagcuag..................................................................    1      1      seq
............uauugAagauccaugagcuag..................................................................    1      1      seq
............uauugcagauccGugagcuag..................................................................    2      1      seq
............uauugcagaucAaugagcuagu.................................................................    4      1      seq
............uauugcagauccaugagcuagu.................................................................    1383   0      seq
............uauugcagauccaGgagcuagu.................................................................    1      1      seq
............uauugcagauccGugagcuagu.................................................................    4      1      seq
............uauugcagauccaugGgcuagu.................................................................    13     1      seq
............uauugcagauccaugagcuaCu.................................................................    1      1      seq
............uauugcGgauccaugagcuagu.................................................................    14     1      seq
............uauugcagauccaugagcuUgu.................................................................    2      1      seq
............uauugcagauccaugCgcuagu.................................................................    4      1      seq
............uauugcagauccUugagcuagu.................................................................    1      1      seq
............uaGugcagauccaugagcuagu.................................................................    1      1      seq
............uauugcagauccaugagcuCgu.................................................................    1      1      seq
............uauugcagGuccaugagcuagu.................................................................    8      1      seq
............uauugcagaCccaugagcuagu.................................................................    13     1      seq
............uauugcagauccaugagcuagG.................................................................    17     1      seq
............uauugAagauccaugagcuagu.................................................................    1      1      seq
............uauugcagauccaugagUuagu.................................................................    6      1      seq
............uaCugcagauccaugagcuagu.................................................................    4      1      seq
............uauugcagaucUaugagcuagu.................................................................    8      1      seq
............uauugcagauccaugagcuGgu.................................................................    13     1      seq
............uauGgcagauccaugagcuagu.................................................................    1      1      seq
............uauugcagauccauAagcuagu.................................................................    3      1      seq
............uauugcagauUcaugagcuagu.................................................................    3      1      seq
............uauugcagauccaugagAuagu.................................................................    1      1      seq
............uauugcagauccaugagcuagC.................................................................    163    1      seq
............Cauugcagauccaugagcuagu.................................................................    4      1      seq
............uauugcagauccaugUgcuagu.................................................................    1      1      seq
............uauuCcagauccaugagcuagu.................................................................    1      1      seq
............uauugcaAauccaugagcuagu.................................................................    1      1      seq
............uauuAcagauccaugagcuagu.................................................................    2      1      seq
```

```
            novel-nve-miR-36-1_guide
                                                       novel-nve-miR-36-1_star
auaagcauuacuuauugcagauccaugagcuaguuuguauguacuuacuuaaaggcacuuacaaacaagcuuguuggucugcaauaaguaaacaggcau

............uauCgcagauccaugagcuagu.................................................................   5    1    seq
............uauugcagauccaAgagcuagu.................................................................   1    1    seq
............uauugcaUauccaugagcuagu.................................................................   1    1    seq
............uauugcagauccaugagcCagu.................................................................   3    1    seq
............uauugcagauccaugagcuagA.................................................................   28   1    seq
............Gauugcagauccaugagcuagu.................................................................   3    1    seq
............uauAgcagauccaugagcuagu.................................................................   2    1    seq
............uauugUagauccaugagcuagu.................................................................   12   1    seq
............uaAugcagauccaugagcuagu.................................................................   4    1    seq
............uauugcagauccaugagcGagu.................................................................   1    1    seq
............uauugcUgauccaugagcuagu.................................................................   2    1    seq
............uGuugcagauccaugagcuagu.................................................................   2    1    seq
............uauugcagauccaCgagcuagu.................................................................   6    1    seq
............uauugcagauAcaugagcuagu.................................................................   1    1    seq
............Aauugcagauccaugagcuagu.................................................................   17   1    seq
............uauugcagauccaugagcuaguu................................................................   19   0    seq
............uauugcagauccaugagcuaguG................................................................   1    1    seq
............uauugcagauccaugagcuaguA................................................................   7    1    seq
............uauugcagauccaugagcuaguC................................................................   3    1    seq
............uauugcagauccGugagcuaguu................................................................   1    1    seq
............uauugcagauAcaugagcuaguu................................................................   1    1    seq
............uauugUagauccaugagcuaguu................................................................   1    1    seq
............uauugcagauccaugagcuGguu................................................................   1    1    seq
............uauugcagaucUaugagcuaguuu...............................................................   4    1    seq
............uauugcagauccaugagcuaguuu...............................................................   2    0    seq
............uauugcagauccaugagcuaguuuU..............................................................   2    1    seq
............uauugcagauccaugagcuaguuuAu.............................................................   2    1    seq
............uauugcagauccaugagcuaguuuUu.............................................................   14   1    seq
.............auugcagauccaugagcuag..................................................................   1    0    seq
.............auugcagauccaugagcuagu.................................................................   5    0    seq
.............auugcagauccaugagcuaguA................................................................   2    1    seq
```

miRBase precursor : novel-nve-miR-38
Total read count : 720
novel-nve-miR-38_guide read count : 720
novel-nve-miR-38_star read count : 0
remaining reads : 0

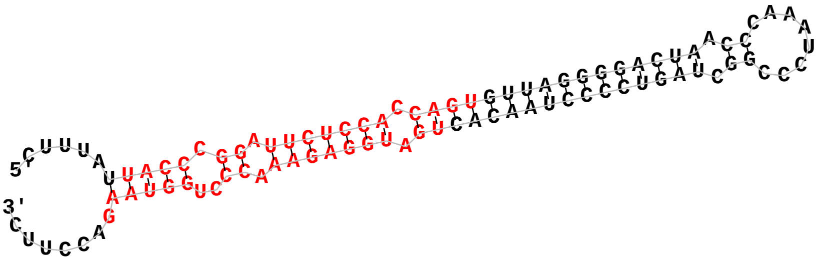

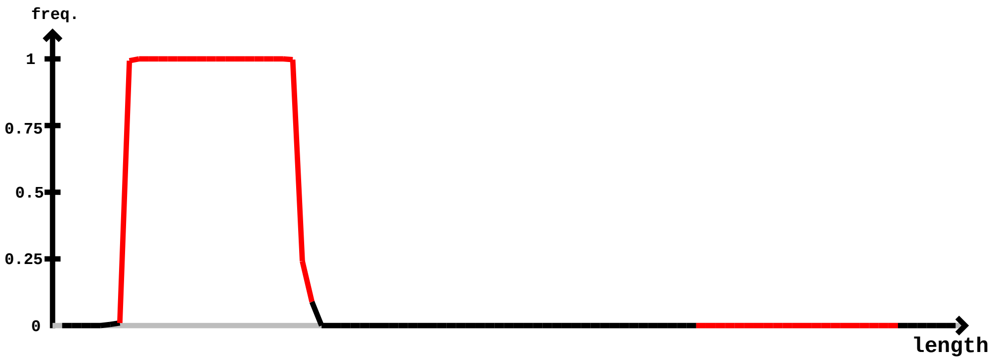


```
                                                                        novel-nve-miR-38_star
        novel-nve-miR-38_guide
5'- cuuuauuacccggauucuccaccaguguuaggggacuaacccaaaucccggcuaguccccuaacacugauggagaaacccugguaagaccuuc   -3'    exp
    .....(((((.((.(((((((.((((((((((((((((.((........)).)))))))))))))))).))))))).))..))))).......    reads      mm        sample
    ....Guuacccggauucuccacc......................................................................    1          1         seq
    ....Guuacccggauucuccacca.....................................................................    2          1         seq
    .....uuacccUgauucuccacc......................................................................    1          1         seq
    .....uuacccggauucuccacca.....................................................................    2          0         seq
    .....Guacccggauucuccacca.....................................................................    1          1         seq
    ......uacccggauucuccaUca.....................................................................    1          1         seq
    ......uacccggauucuccacca.....................................................................    163        0         seq
    ......uacccggauuUuccacca.....................................................................    2          1         seq
    ......Aacccggauucuccacca.....................................................................    1          1         seq
    ......uCcccggauucuccacca.....................................................................    7          1         seq
    ......uacccggauucuccaccU.....................................................................    26         1         seq
    ......uaUccggauucuccacca.....................................................................    4          1         seq
    ......uacccggauucuccacAa.....................................................................    1          1         seq
    ......uacAcggauucuccacca.....................................................................    1          1         seq
    ......uacccggauucuccGcca.....................................................................    1          1         seq
    ......uacccggauucuccaccC.....................................................................    7          1         seq
    ......Cacccggauucuccacca.....................................................................    2          1         seq
    ......uacccggaCucuccacca.....................................................................    2          1         seq
    ......uacccggauucuccaccG.....................................................................    245        1         seq
    ......uUcccggauucuccacca.....................................................................    55         1         seq
    ......uGcccggauucuccacca.....................................................................    1          1         seq
    ......uacccggGuucuccacca.....................................................................    16         1         seq
    ......uacccggauucCccacca.....................................................................    1          1         seq
    ......uacccggauCcuccacca.....................................................................    1          1         seq
    ......uacccggauucuAcacca.....................................................................    1          1         seq
    ......uacccggauucuccUcca.....................................................................    1          1         seq
    ......uaccUggauucuccaccag....................................................................    1          1         seq
    ......uacccggGuucuccaccag....................................................................    1          1         seq
    ......uCcccggauucuccaccag....................................................................    1          1         seq
    ......uacccggauucuccaccaC....................................................................    14         1         seq
    ......uacccggauucuccaccGg....................................................................    1          1         seq
    ......uacccggauuUuccaccag....................................................................    1          1         seq
    ......uacccggauucuccaccag....................................................................    13         0         seq
    ......uacccggauucuccacUag....................................................................    1          1         seq
```

```
                                                                  novel-nve-miR-38_star
      novel-nve-miR-38_guide
cuuuauuacccggauucuccaccaguguuaggggacuaacccaaaucccggcuaguccccuaacacugauggagaaacccugguaagaccuuc

......uacccggauucuccaccCg.....................................................................    1    1    seq
......uacccggauucuccaccaA....................................................................    7    1    seq
......uUcccggauucuccaccag....................................................................    12   1    seq
......uacccggauucuccaccaU....................................................................    48   1    seq
......uacccggauucuccaccUg....................................................................    8    1    seq
......uacccggauucucUaccag....................................................................    1    1    seq
......uacccggauucuccaccaUu...................................................................    52   1    seq
......uacccggauucuccaccaCu...................................................................    7    1    seq
.......acccggauucuccaccaCu...................................................................    1    1    seq
.......acccggauucuccaccaUu...................................................................    4    1    seq
```

| | |
|---|---|
| miRBase precursor | : novel-nve-miR-39-1 |
| Total read count | : 1113 |
| novel-nve-miR-39-1_guide read count | : 1112 |
| novel-nve-miR-39-1_star read count | : 0 |
| remaining reads | : 1 |

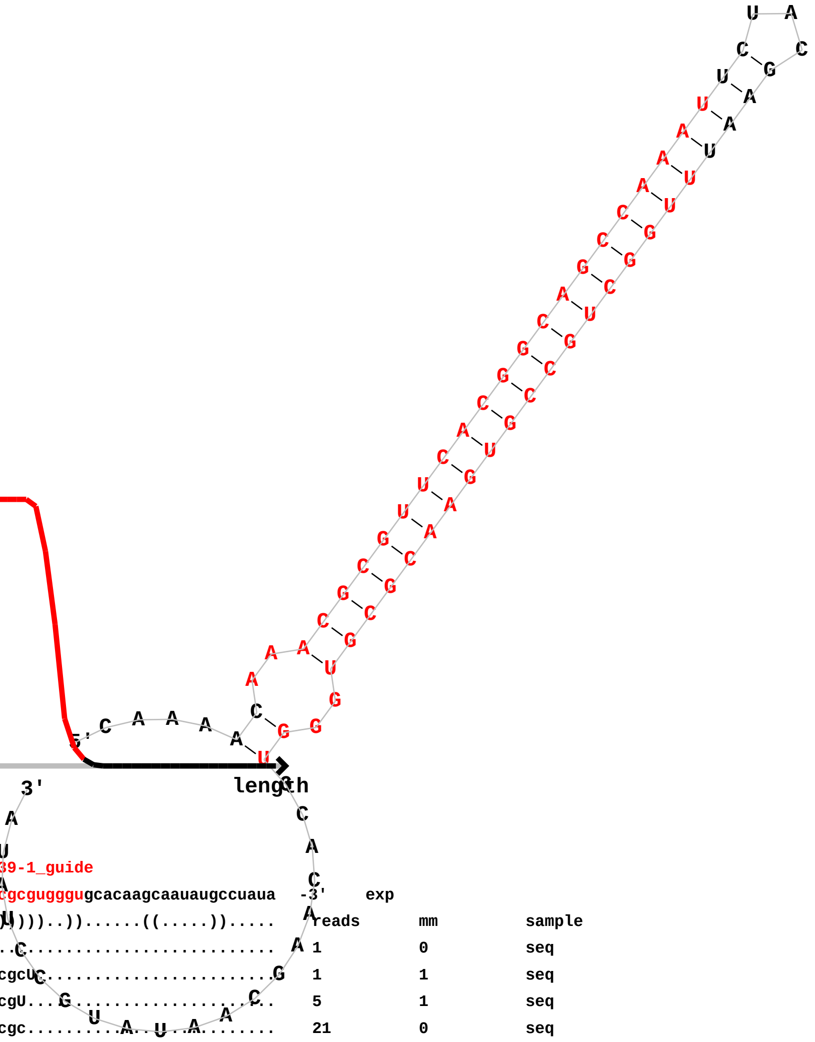


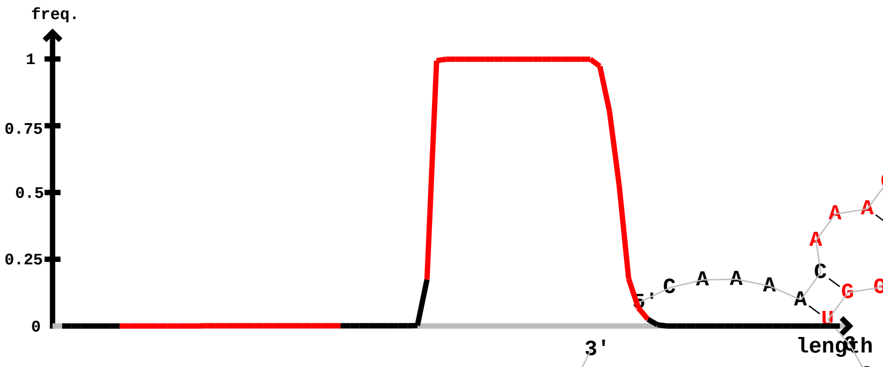


novel-nve-miR-39-1_star

novel-nve-miR-39-1_guide

| 5'- caaaacaaacgcguucacggcagccaaauucuacgaauuuggcugccgugaacgcgugggugcacaagcaauaugccuaua -3' | exp | | |
|---|---|---|---|
| ....((..(((((((((((((((((((((((...)))))))))))))))))))))))..))......((.....))..... | reads | mm | sample |
| ..............ucacggcagccaaauucuacgaau........................................... | 1 | 0 | seq |
| ....................................auuuggcugccgugaacgcU......................... | 1 | 1 | seq |
| .....................................uuuggcugccgugaacgU.......................... | 5 | 1 | seq |
| .....................................uuuggcugccgugaacgc.......................... | 21 | 0 | seq |
| .....................................uuuggcugccgAgaacgc.......................... | 1 | 1 | seq |
| .....................................uuuggcugccgugaacgA.......................... | 2 | 1 | seq |
| .....................................uuugCcugccgugaacgcg......................... | 1 | 1 | seq |
| .....................................uuuggcugccgugaacgcA......................... | 7 | 1 | seq |
| .....................................uCuggcugccgugaacgcg......................... | 2 | 1 | seq |
| .....................................uuuggcugccgugaacgcg......................... | 39 | 0 | seq |
| .....................................uuuggcugcUgugaacgcg......................... | 1 | 1 | seq |
| .....................................uuuggcugccgugaacgcU......................... | 1 | 1 | seq |
| .....................................uuuggcugccgugaacgcgu........................ | 50 | 0 | seq |
| .....................................uuuggcugccgugGacgcgu........................ | 1 | 1 | seq |
| .....................................Auuggcugccgugaacgcgu........................ | 1 | 1 | seq |
| .....................................uuuggcugccgugaacgcgA........................ | 1 | 1 | seq |
| .....................................uuuggcugccgugaacgcgG........................ | 1 | 1 | seq |
| .....................................uuuggcugccgugaacgcgC........................ | 9 | 1 | seq |
| .....................................uuuggcugccgugaGcgcgu........................ | 1 | 1 | seq |
| .....................................uuuggAugccgugaacgcgu........................ | 1 | 1 | seq |
| .....................................uuuggcugccgugaacgcgug....................... | 10 | 0 | seq |
| .....................................uuuggcugccgugaacgcguC....................... | 2 | 1 | seq |
| .....................................uuuggcugccgugaacgcguU....................... | 21 | 1 | seq |
| .....................................uuuggcugccgugaacgcguA....................... | 3 | 1 | seq |
| .....................................uuuggcugccgugaacgcgCg....................... | 1 | 1 | seq |
| .....................................uuuggcugccgugaacgcgugU...................... | 4 | 1 | seq |
| .....................................uuuggcugccgugaacgcgugg...................... | 3 | 0 | seq |
| .....................................uuuggcugccgugaacgcgugA...................... | 1 | 1 | seq |
| .....................................uuuggcugccgugaacgcgugggC.................... | 1 | 1 | seq |
| .....................................uuuggcugccgugaacgcgugggu.................... | 1 | 0 | seq |
| ......................................Cuggcugccgugaacgcg......................... | 1 | 1 | seq |
| ......................................uuggcugccgugaCcgcg......................... | 1 | 1 | seq |
| ......................................uuggcugccgugaacgcU......................... | 4 | 1 | seq |
| ......................................uuggcugccgugaacgcA......................... | 15 | 1 | seq |

novel-nve-miR-39-1_star
novel-nve-miR-39-1_guide

```
caaaacaaacgcguucacggcagccaaauucuacgaauuuggcugccgugaacgcgugggugcacaagcaauaugccuaua

......................................uuggcugUcgugaacgcg.........................   1     1   seq
......................................uuggcugccgCgaacgcg.........................   1     1   seq
......................................uuggcugccgugaGcgcg.........................   2     1   seq
......................................uuggcugccgugaacgcg.........................   102   0   seq
......................................uuggcugccgugaUcgcg.........................   1     1   seq
......................................uuggcugccgugaacgcC.........................   3     1   seq
......................................uuggcCgccgugaacgcg.........................   1     1   seq
......................................Auggcugccgugaacgcg.........................   1     1   seq
......................................uuggcugcAgugaacgcg.........................   1     1   seq
......................................uuUgcugccgugaacgcg.........................   1     1   seq
......................................uuggcugcUgugaacgcgu........................   1     1   seq
......................................uuggcAgccgugaacgcgu........................   1     1   seq
......................................uuggAugccgugaacgcgu........................   1     1   seq
......................................uuggcugccgugaGcgcgu........................   2     1   seq
......................................uuAgcugccgugaacgcgu........................   1     1   seq
......................................uuggcugccgugaacgcgC........................   30    1   seq
......................................uuggcCgccgugaacgcgu........................   1     1   seq
......................................uuggcugccgugaacgGgu........................   2     1   seq
......................................uuggcugccgugaacgcgu........................   181   0   seq
......................................uCggcugccgugaacgcgu........................   1     1   seq
......................................Auggcugccgugaacgcgu........................   1     1   seq
......................................uuggcugccgugaacAcgu........................   1     1   seq
......................................uuggcugccgugaacgcgA........................   11    1   seq
......................................uuggcugccgugaacgcgG........................   5     1   seq
......................................uuggcugccgugaacgcgug.......................   262   0   seq
......................................uuggcAgccgugaacgcgug.......................   2     1   seq
......................................uuggcugccgugaacgcguA.......................   28    1   seq
......................................uuggUugccgugaacgcgug.......................   1     1   seq
......................................uuggAugccgugaacgcgug.......................   2     1   seq
......................................uuggcugccgugaacgcgCg.......................   3     1   seq
......................................uugUcugccgugaacgcgug.......................   1     1   seq
......................................uuggcCgccgugaacgcgug.......................   1     1   seq
......................................uAggcugccgugaacgcgug.......................   1     1   seq
......................................uuggcugccgugaaUgcgug.......................   1     1   seq
......................................uuggcugccgugaacgcguU.......................   36    1   seq
......................................uuggcugccguUaacgcgug.......................   2     1   seq
......................................uuggcugccgCgaacgcgug.......................   4     1   seq
......................................uuggcugccgGgaacgcgug.......................   1     1   seq
......................................uuggcugccgugaacgcgGg.......................   2     1   seq
......................................uuggcugccgugaacgcguC.......................   2     1   seq
......................................uuggcugccgugaGcgcgug.......................   1     1   seq
......................................Guggcugccgugaacgcgug.......................   2     1   seq
......................................Cuggcugccgugaacgcgug.......................   1     1   seq
......................................uuggcugccguCaacgcgug.......................   1     1   seq
......................................Auggcugccgugaacgcgug.......................   2     1   seq
......................................uuggcugccgugaaUgcgugg......................   1     1   seq
......................................uuggcugcAgugaacgcgugg......................   1     1   seq
......................................uuggcugccgugaacgcgugg......................   71    0   seq
......................................uuggcugccgugaacgcgugC......................   6     1   seq
......................................uuggcugcUgugaacgcgugg......................   1     1   seq
......................................uuggcugccgugaacgcgugA......................   13    1   seq
......................................uuggcCgccgugaacgcgugg......................   1     1   seq
......................................uuggcugccgugaacgcgugU......................   15    1   seq
......................................uuggcugccgugaacgUgugg......................   1     1   seq
......................................uuggcugccgugaacgcgugCg.....................   1     1   seq
......................................uuggcugccgugaacgcguggU.....................   15    1   seq
......................................uuggcugccgugaacgcguggg.....................   24    0   seq
......................................uuggcugcAgugaacgcguggg.....................   1     1   seq
......................................uuggcugccgugaacgcguggA.....................   4     1   seq
......................................uuggcugccgugaacgcguggC.....................   2     1   seq
......................................uuggcugccgugaacgcgugggC....................   6     1   seq
......................................uuggcugcUgugaacgcgugggu....................   1     1   seq
......................................uuggcugccgugaacgcgugggA....................   2     1   seq
......................................uuggcugccgugaacgcgugggu....................   12    0   seq
......................................uuggcugccgugaacgcgugggCg...................   1     1   seq
......................................uuggcugccgugaacgcguggguU...................   2     1   seq
......................................uuggcugccgugaacgcgugggug...................   1     0   seq
......................................uuggcugccgugaacgcguggguA...................   1     1   seq
.......................................uggcugccgugaacgcgA........................   1     1   seq
```

```
      novel-nve-miR-39-1_star
                                      novel-nve-miR-39-1_guide
caaaacaaacgcguucacggcagccaaauucuacgaauuuggcugccgugaacgcgugggugcacaagcaauaugccuaua

.......................................uggcugccgugaacgcguU.......................    1    1    seq
.......................................uggcugccgugaacgcgugg......................    2    0    seq
.......................................uggcugccgugaacgcgugA......................    1    1    seq
.......................................uggcugccgugaacgcguggC.....................    1    1    seq
```

miRBase precursor : novel-nve-miR-44-1
Total read count : 2038
novel-nve-miR-44-1_guide read count : 2038
novel-nve-miR-44-1_star read count : 0
remaining reads : 0

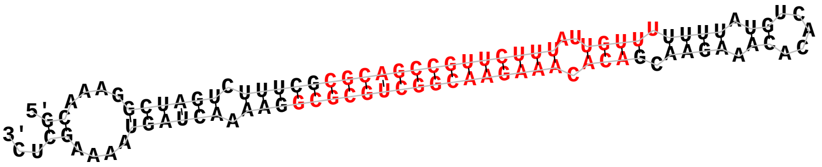

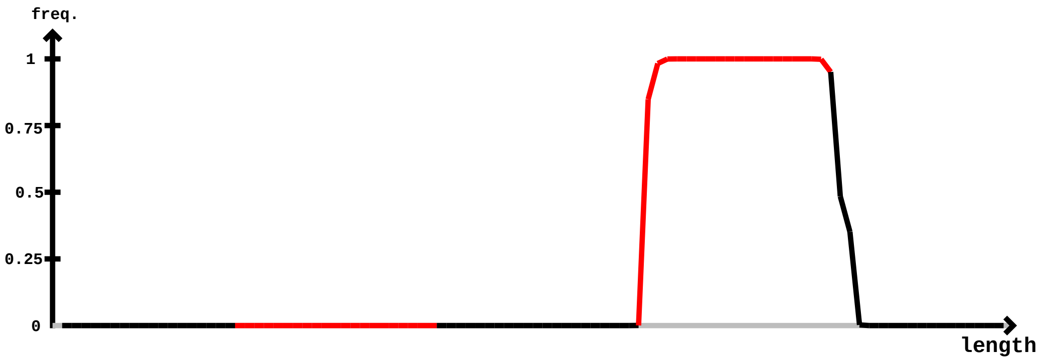


```
                                   novel-nve-miR-44-1_star
                                                                         novel-nve-miR-44-1_guide
5'- gcaaaggcuagucuuucgcgcagccguucuuuauuguuuuuuuaugucacacaaagaacgacacaaagaacggcugcgcggaaaacuaguaaaagcuc  -3'   exp
    ((....((((((.((((((((((((((((((..((((.((((.((.....)).)))).)))).))))))))))))))))))).)))))).....))..   reads   mm   sample
    ...........................................................Cacacaaagaacggcugcgcg..................   1       1    seq
    ............................................................acacaaagaacggcugcU....................   1       1    seq
    ............................................................acacaaagaacggcugcg....................   2       0    seq
    ............................................................acacaaagaacggcugcUc...................   1       1    seq
    ............................................................acacaaagaacggcugcgU...................   22      1    seq
    ............................................................acacaaUgaacggcugcgc...................   1       1    seq
    ............................................................acacaaagaacggcugcgG...................   2       1    seq
    ............................................................acacaaagaacggcugcgc...................   55      0    seq
    ............................................................Ucacaaagaacggcugcgc...................   1       1    seq
    ............................................................acacGaagaacggcugcgc...................   1       1    seq
    ............................................................acacaaagaacggcugcgA...................   3       1    seq
    ............................................................acacaaagCacggcugcgcg..................   2       1    seq
    ............................................................acacaaagaGcggcugcgcg..................   1       1    seq
    ............................................................aUacaaagaacggcugcgcg..................   1       1    seq
    ............................................................acacaaagaacggcugcgcg..................   575     0    seq
    ............................................................acacaaagaacggcugUgcg..................   1       1    seq
    ............................................................acacaaaCaacggcugcgcg..................   1       1    seq
    ............................................................acacaaagaacggUugcgcg..................   1       1    seq
    ............................................................acacaaagaCcggcugcgcg..................   1       1    seq
    ............................................................acacaaagaaAggcugcgcg..................   1       1    seq
    ............................................................acGcaaagaacggcugcgcg..................   1       1    seq
    ............................................................acacaaagaacggcugcgcC..................   11      1    seq
    ............................................................acacaaagaacggAugcgcg..................   1       1    seq
    ............................................................acacaaagaaUggcugcgcg..................   2       1    seq
    ............................................................acUcaaagaacggcugcgcg..................   1       1    seq
    ............................................................acaAaaagaacggcugcgcg..................   1       1    seq
    ............................................................Ccacaaagaacggcugcgcg..................   3       1    seq
    ............................................................acacaaagaacggcuAcgcg..................   1       1    seq
    ............................................................acacaaagaacggcugAgcg..................   1       1    seq
    ............................................................acacaaagaacggcugcgcA..................   147     1    seq
    ............................................................acacaaagaacgCcugcgcg..................   1       1    seq
    ............................................................acacaaagaacggcAgcgcg..................   1       1    seq
    ............................................................acacaaagaacgUcugcgcg..................   1       1    seq
    ............................................................acacaaagaacggcugcgcU..................   34      1    seq
```

```
                  novel-nve-miR-44-1_star
                                                             novel-nve-miR-44-1_guide
gcaaaggcuagucuuucgcgcagccguucuuuauuguuuuuuuaugucacacaaagaacgacacaaagaacggcugcgcggaaaacuaguaaaagcuc

............................................................aAacaaagaacggcugcgcg..................   1   1   seq
............................................................acacaaagaacggcugcgAg..................   2   1   seq
............................................................acacaaagaacggcugcCcg..................   1   1   seq
............................................................acacaaagaacggcugcUcg..................   1   1   seq
............................................................acacaaUgaacggcugcgcg..................   2   1   seq
............................................................acacaaagaacggcugcAcg..................   3   1   seq
............................................................acacaaagaacgAcugcgcg..................   3   1   seq
............................................................acacaaGgaacggcugcgcg..................   2   1   seq
............................................................acacaaagaacggcugcgUg..................   2   1   seq
............................................................acacaGagaacggcugcgcg..................   3   1   seq
............................................................acacaaagaacggcugcgGg..................   4   1   seq
............................................................acacaaagaaAggcugcgcgg.................   1   1   seq
............................................................acGcaaagaacggcugcgcgg.................   1   1   seq
............................................................acacaaagaacggcugcgcgU.................   12  1   seq
............................................................acacaaagaacggcugcgUgg.................   1   1   seq
............................................................acacaaagaacggcugcgcgg.................   140 0   seq
............................................................acacaaagaacggcugcgcgC.................   9   1   seq
............................................................acacGaagaacggcugcgcgg.................   1   1   seq
............................................................acacaaagCacggcugcgcgg.................   1   1   seq
............................................................acacaaGgaacggcugcgcgg.................   1   1   seq
............................................................acacaaagaacgCcugcgcgg.................   1   1   seq
............................................................acacaaagaacggcugcgGgg.................   2   1   seq
............................................................acacaaagaacggcugcgcgA.................   34  1   seq
............................................................acaAaaagaacggcugcgcgg.................   1   1   seq
............................................................acUcaaagaacggcugcgcgga................   1   1   seq
............................................................acacaCagaacggcugcgcgga................   1   1   seq
............................................................acacaaagaacggcugcgcggU................   129 1   seq
............................................................acacaaaAaacggcugcgcgga................   1   1   seq
............................................................acaAaaagaacggcugcgcgga................   1   1   seq
............................................................acacaaagaacggcugcgAgga................   2   1   seq
............................................................acacaaagaacggcugcgcggC................   64  1   seq
............................................................acacaGagaacggcugcgcgga................   6   1   seq
............................................................acacaaagaacggcugcgcgUa................   3   1   seq
............................................................acacaaaCaacggcugcgcgga................   1   1   seq
............................................................acacaaagaacggcugcCcgga................   1   1   seq
............................................................acacaaagaacAgcugcgcgga................   2   1   seq
............................................................acacaaagaacggAugcgcgga................   1   1   seq
............................................................acacaaagaacgCcugcgcgga................   2   1   seq
............................................................Gcacaaagaacggcugcgcgga................   1   1   seq
............................................................acacaaagaacggcugcgGgga................   2   1   seq
............................................................Ccacaaagaacggcugcgcgga................   1   1   seq
............................................................acGcaaagaacggcugcgcgga................   1   1   seq
............................................................aAacaaagaacggcugcgcgga................   1   1   seq
............................................................acacaaagaacggGugcgcgga................   1   1   seq
............................................................acacaaGgaacggcugcgcgga................   1   1   seq
............................................................acacaaagaacggcAgcgcgga................   1   1   seq
............................................................acacaaagaacggUugcgcgga................   1   1   seq
............................................................acacaaagaGcggcugcgcgga................   1   1   seq
............................................................acacaaagaacggcuAcgcgga................   1   1   seq
............................................................acacaaagaacggcCgcgcgga................   2   1   seq
............................................................acacaaagGacggcugcgcgga................   1   1   seq
............................................................acacaaagaacggcugcgUgga................   2   1   seq
............................................................acacaaagaacCgcugcgcgga................   1   1   seq
............................................................Ucacaaagaacggcugcgcgga................   1   1   seq
............................................................acacaaagaacggcugcgcgCa................   3   1   seq
............................................................acacaaagaaUggcugcgcgga................   2   1   seq
............................................................acacaaagaacUgcugcgcgga................   2   1   seq
............................................................acacaaagaacggcugcgcgga................   373 0   seq
............................................................acacaaagaacggcugAgcgga................   2   1   seq
............................................................acacaaagaacggcugUgcgga................   1   1   seq
............................................................acacaaagaacggcugcgcggaG...............   1   1   seq
............................................................acacaaagaacggcugcgcggCa...............   1   1   seq
............................................................acacaaagaacggcugcgcggUa...............   1   1   seq
.............................................................cacaaagaacggcugcgc...................   7   0   seq
.............................................................cacaaagaacggcugcgU...................   3   1   seq
.............................................................cacaaagaacggcugcgA...................   1   1   seq
.............................................................cacaaagaacggcugcgcg..................   90  0   seq
.............................................................cacaaagaacggcugcgGg..................   2   1   seq
.............................................................cacaaagaacggcugcgUg..................   1   1   seq
```

```
                  novel-nve-miR-44-1_star
                                                                                       novel-nve-miR-44-1_guide
gcaaaggcuagucuuucgcgcagccguucuuuauuguuuuuuuaugucacacaaagaacgacacaaagaacggcugcgcggaaaacuaguaaaagcuc

.............................................................cacaaagaacggcugcgcU..................    3     1     seq
.............................................................cacaaagaCcggcugcgcg..................    1     1     seq
.............................................................cacaaagGacggcugcgcg..................    1     1     seq
.............................................................cacUaagaacggcugcgcg..................    1     1     seq
.............................................................Uacaaagaacggcugcgcg..................    1     1     seq
.............................................................cacaaagaacggcugcgcA..................    18    1     seq
.............................................................cacaaagaacggcugcgcgC.................    3     1     seq
.............................................................cacaaagaacggcugcgcgA.................    12    1     seq
.............................................................cacaaagaacggcugcgcgg.................    36    0     seq
.............................................................cacaaagaacggcugcgcgU.................    2     1     seq
.............................................................cacaGagaacggcugcgcgg.................    1     1     seq
.............................................................cacUaagaacggcugcgcgga................    1     1     seq
.............................................................cacaaagaacggcugcgcgga................    46    0     seq
.............................................................cacaaagaacggcugcgUgga................    2     1     seq
.............................................................cacaaGgaacggcugcgcgga................    2     1     seq
.............................................................cacaaagaacggcugcgcgUa................    1     1     seq
.............................................................cacaaagaacggcugcgcggC................    7     1     seq
.............................................................cGcaaagaacggcugcgcgga................    1     1     seq
.............................................................cacaaagaacggcugcAcgga................    1     1     seq
.............................................................cacaaagaacggcugcgcggU................    26    1     seq
.............................................................cacaaagaacggcugcgcUga................    1     1     seq
.............................................................cacaaagaaAggcugcgcgga................    1     1     seq
.............................................................caUaaagaacggcugcgcgga................    1     1     seq
.............................................................cacaaagaacggcugcgcggCa...............    1     1     seq
..............................................................Ccaaagaacggcugcgcg..................    1     1     seq
..............................................................acaaaCaacggcugcgcg..................    1     1     seq
..............................................................acaaagaacggcugcgcA..................    5     1     seq
..............................................................acaaagaacggcugcgcg..................    11    0     seq
..............................................................acaaagaacggcugcgcgA.................    4     1     seq
..............................................................acaaagaacggcugcgcgU.................    3     1     seq
..............................................................acaaagaacggcugcgcgg.................    4     0     seq
..............................................................acaaagaacggcugcgcggC................    1     1     seq
..............................................................acaaagaacggcugcgcggU................    1     1     seq
..............................................................acaaagaacggcugcgcgga................    2     0     seq
..............................................................acaaagaacggcugcgcggaU...............    1     1     seq
...............................................................caaagaacggcugcgcggU................    1     1     seq
```

miRBase precursor : novel-nve-miR-44-2
Total read count : 13570
novel-nve-miR-44-2_guide read count : 13556
novel-nve-miR-44-2_star read count : 0
remaining reads : 14

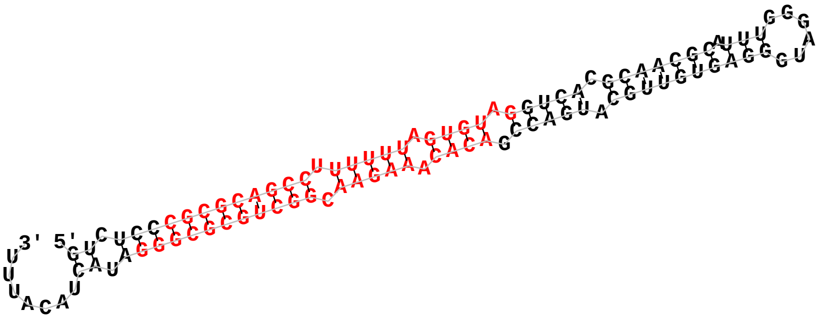

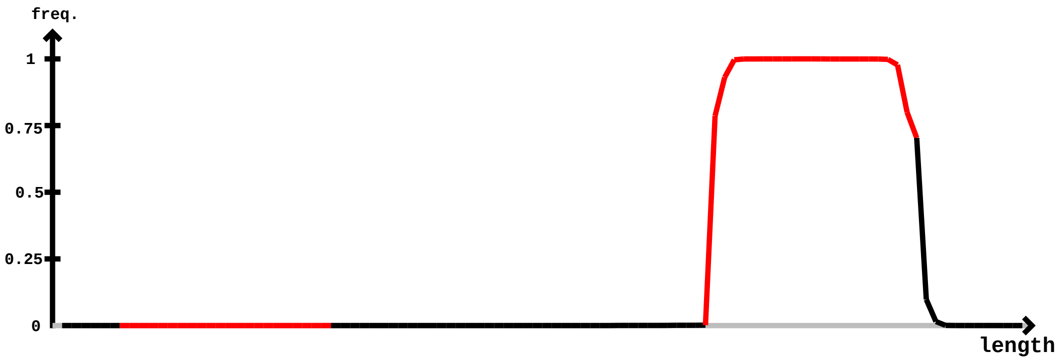


```
                                                                 novel-nve-miR-44-2_guide
         novel-nve-miR-44-2_star
5'- gucucccgcgcagccuuuuuuaguguaggucacgcaacgcauuugggaugggaguguugcaugaccgacacaaagaacggcugcgcgggauacuacauuu  -3'   exp
    ((.((((((((((((.(((((.((((.(((((.(((((((.(((......))))))))))).))))).)))).))))).))))))))))).)).......   reads      mm        sample
    .........................................................ugcGugaccgacacaaagaac......................   1          1         seq
    .........................................................ugcGugaccgacacaaagaacg.....................   4          1         seq
    ..............................................................gaccgacacaaagaacggcugcgcgg............   1          0         seq
    ..............................................................gaccgacacaaagaacggcugcgcUgg...........   1          1         seq
    ...............................................................acUgacacaaagaacggcug.................   1          1         seq
    ...............................................................accgacacaaagaacggcugcg...............   1          0         seq
    ...............................................................accgacacaaagaacggcugcgU..............   4          1         seq
    .................................................................cgacacaaagaacggcugcg...............   1          0         seq
    .................................................................cgacacaaagaacggcugcgG..............   1          1         seq
    ..................................................................gacacaaagaacggcugc................   1          0         seq
    ..................................................................gacacaaagGacggcugcgc..............   2          1         seq
    ..................................................................Cacacaaagaacggcugcgcg.............   1          1         seq
    ..................................................................Cacacaaagaacggcugcgcggg...........   3          1         seq
    ...................................................................acacaaagaacggcugcA...............   2          1         seq
    ...................................................................acacaaagaacggcugcU...............   1          1         seq
    ...................................................................acacaaagaacggcugcg...............   14         0         seq
    ...................................................................acacaaagaacggcAgcgc..............   1          1         seq
    ...................................................................acacGaagaacggcugcgc..............   1          1         seq
    ...................................................................acacaaagGacggcugcgc..............   2          1         seq
    ...................................................................acacaaagaacggUugcgc..............   1          1         seq
    ...................................................................acacaaagaacggcuAcgc..............   1          1         seq
    ...................................................................acacaaagaaUggcugcgc..............   1          1         seq
    ...................................................................acacaaagaacggcugcgU..............   52         1         seq
    ...................................................................Ucacaaagaacggcugcgc..............   1          1         seq
    ...................................................................acacaaagaacggcugcgc..............   142        0         seq
    ...................................................................acacaaagaacggcugcUc..............   1          1         seq
    ...................................................................acacaaagaacggcugcgA..............   11         1         seq
    ...................................................................acacaaagaacggcugcgG..............   5          1         seq
    ...................................................................acacaaUgaacggcugcgc..............   1          1         seq
    ...................................................................acacaaagaacggcugcCcg.............   1          1         seq
    ...................................................................acacaaagaacggcugcgcA.............   336        1         seq
    ...................................................................acacaaagaacggcugcgcC.............   21         1         seq
    ...................................................................acacaaagaacggcCgcgcg.............   1          1         seq
    ...................................................................acacaaagaacggcugcgUg.............   5          1         seq
```

```
                                                                  novel-nve-miR-44-2_guide
      novel-nve-miR-44-2_star
gucucccgcgcagccuuuuuuaguguaggucacgcaacgcauuugggaugggaguguugcaugaccgacacaaagaacggcugcgcgggauacuacauuu

...................................................................acacaaagaacggcugGgcg.............  1      1   seq
...................................................................acacaGagaacggcugcgcg.............  6      1   seq
...................................................................acacaaagaacggcugcAcg.............  4      1   seq
...................................................................acacaaagCacggcugcgcg.............  3      1   seq
...................................................................acacaaagaacggcugcUcg.............  1      1   seq
...................................................................acacGaagaacggcugcgcg.............  3      1   seq
...................................................................acacaaagaacggcugcgcg.............  1461   0   seq
...................................................................Ccacaaagaacggcugcgcg.............  4      1   seq
...................................................................acacaaagaacggcAgcgcg.............  1      1   seq
...................................................................acacaaagaacggcugcgGg.............  4      1   seq
...................................................................acacaaagaaAggcugcgcg.............  1      1   seq
...................................................................acacaaagaacggcugAgcg.............  2      1   seq
...................................................................acacaaagaacAgcugcgcg.............  1      1   seq
...................................................................acacaaagaacggcuAcgcg.............  2      1   seq
...................................................................acacaaagGacggcugcgcg.............  4      1   seq
...................................................................acUcaaagaacggcugcgcg.............  2      1   seq
...................................................................acacaaagaGcggcugcgcg.............  5      1   seq
...................................................................acacaaagaacggcugcgcU.............  70     1   seq
...................................................................acacaaagaCcggcugcgcg.............  1      1   seq
...................................................................acacaaagaacggcugUgcg.............  5      1   seq
...................................................................acacaaUgaacggcugcgcg.............  2      1   seq
...................................................................aAacaaagaacggcugcgcg.............  4      1   seq
...................................................................aUacaaagaacggcugcgcg.............  4      1   seq
...................................................................acacaaagaacggAugcgcg.............  2      1   seq
...................................................................acacaaagaacgCcugcgcg.............  1      1   seq
...................................................................acaUaaagaacggcugcgcg.............  4      1   seq
...................................................................acacaaagaacgAcugcgcg.............  4      1   seq
...................................................................acacaaagaacggUugcgcg.............  3      1   seq
...................................................................acacaaagaacggcugcgAg.............  6      1   seq
...................................................................acaAaaagaacggcugcgcg.............  4      1   seq
...................................................................acacaaagaacCgcugcgcg.............  3      1   seq
...................................................................acacaaaCaacggcugcgcg.............  1      1   seq
...................................................................acCcaaagaacggcugcgcg.............  1      1   seq
...................................................................acacaaCgaacggcugcgcg.............  1      1   seq
...................................................................acacaaagaaUggcugcgcg.............  4      1   seq
...................................................................acGcaaagaacggcugcgcg.............  6      1   seq
...................................................................acacaaagaacgUcugcgcg.............  1      1   seq
...................................................................acacaaGgaacggcugcgcg.............  3      1   seq
...................................................................acacaaagaacggcugcgGgg............  4      1   seq
...................................................................Ccacaaagaacggcugcgcgg............  1      1   seq
...................................................................acacaaagaacggcugcgcgA............  128    1   seq
...................................................................acacaaagaacggcugAgcgg............  1      1   seq
...................................................................acacaaagaaAggcugcgcgg............  2      1   seq
...................................................................acacaaagaacggcugcgcgU............  60     1   seq
...................................................................acacaaGgaacggcugcgcgg............  3      1   seq
...................................................................acacaaagaacgCcugcgcgg............  1      1   seq
...................................................................acacaaagaacggcugcgcUg............  2      1   seq
...................................................................aAacaaagaacggcugcgcgg............  1      1   seq
...................................................................acacaaagaacggcugcgUgg............  4      1   seq
...................................................................acacaaagaacggAugcgcgg............  2      1   seq
...................................................................acacaGagaacggcugcgcgg............  2      1   seq
...................................................................acGcaaagaacggcugcgcgg............  5      1   seq
...................................................................acacaaagaacggcugcgcCg............  2      1   seq
...................................................................acacaaagaacggcugUgcgg............  1      1   seq
...................................................................acacaaagaGcggcugcgcgg............  1      1   seq
...................................................................acacaaagaacggcCgcgcgg............  2      1   seq
...................................................................acacaaagaacggcugcgcAg............  2      1   seq
...................................................................acacaaagaacggcugcgcgg............  672    0   seq
...................................................................acacaaagCacggcugcgcgg............  2      1   seq
...................................................................acaAaaagaacggcugcgcgg............  1      1   seq
...................................................................acacGaagaacggcugcgcgg............  1      1   seq
...................................................................acacaaagaacggcugcAcgg............  1      1   seq
...................................................................acacaaaUaacggcugcgcgg............  1      1   seq
...................................................................Ucacaaagaacggcugcgcgg............  1      1   seq
...................................................................acacaaagaacggcugcgcgC............  22     1   seq
...................................................................acacaaagaaUggcugcgcgg............  1      1   seq
...................................................................acacaaagaacggUugcgcgg............  3      1   seq
...................................................................acacaaagaCcggcugcgcgg............  1      1   seq
...................................................................acacaaagaacggcugcgcgAg...........  15     1   seq
```

```
                                                                   novel-nve-miR-44-2_guide
      novel-nve-miR-44-2_star
gucucccgcgcagccuuuuuuaguguaggucacgcaacgcauuugggaugggaguguugcaugaccgacacaaagaacggcugcgcgggauacuacauuu

...................................................................acacaaagaaUggcugcgcggg...........   12      1       seq
...................................................................acacaaagaacggcugcgAggg...........   13      1       seq
...................................................................acacaaaUaacggcugcgcggg...........   1       1       seq
...................................................................acacaaagaacggcugcCcggg...........   4       1       seq
...................................................................acacaaagGacggcugcgcggg...........   44      1       seq
...................................................................acacaaagaacggcugcgcAgg...........   20      1       seq
...................................................................acacaGagaacggcugcgcggg...........   20      1       seq
...................................................................acacaaagaacggcugcgUggg...........   6       1       seq
...................................................................acacaaagaGcggcugcgcggg...........   18      1       seq
...................................................................acacaaagaacgAcugcgcggg...........   2       1       seq
...................................................................acacaaagaacggcugGgcggg...........   2       1       seq
...................................................................acacaaagaaGggcugcgcggg...........   1       1       seq
...................................................................acacaaagaacggcCgcgcggg...........   13      1       seq
...................................................................acacaaagaacggcugcgcggg...........   5999    0       seq
...................................................................acacaaagaacggAugcgcggg...........   5       1       seq
...................................................................acacaaagaacggcugcUcggg...........   4       1       seq
...................................................................acacaaagaacggcugcgcggU...........   307     1       seq
...................................................................acacaaagaacgUcugcgcggg...........   8       1       seq
...................................................................aAacaaagaacggcugcgcggg...........   7       1       seq
...................................................................aGacaaagaacggcugcgcggg...........   2       1       seq
...................................................................acacUaagaacggcugcgcggg...........   5       1       seq
...................................................................Ucacaaagaacggcugcgcggg...........   10      1       seq
...................................................................aUacaaagaacggcugcgcggg...........   17      1       seq
...................................................................acacaaaCaacggcugcgcggg...........   4       1       seq
...................................................................acacaaCgaacggcugcgcggg...........   2       1       seq
...................................................................acacaaaAaacggcugcgcggg...........   3       1       seq
...................................................................acacaaagaacggcugcgcCgg...........   10      1       seq
...................................................................acacaUagaacggcugcgcggg...........   7       1       seq
...................................................................acacaaagaacggcAgcgcggg...........   7       1       seq
...................................................................acacaaagaacggcugUgcggg...........   27      1       seq
...................................................................acacaaagaacgCcugcgcggg...........   8       1       seq
...................................................................acacaaagCacggcugcgcggg...........   10      1       seq
...................................................................acacaaagaacggcuCcgcggg...........   2       1       seq
...................................................................acacaaagaacggcuUcgcggg...........   9       1       seq
...................................................................acacaaagUacggcugcgcggg...........   3       1       seq
...................................................................acacaaagaacggcugcgGggg...........   4       1       seq
...................................................................acacaaagaacggcugcgcUgg...........   2       1       seq
...................................................................acacGaagaacggcugcgcggg...........   31      1       seq
...................................................................acCcaaagaacggcugcgcggg...........   1       1       seq
...................................................................acacaaagaCcggcugcgcggg...........   4       1       seq
...................................................................acacaaagaacggcugcgcgUg...........   26      1       seq
...................................................................acUcaaagaacggcugcgcggg...........   6       1       seq
...................................................................acacaaagaacCgcugcgcggg...........   7       1       seq
...................................................................acacaaagaacggcGgcgcggg...........   1       1       seq
...................................................................acGcaaagaacggcugcgcggg...........   26      1       seq
...................................................................acacaaagaacAgcugcgcggg...........   15      1       seq
...................................................................acacaaagaacggcuAcgcggg...........   7       1       seq
...................................................................acacaaagaUcggcugcgcggg...........   4       1       seq
...................................................................acacaaUgaacggcugcgcggg...........   3       1       seq
...................................................................acacaaagaacggUugcgcggg...........   8       1       seq
...................................................................Ccacaaagaacggcugcgcggg...........   22      1       seq
...................................................................acacaaGgaacggcugcgcggg...........   23      1       seq
...................................................................acacaaagaacggcugAgcggg...........   10      1       seq
...................................................................acaUaaagaacggcugcgcggg...........   13      1       seq
...................................................................acaAaaagaacggcugcgcggg...........   12      1       seq
...................................................................acacaaagaacggcugcAcggg...........   9       1       seq
...................................................................acacaaagaacggcugcgcggC...........   125     1       seq
...................................................................acacaaagaacUgcugcgcggg...........   6       1       seq
...................................................................acacaaagaacggcugcgcgCg...........   39      1       seq
...................................................................acacaaagaaAggcugcgcggg...........   12      1       seq
...................................................................acaUaaagaacggcugcgcggga..........   2       1       seq
...................................................................acacaaagaacggcugcgcgUga..........   1       1       seq
...................................................................acacaaagaGcggcugcgcggga..........   1       1       seq
...................................................................acacaaagaacggcugcgUggga..........   1       1       seq
...................................................................acacaaagaacggcugcgcgggU..........   162     1       seq
...................................................................acacaGagaacggcugcgcggga..........   1       1       seq
...................................................................acacaaagaacggcugcgcgggG..........   14      1       seq
...................................................................acacaaagaacggcugcgcggUa..........   4       1       seq
...................................................................acacaaagaacAgcugcgcggga..........   2       1       seq
```

```
                                                                   novel-nve-miR-44-2_guide
      novel-nve-miR-44-2_star
gucucccgcgcagccuuuuuuaguguaggucacgcaacgcauuugggaugggaguguugcaugaccgacacaaagaacggcugcgcgggauacuacauuu

...................................................................acacaaagCacggcugcgcggga..........    1      1      seq
...................................................................acacaaagaacggcugcgcggCa..........    1      1      seq
...................................................................acacaaagaacggcugcgcgggC..........    52     1      seq
...................................................................acacaaagaacggcugcgcggga..........    151    0      seq
...................................................................acGcaaagaacggcugcgcggga..........    2      1      seq
...................................................................acacaaagaacggcuAcgcggga..........    1      1      seq
...................................................................acacaaGgaacggcugcgcggga..........    1      1      seq
...................................................................acacaaagaacggcugcgcgggaA.........    3      1      seq
...................................................................acacaaagaacggcugcgcgggaG.........    1      1      seq
...................................................................acacaaagaacggcugcgcgggUu.........    28     1      seq
...................................................................acacaaagaacggcugcgcgggau.........    2      0      seq
...................................................................acacaaagaacggcugcgcgggauU........    1      1      seq
...................................................................acacaaagaacggcugcgcgggauC........    1      1      seq
...................................................................acacaaagaacggcugcgcgggaGa........    1      1      seq
...................................................................acacaaagaacggcugcgcgggauUc.......    1      1      seq
...................................................................acacaaagaacggcugcgcgggauaU.......    1      1      seq
....................................................................cacaaagaacggAugcgc..............    1      1      seq
....................................................................cacaaagaacggcugcgA..............    3      1      seq
....................................................................cacaaagaacggcugcgc..............    42     0      seq
....................................................................cacaaGgaacggcugcgc..............    1      1      seq
....................................................................cacaaagaacggcugcgU..............    7      1      seq
....................................................................cacaaagaacggcugcgGg.............    2      1      seq
....................................................................cacaaagaacggcugcgcU.............    11     1      seq
....................................................................cacaaagaacggcCgcgcg.............    1      1      seq
....................................................................Uacaaagaacggcugcgcg.............    2      1      seq
....................................................................cacaGagaacggcugcgcg.............    1      1      seq
....................................................................cacaaagaacggcugcgcC.............    6      1      seq
....................................................................cacGaagaacggcugcgcg.............    1      1      seq
....................................................................cGcaaagaacggcugcgcg.............    1      1      seq
....................................................................cacaaagGacggcugcgcg.............    1      1      seq
....................................................................cacaaGgaacggcugcgcg.............    1      1      seq
....................................................................cacaaagaacggcugcCcg.............    1      1      seq
....................................................................cacaaagaacggcugcgcg.............    271    0      seq
....................................................................cacaaagaacggcugcgUg.............    1      1      seq
....................................................................cacaaagaGcggcugcgcg.............    1      1      seq
....................................................................caAaaagaacggcugcgcg.............    1      1      seq
....................................................................cacaaagaacggcugUgcg.............    2      1      seq
....................................................................cacUaagaacggcugcgcg.............    1      1      seq
....................................................................cacaaagaacggcugcgcA.............    60     1      seq
....................................................................cacaaagaCcggcugcgcg.............    3      1      seq
....................................................................cacaaagaacggcugcgcgA............    43     1      seq
....................................................................cacaaagaacCgcugcgcgg............    1      1      seq
....................................................................cacaaagCacggcugcgcgg............    1      1      seq
....................................................................cacaaagaaAggcugcgcgg............    1      1      seq
....................................................................cacaaagaacggcugcgcgg............    240    0      seq
....................................................................cacaaagaUcggcugcgcgg............    1      1      seq
....................................................................cUcaaagaacggcugcgcgg............    1      1      seq
....................................................................cacaaagaacggcugUgcgg............    2      1      seq
....................................................................cGcaaagaacggcugcgcgg............    1      1      seq
....................................................................cacaaagGacggcugcgcgg............    1      1      seq
....................................................................cacaaagaacggcugcgcgU............    11     1      seq
....................................................................cacaaagaacggcugcgGgg............    2      1      seq
....................................................................cacaaagaacggcugcgcAg............    1      1      seq
....................................................................cacaaagaacggcuAcgcgg............    1      1      seq
....................................................................cacaaagaacggcugcgcUg............    1      1      seq
....................................................................cacaGagaacggcugcgcgg............    1      1      seq
....................................................................cacaaagaacggcugcgcgC............    6      1      seq
....................................................................Uacaaagaacggcugcgcgg............    1      1      seq
....................................................................Uacaaagaacggcugcgcggg...........    2      1      seq
....................................................................cacaaagaacggcugUgcggg...........    4      1      seq
....................................................................caGaaagaacggcugcgcggg...........    1      1      seq
....................................................................cacaaagaacggcAgcgcggg...........    1      1      seq
....................................................................cacaaagaacggcugcgcggC...........    20     1      seq
....................................................................cacaaGgaacggcugcgcggg...........    2      1      seq
....................................................................cacaaagaGcggcugcgcggg...........    4      1      seq
....................................................................cacaaagGacggcugcgcggg...........    3      1      seq
....................................................................cacaaagaacggcuCcgcggg...........    1      1      seq
....................................................................cacaaagaacggAugcgcggg...........    1      1      seq
....................................................................cUcaaagaacggcugcgcggg...........    4      1      seq
```

```
                                                                  novel-nve-miR-44-2_guide
      novel-nve-miR-44-2_star
gucucccgcgcagccuuuuuuaguguaggucacgcaacgcauuugggaugggaguguugcaugaccgacacaaagaacggcugcgcgggauacuacauuu

....................................................................Aacaaagaacggcugcgcggg...........    2    1    seq
....................................................................cacUaagaacggcugcgcggg...........    1    1    seq
....................................................................cacGaagaacggcugcgcggg...........    3    1    seq
....................................................................cacaaagaUcggcugcgcggg...........    1    1    seq
....................................................................cacaaagaacggcugcCcggg...........    2    1    seq
....................................................................cacaaagaacggcugcgAggg...........    3    1    seq
....................................................................caUaaagaacggcugcgcggg...........    1    1    seq
....................................................................cacaaagaacggcugcgcAgg...........    4    1    seq
....................................................................cacaaagaacggcugcgcgCg...........    6    1    seq
....................................................................cacaaagaacggcugcgcggU...........    62   1    seq
....................................................................cacaaagaacggcugcgcCgg...........    1    1    seq
....................................................................cacaaagaacggcugcgcggg...........    922  0    seq
....................................................................cacaaagaacgAcugcgcggg...........    2    1    seq
....................................................................cacaaagaacggcugAgcggg...........    1    1    seq
....................................................................cacaaagaacggcugcAcggg...........    1    1    seq
....................................................................cacaaagaacggUugcgcggg...........    3    1    seq
....................................................................cacaaagaCcggcugcgcggg...........    2    1    seq
....................................................................cacaaagaacUgcugcgcggg...........    1    1    seq
....................................................................cCcaaagaacggcugcgcggg...........    1    1    seq
....................................................................cacaGagaacggcugcgcggg...........    2    1    seq
....................................................................cacaaagaacggcugcgcgUg...........    4    1    seq
....................................................................cacaaagaacggcugcgcgAg...........    2    1    seq
....................................................................cacaaaAaacggcugcgcggg...........    1    1    seq
....................................................................cGcaaagaacggcugcgcggg...........    4    1    seq
....................................................................cacaaagaGcggcugcgcggga..........    1    1    seq
....................................................................cacaaagaacggcugcgcggga..........    40   0    seq
....................................................................cacaaagaacggcugcgcgggG..........    3    1    seq
....................................................................cacaaagaacggcugcgcgggU..........    62   1    seq
....................................................................cacaaagaacggcuAcgcggga..........    1    1    seq
....................................................................cacaaagaacggcugcgcgggC..........    19   1    seq
....................................................................caUaaagaacggcugcgcggga..........    1    1    seq
....................................................................cacaaagaacggcugcgcggUa..........    1    1    seq
....................................................................cacaaagaacggcugcgcggCa..........    1    1    seq
....................................................................cacaGagaacggcugcgcggga..........    1    1    seq
....................................................................Gacaaagaacggcugcgcgggau.........    1    1    seq
....................................................................cacaaagaacggcugcgcgggau.........    1    0    seq
....................................................................cacaaagaacggcugcgcgggUu.........    17   1    seq
.....................................................................acaaagaacAgcugcgcg.............    1    1    seq
.....................................................................acaaagaacggcugcgcA.............    8    1    seq
.....................................................................acaaagaacggcugcgcg.............    30   0    seq
.....................................................................acaaaCaacggcugcgcg.............    1    1    seq
.....................................................................acaaagaacggcugcgcC.............    2    1    seq
.....................................................................Ccaaagaacggcugcgcg.............    1    1    seq
.....................................................................acaaagaacggcugcgcU.............    1    1    seq
.....................................................................acaaagaacggcugcgcgC............    1    1    seq
.....................................................................acaaagaacggcugcgcgA............    12   1    seq
.....................................................................acaaagaacggcugcgcgU............    7    1    seq
.....................................................................Gcaaagaacggcugcgcgg............    1    1    seq
.....................................................................acaaagaacUgcugcgcgg............    1    1    seq
.....................................................................acaaagaacggcugcCcgg............    1    1    seq
.....................................................................aUaaagaacggcugcgcgg............    1    1    seq
.....................................................................acaaagaacggcugcgcgg............    30   0    seq
.....................................................................acaaagaacggcugcgGggg...........    1    1    seq
.....................................................................acaaagaacggcugcAcggg...........    1    1    seq
.....................................................................acaaagaacggcugcgcggC...........    3    1    seq
.....................................................................acaaagaacAgcugcgcggg...........    1    1    seq
.....................................................................acaaagaacggcugcgcggU...........    4    1    seq
.....................................................................acGaagaacggcugcgcggg...........    2    1    seq
.....................................................................acaaagaacggcugUgcggg...........    1    1    seq
.....................................................................acaaagaacggcugcgcggg...........    75   0    seq
.....................................................................acaCagaacggcugcgcggg...........    1    1    seq
.....................................................................Gcaaagaacggcugcgcggg...........    1    1    seq
.....................................................................acaaagaacggAugcgcggg...........    1    1    seq
.....................................................................acaaagaacgAcugcgcggg...........    1    1    seq
.....................................................................acaaagaacAgcugcgcggga..........    1    1    seq
.....................................................................acaaagaacggcugcgcgggG..........    11   1    seq
.....................................................................Ccaaagaacggcugcgcggga..........    4    1    seq
.....................................................................acaaagaacggcugcgcgggU..........    385  1    seq
.....................................................................acaaagaacggcugcgcggga..........    72   0    seq
```

novel-nve-miR-44-2_guide

novel-nve-miR-44-2_star

gucucccgcgcagccuuuuuuaguguaggucacgcaacgcauuugggaugggaguguugcaugaccgacacaaagaacggcugcgcgggauacuacauuu

| | | | |
|---|---|---|---|
| .....................................................................acaaagaacggcCgcgcggga.......... | 1 | 1 | seq |
| .....................................................................acaaagaacggcugcgcgggC.......... | 89 | 1 | seq |
| .....................................................................acaaagaaAggcugcgcggga.......... | 1 | 1 | seq |
| .....................................................................acaaagaaUggcugcgcggga.......... | 1 | 1 | seq |
| .....................................................................Gcaaagaacggcugcgcggga.......... | 1 | 1 | seq |
| .....................................................................acaaagaacggcugcgcgggCu......... | 2 | 1 | seq |
| .....................................................................acaaagaacggcugcgcgggUu......... | 123 | 1 | seq |
| .....................................................................acaaagaacggcugcgcgggau......... | 9 | 0 | seq |
| .....................................................................acaaagaacggcugcgcgggaA......... | 1 | 1 | seq |
| .....................................................................acaaagaacggcugcgcgggUua........ | 3 | 1 | seq |
| .....................................................................acaaagaacggcugcgcgggauC........ | 1 | 1 | seq |
| ......................................................................caaagaUcggcugcgcggg........... | 1 | 1 | seq |
| ......................................................................caaagaacggcugcgcggU........... | 3 | 1 | seq |
| ......................................................................caaagaacggcugcgcggg........... | 4 | 0 | seq |
| ......................................................................caaagaacggcugcgcggC........... | 1 | 1 | seq |
| ......................................................................caaagaacggUugcgcggga.......... | 1 | 1 | seq |
| ......................................................................caaagaacAgcugcgcggga.......... | 1 | 1 | seq |
| ......................................................................caaagaacggcugcgcgggC.......... | 4 | 1 | seq |
| ......................................................................caaagaacggcugcgcggga.......... | 7 | 0 | seq |
| ......................................................................caaagaacggcugcgcgggG.......... | 1 | 1 | seq |
| ......................................................................caaagaacggcugcgcgggU.......... | 11 | 1 | seq |
| ......................................................................caaagaacggcugcgcgggUu......... | 3 | 1 | seq |
| ......................................................................caaagaacggcugcgcgggaC......... | 1 | 1 | seq |
| ......................................................................caaagaacggcugcgcgggau......... | 2 | 0 | seq |
| .......................................................................aaagaacggcugcgcggg........... | 1 | 0 | seq |
| .......................................................................aaagaacggcugcgcgggUu......... | 1 | 1 | seq |
| ........................................................................aagaacggcuUcgcggga.......... | 3 | 1 | seq |
| ............................................................................acggcugcgcgggaGacuaca... | 1 | 1 | seq |

miRBase precursor : novel-nve-miR-50
Total read count : 998
novel-nve-miR-50_guide read count : 998
novel-nve-miR-50_star read count : 0
remaining reads : 0

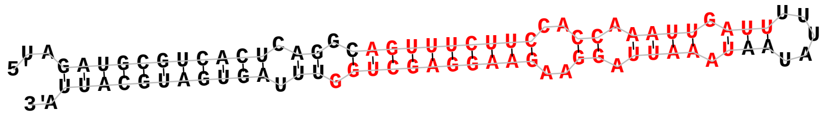

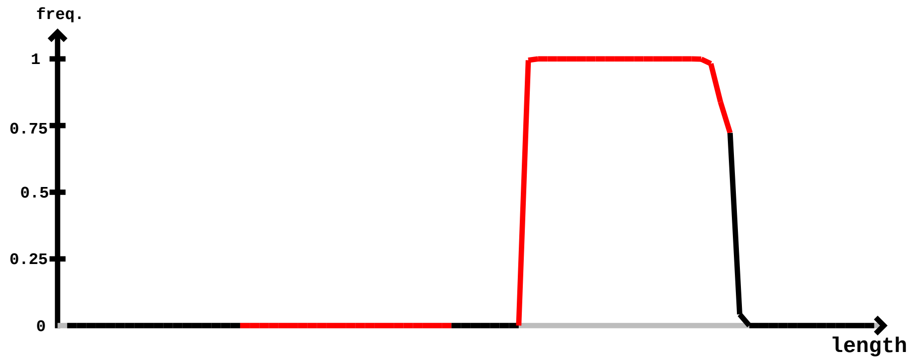


```
                                  novel-nve-miR-50_star
                                                            novel-nve-miR-50_guide
5'- uagaugcgucacucaggcaguuucuuccaccaaauugauuuuuauaauaaauuaggaagaaggagcugguuuagugaugcauua   -3'    exp
    ..(((((((((((.((.((((((((((..((.((((.(((.....))).)))).))..)))))))))).)).))))))))))).    reads       mm          sample
    ..............................................uaaauuaggaagaaggag....................    1           0           seq
    ..............................................uaaauuaggaagaaggagU...................    4           1           seq
    ..............................................uaaauuaggaagaaggagc...................    12          0           seq
    ..............................................uaaauuaggaagaaggagcA..................    13          1           seq
    ..............................................uGaauuaggaagaaggagcu..................    1           1           seq
    ..............................................uaaauuaggaagaaggUgcu..................    1           1           seq
    ..............................................uaGauuaggaagaaggagcu..................    1           1           seq
    ..............................................uaaauuaggaagaaggagcC..................    13          1           seq
    ..............................................uaaauuaggaagaaggGgcu..................    1           1           seq
    ..............................................Aaaauuaggaagaaggagcu..................    2           1           seq
    ..............................................uaaauuaggaGgaaggagcu..................    4           1           seq
    ..............................................uaaGuuaggaagaaggagcu..................    1           1           seq
    ..............................................uaaauuaggaagaaggagcu..................    100         0           seq
    ..............................................uUaauuaggaagaaggagcu..................    1           1           seq
    ..............................................uaaauuaggaagaaggagcG..................    2           1           seq
    ..............................................uaaaCuaggaagaaggagcu..................    1           1           seq
    ..............................................uaaauAaggaagaaggagcu..................    1           1           seq
    ..............................................uaaauuaggaagaaggagcCg.................    3           1           seq
    ..............................................uaaauuaggaagaaggagcuU.................    3           1           seq
    ..............................................uaaauuaggaagaGggagcug.................    1           1           seq
    ..............................................uaaauuaggaagaaggagcuC.................    2           1           seq
    ..............................................uaaauuaggaagaaggGgcug.................    3           1           seq
    ..............................................Aaaauuaggaagaaggagcug.................    2           1           seq
    ..............................................Caaauuaggaagaaggagcug.................    1           1           seq
    ..............................................uaaauuaggaagaaggagcuA.................    14          1           seq
    ..............................................uaaauuaggaagGaggagcug.................    1           1           seq
    ..............................................uaaaAuaggaagaaggagcug.................    1           1           seq
    ..............................................uaaauuaggaagaaggaUcug.................    1           1           seq
    ..............................................uaaauuaggaagaaggagcug.................    81          0           seq
    ..............................................uaaauuaggaagaaggagcGg.................    2           1           seq
    ..............................................uaaGuuaggaagaaggagcugg................    1           1           seq
    ..............................................Gaaauuaggaagaaggagcugg................    2           1           seq
    ..............................................uaaauuaggaagaaggGgcugg................    2           1           seq
    ..............................................uaaauuGggaagaaggagcugg................    2           1           seq
```

```
                  novel-nve-miR-50_star
                                               novel-nve-miR-50_guide
uagaugcgucacucaggcaguuucuuccaccaaauugauuuuuauaauaaauuaggaagaaggagcugguuuagugaugcauua

...............................................uaaauuaggaagGaggagcugg...............    3          1          seq
...............................................uGaauuaggaagaaggagcugg...............    1          1          seq
...............................................Caaauuaggaagaaggagcugg...............    1          1          seq
...............................................Aaaauuaggaagaaggagcugg...............    9          1          seq
...............................................uaaaAuaggaagaaggagcugg...............    1          1          seq
...............................................uaaauuaggaagaGggagcugg...............    1          1          seq
...............................................uaaauuaggaagaaggagcCgg...............    2          1          seq
...............................................uaaauuaggaagaaggagcugU...............    35         1          seq
...............................................uaaCuuaggaagaaggagcugg...............    1          1          seq
...............................................uaaauuaggaagaaggagcugC...............    121        1          seq
...............................................uaaauuaggaagaaggagcugA...............    43         1          seq
...............................................uaaauuaggaagaaggagUugg...............    1          1          seq
...............................................uaaauuaggGagaaggagcugg...............    2          1          seq
...............................................uaaauuaAgaagaaggagcugg...............    2          1          seq
...............................................uaaauuaggaGgaaggagcugg...............    17         1          seq
...............................................uaGauuaggaagaaggagcugg...............    2          1          seq
...............................................uaaauAaggaagaaggagcugg...............    1          1          seq
...............................................uaaauuagCaagaaggagcugg...............    3          1          seq
...............................................uaaauuaggaagaaggagcugg...............    421        0          seq
...............................................uaaauuagAaagaaggagcugg...............    1          1          seq
...............................................uaaaGuaggaagaaggagcugg...............    1          1          seq
...............................................uaaauuaggaagaaggagcAgg...............    1          1          seq
...............................................uaaauuaggaagaaggagcugCu..............    28         1          seq
...............................................uaaauuaggaagaagUagcuggu..............    1          1          seq
...............................................uaaauuaggaagaaggagcuggu..............    11         0          seq
...............................................uaaauuaggaagaaggagcuggA..............    1          1          seq
...............................................uaaauuaggaagaaggagcuggC..............    1          1          seq
................................................aaauuaggaagaaggagU..................    1          1          seq
................................................Uaauuaggaagaaggagcu.................    1          1          seq
................................................aaauuaggaagaaggagcug................    1          0          seq
................................................aaauuaggaagaaggagcugg...............    2          0          seq
```

miRBase precursor : novel-nve-miR-67
Total read count : 353
novel-nve-miR-67_guide read count : 353
novel-nve-miR-67_star read count : 0
remaining reads : 0

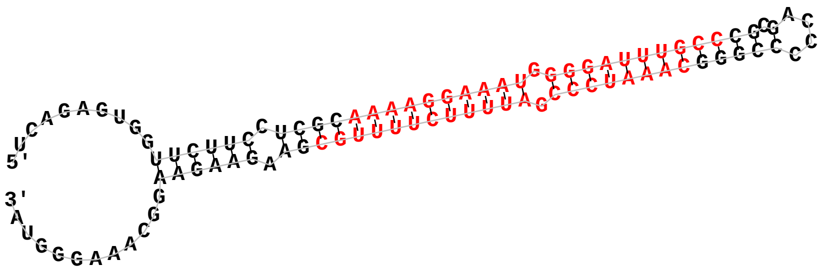

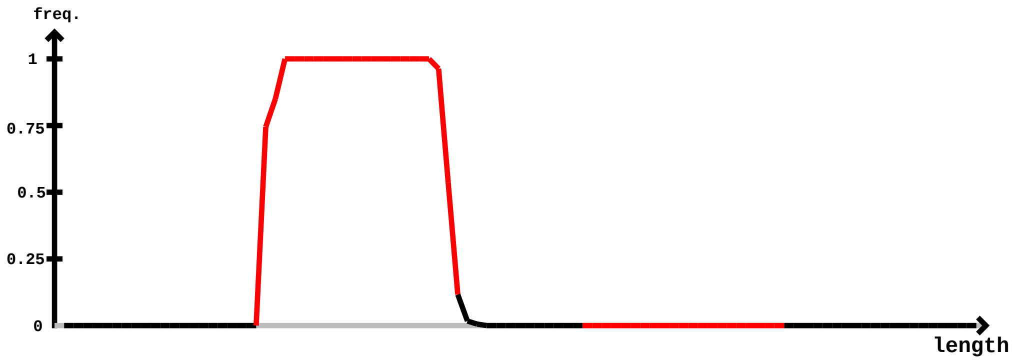


```
                                                                 novel-nve-miR-67_star
                         novel-nve-miR-67_guide
5'- ucagagugguucuuccucgcaaaaggaaauggggauuugcccgcgacccccgggcaaaucccgauuuucuuuugcgaagaagaaggcaaagggua  -3'   exp
    .....((..((((((.((((((((((((((.(((((((((((((.(....))))))))))))))).))))))))))))))).)))))).))......    reads   mm   sample
    ....................aaaaggaaauggggauuG.........................................................    1       1    seq
    ....................aaaaggaaauCgggauuu.........................................................    12      1    seq
    ....................aaaaggaaauggggauuug........................................................    2       0    seq
    ....................aaaaggaaauCgggauuug........................................................    123     1    seq
    ....................aaaaggaaauggggauuuA........................................................    1       1    seq
    ....................aaaaggaaauUgggauuugc.......................................................    1       1    seq
    ....................aaaaggaaauggggauuugc.......................................................    2       0    seq
    ....................aaaaggaaauCgggauuugc.......................................................    99      1    seq
    ....................aaaaggaaauggggauuugU.......................................................    1       1    seq
    ....................aaaaggaaauggggauuugcc......................................................    2       0    seq
    ....................aaaaggaaauCgggauuugcc......................................................    16      1    seq
    ....................aaaaggaaauggggauuugcU......................................................    1       1    seq
    ....................aaaaggaaauCgggauuugccc.....................................................    2       1    seq
    .....................aaaggaaauggggauuCg........................................................    2       1    seq
    .....................aaaggaaauCgggauuug........................................................    16      1    seq
    .....................aaaggaaauggggauuug........................................................    7       0    seq
    .....................aaaggaaauCgggauuugc.......................................................    3       1    seq
    .....................aaaggaaauggggauuugc.......................................................    4       0    seq
    .....................aaaggaaauCgggauuugcc......................................................    2       1    seq
    .....................aaaggaaauggggauuugcG......................................................    1       1    seq
    .....................aaaggaaauggggauuugccU.....................................................    1       1    seq
    .....................aaaggaaauggggauuugccG.....................................................    1       1    seq
    ......................aaggaaauCgggauuugc.......................................................    37      1    seq
    ......................aaggaaauggggauuugG.......................................................    1       1    seq
    ......................aaggaaauCgggauuugcc......................................................    13      1    seq
    ......................aaggaaauCgggauuugcccg....................................................    1       1    seq
    ......................aaggaaauggggauuugcccg....................................................    1       0    seq
```

miRBase precursor : novel-nve-miR-70
Total read count : 122
novel-nve-miR-70_guide read count : 122
novel-nve-miR-70_star read count : 0
remaining reads : 0

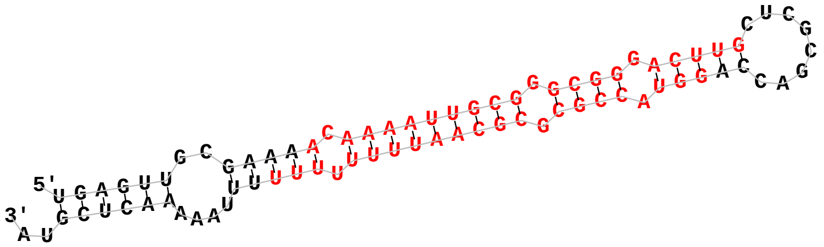

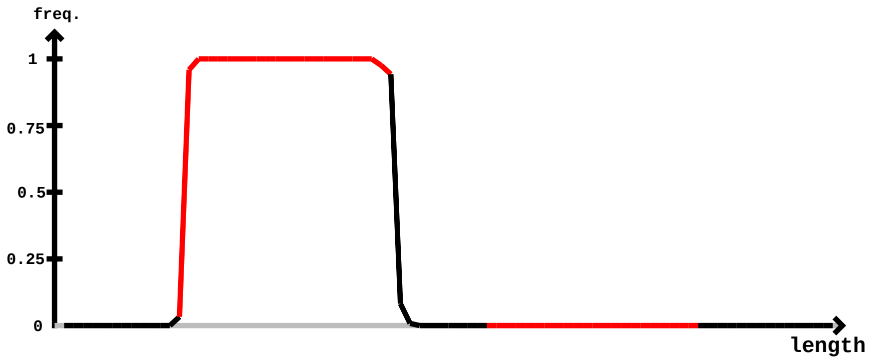


```
                                                      novel-nve-miR-70_star
                 novel-nve-miR-70_guide
5'- ugaguugcgaaaacaaaauugcgggcgggacuugcucgcgaccaggguaccgcgcgcaauuuuuuuuuuuaaaaacucgua   -3'    exp
    .(((((..(((((.(((((((((.((((.(((((........))))).)))).))))))))).)))))....)))))...    reads      mm         sample
    ...........aacaaaauugcgggAgggacuug..............................................    1          1          seq
    ...........Gacaaaauugcgggcgggacuug..............................................    1          1          seq
    ...........aacaaaauugcgggcgggacuug..............................................    2          0          seq
    ............acaaaauugcgggcgggacu................................................    2          0          seq
    ............acaaaauugcgggcgggacA................................................    1          1          seq
    ............acaaaauugcgggcgggacuu...............................................    4          0          seq
    ............acaaaauugcgggcgggacuuU..............................................    1          1          seq
    ............acGaaauugcgggcgggacuug..............................................    1          1          seq
    ............acaaaauugcgggcgggacuuA..............................................    11         1          seq
    ............acaaaauugcgggcgggacuug..............................................    81         0          seq
    ............Gcaaaauugcgggcgggacuug..............................................    2          1          seq
    ............acaGaauugcgggcgggacuug..............................................    3          1          seq
    ............acaaaGuugcgggcgggacuug..............................................    1          1          seq
    ............Ucaaaauugcgggcgggacuug..............................................    1          1          seq
    ............acaaaauugcgggcgggacuugA.............................................    1          1          seq
    ............acaaaauugcgggcgggacuugU.............................................    3          1          seq
    ............acaaaauugcgggcgggacuugcu............................................    1          0          seq
    .............caaaauugcgggcgggacuugU.............................................    2          1          seq
    .............caaaauugcgggcgggacuugc.............................................    3          0          seq
```

miRBase precursor : novel-nve-miR-74
Total read count : 199
novel-nve-miR-74_guide read count : 199
novel-nve-miR-74_star read count : 0
remaining reads : 0

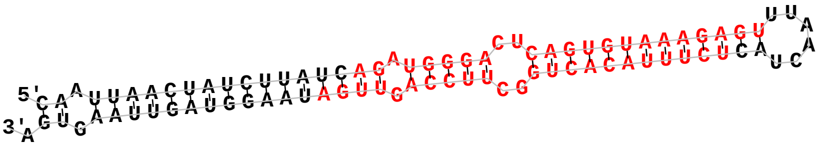

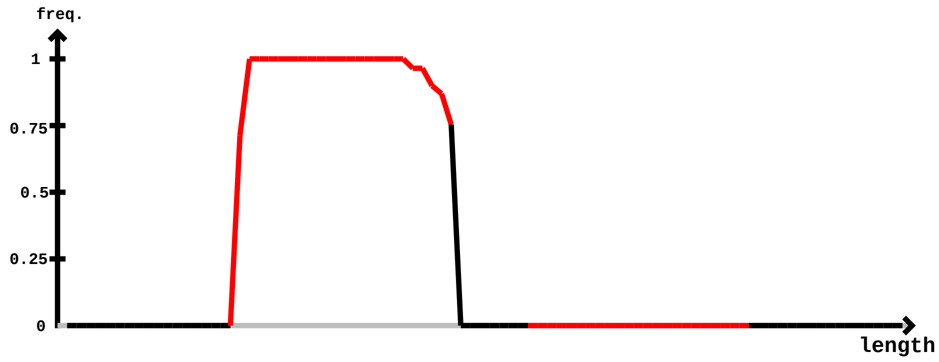


```
                                  novel-nve-miR-74_guide
                                                              novel-nve-miR-74_star
5'- caauuaacuaucuuaucagaugggacucaguguaaagaguuuaacuacucuuuacacuggcuuccaguugauaagguaguuaaguga  -3'    exp
    ((.((((((((((((((((.(((((..((((((((((((......)))))))))))))..))))).))))))))))))))))).)).    reads   mm   sample
    .................agaugggacucaguguaa....................................................    7       0    seq
    .................agaugggacucaguguaaag..................................................    4       0    seq
    .................agaugggacucaguguaGaga.................................................    1       1    seq
    .................agaugggacucaguguaaaga.................................................    5       0    seq
    .................agaugggacucaguguaaagag................................................    14      0    seq
    .................agaugggacuGaguguaaagag................................................    1       1    seq
    .................agGugggacucaguguaaagag................................................    1       1    seq
    .................agaugggacCcaguguaaagag................................................    1       1    seq
    .................agaugggacucaguguaaagagC...............................................    14      1    seq
    .................Ggaugggacucaguguaaagagu...............................................    1       1    seq
    .................agaugggacGcaguguaaagagu...............................................    1       1    seq
    .................agauAggacucaguguaaagagu...............................................    1       1    seq
    .................agaugggacucaguguaaagagu...............................................    82      0    seq
    .................agaugggacucaguguaaagagA...............................................    4       1    seq
    .................agaAgggacucaguguaaagagu...............................................    2       1    seq
    .................agaugggacucaguguaaGgagu...............................................    1       1    seq
    .................agGugggacucaguguaaagagu...............................................    1       1    seq
    .................agaugggacucagugCaaagagu...............................................    1       1    seq
    ..................gaugggacucaguguaaag..................................................    9       0    seq
    ..................gaugggacucaguguaaagag................................................    5       0    seq
    ..................gauCggacucaguguaaagag................................................    1       1    seq
    ..................gaugggacucaguguaaagagC...............................................    4       1    seq
    ..................gaugggacucaguguaaagagu...............................................    32      0    seq
    ..................gaugggacucagAguaaagagu...............................................    1       1    seq
    ..................gaugggacuUaguguaaagagu...............................................    3       1    seq
    ..................gaugggacucaguguGaagagu...............................................    1       1    seq
    ..................Aaugggacucaguguaaagagu...............................................    1       1    seq
```

miRBase precursor : novel-nve-miR-78
Total read count : 87
novel-nve-miR-78_guide read count : 87
novel-nve-miR-78_star read count : 0
remaining reads : 0

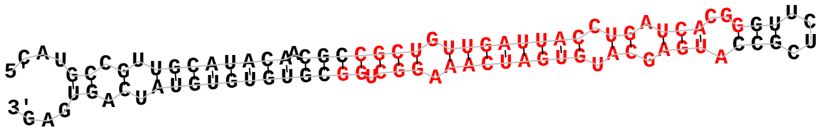

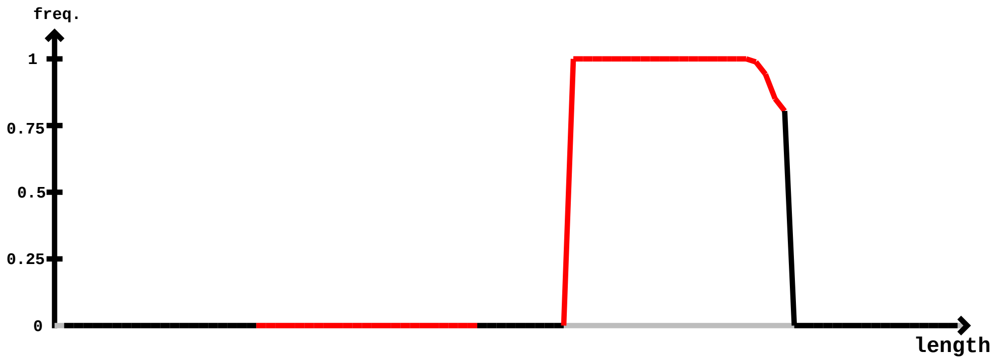


novel-nve-miR-78_star

novel-nve-miR-78_guide

```
5'- caugccguugcauacaacgccgcuguugauuaccugaucacggguucucgccaugagcaugugaucaaaggcuggcgugugugguaucagugag   -3'    exp
    (((.....((((((((.(((((((.((((((((.((.(((..(((....))).))).)).)))))))).))).))))))))))))...)))..      reads      mm         sample
    ....................................................augagcaugugaucaaagg......................      1          0          seq
    ....................................................augagcaugugaucaaaggc.....................      3          0          seq
    ....................................................augGgcaugugaucaaaggc.....................      1          1          seq
    ....................................................augagcaugugaucaaaggcu....................      6          0          seq
    ....................................................augagcaugugaucaaaggcC....................      1          1          seq
    ....................................................Gugagcaugugaucaaaggcu....................      1          1          seq
    ....................................................augagcaugugaucaaaggcuU...................      1          1          seq
    ....................................................augagcaugugaucaaaggcug...................      3          0          seq
    ....................................................augUgcaugugaucaaaggcugg..................      1          1          seq
    ....................................................augagcGugugaucaaaggcugg..................      1          1          seq
    ....................................................augagcaugugaucaaaggcugC..................      1          1          seq
    ....................................................augagcaugugaucaaagAcugg..................      1          1          seq
    ....................................................augagcaugugaucaaaggcugU..................      3          1          seq
    ....................................................augagcaugugGucaaaggcugg..................      1          1          seq
    ....................................................augagcaugugaucaaaggcugA..................      8          1          seq
    ....................................................augagcaugugaucaaaggcugg..................      54         0          seq
```

miRBase precursor : novel-nve-miR-80
Total read count : 121
novel-nve-miR-80_guide read count : 121
novel-nve-miR-80_star read count : 0
remaining reads : 0

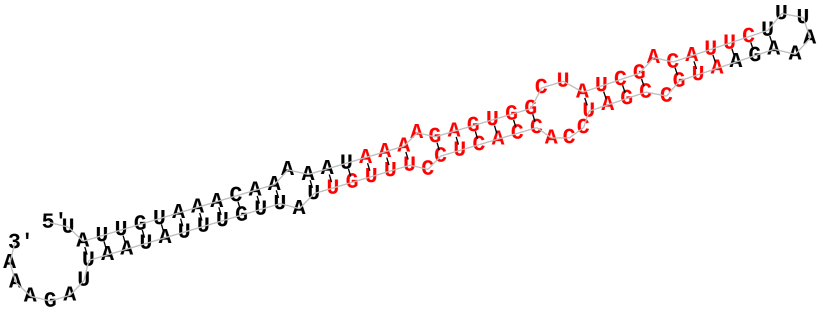

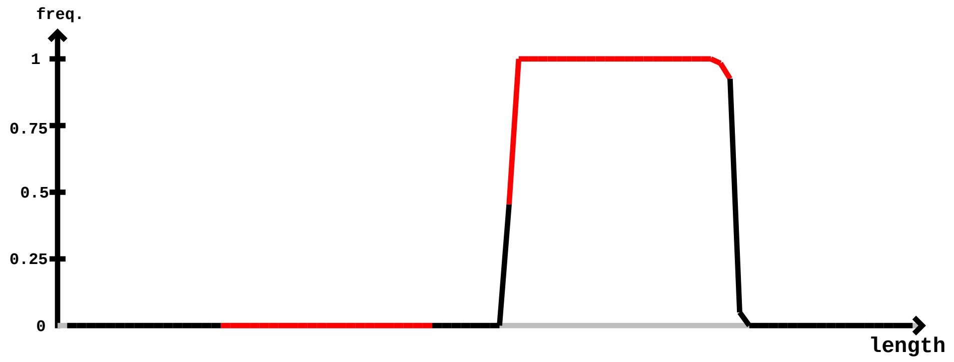


```
                 novel-nve-miR-80_star
                                             novel-nve-miR-80_guide
5'- uauuguaaacaaaaauaaaagaguggcuaucgacauucuuuaaagaaugccgauccaccacuccuuuguuauuguuuauaauuagaaa   -3'    exp
    .(((((((((((.((((((.((((((..((((.((((((....)))))).))))...)))))).)))))).)))))))))))......    reads      mm         sample
    .............................................aaugccgauccaccacuccuuu.....................    2          0          seq
    .............................................aaugccgauccaccacuccuuug....................    3          0          seq
    .............................................aaugccgauccaccacuccuuuA....................    1          1          seq
    .............................................aaugccgauccaccacuccuuCgu...................    1          1          seq
    .............................................aaugccgauccaccacuccuuugG...................    1          1          seq
    .............................................aaugccgauccaccacuccuuugC...................    10         1          seq
    .............................................aaugccgauccaccacuccuuugA...................    1          1          seq
    .............................................aaugccgauccaccacuUcuuugu...................    1          1          seq
    .............................................aaugccgauccaccacuccuuugu...................    35         0          seq
    ..............................................augccgauccaccacuccuuGg....................    1          1          seq
    ..............................................augccgauccaccacuccuuug....................    2          0          seq
    ..............................................augccgauccaccacuccuuugC...................    8          1          seq
    ..............................................augccgauccaccacuccuuugG...................    2          1          seq
    ..............................................augccgaucGaccacuccuuugu...................    1          1          seq
    ..............................................augccgauccaccacuccuuugA...................    6          1          seq
    ..............................................augccgauccaccacuccuuugu...................    36         0          seq
    ..............................................aCgccgauccaccacuccuuugu...................    1          1          seq
    ..............................................augccgauccaccacCccuuugu...................    1          1          seq
    ..............................................Gugccgauccaccacuccuuugu...................    2          1          seq
    ..............................................augccgauccaccacuccuuuguu..................    5          0          seq
    ..............................................augccgauccaAcacuccuuuguu..................    1          1          seq
```

miRBase precursor : novel-nve-miR-81
Total read count : 90
novel-nve-miR-81_guide read count : 90
novel-nve-miR-81_star read count : 0
remaining reads : 0

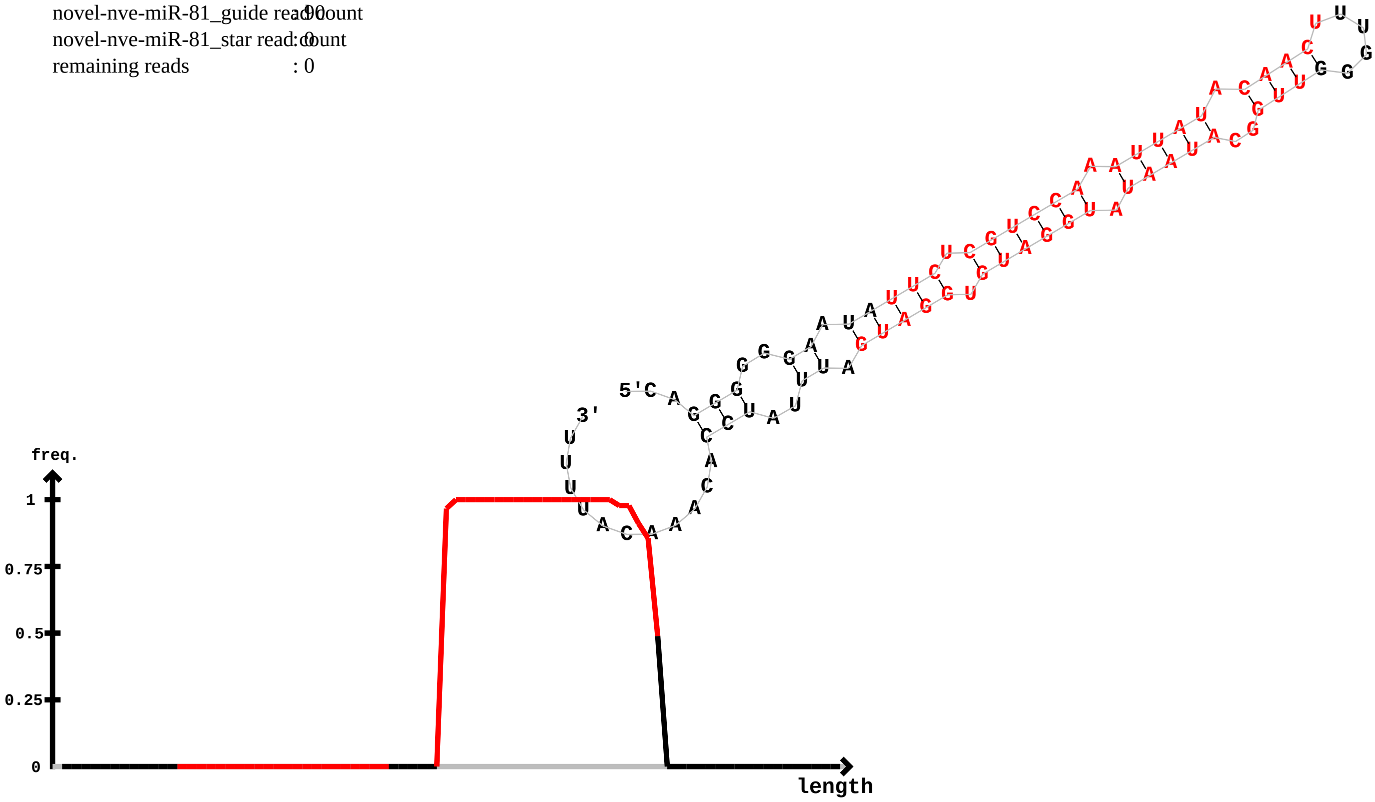


```
                 novel-nve-miR-81_star
                                          novel-nve-miR-81_guide
5'- caggggggaauauucucguccaaauuauacaacuuuggguuggcauaauauggauguggaugauuuauccacaaacauuuu   -3'    exp
    ..(((..((.(((((.((((((.(((((.((((.....))))..))))).)))))).))))).))..))).........   reads      mm         sample
    .......................................uuggUauaauauggaugu........................   1          1          seq
    .......................................uuggcauaauauggaugu........................   1          0          seq
    .......................................uuggcauaauauggaugugg......................   5          0          seq
    .......................................uuggcauaauauggaugugA......................   1          1          seq
    .......................................uuggcauUauauggaugugga.....................   3          1          seq
    .......................................uuggcauaauauggaugugga.....................   2          0          seq
    .......................................uuCgcauaauauggauguggau....................   1          1          seq
    .......................................uuAgcauaauauggauguggau....................   1          1          seq
    .......................................uuggcauaauauggauguggau....................   23         0          seq
    .......................................uuggcauaGuauggauguggau....................   1          1          seq
    .......................................uuggcauaauauggauguggaC....................   4          1          seq
    .......................................uuggcauaauauggauguggauC...................   10         1          seq
    .......................................uuggcauaauauggauguggauU...................   34         1          seq
    ........................................uggcauaauauggauAuggau....................   1          1          seq
    ........................................uggcauaauauggauguggau....................   2          0          seq
```

miRBase precursor : novel-nve-miR-82
Total read count : 226
novel-nve-miR-82_guide read count : 226
novel-nve-miR-82_star read count : 0
remaining reads : 0

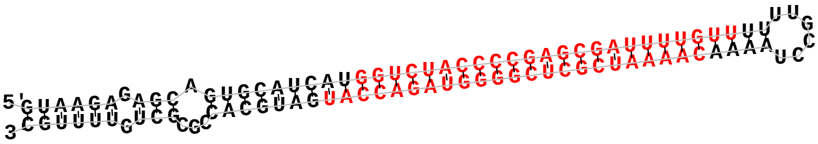

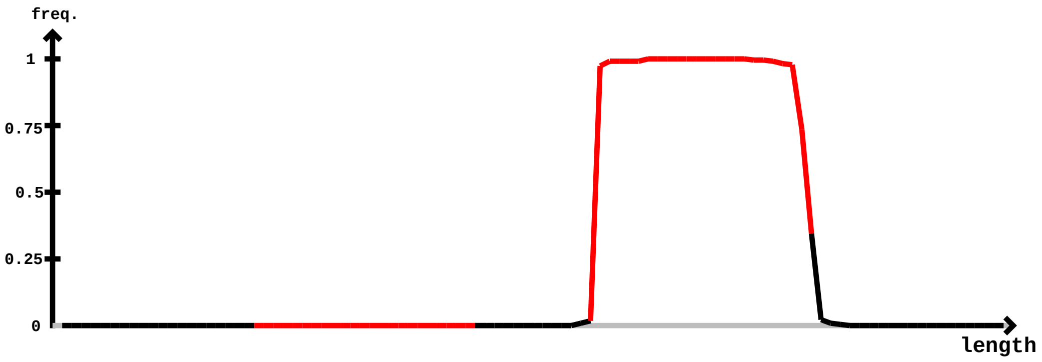


```
                                                                  novel-nve-miR-82_guide
                         novel-nve-miR-82_star
5'- guaagagagcagugcaucauggucuaccccgagcgauuuuguuuuuugccuaaaacaaaaucgcucgggguagaccaugaugcaccgcgcuguuuugc  -3'  exp
    ((((((.(((.((((((((((((((((((((((((((((((((((......)))))))))))))))))))))))))))))))))...))).))))))   reads  mm  sample
    .....................................................aacaaaaucgcucggggu...........................   1   0   seq
    .....................................................aacaaaaucgcucgggguag.........................   1   0   seq
    ......................................................acaaaaucgcucgggguagacU......................   1   1   seq
    ......................................................acaaaaucgcucgggguagacA......................   1   1   seq
    .......................................................caaaaucgcucgggguaga........................   2   0   seq
    .......................................................caaaaucgcucgggguagac.......................   1   0   seq
    .......................................................caaaaucgcuUgggguagacc......................   2   1   seq
    .......................................................caaaaucgcucgggguagacA......................   5   1   seq
    .......................................................caaaaucgcucgggguagacG......................   2   1   seq
    .......................................................caaaaucgcucggggguGgacc.....................   1   1   seq
    .......................................................caaaaucgcucgggguagacc......................   17  0   seq
    .......................................................caaaaucgcucgggguagacU......................   25  1   seq
    .......................................................caaaaucgcucgggguagacca.....................   69  0   seq
    .......................................................caaaaucgcucgggguagaccC.....................   2   1   seq
    .......................................................caaaaucgcucgggguagCcca.....................   1   1   seq
    .......................................................caaaaucgcucgggguagaccU.....................   4   1   seq
    .......................................................caaaaucUcucgggguagacca.....................   1   1   seq
    .......................................................caaaaucgcucgggguGgacca.....................   4   1   seq
    .......................................................Uaaaaucgcucgggguagacca.....................   1   1   seq
    .......................................................caaaaucgcucCggguagacca.....................   1   1   seq
    .......................................................caaaaucgcucgggguagaccG.....................   3   1   seq
    .......................................................caaaaucgcAcgggguagaccau....................   1   1   seq
    .......................................................caaaaucgcucgggguagaccaC....................   7   1   seq
    .......................................................caaaaucgcGcgggguagaccau....................   1   1   seq
    .......................................................caaaaucgcCcgggguagaccau....................   1   1   seq
    .......................................................caaaaucgcucgggguagGccau....................   1   1   seq
    .......................................................caaaaucgcucgggguagaccau....................   56  0   seq
    .......................................................caaaGucgcucgggguagaccau....................   2   1   seq
    .......................................................caaaaucgcucgggguGgaccau....................   3   1   seq
    .......................................................caaaaucgcucgggguagaccauU...................   2   1   seq
    .......................................................caaaaucgcucgggguagaccauC...................   1   1   seq
    ........................................................Caaaucgcucgggguagacc......................   1   1   seq
    ........................................................aaaaucgcucgggguagacca.....................   1   0   seq
    ........................................................Caaaucgcucgggguagacca.....................   1   1   seq
```

```
                                                       novel-nve-miR-82_guide
                    novel-nve-miR-82_star
guaagagagcagugcaucauggucuaccccgagcgauuuuguuuuuugccuaaaacaaaaucgcucgggguagaccaugaugcaccgcgcuguuuugc

........................................................aaaaucgcucgggguagaccau....................    1    0    seq
............................................................ucgcucgggguagaccauga..................    1    0    seq
............................................................ucgcucgggguagaccaugau.................    1    0    seq
```

miRBase precursor : novel-nve-miR-93
Total read count : 242
novel-nve-miR-93_guide read count : 242
novel-nve-miR-93_star read count : 0
remaining reads : 0

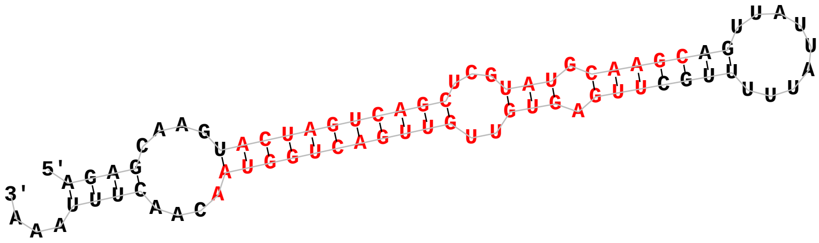

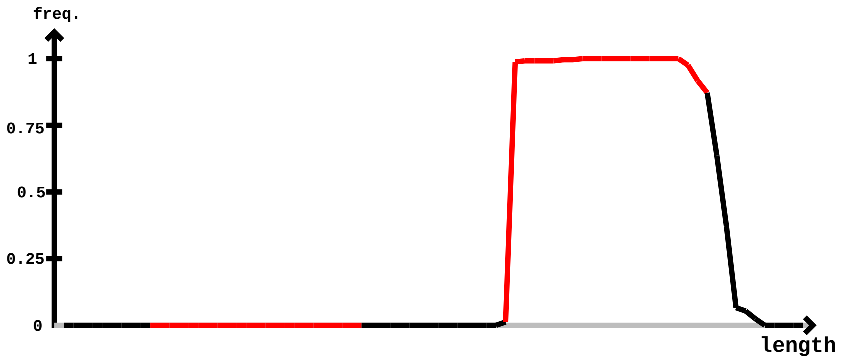


```
                                   novel-nve-miR-93_star
                                                                      novel-nve-miR-93_guide
5'- agagcaaguacuagucagcucguaugcaagcaguuauuauuuuugcuugaguguuguugacugguaacaacuuuaaa  -3'    exp
    ((((....((((((((((...(((.(((((((.........))))))).)))..))))))))))....))))...     reads     mm     sample
    .............................................cuugaguguuguugacugU.............     1         1      seq
    .............................................cuugaguguuguugacugguaa..........     1         0      seq
    .............................................cuugaguguuguugacugguaU..........     1         1      seq
    ..............................................uugaguguuguugacugg.............     2         0      seq
    ..............................................uugaguguuguugacugU.............     1         1      seq
    ..............................................uugaguguuguugacugA.............     2         1      seq
    ..............................................uugaguguuguugacuggC............     4         1      seq
    ..............................................uugaguguuguugacuggA............     1         1      seq
    ..............................................uugaguguuguugacuggu............     9         0      seq
    ..............................................uugaguguuguugacuggua...........     10        0      seq
    ..............................................uugaguguuguugacugUua...........     1         1      seq
    ..............................................uugaguguuguugacugguaU..........     5         1      seq
    ..............................................uugaguguuguuUacugguaa..........     1         1      seq
    ..............................................Gugaguguuguugacugguaa..........     1         1      seq
    ..............................................uugaguguuguugacugguaC..........     1         1      seq
    ..............................................uugGguguuguugacugguaa..........     1         1      seq
    ..............................................uugagugCuguugacugguaa..........     1         1      seq
    ..............................................uugaguguuguugacugguaa..........     40        0      seq
    ..............................................uugaguguuguugacugguaG..........     3         1      seq
    ..............................................uugaguguuguugacugCuaa..........     1         1      seq
    ..............................................uCgaguguuguugacugguaa..........     1         1      seq
    ..............................................uugaguguuAuugacugguaac.........     1         1      seq
    ..............................................uugaguguuguugacugguaaU.........     13        1      seq
    ..............................................uCgaguguuguugacugguaac.........     1         1      seq
    ..............................................uugaguguuguugacugguGac.........     1         1      seq
    ..............................................uugaguguuguugacugguaac.........     46        0      seq
    ..............................................uugaguguuguugacugguaUc.........     1         1      seq
    ..............................................uAgaguguuguugacugguaac.........     1         1      seq
    ..............................................uugaguguuguugacugguaacC........     9         1      seq
    ..............................................uugaguguuguugacugguaacG........     4         1      seq
    ..............................................uugaguguuguugacugguaaca........     27        0      seq
    ..............................................uugaguguuguugacugguaacU........     33        1      seq
    ..............................................uugaguguuguugacugguaacaG.......     1         1      seq
    ..............................................uugaguguuguugacugguaacaa.......     1         0      seq
```

```
         novel-nve-miR-93_star
                                                novel-nve-miR-93_guide
agagcaaguacuagucagcucguaugcaagcaguuauuauuuuugcuugaguguuguugacugguaacaacuuuaaa

.............................................uugaAuguuguugacugguaacaa.......    1          1          seq
.............................................uugaguguuguugacugguaacaaU......    5          1          seq
.............................................uugaguguuguugacugguaacaac......    2          0          seq
.............................................uugaguguuguugacugguaacaacA.....    1          1          seq
.............................................uugaguguuguugacugguaacaacu.....    3          0          seq
..............................................ugaguguuguugacugguaacaacu.....    1          0          seq
..................................................uguuguugacugguaaca........    1          0          seq
....................................................uuguugacugguaacaacu.....    1          0          seq
```

miRBase precursor : novel-nve-miR-95
Total read count : 85
novel-nve-miR-95_guide read count : 85
novel-nve-miR-95_star read count : 0
remaining reads : 0

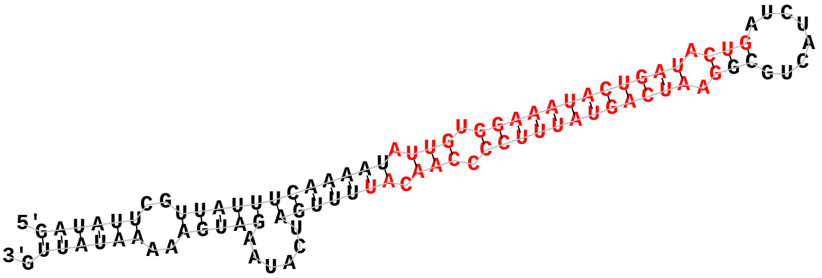

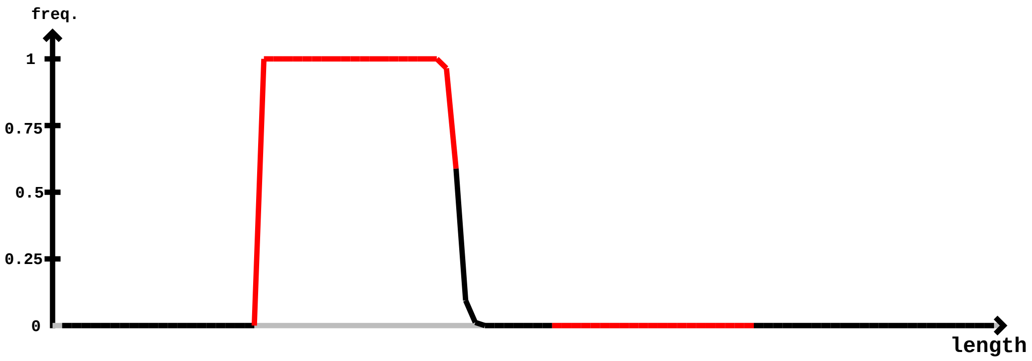


```
                                                                      novel-nve-miR-95_star
                           novel-nve-miR-95_guide
5'- gauauucguuauuucaaaauauuguggaaauacugauacugaucuacugcggaaucaguauuuccccaacauuuugaucauaagaugaaaaauauug   -3'   exp
    (((((..(((((((((((.(((.(((((((((((.(((........))).))))))))))).))).)))))))......)))))..)))))).   reads   mm   sample
    ....................auuguggaaauacugauaU..........................................................   1   1   seq
    ....................auuguggaaauacugauac..........................................................   2   0   seq
    ....................auuguggaaauacugauacu.........................................................   23   0   seq
    ....................auuguggaaauacugauacA.........................................................   3   1   seq
    ....................auuguggaaauacugauacC.........................................................   5   1   seq
    ....................Guuguggaaauacugauacu.........................................................   1   1   seq
    ....................auuguggaaauacugauacug........................................................   33   0   seq
    ....................auuguggGaauacugauacug........................................................   1   1   seq
    ....................auuguggaaauacugauacCg........................................................   1   1   seq
    ....................auuguggaaauacugauacuA........................................................   7   1   seq
    ....................auuguggaaauacugauacugC.......................................................   1   1   seq
    ....................auuguggaaauacugauacugU.......................................................   3   1   seq
    ....................auuguggaaauacugauacuga.......................................................   3   0   seq
    ....................auuguggaaauacugauacugau......................................................   1   0   seq
```

miRBase precursor : novel-nve-miR-97
Total read count : 42
novel-nve-miR-97_guide read count : 42
novel-nve-miR-97_star read count : 0
remaining reads : 0

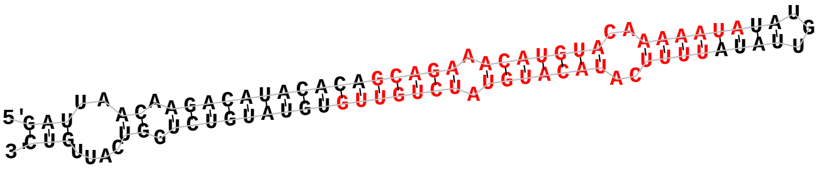

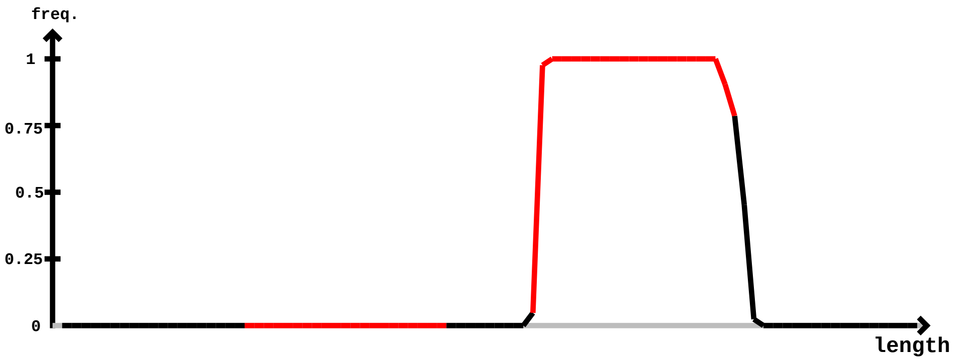


```
                                   novel-nve-miR-97_star
                                                          novel-nve-miR-97_guide
5'- gauuaacaagacauacacagcagaaacauguacaaaaauauauguuauauuucauacauguaucuguuguguaugucuggucauuguc   -3'    exp
    ((((....(((((((((((((((.(((((((..((((((((...))))))))..))))))).))))))))))))))))))).......    reads      mm         sample
    ................................................auuuucauacauguaucugC....................    1          1          seq
    ................................................auuuucauacauguaucuguu...................    1          0          seq
    .................................................uuuucauacauguaucugA....................    1          1          seq
    .................................................uuuucauacauguaucugu....................    1          0          seq
    .................................................uuuucauacauguaucugC....................    1          1          seq
    .................................................uuuuUauacauguaucuguu...................    1          1          seq
    .................................................uuuucauacauguaucuguu...................    3          0          seq
    .................................................uuuuUauacauguaucuguug..................    1          1          seq
    .................................................uuuucauacauguaucuguug..................    11         0          seq
    .................................................uuuucauacauguaucuguuU..................    2          1          seq
    .................................................uuuucauacauguaucuguugu.................    10         0          seq
    .................................................uuuucauacauguaucuguugA.................    2          1          seq
    .................................................uuuucauacauguaucuguuUu.................    1          1          seq
    .................................................uuuucauacauguaucuguugC.................    4          1          seq
    .................................................uuuucauacauguaucuguuguU................    1          1          seq
    ..................................................uuucauacauguaucuguugu.................    1          0          seq
```

| | | |
|---|---|---|
| Provisional ID | | novel-nve-miR-98 |
| Score total | : | 1.6 |
| Score for star read(s) | : | -1.3 |
| Score for read counts | : | 0 |
| Score for mfe | : | 1.9 |
| Score for randfold | : | 1.6 |
| Score for cons. seed | : | -0.6 |
| Total read count | : | 3226 |
| Mature read count | : | 3226 |
| Loop read count | : | 0 |
| Star read count | : | 0 |

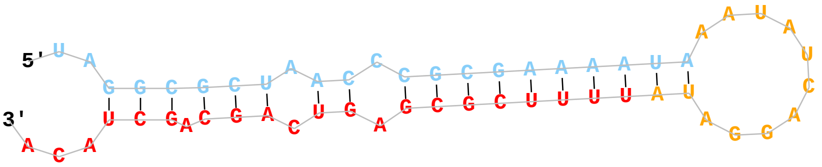


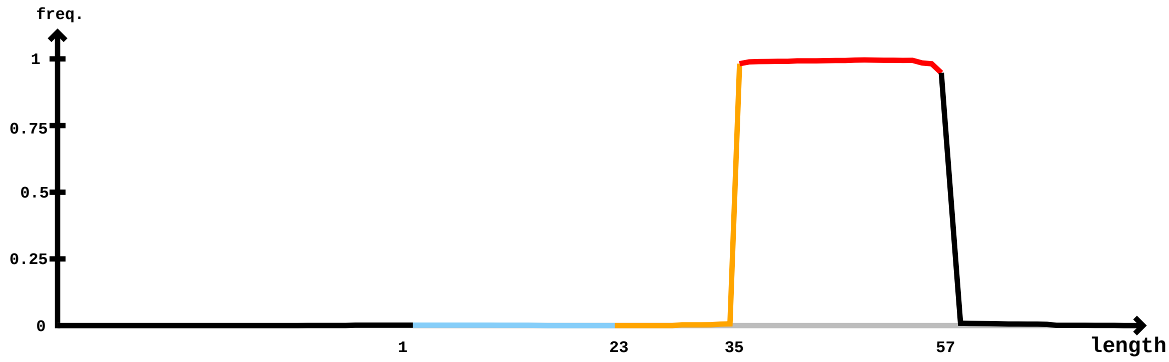


Star                                Mature

```
5'- gaagcacagauuugauuaccaacauagccuacguguaggcgcuaacccgcgaaaauaaauaucaggauauuuucgcgagucagcagcuacacguaggcuauuaccaacaca   -3'    exp
    .......................(((((((((((((((..(((.((.((((((((((..........)))))))))).)).)))..))))))))))))))).......... reads    mm    sample
    ........................uagccuacguguagCcgcua...................................................................   1       1     seq
    .............................uacguguaggcgcuaaccG...............................................................   2       1     seq
    .............................uacguguaggcgcuaacccA..............................................................   2       1     seq
    ...............................................................aggauauuuucgcgaguc..............................   4       0     seq
    ...............................................................aggauauuuucgcgagucag............................   1       0     seq
    ...............................................................aggauauuuucgcgagucaA.............................   1       1     seq
    ...............................................................aggauauuuucgcgagucaC.............................   1       1     seq
    ...............................................................aggauauuuucgcgagucagc...........................   1       0     seq
    ...............................................................aggauauuuucgcgagucagU...........................   1       1     seq
    ..................................................................auauuuucgcgagucagcagcu.......................   1       0     seq
    ...................................................................uauuuucgcgagucagcGg.........................   2       1     seq
    ...................................................................uauuuucgcgagucagcGgc........................   1       1     seq
    ...................................................................uauuuucgcgagucagcGgcu.......................   5       1     seq
    ....................................................................auuuucgcgagucagcagcuac.....................   1       0     seq
    ....................................................................auuuucgcgagucagcagcuacU....................   2       1     seq
    .....................................................................uuuucgcgagucagcagU........................   1       1     seq
    .....................................................................uuuucgcgagucagcCgcu.......................   1       1     seq
    .....................................................................uuuucgcgagucagcagcu.......................   15      0     seq
    .....................................................................uAuucgcgagucagcagcu.......................   1       1     seq
    .....................................................................uuuCcgcgagucagcagcu.......................   1       1     seq
    .....................................................................uuuucgcgagucagcagcA.......................   3       1     seq
    .....................................................................uuuucgcgagucGgcagcu.......................   1       1     seq
    .....................................................................uuuucgcgagucagcagcG.......................   4       1     seq
    .....................................................................uuuucgcgagucagcagcuU......................   1       1     seq
    .....................................................................uuuucgcgagucagcagcuC......................   1       1     seq
    .....................................................................uuuucgcgagucagcagcua......................   7       0     seq
    .....................................................................uuuucgcgagucagcagcGa......................   1       1     seq
    .....................................................................uuuucgcgagucagcGgcuac.....................   1       1     seq
    .....................................................................uuuucgcgagucagcagcuaA.....................   5       1     seq
    .....................................................................uuuucgcAagucagcagcuac.....................   1       1     seq
    .....................................................................uuuucgcgagCcagcagcuac.....................   1       1     seq
    .....................................................................uuuucgcgagucagcagcuaU.....................   44      1     seq
    .....................................................................uuuucgcgagucagcagcuac.....................   56      0     seq
    .....................................................................uuuucgcgagucagcagcuaG.....................   1       1     seq
```

```
                                   Star                              Mature
gaagcacagauuugauuaccaacauagccuacguguaggcgcuaacccgcgaaaauaaauaucaggauauuuucgcgagucagcagcuacacguaggcuauuaccaacaca
.....................................................................uuuucgcgagucagcagcuacG....................   257    1   seq
.....................................................................uuuucgcgagucagcagcuaca....................   200    0   seq
.....................................................................uuuucgcgagucagcGgcuaca....................   1      1   seq
.....................................................................uuuuGgcgagucagcagcuaca....................   1      1   seq
.....................................................................uuuucgcgaUucagcagcuaca....................   1      1   seq
.....................................................................Cuuucgcgagucagcagcuaca....................   1      1   seq
.....................................................................uuuucgcgagucagcaAcuaca....................   1      1   seq
.....................................................................uAuucgcgagucagcagcuaca....................   1      1   seq
.....................................................................uuuucgcgagucagcagcuGca....................   1      1   seq
.....................................................................uuuucgcgGgucagcagcuaca....................   4      1   seq
.....................................................................Guuucgcgagucagcagcuaca....................   1      1   seq
.....................................................................uuuucgcgagucagcagcuacU....................   2080   1   seq
.....................................................................uuuucgcgagucUgcagcuaca....................   1      1   seq
.....................................................................uuuucgcgagucagcagcuacC....................   478    1   seq
.....................................................................uuuuAgcgagucagcagcuaca....................   2      1   seq
.....................................................................uuuucgcgagucagcagcuacaU...................   1      1   seq
.....................................................................uuuucgcgagucagcagcuacUc...................   6      1   seq
......................................................................uuucgcgagucagcagcuac.....................   1      0   seq
......................................................................uuucgcgagucagcagcuacG....................   6      1   seq
......................................................................uuucgcgagucagcagcuacU....................   9      1   seq
......................................................................uuucgcgagucagcagcuacC....................   6      1   seq
.......................................................................uucgcgagucagcagcuacU....................   4      1   seq
........................................................................ucgcgagucagcaAcuacacguaggcu............   1      1   seq
.........................................................................cgcgagucagcagcuacacU..................   2      1   seq
...........................................................................cgagucagcaAcuacacgu.................   1      1   seq
...........................................................................cgagucagcagcuaUacgua................   1      1   seq
...........................................................................cgagucagcaAcuacacguag...............   2      1   seq
...........................................................................cgagucagcaAcuacacguagg..............   2      1   seq
..............................................................................gucagcaAcuacacguagg..............   1      1   seq
..............................................................................gucagcGgcuacacguaggcuauu.........   1      1   seq
...............................................................................ucagcGgcuacacguaggcuauu.........   2      1   seq
.................................................................................agcagcuacacguaggcuaC..........   2      1   seq
.................................................................................agcGgcuacacguaggcuauu.........   7      1   seq
..................................................................................gcGgcuacacguaggcuauu.........   2      1   seq
...................................................................................cUgcuacacguaggcuau..........   1      1   seq
.......................................................................................uacacguaggcuauuaccU.....   1      1   seq
.......................................................................................uacacguaggcuauuaccaaca..   1      0   seq
.......................................................................................uacacguaggcuauuaccaacU..   1      1   seq
```

miRBase precursor : novel-nve-miR-99-1
Total read count : 75
novel-nve-miR-99-1_guide read count : 75
novel-nve-miR-99-1_star read count : 0
remaining reads : 0

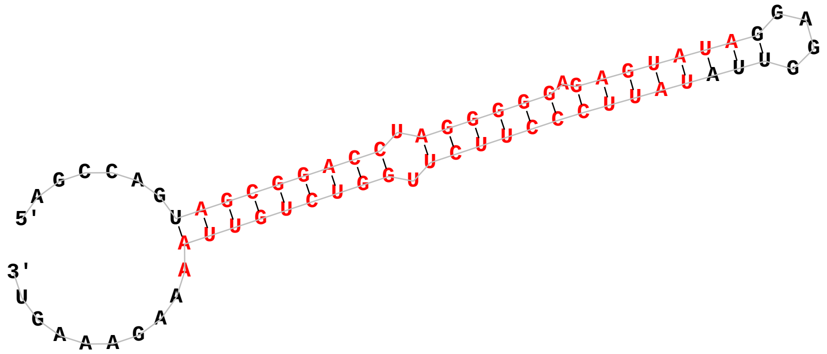

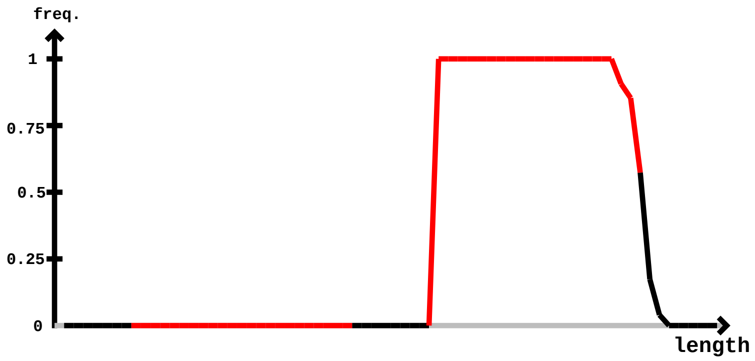


```
                                     novel-nve-miR-99-1_guide
         novel-nve-miR-99-1_star
5'- agccaguagcggaccuagggggagaguauaggagguuauauucccuucuuggucuguuaaaagaaagu   -3'    exp
    ......(((((((((.((((((.((((((((....)))))))))))))).)))))))))).........    reads      mm         sample
    ......................................uauucccuucuuggucugC...........    2          1          seq
    ......................................uauuUccuucuuggucugu...........    1          1          seq
    ......................................uauucccuucuuggucugu...........    4          0          seq
    ......................................uauucccuucuuggucuguu..........    4          0          seq
    ......................................uauucccuucCuggucuguua.........    1          1          seq
    ......................................uauucccuucuuggucuguuC.........    3          1          seq
    ......................................uauucccuucuuggucuguua.........    7          0          seq
    ......................................uauucccuucuuggucuguuU.........    9          1          seq
    ......................................uCuucccuucuuggucuguua.........    1          1          seq
    ......................................uauucccuucuuggucuguuaU........    6          1          seq
    ......................................uauucccuucuuggCcuguuaa........    1          1          seq
    ......................................uauucccuucuuggucuguuUa........    1          1          seq
    ......................................uauucccuucuuggucuguuaa........    22         0          seq
    ......................................uauucccuucuuggucuguuaaU.......    4          1          seq
    ......................................uauucccuucuuggucuguuaaa.......    3          0          seq
    ......................................uauucccuucuuggucuguuaaC.......    1          1          seq
    ......................................uauucccuucuuggucuguuaUa.......    1          1          seq
    ......................................uaGucccuucuuggucuguuaaa.......    1          1          seq
    ......................................uauucccuucuuggucuguuaaaU......    3          1          seq
```

miRBase precursor : novel-nve-miR-100-1
Total read count : 141
novel-nve-miR-100-1_guide read count : 141
novel-nve-miR-100-1_star read count : 0
remaining reads : 0

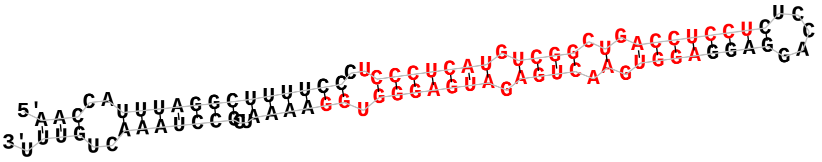

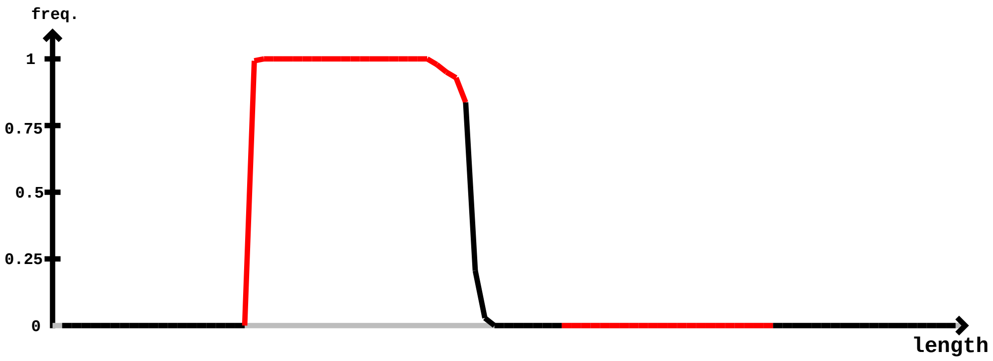


novel-nve-miR-100-1_guide

novel-nve-miR-100-1_star

```
5'- aaccauuuaggcuuuucccucccucaugucggcugaccuccucuccaggaggaggugaacugagaugagggugg aaaaucgccuaaacuguuu   -3'    exp
    (((..(((((((((((((..(((((((.((((.(.((((((((.....)))))))).).)))).))))))).))))))..)))))))..))).    reads      mm         sample
    ...................ucccucaugucggcugacA.......................................................    1          1          seq
    ...................ucccucaugucggcugacU.......................................................    1          1          seq
    ...................ucccucaugucggcugacc.......................................................    1          0          seq
    ...................ucccucaugucggcugaccu......................................................    3          0          seq
    ...................ucccucaugucggcugaAcu......................................................    1          1          seq
    ...................ucccucaugucggcugaccuc.....................................................    2          0          seq
    ...................ucccucaugucggcugaccuU.....................................................    1          1          seq
    ...................ucccucaugucggcugaccucc....................................................    7          0          seq
    ...................ucccucaugucggcugaccucA....................................................    2          1          seq
    ...................ucccucaugucggcugCccucc....................................................    1          1          seq
    ...................ucccucaugucggcugaccucG....................................................    1          1          seq
    ...................ucccucaugucggcugaccucU....................................................    2          1          seq
    ...................ucccucaugucAgcugaccuccu...................................................    2          1          seq
    ...................ucccucaugucggcugaccuccA...................................................    6          1          seq
    ...................ucccucaugucggcugaccuccG...................................................    5          1          seq
    ...................ucccucaugucggcugaccuccu...................................................    59         0          seq
    ...................ucccucaugucggcugaccuGcu...................................................    1          1          seq
    ...................ucccucaugucggcugaccuccC...................................................    15         1          seq
    ...................ucccucaugucggcugCccuccuc..................................................    1          1          seq
    ...................ucccucaugucAgcugaccuccuc..................................................    1          1          seq
    ...................ucccucaugucggcugaccuccuc..................................................    8          0          seq
    ...................ucccucaugucggcugaccuccuU..................................................    15         1          seq
    ...................ucccucaugucggcugaccuccuUu.................................................    4          1          seq
    ....................Uccucaugucggcugaccuccu...................................................    1          1          seq
```

miRBase precursor : novel-nve-miR-101-1
Total read count : 86
novel-nve-miR-101-1_guide read count : 86
novel-nve-miR-101-1_star read count : 0
remaining reads : 0

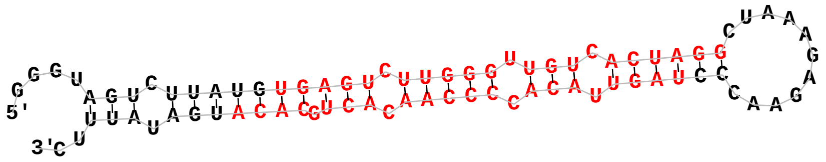

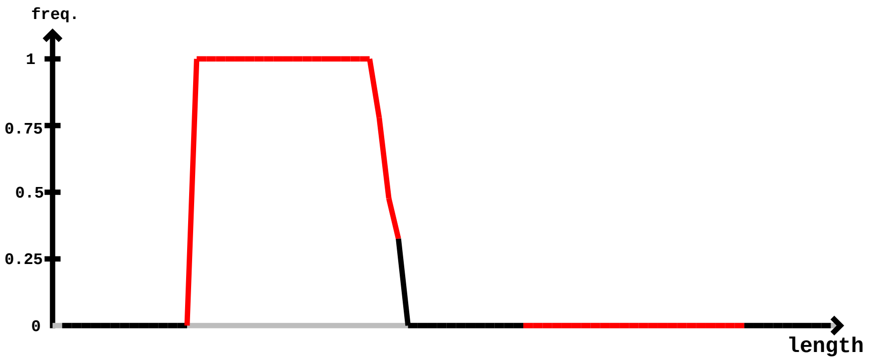


novel-nve-miR-101-1_star
novel-nve-miR-101-1_guide

```
5'- ggguagucuuaugugagucuuggguugucacuaggcuaaagagaacccuaguuacaccccaacacugcacaugauauuuc  -3'    exp
    ........((((((((((.(((((.(((.((((((...........)))))).))).))))).))).)))))))......     reads      mm         sample
    ............ugagucuuggguugucacC.............................................     1          1          seq
    ............ugagucuuggguugucacA.............................................     2          1          seq
    ............ugagucuAggguugucacu.............................................     1          1          seq
    ............ugagucuuggguugucacG.............................................     1          1          seq
    ............ugagucuuggguugucacu.............................................     14         0          seq
    ............Cgagucuuggguugucacua............................................     1          1          seq
    ............ugagucuuggguuAucacua............................................     1          1          seq
    ............ugagucuuggguugucacuC............................................     1          1          seq
    ............ugagucuuggguugucacua............................................     20         0          seq
    ............ugagucuuggguugucacuU............................................     1          1          seq
    ............ugagucuuggguCgucacua............................................     1          1          seq
    ............ugagucuuggguAgucacua............................................     1          1          seq
    ............ugagucuuggguugucacuaU...........................................     2          1          seq
    ............ugagucuuggguugucacuag...........................................     11         0          seq
    ............ugagucuuggguugucacuagA..........................................     2          1          seq
    ............ugagucuCggguugucacuagg..........................................     1          1          seq
    ............ugagucuuggguugucacuGgg..........................................     1          1          seq
    ............ugagucuuggguugucacuagg..........................................     23         0          seq
    ............ugagucuuggguugucacuagU..........................................     1          1          seq
```
