## Supplementary table 7 for "Unravelling the developmental and functional significance of an ancient Argonaute duplication"

| **AGO1 miRNAs** | **mRNA_targets** | **RE_targets** | **miR_homology_to_target** |
| --- | --- | --- | --- |
| missing: 9446 | 13 | 199 | NO |
| nve-miR-2022-3p | 10 | 4 | NO |
| novel-nve-miR-23 | 10 | 0 | NO |
| novel-nve-miR-103-1 | 8 | 0 | NO |
| novel-nve-miR-47-2 | 7 | 47 | NO |
| nve-miR-9424a | 6 | 0 | YES:NVE3874 |
| nve-miR-9422 | 6 | 2 | YES:NVE7449 |
| nve-miR-2030-5p | 6 | 0 | NO |
| novel-nve-miR-6 | 6 | 1 | NO |
| nve-miR-9413 | 5 | 15 | NO |
| nve-miR-2026-5p | 5 | 0 | NO |
| novel-nve-miR-66 | 5 | 41 | YES:Sattelite: SAT-1_NV |
| novel-nve-miR-34 | 4 | 0 | NO |
| novel-nve-miR-73 | 4 | 0 | YES:NVE23031 |
| novel-nve-miR-78 | 4 | 0 | NO |
| novel-nve-miR-9 | 4 | 0 | NO |
| novel-nve-miR-48-1 | 4 | 9 | NO |
| novel-nve-miR-50 | 4 | 9 | NO |
| novel-nve-miR-79 | 4 | 3 | YES:NVE17255 |
| missing: 48-2 | 4 | 9 | NO |
| nve-miR-9435 | 3 | 1 | NO |
| nve-miR-2025-3p | 3 | 1 | NO |
| novel-nve-miR-28 | 3 | 202 | NO |
| novel-nve-miR-58 | 3 | 17 | YES:NVE20313 |
| nve-miR-2037-3p | 2 | 1 | NO |
| nve-miR-2023-3p | 1 | 1 | NO |
| nve-miR-2040b-3p | 1 | 0 | NO |
| nve-miR-9425 | 1 | 0 | YES:NVE9231 |
| nve-miR-9455 | 1 | 0 | NO |
| novel-nve-miR-18 | 1 | 0 | NO |
| novel-nve-miR-74 | 1 | 1 | NO |
| novel-nve-miR-71 | 1 | 1 | YES:NVE12137 |
| missing: 2040a | 1 | 1 | NO |
| nve-miR-2024g-5p | 1 | 4 | NO |
| nve-miR-100-5p | 0 | 0 | NO |
| novel-nve-miR-100-1 | 0 | 0 | NO |
| novel-nve-miR-69 | 0 | 0 | NO |
| novel-nve-miR-8 | 0 | 0 | NO |
| novel-nve-miR-80 | 0 | 19 | NO |
| novel-nve-miR-36-2 | 0 | 0 | NO |

| **AGO2 miRNAs** | **mRNA targets** | **RE targets** | **miR_homology_to_target** |
| --- | --- | --- | --- |
| nve-miR-9445 | 37 | 0 | YES:NVE19200 |
| novel-nve-miR-89 | 14 | 7 | NO |
| novel-nve-miR-32 | 14 | 33 | **NO** |
| nve-miR-9446 | 13 | 199 | NO |
| novel-nve-miR-98 | 9 | 2 | YES:NVE1786 |
| nve-miR-2024f-3p | 8 | 1 | YES:NVE21566 |
| novel-nve-miR-96 | 6 | 23 | NO |
| novel-nve-miR-49 | 6 | 29 | NO |
| novel-nve-miR-26 | 6 | 4 | YES:NVE7145 |
| nve-miR-9456 | 5 | 1 | YES:NVE19966 |
| novel-nve-miR-5 | 5 | 11 | NO |
| novel-nve-miR-46 | 5 | 75 | YES:NVE20003 |
| novel-nve-miR-42-b | 5 | 3 | NO |
| novel-nve-miR-42-a | 5 | 1 | NO |
| novel-nve-miR-33 | 5 | 3 | NO |
| novel-nve-miR-20 | 5 | 203 | NO |
| nve-miR-9414-5p | 4 | 87 | YES:NVE17823 |
| nve-miR-2042-3p | 4 | 27 | NO |
| nve-miR-2041a-3p | 4 | 0 | YES:NVE1344 |
| nve-miR-2024e-3p | 4 | 1 | NO |
| novel-nve-miR-93 | 4 | 3 | YES:NVE7053 |
| novel-nve-miR-75 | 4 | 25 | NO |
| novel-nve-miR-57 | 4 | 8 | YES:NVE26001 |
| novel-nve-miR-44-1 | 4 | 5 | NO |
| nve-miR-9433 | 3 | 22 | NO |
| nve-miR-2024a-3p | 3 | 1 | -- |
| novel-nve-miR-39-1 | 3 | 0 | YES:NVE17602 |
| novel-nve-miR-35 | 3 | 4 | YES:NVE10979 OR RE: scaffold_3:2562609-2562933 |
| novel-nve-miR-19 | 3 | 175 | NO |
| novel-nve-miR-10 | 3 | 9 | NO |
| nve-miR-9431 | 2 | 5 | YES:RE: scaffold_73:490409-493015 |
| nve-miR-2045-5p | 2 | 1 | NO |
| nve-miR-2041b-3p | 2 | 3 | NO |
| nve-miR-2036-3p | 2 | 0 | NO |
| nve-miR-2035-5p | 2 | 0 | YES:NVE13698 |
| nve-miR-2027-5p | 2 | 16 | NO |
| novel-nve-miR-55 | 2 | 0 | YES:NVE12804 |
| novel-nve-miR-53 | 2 | 0 | NO |
| novel-nve-miR-38 | 2 | 0 | NO |
| nve-miR-9428 | 2 | 64 | NO |
| nve-miR-2047-3p | 1 | 0 | NO |
| nve-miR-2046-5p | 1 | 1 | NO |
| nve-miR-2039-3p | 1 | 5 | NO |
| nve-miR-2029-3p | 1 | 1 | YES:Kolobok-N5_NV scaffold_319:50509-50950 |
| nve-miR-2028-5p | 1 | 0 | NO |
| novel-nve-miR-85 | 1 | 4 | YES:NVE13163 |
| novel-nve-miR-83 | 1 | 0 | YES:NVE1474 |
| novel-nve-miR-62 | 1 | 3 | NO |
| novel-nve-miR-59 | 1 | 1 | NO |
| novel-nve-miR-54 | 1 | 1 | NO |
| novel-nve-miR-16 | 1 | 25 | NO |
| novel-nve-miR-11 | 1 | 0 | **NO** |
| nve-miR-2040a | 1 | 1 | NO |
| nve-miR-9448 | 0 | 0 | NO |
| nve-miR-2032a-3p | 0 | 3 | NO |
| nve-miR-2031-5p | 0 | 1 | NO |
| novel-nve-miR-82 | 0 | 1 | YES:RE: scaffold_130:31251-31483 |
| novel-nve-miR-56 | 0 | 0 | NO |
| novel-nve-miR-37 | 0 | 1 | NO |
| nve-miR-2032b | 0 | 0 | NO |
